# Exploiting Substrate Specificities of 6-*O*-Sulfotransferases to Enzymatically Synthesize Keratan Sulfate Oligosaccharides

**DOI:** 10.1101/2023.08.20.554027

**Authors:** Yunfei Wu, Gael Vos, Chin Huang, Digantkumar Chapla, Anne L.M. Kimpel, Kelley W. Moremen, Robert P. de Vries, Geert-Jan Boons

## Abstract

Keratan sulfate (KS) is a glycosaminoglycan that is widely expressed in the extracellular matrix of various tissue types where it is involved in many biological processes. Herein, we describe a chemo-enzymatic approach to prepare well-defined KS oligosaccharides by exploiting known and newly discovered substrate specificities of relevant sulfotransferases. The premise of the approach is that recombinant GlcNAc-6-O-sulfotransferases (CHST2) only sulfates terminal GlcNAc moieties to give GlcNAc6S that can be galactosylated by B4GalT4. Furthermore, CHST1 can modify internal galactosides of a poly-LacNAc chain, however, it was found that a GlcNAc6S residue greatly increases the reactivity of CHST1 of a neighboring and internal galactoside. The presence of a 2,3-linked sialoside further modulates the site of modification by CHST1, and a galactoside flanked by 2,3-Neu5Ac and GlcNAc6S is preferentially sulfated over other Gal residues. The substrate specificities of CHST1 and 2 were exploited to prepare a panel of KS oligosaccharides including selectively sulfated *N*-glycans. The compounds and several other reference derivatives were used to construct a microarray that was probed for binding by several plant lectins, Siglec proteins and hemagglutinins of influenza viruses. It was found that not only the sulfation pattern but also presentation of epitopes as part of an *O*- or *N*-glycan determines binding properties.

## INTRODUCTION

Keratan sulfates (KS) are *N*- and *O*-linked glycans that occur in the extracellular matrix of many tissue types where they can interact with a multitude of proteins thereby controlling physiological and disease processes.^1-4^ One of the antennae of these *N*- and *O*-glycans is composed of poly-*N*-acetyl-lactosamine (poly-LacNAc) that can be modified by sulfation at C-6 positions of *N*-acetyl-glucosamine (GlcNAc) and galactose (Gal) (Fig. 1). Certain classes of KS can be further modified by 1,3-linked fucosides. KS has a modular architecture, and its backbone is composed of differently sulfated and fucosylated LacNAc moieties and the termini can additionally be modified by 2,3- and 2,6-linked sialic acids.

**Figure 1.**
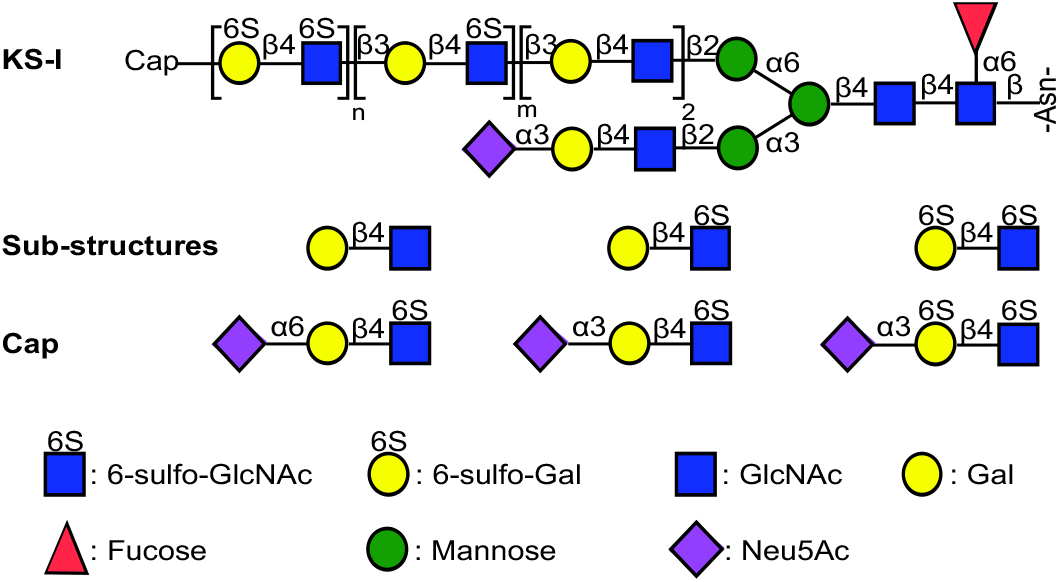
Structures of keratan sulfate oligosaccharides including sub-structures of repeating LacNAc chain and structures of capping epitopes.

The sulfated poly-LacNAc moieties of KS are assembled by a collection of glycosyltransferases and sulfotransferases.^5^ The consecutive action of β(1,3)-N-acetylglucosaminyltransferases (B3GnT) and β(1,4)-galactosyl transferases (B4GalT) results in the formation of the LacNAc backbone of KS. During the assembly of this chain, C-6 hydroxyls of terminal GlcNAc moieties can be sulfated by GlcNAc-6-O-sulfotransferases 2 and 6 (CHST2 and 6).^6-11^ The enzyme B4GalT4 can attach a 1,4-linked galactoside to 6-sulfo-GlcNAc residues whereas B4GalT1 and B4GalT7 can extend unmodified GlcNAc residues. After assembly of the LacNAc chain, C-6 hydroxyls of galactosides can be sulfated by keratan sulfate galactose 6-sulfotransferase (KSGal6ST or CHST1) or chondroitin sulfotransferase-1 (C6ST1).^12,13^ β-Galactoside α-2,3-sialyltransferase 4 (ST3Gal4) and β-galactoside-α-2,6-sialyltransferase 1 (ST6Gal1) can install terminal 2,3- and 2,6-linked sialosides. Although not fully understood, the KS biosynthetic enzymes cooperate to create specific epitopes that can recruit KS-binding proteins. A lack of well-defined KS structures has made it difficult to uncover the molecular basis by which it regulates biological processes.

Structural heterogeneity makes it challenging to obtain well-defined KS oligosaccharides from natural sources. Chemical approaches have been reported for the preparation of sulfated LacNAc derivatives, however, due to the need to perform time consuming and demanding protecting manipulations and glycosylations, it has resulted only in relatively small structural motifs such as di- and tetrasaccharides.^14-18^ Sulfated LacNAc derivatives have been chemically synthesized that were further modified by fucosylation and sialylation.^19^ In an interesting approach, chemically synthesized oxazolines were linked together by *trans*-glycosylation using a mutant form of keratanase II from *Bacillus* to give several oligosaccharides. The scope of this approach is restricted, however, due to a limited substrate tolerance and cannot provide compounds larger than a hexasaccharide.^20,21^

Herein, we describe a chemo-enzymatic approach to prepare well-defined KS oligosaccharides by exploiting known and newly discovered substrate specificities of relevant sulfotransferases. The premise of the approach is that recombinant CHST2^22^ only sulfates terminal GlcNAc moieties to give GlcNAc6S that can be galactosylated by B4GalT4 to provide βGal(1,4)GlcNAc6S. Furthermore, CHST1 can modify internal galactosides of a poly-LacNAc chain, however, it was found that a GlcNAc6S residue greatly increases the reactivity of neighboring and internal galactoside. The presence of a 2,3-linked sialoside further modulates the site of modification by CHST1, and a galactoside that is flanked by 2,3-Neu5Ac and GlcNAc6S is preferentially sulfated over other galactosyl residues. The substrate specificities of CHST1 and 2 were exploited for the preparation of a panel of KS oligosaccharides including selectively sulfated *N*-glycans. The resulting and several other reference compounds were used to construct a microarray that was probed with lectins, glycan binding proteins and hemagglutinins of influenza A viruses. It was found that sulfation can modulate recognition and can either be tolerated, enhance, or reduce binding. Furthermore, we discovered that the presentation of sulfated epitopes in the context of *N*- and *O*-glycan can greatly impact binding.

LacNAc derivatives **4**-**6** were prepared to examine the substrate specificities of CHST1 (Scheme 1a). In addition, compounds **11, 8**, and **10** were used to examine the influence of GlcNAc6S on sialylation, Finally, compounds **10, 20**, and **15** were employed to examine in which way sulfation at the C-6 position of GlcNAc and 2,3-sialylation influences the regioselectivity of CHST1 (Schemes 2b and 2c). We exploited the ecto-domains of recombinantly B4GalT1 and B3GnT2^22^ in combination with UDP-Gal and UDP-GlcNAc to assemble the unmodified oligo-LacNAc moieties **4**-**6** (Scheme 1a) starting from chemically synthesized LacNAc derivative **1**. The latter compound has a benzyloxycarbonyl (Cbz) protected aminopentyl spacer at the anomeric center, which after deprotection, facilitates glycan microarray construction. Thus, treatment of **1** with B3GnT2 and UDP-GlcNAc gave **2** which was further treated with B4GalT1 in the presence of UDP-Gal to provide **3**. Another cycle of enzymatic modification by B3GnT2 an B4GalT1 gave access to **5** and **6**. The compounds were purified by size exclusion column chromatography over Bio-Gel P2 or P6 and fully characterized by homo- and heteronuclear two-dimensional NMR experiments and by LC-MS.

Next, attention was focused on sulfation and further elongation of compounds **2** and **4**. Treatment of these compounds with CHST2 in the presence of PAPS resulted in the facile formation of **7** and **9**, respectively. Detailed NMR analysis confirmed that sulfation had occurred at the terminal GlcNAc moiety. For example, 1D ^1^H NMR and 2D ^13^C–^1^H HSQC spectra of compound **7** (see SI) made it possible to assign all proton and carbon signals. The 6-carbon of the terminal GlcNAc moiety had shifted downfield (δ 60.2 → δ 67.2) and the corresponding protons also exhibited a chemical shift difference (H6a δ 3.98 → 4.34, H6b 3.83 → 4.23), which confirmed the regioselectivity of sulfation. The inter-residue connectivity was confirmed by a NOESY spectrum, which showed interactions of H-1 of GlcNAc-C with H-3 Gal-B, and H-1 of Gal-B with H-4 of GlcNAc-A in accordance with C(1→3)B, B(1→4)A linkages, respectively.

Surprisingly, during the preparation of **7** a trace amount of di-sulfated product was detected by LC-MS, which could readily be removed by diethylaminoethyl (DEAE) ion exchange column chromatography. The main product was, however, terminally sulfated confirming the substrate specificity of CHST2 (Scheme S1 in SI). Compounds **7** and **9** could readily be galactosylated using B4GalT4 in the presence of UDP-Gal to provide compounds **8** and **10**, respectively. Other galactosyl transferases were also examined and for example B4GalT1 was able to modify **7** and **9** but the rate of transfer was very slow, and the reaction could not be driven to completion. HpGalT^23^ was able to fully modify **7** and **9** but the rate of transfer was substantially slower than for B4GalT4. The presence of a sulfate at the terminal LacNAc moieties of compounds **11, 8**, and **10** did not impede sialylation by ST3Gal4 and ST6Gal1 and compounds **12**-**15** could readily be prepared.

**Scheme 1.**
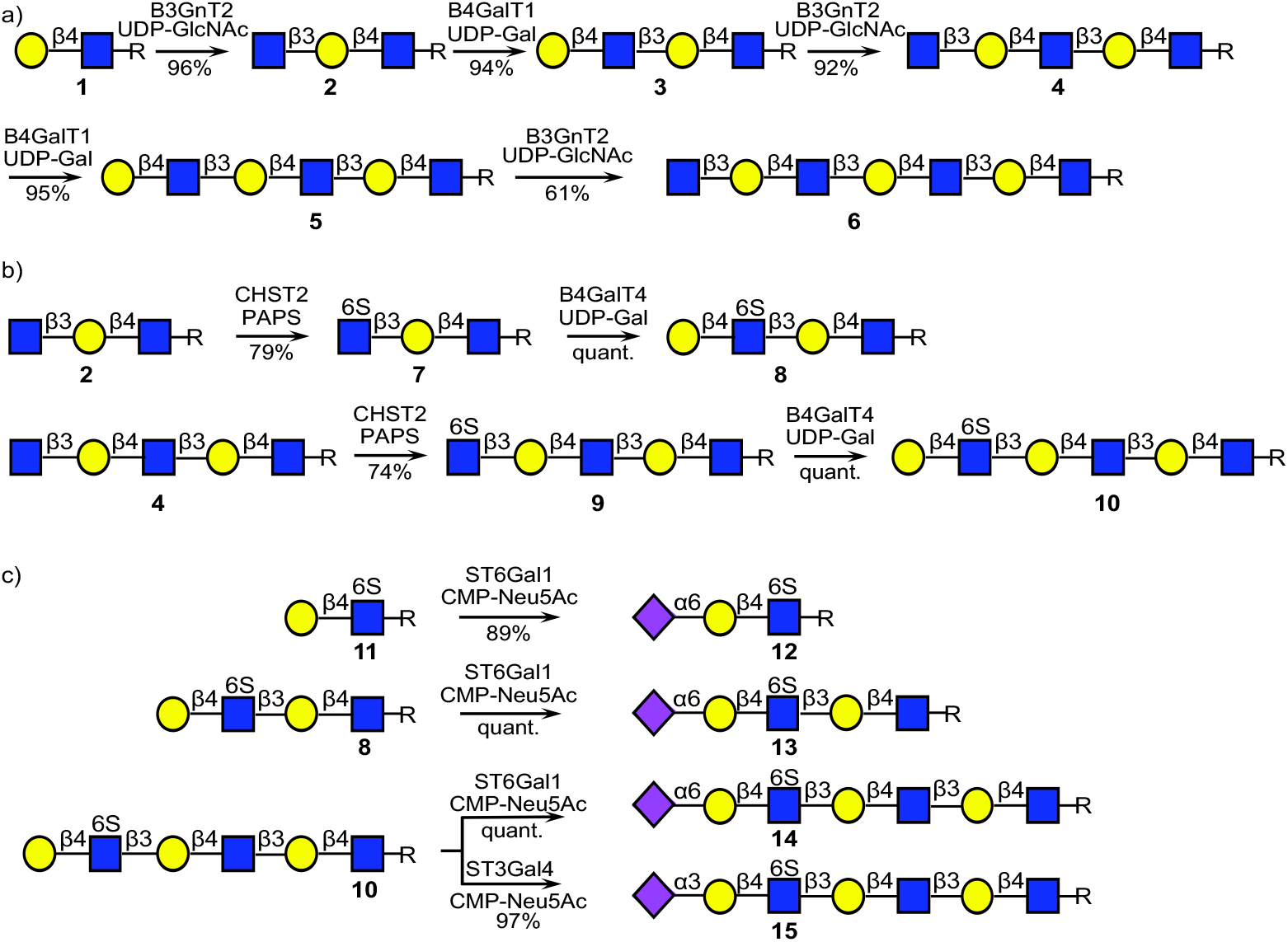
Enzymatic synthesis of linear keratan sulfate oligosaccharides. a) assembly of polyLacNAc backbone. b) elongation of GlcNAcS6. c) cap introduction of keratan sulfate. R= O(CH_2_)_5_NHCbz

Compounds **4**-**6** and **10, 20, 15** were used to explore the substrate specificities of CHST1 (Scheme 2). First, the linear LacNAc substrates **4, 5** and **6** were treated with CHST1 in the presence of PAPS (1.5 eq. per galactoside) and the progress of the reactions were monitored by LC-MS. Although the reactions were slow, all galactosyl moieties of compounds **4** and **6** were sulfated at C-6 to provide, after purification by size exclusion column (P6) and ion exclusion column (DEAE) chromatography, the sulfates **16** and **18**, respectively. On the other hand, hexasaccharide **5** was only sulfated at the internal galactosides to give di-sulfate **17**.

To examine influence of 6-*O*-sulfation of GlcNAc, compound **10** was incubated with CHST1 in the presence of PAPS, which resulted in the formation of a single compound (Scheme 2b). Detailed NMR analysis demonstrated sulfation had occurred at the -1 site to give compound **19**. For example, 1D ^1^H NMR and 2D ^13^C–^1^H HSQC spectra of compound **19** (SI) made it possible to assign all proton and carbon signals. The 6-carbon of the internal sulfated galactose moiety had substantially shifted downfield (δ 61.3 → δ 67.9) and the corresponding protons also exhibited a chemical shift difference (H6 δ 3.76 → 4.20), which confirmed the regioselectivity of sulfation. The inter-residue connectivity was confirmed by a NOESY spectrum, the inter-residue connectivities Gal-F H-1, GlcNAc-E H-4, GlcNAc-E H-1, Gal-D H-3, Gal-D H-1, GlcNAc-C H-4, GlcNAc-C H-1, Gal-B H-3 and Gal-B H-1, GlcNAc-A H-4 are in accordance with F(1→4)E, E(1→3)D, D(1→4)C, C(1→3)B, B(1→4)A linkages, respectively. Surprisingly, sialoside **20** was not sulfated by CHST1 indicating that sialic acid deactivates all the galactosides from modification (Scheme 2b). It implies that CHST1 senses the modifications at the non-reducing terminus of the polymer impacting modification of internal sites. A combination of GlcNAc6S and a 2,3-linked sialoside at a terminal LacNAc moiety, as in compound **15** (Scheme 2c), changed the site of sulfation and in this case treatment with CHST1 in the presence of PAPS (1.6 eq.) resulted in the formation of **21** as the major product which has a sulfate at the Gal moiety that is flanked by the sialoside and GlcNAc6S. A small amount of additionally sulfated **22** was formed, which could easily be removed by DEAE ion exchange column chromatography. Further incubation of **21** with further PAPS for a prolonged period of time resulted in further sulfation of the Gal moiety at the - 1 site providing compound **22**. These results highlight a complex interplay between sialylation and sulfation to introduce terminal epitopes and only the presence of a 2,3-linked sialoside and GlcNAc6S activates the C-6 of the galactose of the terminal LacNAc moiety for sulfation by CHST1. The latter sulfotransferase can also modify the galactoside at -1 site albeit at a lower rate of modification.

**Scheme 2.**
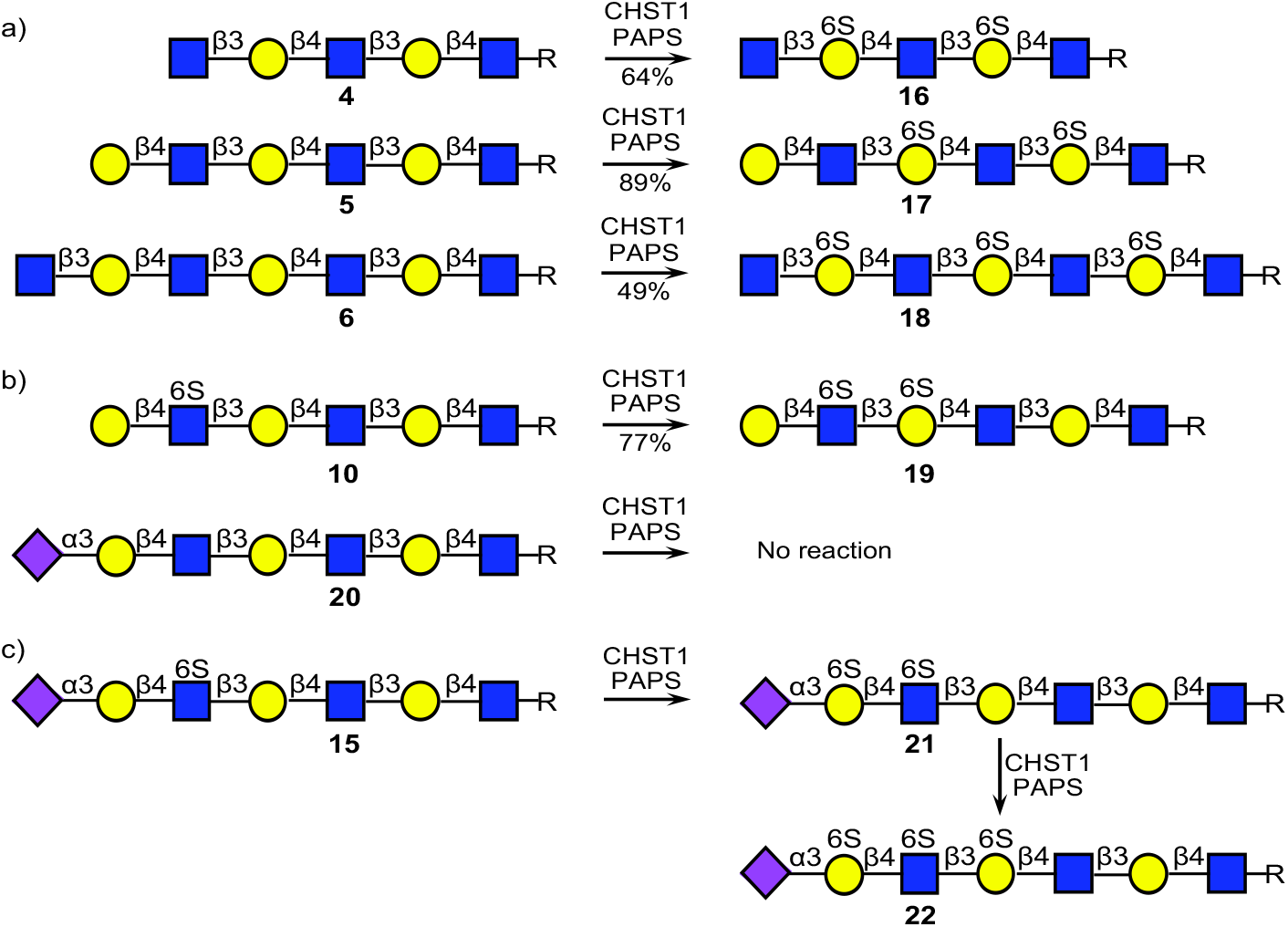
Using well-defined oligosaccharides to examine the substrate specificity of CHST1. R= O(CH_2_)_5_NHCbz

The newly discovered selectivities of CHST1 and CHST2 were exploited to enzymatically prepare several *N*-linked glycans. Sialoglycopeptide (SGP, Scheme 3), which can be isolated in multi-gram quantities from egg yolk powder, was converted into bi-antennary glycosyl asparagine **23** by subsequent pronase treatment to remove the peptide moiety and hydrolysis of the sialosides with 2 M AcOH.^24,25^ Next, ST6Gal1, which has a preference for the MGAT1 arm, was used to desymmetrize **23** to give mono-sialoside **24**.^26^ The terminal galactoside of **24** was extended by subsequent modification by B3GNT2 (→**25**), B4GalT1 (→ **26**) and B3GNT2 to provide **27**. As expected, the terminal GlcNAc moiety of **27** could selectively be sulfated by CHST2 in the presence of PAPS to give **28**, which was further treated with B4GalT4 and UDP-Gal to provide compound **29**. As expected, the terminal Gal of **29** could readily be sialylated by ST6Gal1 or ST3Gal4 resulting in the formation of compounds **30** and **31**, respectively. Finally, **31** was subjected to CHST1 and PAPS (1.5 eq.), which gave **32** as the major compound and a small amount of **33**. The two compounds could readily be separated by DEAE ion exchange column chromatography and were fully characterized by NMR and LC-MS.

**Scheme 3.**
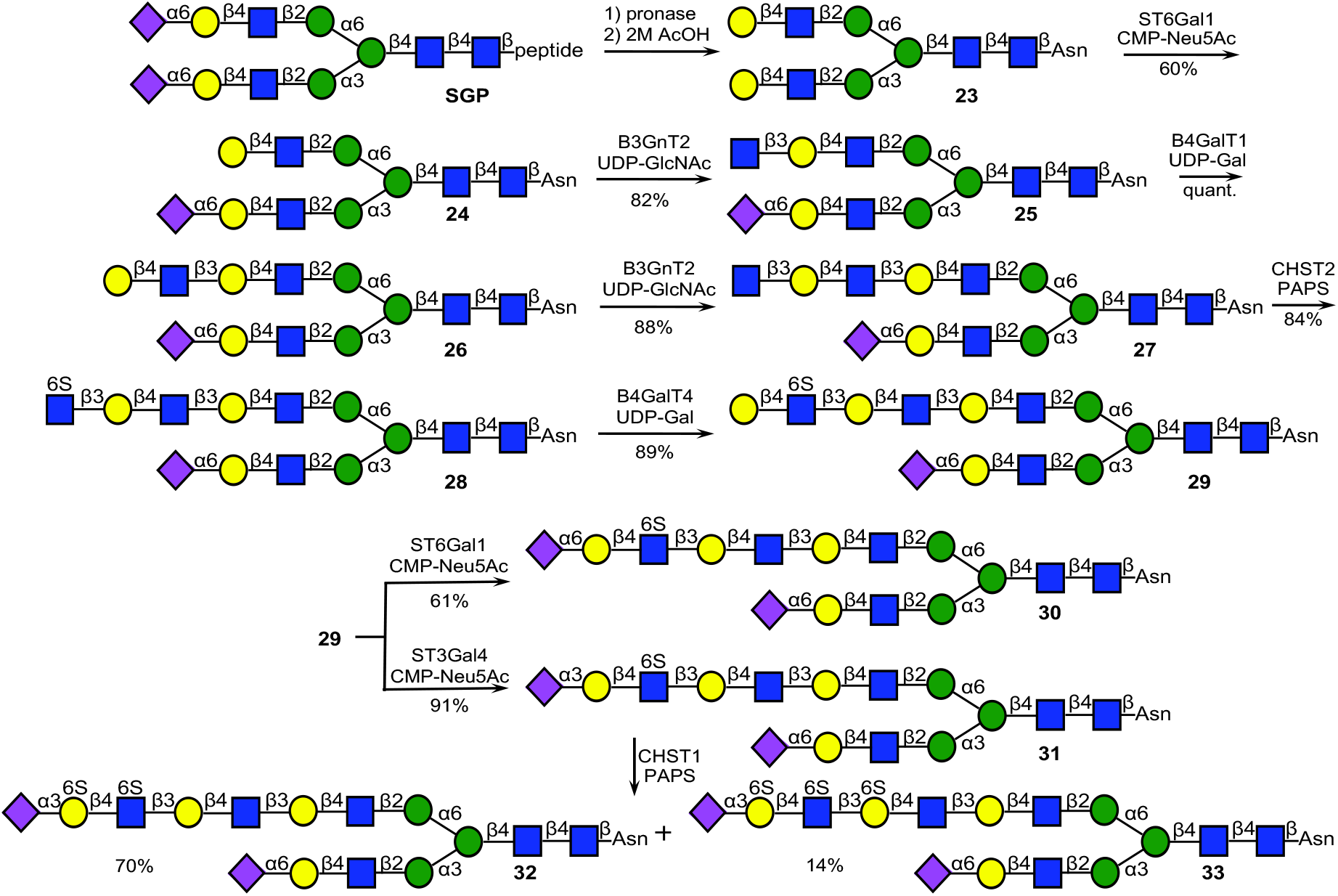
Chemoenzymatic synthesis of KS-I oligosaccharides by enzymatic modification of an *N*-linked glycan obtained from egg yolk powder and exploitation of the regioselectivities of CHST1 and CHST2.

The newly synthesized glycans and several reference compounds, which include several *O*-glycans presenting relevant epitopes (**Q**-**S**), were employed to construct a glycan microarray. The linear derivatives have an anomeric aminopentyl and the *N*-glycans an asparagine moiety, which facilitated printing on amine reactive *N*-hydroxysuccinimide (NHS) activated glass slides to give a glycan microarray (Fig. 2a). The array was probed by several plant lectins and galectin-3 (Gal-3) (Fig. 2b), sialic acid binding immunoglobulin type lectins (Siglecs) (Fig. 2c) and recombinant hemagglutinins (HAs) of animal and human influenza viruses (Fig. 2d-g). As expected, MAL-I and MAL-II bound 2,3-sialylated LacNAc containing structures (**J, M**, and **Q**). In the case of MAL-1, sulfation at GlcNAc (**K, N**, and **R**) substantially reduced the responsiveness, whereas for MAL-II it was tolerated. Further sulfation at Gal (**L, O**, and **S**) was not allowed by these lectins. SNA recognizes 2,6-linked sialosides and as expected compound (**D**-**I**) having such a structural element were well recognized by this lectin and it appears that sulfation of GlcNAc as in compound **G, H**, and **I** did not impede on binding. Compounds **N**-**P** gave relatively low responses indicating that the 2,6-sialoside at the MGAT1 arm is not well recognized by this lectin possibly through interference by the other arm.

**Figure 2.**
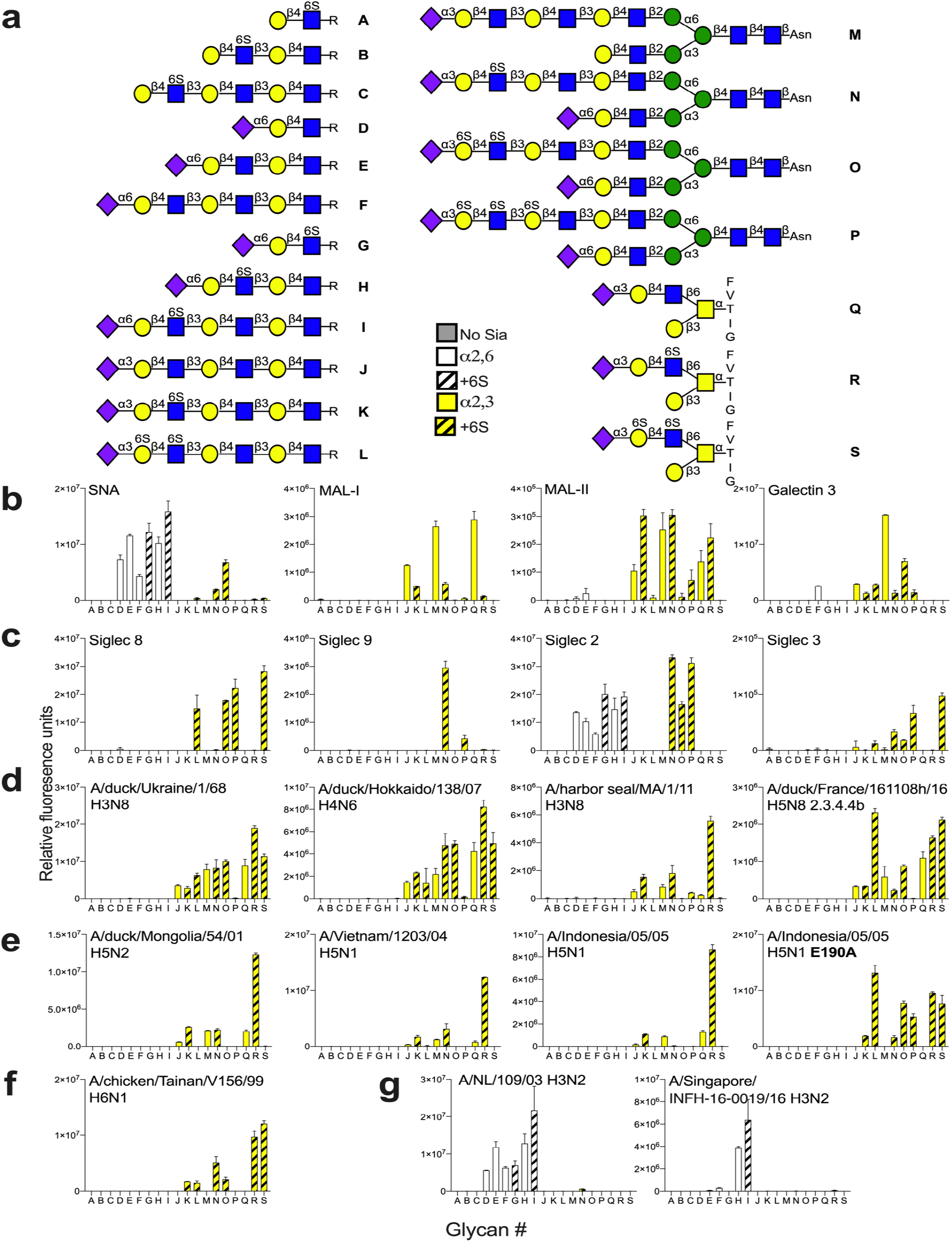
Probing glycan binding properties of lectins, glycan binding proteins and influenza A hemagglutinins. **A**) Collection of glycans printed on succinimide reactive microarray slides. Glycan binding data of **B**) lectins; **C**) Siglec proteins; and **D**) HAs of various flu viruses. Bars represent the background-subtracted average relative fluorescence units (RFU) of four replicates ± SD.

Siglecs are receptors expressed by cells of the immune system that can bind specific sialic acid containing glycoconjugates thereby modulating immune responses.^27^ The sialic acid–Siglec axis plays an important role in the balance between self- and non-self and is disturbed in diseases such as cancer, auto-immunity and allergy.^28^ Genetic engineering of cells has indicated that sulfation can regulate Siglec binding, however, the precise molecular mechanisms of this regulation are not well understood.^29,30^

Histological studies have shown that airway tissues express high molecular weight ligands for Siglec-8 and 9.^31^ Treatment of these tissues with keratanase-I abolished binding of recombinant Siglec-8. There is evidence to supports that in inflamed tissue human eosinophils and mast cells, which express Siglec-8, bind to sialoglycans to resolve inflammation and limit tissue damage.^31^ Previous glycan array studies indicated that Siglec-8 binds to glycans having a terminal Neu5Acα2,3(6-sulfo)-Gal moiety.^32^ The enzymatic studies described here have indicated that sulfation of Gal most like occurs in the context of a neighboring (6-sulfo)-GlcNAc moiety, and thus these ligands are the most likely natural candidates for Siglec-8. Indeed, when the microarray was probed with recombinant Siglec-8, binding was only observed to compounds **L, O, P**, and **S**, which have a sialoside 2,3-linked to a LacNAc moiety sulfated at Gal and GlcNAc (Fig 2c). It appears that presentation in the context of an *N*- or *O*-glycan does not modulate binding and compound **O** and **S** gave similar responsiveness. Siglec-8 tolerates sulfation of the subsequent Gal, as in compound **P**, which is in agreement with the finding that this Siglec binds to KS.^32^ Siglec-9 exhibited a different binding pattern and preferentially bound to compound **N** which has a Neu5Acα2,3Gal β1,4GlcNAc6S moiety presented in the context of an *N*-glycan (Fig. 2c). When this epitope was part of an *O*-glycan, as in compound **R**, binding was diminished. Siglec-2 (CD22) is expressed by B-cells and recognizes 2,6-linked sialoglycans. The array data showed that it tolerates sulfation at the GlcNAc moiety (**D** *vs*. **G, E** *vs*. **H, F**, *vs*. **I**). Siglec-3 (CD33) controls activation of microglial, however, in Alzheimer disease it is overactivated due to the presence of amyloid and tau proteins. Siglec-3 showed a binding pattern that is like that of Siglec-8 (Fig. 2c). Finally, we investigated binding properties of Gal-3, which prefers extended LacNAc moieties, and as expected compound **M** was well recognized whereas sulfation of the extended LacNAc chain resulted in substantial reduced binding (Fig. 2b).

The expression of KS in airway tissues makes these biomolecules potential candidates as receptors for respiratory viruses. Other GAGs such as heparan sulfate have been implicated in viral infections,^33^ however, the role of KS in such processes has received little attention. Hemagglutinin (HA) of influenza A viruses binds to sialoglycans of the host for cell entry. Avian viruses preferentially bind 2,3-linked sialic acids whereas human viruses recognize α2,6-linked sialic acids. Several human and avian viral HAs can bind to sialoglycans modified by sulfates or fucosides and such binding preferences may represent species barriers.^34-37^ We used the array to examine binding selectivities of several recombinant HAs that are derived from viruses that infect different host species such as H3N8 and H5Nx viruses. Additionally, HAs derived from other subtypes that are mostly maintained in avian species were also examined.

Recombinant HAs derived from duck H3N8 and H4N6 viruses bound 2,3-linked sialoside and tolerated sulfation at Gal as well as GlcNAc, and the sulfates had relatively little impact on the responsiveness (Fig. 2d). Further sulfation of Gal, as in compound **P**, was not tolerated. Another duck virus (A/duck/France/161108h/16, H5N8), which is representative of current epizootic outbreaks,^38^ exhibited a similar promiscuous binding behavior (Fig. 2d). HA derived from a harbor seal H3N8 virus showed a different binding pattern and allowed only sulfation at GlcNAc (Fig. 2d). Presentation of the Neu5Ac2,3-Gal-GlcNAc6S epitope in the context of an *O*-glycan resulted in the strongest binding (**Q** *vs*. **R**).

Several H5 proteins (A/duck/Mongolia, A/Indonesia and A/Vietnam) also showed a strong preference for an *O*-glycan presenting the Neu5Ac2,3-Gal-GlcNAc6S epitope (**R**) (Fig. 2e). In a previous study, we analyzed amino acid mutations that occurred during a human infection, one of these diminished binding to sialylated LacNAc (A/Vietnam E190A).^39^ Here, we observed that this mutant HA exclusively binds receptors that include a 6-sulfated GlcNAc.

An HA from a chicken H6N1 virus had an obligatory requirement for sulfation of GlcNAc (**K, N**, and **R**) and tolerated further sulfation of Gal (**L, O**, and **S**) (Fig. 2f).^40^ The highest responsiveness was observed when the sulfated epitope is presented as part of an *O*-glycan. We also examined receptor specificities of two human H3N2 viruses (Fig. 2g). As expected, these viruses only bound to 2,6-linked sialosides. The more recent A/H3N2/Singapore required the presentation of the 2,6-linked sialoside at an extended LacNAc moiety (**H**), which agrees with previous observations.^41^ Interestingly, both viruses tolerated a GlcNAc6S moiety (**F** *vs*. **G** and **H** *vs*. **I**). Collectively, the microarray screening indicate that not only sialic linkage type but also sulfation and presentation of an epitope as part of an *N*- or *O*-glycan can modulate receptor specificity.

## CONCLUSIONS

The results presented here show that the sites that will be sulfated by CHST1 is due to an intricate interplay of enzyme-substrate specificities for sulfated and sialylated structures. In particular, a GlcNAc6S moiety and 2,3-sialylation determines the site of sulfation and in combination with the regioselectivity of CHST2, which only modifies terminal GlcNAc moieties, made it possible to prepare a range of well-defined KS oligosaccharides. The compounds and several reference derivatives were printed as a glycan microarray which uncovered binding selectivities of a range of lectins, glycan binding proteins and HAs from influenza A viruses. It was found that sulfation can greatly influence recognition and can either be tolerated, reduce, or increase binding. Modification of Gal and GlcNAc exert differential effects, which is further influenced by the presentation of a sulfated epitope as part of an *N*- or *O*-glycan. Thus these structure-binding relationships highlight that glycan complexity can modulate protein binding properties. Future efforts will focus on the preparation of a broader range of KS oligosaccharides to examine effects of sulfation on protein binding.

## ASSOCIATED CONTENT

### Supporting Information

The Supporting Information is available free of charge at https://pubs.acs.org. Synthetic protocols, compound characterization, experimental procedure for enzyme and protein expression, microarray screening, and copies of NMR spectra (PDF).

## Author Contributions

G.J.B., Y.W. and GV designed the project. G.J.B. was responsible for overall project management. Y.W. performed all chemical and enzymatic synthesis. C.H. and D.C expressed the glycosyltransferases and CHST1 and 2, which were supervised by K.W.M. Siglec proteins were expressed by A.L.M.K under supervision of R.P.dV. Microarray screening and analysis was performed by R.P.dV. The manuscript was written by G.J.B. and Y.W. All authors provided feedback on the manuscript.

## Note

The authors declare no competing financial interest.

## ACKNOWLEDGMENTS

This research was supported by the European Commission (grants 101020769 and 802780) to G.J.B. and R.P.dV, respectively and the Chinese Scholarship Council (to Y.W).

## Table of Contents Graphic

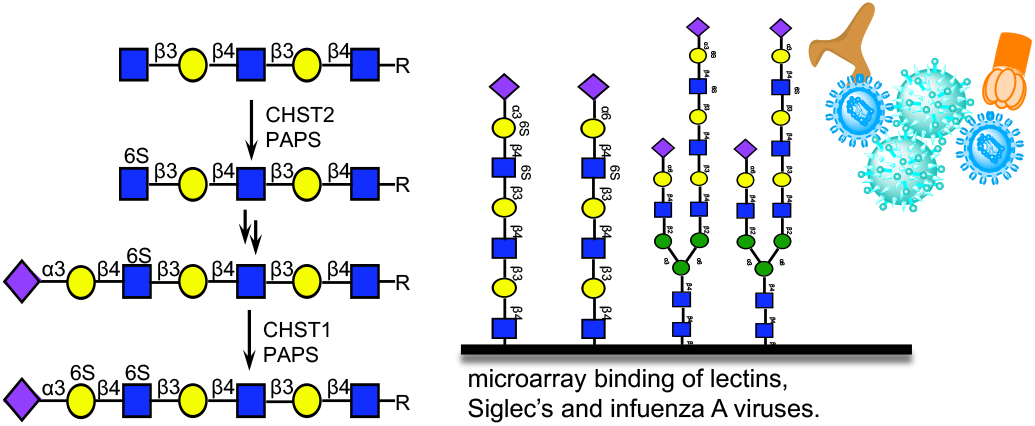

### Table of Contents Text

We describe a chemoenzymatic strategy to prepare well-defined keratan sulfate oligosaccharides by exploiting substrate specificities of relevant sulfotransferases. The KS oligosaccharides were printed as a microarray to explore binding preferences of lectins, Siglec proteins and influenza A viruses.

