## Supplementary Information for "Exploiting Substrate Specificities of 6-*O*-Sulfotransferases to Enzymatically Synthesize Keratan Sulfate Oligosaccharides"

#### 1) Supplementary Scheme

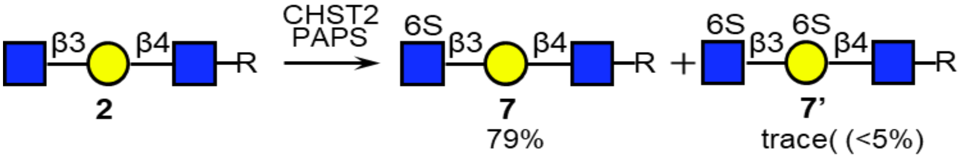

Scheme S1. substrate specificity of CHST2.

#### 2) Materials and Methods

##### Materials

Reagents were purchased from Sigma-Aldrich. Uridine 5'-diphosphogalactose (UDP-Gal), uridine 5'-diphospho-N-acetyl-glucosamine (UDP-GlcNAc) and cytidine-5'-monophospho-N-acetylneuraminic acid (CMP-Neu5Ac) were obtained from Roche Diagnostics [UDP-Gal: Cat# 07703562103; UDP-GlcNAc: Cat# 06369855103; CMPNeu5Ac: Cat# 05974003103]. Adenosine 3′-Phosphate 5′-Phosphosulfate (PAPS) were obtained from Merck [Cat# 118410, Purity⩾80% by HPLC]. Progress of reactions was monitored by liquid chromatography mass spectrometry system (LC-MS) from Shimadzu (system controller: SCL10A-VP; HPLC pumps: LC10AD-VP; injector: SIL10AD-VP) using a ZIC HILIC column (ZeQuant, PEEK coated guard HPLC column, 3.5 µm particle size, 20x 2.1 mm). The LC system was attached to a Bruker Daltonics micro TOF-Q mass spectrometer. Mass spectra were recorded on either on an Applied Biosystems SCIEX MALDI TOF/TOF 5800 mass spectrometer, a Shimadzu Biotech Axima-CFR MALDI-TOF, or a high-resolution Shimadzu LCMS-IT-TOF mass spectrometer. Reaction mixtures were purified using a size exclusion Biogel (P2) and Biogel (P6) from BioRad in Econo glass columns (0.7 x 30 cm / 1.5 x 30 cm / 1.5 x 50 cm/ 1.5 x 120 cm) coupled to a BioFrac fraction collector (BioRad). Carbohydrate-containing fractions were detected by thin layer chromatography and an appropriate staining reagent (15 mL AcOH and 3.5 mL p-anisaldehyde in 350 mL EtOH and 50 mL $H_2SO_4$). If needed further purification was performed by LC-MS using a ZIC HILIC column.

##### Expression and purification of recombinant human glycosyltransferases and sulfotransferases

Expression constructs were generated encoding the truncated catalytic domains of human glycosyltransferases (B4GALT1, B4GALT4, B3GNT2, ST6GAL1, and ST3GAL4) and sulfotransferases (CHST1 and 2) as $NH_2$-terminal fusion proteins in the pGEn2 expression vector essentially as described in prior studies.[1] Briefly, the fusion protein coding regions were comprised of a 25-amino acid signal sequence, an $His_8$ tag, AviTag, the "superfolder" GFP coding region, the 7-amino acid recognition sequence of the tobacco etch virus (TEV) protease followed by the respective catalytic domain regions (for human CHST1 (Uniprot ID: O43916) and CHST2 (Uniprot ID: Q9Y4C5) catalytic domain region comprising of 388 and 454 amino acid residues, respectively). The recombinant human glycosyltransferases and sulfotransferases were expressed as soluble secreted proteins by transient transfection of suspension culture HEK293-F cells (FreeStyle™ 293-F cells, Thermo Fisher Scientific,

Waltham MA) and purified by $Ni^{2+}$-NTA chromatography as previously described.[1,2] Each protein was concentrated to approximately 3 mg/mL using an ultrafiltration pressure cell (Millipore, Billerica, MA) with a 10-kDa molecular mass cutoff membrane. The enzymes were further purified by gel filtration on a Superdex G-75 column (GE Healthcare) preconditioned with a buffer containing 20 mM HEPES, 150 mM NaCl, 0.05% sodium azide, pH 7.0. Peak fractions of recombinant human enzymes were pooled, respectively, concentrated at 1 mg/mL and buffer exchanged with 20 mM HEPES, 100 mM NaCl, 0.05% sodium azide, pH 7.0, 10% glycerol. The final protein preparations were aliquoted and stored at -80 °C until use.

**Extraction and isolation of a sialoglycopeptide (SGP) from egg yolk powder**

SGP was extracted according to our previously reported procedure.[3] In short, commercially available egg yolk powder (Natural Foods, Inc., 2.27 Kg) was suspended twice in 95% ethanol (4 L) and mechanically stirred for 2 h at room temperature to remove lipids and other organic soluble components. The filtrate was discarded and the insoluble powder was suspended twice in aqueous ethanol (40% v/v ethanol, 3 L) solution. The insoluble material was discarded and the filtrate was concentrated under reduced pressure at 40 °C. The resulting translucent liquid was purified using an active carbon / celite column (500 g of active carbon and 500 g celite). Impurities were removed by flushing the column with 3 L of water (0.1% v/v TFA), 3 L of 5% acetonitrile in water (0.1% v/v TFA), and 3 L 10% acetonitrile in water (0.1% v/v TFA). The desired glycopeptide was released from the column using a solution of 25% acetonitrile in water (0.1% v/v TFA), and fractions containing the product were pooled and dried under reduced pressure. The resulting white powder was subjected to size-exclusion chromatography (Bio-Rad® P-2, fine particle size 45 – 90 μm, column dimensions 5.0 cm x 80 cm, 250 mL fractions) eluting with 0.1 M ammonium bicarbonate to yield SGP as a fluffy, white powder (1.82 g, or 0.8 mg SGP / g egg yolk powder).

**3) General Protocols for Enzymatic Reactions**

**General Procedure for the Installation of β1,3-GlcNAc using B3GnT2**

Glycosyl acceptor (1 eq) and UDP-GlcNAc (1.5 eq) were dissolved to provide a final acceptor concentration of 2 – 5 mM in a HEPES buffered solution (50 mM, pH 7.3) containing KCl (25 mM), $MgCl_2$ (2 mM) and DTT (1 mM). Calf intestine alkaline phosphotase (CIAP, 1% total volume, 1 kU/mL) and B3GnT2 (1% wt/wt relative to acceptor substrate) were added, and the reaction mixture was incubated overnight at 37 °C with gentle shaking. Progress of the reaction was monitored by MALDI-TOF MS or ESI-TOF MS, and if starting material remained after 18 h another portion of B3GnT2 was added until no starting material could be detected. The reaction mixture was centrifuged over a Nanosep® Omega ultrafiltration device (10 kDa MWCO) to remove proteins and the filtrate was lyophilized. The residue was applicated to P2 or P6 size-exclusion column chromatography using Milli–Q water as eluent providing the desired product. High performance liquid chromatography (HPLC) using HILIC column (see materials) was employed when the impurities were detected.

##### General Procedure for the Installation of β1,4-Gal using B4GalT1

Glycosyl acceptor (1 eq) and UDP-Gal (1.5 eq per Gal to be added) were dissolved to provide an acceptor concentration of 2 – 5 mM in a Tris buffered solution (100 mM, pH 7.5) containing $MnCl_2$ (10 mM) and BSA (1% total volume). CIAP (1% volume total) and B4GalT1 (1% wt/wt relative to acceptor substrate) were added, and the reaction mixture was incubated overnight at 37 °C with gentle shaking. Progress of the reaction was monitored by MALDI-TOF MS or ESI-TOF MS, and if starting material was remaining after 18 h, another portion of B4GalT1 was added until no starting material could be detected. The reaction mixture was centrifuged over a Nanosep® Omega ultrafiltration device (10 kDa MWCO) to remove proteins and the filtrate was lyophilized. The residue was applicated to P2 or P6 size-exclusion column chromatography using Milli–Q water as eluent providing the desired product. High performance liquid chromatography (HPLC) using HILIC column (see materials) was employed when impurities were detected.

##### General Procedure for the Installation of β1,4-Gal using B4GalT4

Glycosyl acceptor (1 eq) and UDP-Gal (1.5 eq per Gal to be added) were dissolved to a provide an acceptor concentration of 2 – 5 mM in a Tris buffered solution (100 mM, pH 7.5) containing $MnCl_2$ (10 mM) and BSA (1% total volume). CIAP (1% volume total) and B4GalT4 (1% wt/wt relative to acceptor substrate) were added, and the reaction mixture was incubated overnight at 37 °C with gentle shaking. Reaction progress was monitored by MALDI-TOF MS or ESI-TOF MS, and if starting material remained after 18 h another portion of B4GalT4 was added until no starting material could be detected. The reaction mixture was centrifuged over a Nanosep® Omega ultrafiltration device (10 kDa MWCO) to remove reaction proteins and the filtrate was lyophilized. The residue was applicated to P2 or P6 size-exclusion column chromatography using Milli–Q water as eluent provided the desired product. High performance liquid chromatography (HPLC) using a HILIC column (see materials) was employed when impurities were remaining.

##### General Procedure for the Installation of α2,3-Neu5Ac using ST3Gal4

Glycosyl acceptor (1 eq) and CMP-Neu5Ac (1.5 eq) were dissolved at a final acceptor concentration of 2 –5 mM in a HEPES-buffered solution (50 mM, pH 7.2) containing BSA (1% total volume). CIAP (1% volume total) and ST3Gal4 (1% wt/wt relative to acceptor substrate) were added, and the reaction mixture was incubated overnight at 37 °C with gentle shaking. Progress of the reaction was monitored by ESI-TOF MS, and if starting material remained after 18 h another portion of ST3Gal4 was added until no starting material could be detected. The reaction mixture was centrifuged over a Nanosep® Omega ultrafiltration device (10 kDa MWCO) to remove proteins, and the filtrate was lyophilized. The residue was applicated to P2 or P6 size-exclusion column chromatography using Milli–Q water as eluent to provide the desired product. High performance liquid chromatography (HPLC) using HILIC column (see materials) was employed when impurities were detected.

**General Procedure for the Selective Installation of Terminal α2,6-Neu5Ac using ST6Gal1**

Glycosyl acceptor (1 eq) and CMP-Neu5Ac (1.1 eq) were dissolved at a final acceptor concentration of 2 – 5 mM in a HEPES-buffered solution (100 mM, pH 7.5) containing BSA (1% volume total). CIAP (1% volume total) and ST6GAL1 (1% wt/wt relative to acceptor substrate) were added, and the reaction mixture was incubated overnight at 37 °C with gentle shaking. The reaction mixture was centrifuged over a Nanosep® Omega ultrafiltration device (10 kDa MWCO) to remove reaction proteins, and the filtrate was lyophilized. The residue was applicated to P2 or P6 size-exclusion column chromatography using Milli–Q water as eluent to provide the desired product. High performance liquid chromatography (HPLC) using HILIC column (see materials) was employed when impurities were detected (see **5** for further details).

**General Procedure for the 6-O-Sulfate Installation of Terminal GlcNAc using CHST2**

Glycosyl acceptor (1 eq) and PAPS (1.6 eq) were dissolved at a final acceptor concentration of 2–5 mM in a Tris buffered solution (100 mM, pH 7.5) containing $MgCl_2$ (10 mM). CHST2 (10%-20% wt/wt relative to acceptor substrate) were added, the reaction mixture was incubated overnight at 37 °C with gentle shaking. The reaction mixture was centrifuged over a Nanosep® Omega ultrafiltration device (10 kDa MWCO) to remove reaction proteins, and the filtrate was lyophilized. The residue was applicated to P2 or P6 size-exclusion column chromatography using $NH_4HCO_3$ buffer (50 mM) as eluent to provide the desired product. High performance liquid chromatography (HPLC) using HILIC column (see materials) or DEAE ion exchange column was employed when impurities were detected.

**General Procedure for the 6-O-Sulfate Installation of Internal Galactose using CHST1**

Glycosyl acceptor (1 eq) and PAPS (1.6 eq per galactose) were dissolved at a final acceptor concentration of 2–5 mM in a Tris buffered solution (100 mM, pH 7.5) containing $MgCl_2$ (10 mM). CHST1 (10% wt/wt relative to acceptor substrate) were added, and the reaction mixture was incubated overnight at 37 °C with gentle shaking. The reaction mixture was centrifuged over a Nanosep® Omega ultrafiltration device (10 kDa MWCO) to remove reaction proteins, and the filtrate was lyophilized. The residue was applicated to P2 or P6 size-exclusion column chromatography using $NH_4HCO_3$ buffer (50 mM) as eluent to provide the desired product. High performance liquid chromatography (HPLC) using HILIC column (see materials) was employed when impurities were detected.

**Procedure for Rate-controlled Synthesis of Compound 32 and 33 using CHST1**

Glycosyl acceptor **31** (1 eq) and PAPS (1 eq) were dissolved at a final acceptor concentration of 2 mM in a Tris buffered solution (100 mM, pH 7.5) containing $MgCl_2$ (10 mM). CHST1 (10% wt/wt relative to acceptor substrate) were added, the reaction mixture was incubated overnight at 37 °C with gentle shaking. Another portion of PAPS (0.5 eq) was added followed by incubation at 37 °C with gentle shaking. The progress of the reaction was monitored by ESI-TOF MS until **31** was not further consumed. The reaction mixture was centrifuged over a Nanosep® Omega ultrafiltration device (10 kDa MWCO) to remove proteins, and the filtrate was lyophilized. The residue was applicated to P2 or P6 size-exclusion column chromatography using $NH_4HCO_3$ buffer (50 mM) as eluent provided the desired product. High performance liquid chromatography (HPLC) using HILIC column (see materials) or DEAE ion exchange column was employed when the impurities were remaining.

**4) General Procedure for hydrogenation of Cbz Protecting Group using $Pd(OH)_2$**

Palladium hydroxide on carbon (Degussa type, 20%, 1.5 times the weight of starting material) was added to a solution of the starting material in $H_2O$ (0.1% AcOH as additive). The mixture was placed under an atmosphere of hydrogen until ESI-LC-MS indicated completion of the reaction. The mixture was filtered through a spin filter and the residue was washed with $H_2O$. The filtrate was lyophilized to give the final product. P6 size-exclusion column chromatography was used for purification with 50 mM ammonium bicarbonate as eluent. The fractions containing compound were lyophilized to give the desired product as a white powder.

**5) General Protocols for HILIC-HPLC Purification with MS Detection**

**HILIC-HPLC Purification Conditions**

Semi-preparative HILIC-HPLC was performed on a Shimadzu (LC-20AT, SIL-20A, CBM-20A, SPD-20A, FRC-10A) LC-ESI-IT-TOF with a XBridge HILIC column, 5 µm, 10 x 250 mm at a flow rate of 3.6 mL/min, injection volume of 100 µL (10-20 mg/mL), with 0.2% of the flow is diverted to the ESI-MS detector using a splitter. The purification was done using 10% 10 mM $NH_4HCO_3$ in MeCN (buffer B) and MeCN in 80% 10 mM $NH_4HCO_3$ (buffer A).

The general condition using a linear gradient is as follows:

Linear glycan

| Time (min) | A (%) | B (%) |
|---|---|---|
| 0 | 10 | 90 |
| 90 | 50 | 50 |

N-glycan

| Time (min) | A (%) | B (%) |
|---|---|---|
| 0 | 20 | 80 |
| 60 | 45 | 55 |
| 70 | 45 | 55 |

**6) Analytical Data**

**NMR Nomenclature**

VnmrJ 4 and Topspin 4 were used to collect NMR data. NMR data was obtained at room temperature on a 600 MHz instrument from Bruker. The chemical shift δ is given in parts per million (ppm) and refers to tetramethyl silane and the residual solvent peak [$^1$H-NMR: δ(D2O) =4.79 ppm]. NMR data is given as follows: $^1$H-NMR: chemical shift (multiplicity, coupling constants, relative integral, functional group); $^{13}$C data are extracted from HSQC spectra and given as follows: chemical shift. Multiplicity is defined as follows: s = singlet; d = doublet; t = triplet; m= multiplet. Signals were assigned by numbering the monosaccharide units starting at the reducing end of the oligosaccharide. Monosaccharides attached to the mannose-3 branch are indicated by a“ ’ “ (prime) and those attached to the mannose-2 branch without any mark. The assignment was done by using corresponding 2D-NMR spectra (COSY, HSQC, TOCSY, NOESY). The yield/concentration of the final products was determined by NMR spectroscopy, using n-propanol as an internal standard. High resolution masses were measured on an Agilent 6560 Ion Mobility Q-TOF LC-MS system.

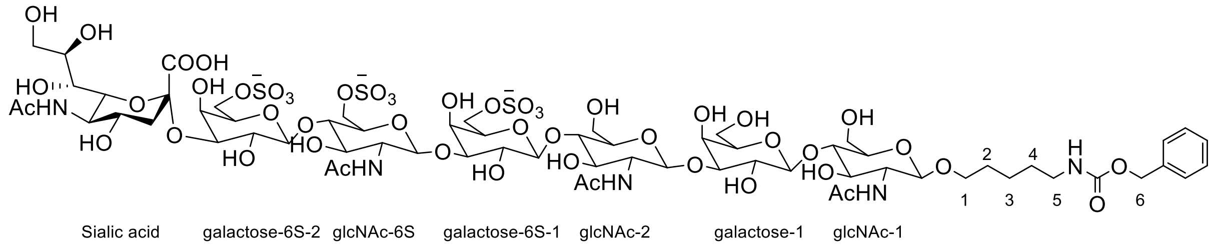

Linear glycan labelling system for peak assignment in NMR data.

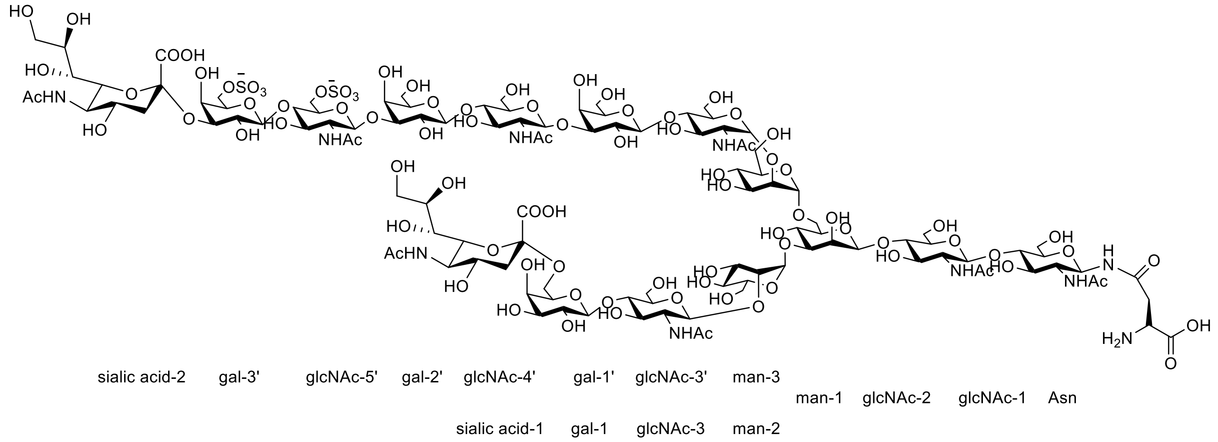

N-glycan labelling system for peak assignment in NMR data.

#### Determination of Sulfate Position

#### Characterization of Terminal GlcNAc 6-O-Sulfate of Compound 7

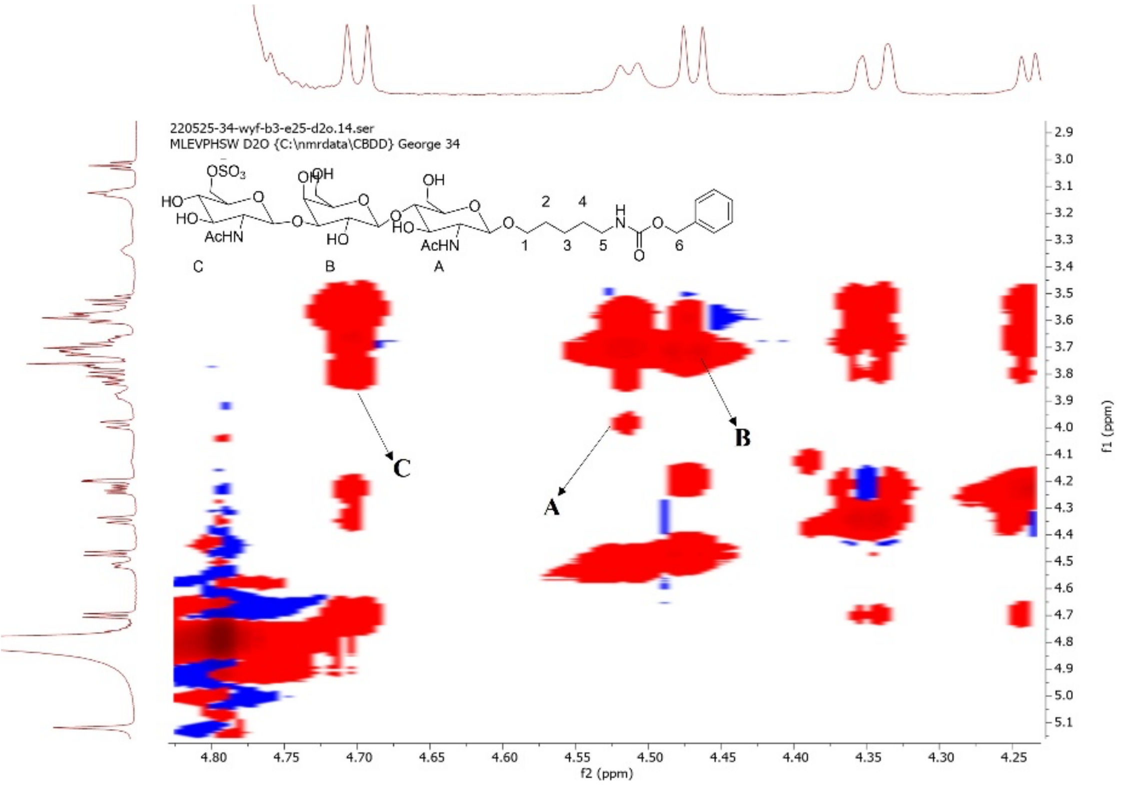

**Figure S1a.** Analysis TOCSY of compound **7**

The position of the sulfate was confirmed by a combination of $^{1}$H, COSY, NOESY, TOCSY and HSQC NMR experiments using compound **7**. The 1D $^{1}$H NMR and 2D $^{13}$C–$^{1}$H HSQC spectra of **7** are depicted in Fig. S1b.

The 1D $^{1}$H NMR spectrum of **7** shows three anomeric signals, correlating to residues A, B and C. The H-1 signal at δH 4.51 (A) is stemming from a reducing-end GlcNAc residue, whereas the H-1 signal at δH 4.47 (B) belongs to the non-reducing βGal. The anomeric signal at δH 4.70 (C) belongs to non-reducing βGlcNAc, respectively.

In the TOCSY spectrum (80 ms, Fig. S1a), the H-1 tracks of A and C show complete spin systems H-1,2,3,4,5,6a,6b, typical for βGlcNAc residues.

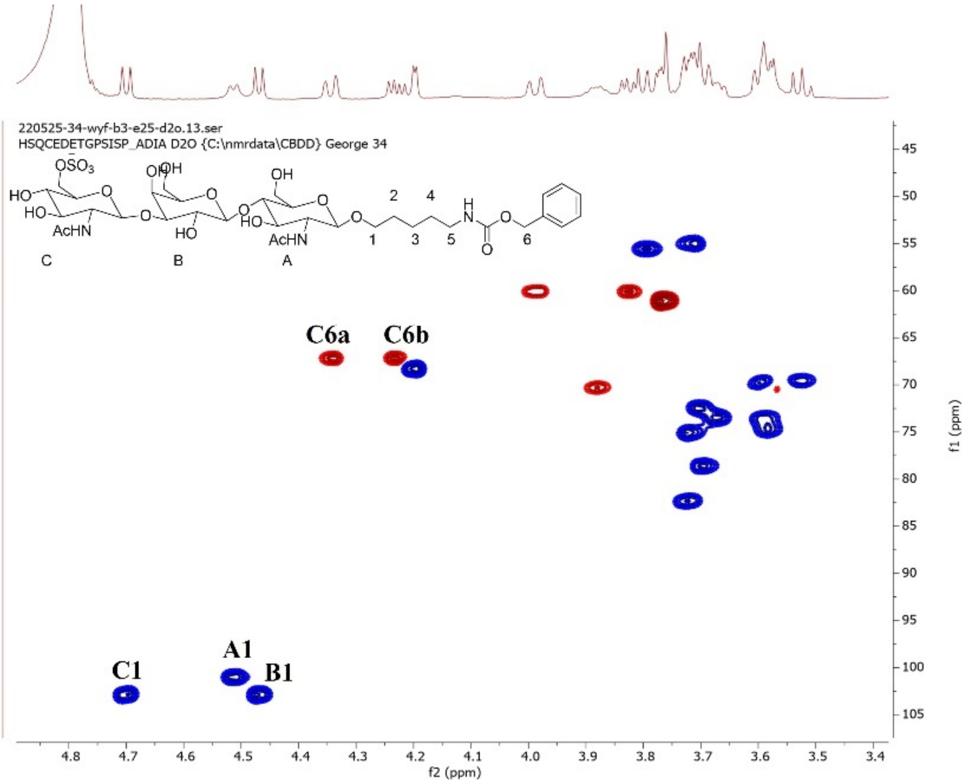

**Figure S1b.** Analysis HSQC of compound **7**

The HSQC spectrum (Fig. S1b) showed downfield shifts for GlcNAc-C6 [δC-6 67.2] from (δ 60.2). The corresponding protons shifted from (H6a δ 3.98, H6b 3.83) to (H6a δ 4.34, H6b 4.23), which is indicative of sulfation. The -1 site Gal B H-4 also shifted from (H4 δ 4.16) to (H4 δ 4.20) indicating nearby sulfation.

In the 2D NOESY spectrum (300 ms, not shown), the inter-residue connectivities GlcNAc-C H-1,Gal-B H-3 and Gal-B H-1, GlcNAc-A H-4 are in accordance with C(1→3)B, B(1→4)A linkages, respectively.

#### Characterization of Internal Galactose 6-O-Sulfation of Compound 19

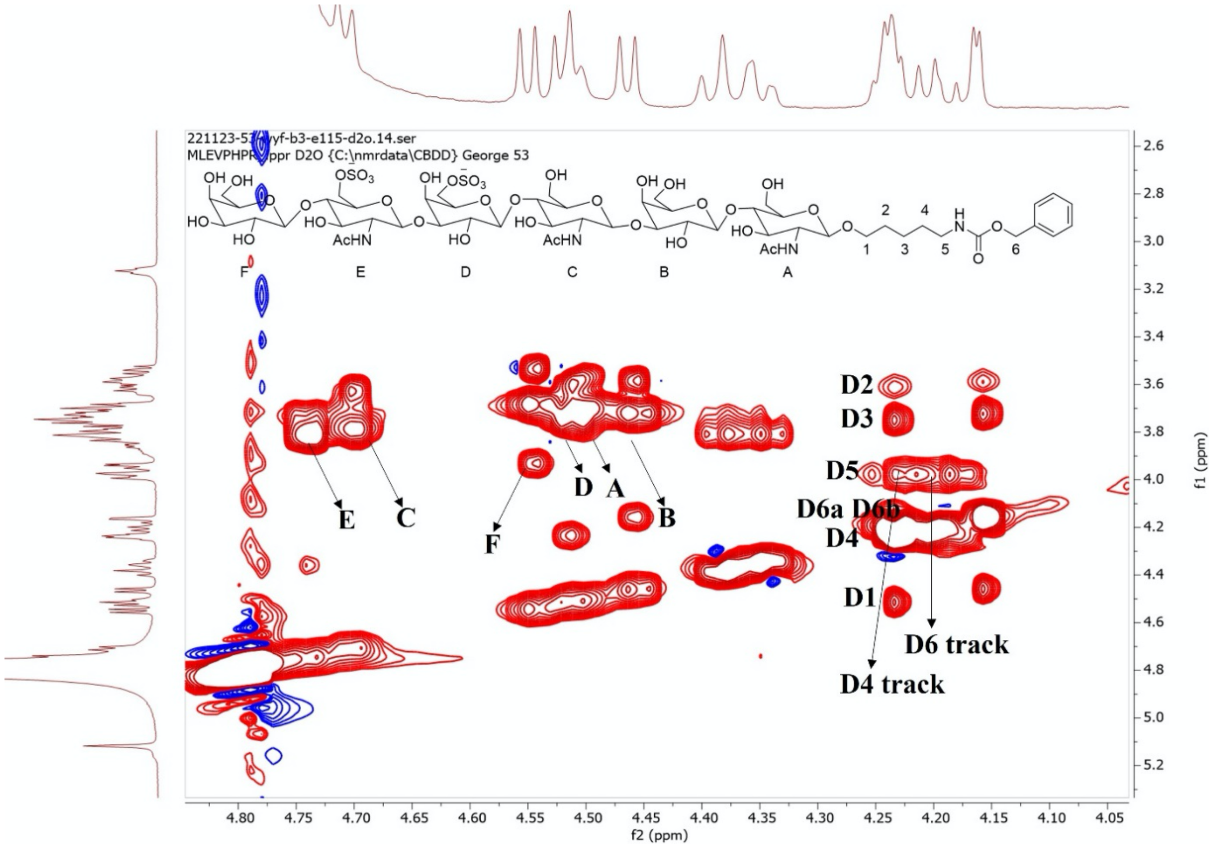

**Figure S2a.** Analysis TOCSY of compound **19**

The position of the sulfate was confirmed by a combination of $^{1}$H, COSY, NOESY, TOCSY and HSQC NMR experiments. The 1D $^{1}$H NMR and 2D $^{13}$C–$^{1}$H HSQC spectra of **19** are depicted in Fig. S2b.

The 1D $^{1}$H NMR spectrum of **19** showed six anomeric signals, correlating with residues A, B, C, D, E and F. The H-1 signals at δH 4.51 (A) stemming from a reducing-end GlcNAc residue whereas the H-1 signal at δH 4.46 (B), δH 4.52 (D) and δH 4.55 (F) belong to non-reducing βGal. The anomeric signals at δH 4.71 (C) and δH 4.74 (E) belong to non-reducing βGlcNAc, respectively.

In the TOCSY spectrum (80 ms, Fig. S2a), the H-1 track of D show spin systems H-1,2,3,4, typical for βGal residues. In the TOCSY and COSY spectra (not shown) the H-6a, 6b track of D allowed the observation of cross-peaks with H-5. Finally, the Gal-D H-4 tracks a cross-peak with D H-5 was found *via* the D H-4 track.

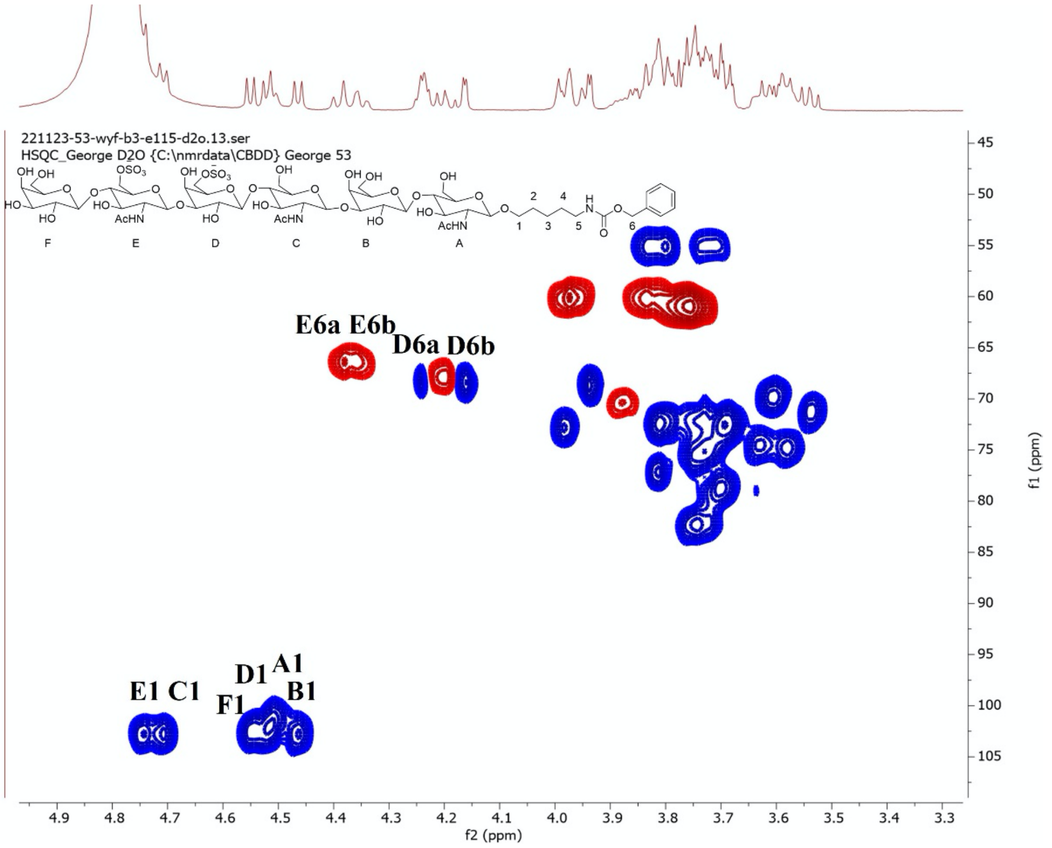

**Figure S2b.** Analysis HSQC of compound **19**

HSQC (Fig. S2b) was used to determine proton-carbon correlations. It showed a downfield shift for Gal-D6 [δC-6 67.9] from (δ 61.3). The corresponding protons shifted from (H6 δ 3.76) to (H6 δ 4.20) indicating sulfation. Gal D H-4 also shifted from (H4 δ 4.20) to (H4 δ 4.24), which indicates a nearby sulfate ester. Gal D H-5 shifted from (H5 δ 3.72) to (H5 δ 3.99), which indicates nearby sulfation. Gal D C-5 also shifted upfield from (δ 75.2) to (δ 72.9), which indicates a nearby sulfate.

In the 2D NOESY spectrum (300 ms, not shown), the inter-residue connectivities Gal-F H-1, GlcNAc-E H-4, GlcNAc-E H-1, Gal-D H-3, Gal-D H-1, GlcNAc-C H-4, GlcNAc-C H-1, Gal-B H-3 and Gal-B H-1, GlcNAc-A H-4 are in accordance with F(1→4)E, E(1→3)D, D(1→4)C, C(1→3)B, B(1→4)A linkages, respectively.

#### 7) Experimental Procedures and Analysis

##### Compound 2

**2** was prepared from **1** (63.0 mg, 104.5 μmol) using the general procedure for the installation of β1,3-GlcNAc with B3GnT2. After P2 purification, **2** was obtained as a white solid (80.8 mg, 96%).

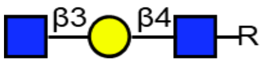

$^{1}H$ (600 MHz, $D_2O$): δ (ppm)

| | H-1 | H-2 | H-3 | H-4 | H-5 | H-6 | NHAc |
|---|---|---|---|---|---|---|---|
| GlcNAc-1 | 4.51 (d, J = 7.7 Hz, 1H) | 3.73 | 3.69 | 3.69 | 3.58 | 3.98 (dd, J = 12.1 Hz, 1H)，3.84 – 3.80 (m, 1H) | 2.07 – 1.98 (m, 6H) |
| Galactose | 4.46 (d, J = 7.9 Hz, 1H) | 3.59 | 3.74 | 4.16 (d, J = 3.3 Hz, 1H) | 3.72 | 3.77 – 3.73 (m, 2H) | - |
| GlcNAc-2 | 4.69 (d, J = 8.5 Hz, 1H) | 3.77 | n/a | n/a | 3.46 | 3.90，3.77 | 2.07 – 1.98 (m, 6H) |

$^{13}C$ (150 MHz, $D_2O$): δ (ppm)

| | C-1 | C-2 | C-3 | C-4 | C-5 | C-6 | NHAc |
|---|---|---|---|---|---|---|---|
| GlcNAc-1 | 101.08 | 55.07 | 72.62 | 78.68 | 74.75 | 60.24 | 22.15 |
| Galactose | 102.93 | 70.05 | 82.18 | 68.50 | 74.96 | 60.76 | - |
| GlcNAc-2 | 102.84 | 55.76 | n/a | n/a | n/a | 60.51 | 22.15 |

| Linker | 1 | 2 | 3 | 4 | 5 | 6 |
|---|---|---|---|---|---|---|
| H | 3.87,3.55 | 1.55 (p, J = 7.0 Hz, 2H) | 1.31 (q, J = 7.3, 6.9 Hz, 2H) | 1.49 (p, J = 7.3 Hz, 2H) | 3.12 (t, J = 6.8 Hz, 2H) | 5.11 (s, 2H) |
| C | 70.54 | 28.35 | 22.47 | 28.58 | 40.71 | 66.78 |

HRMS (ESI-MS): m/z calculated for $C_{35}H_{55}N_3O_{18}$ $[M+Na]^+$: 828.3373; found: 828.3342.

#### Compound 3

**3** was prepared from **2** (80.8 mg, 100.4 μmol) using the general procedure for the installation of β1,4-Gal with B4GalT1 to full conversion. After P2 purification, **3** was obtained as a white solid (92.0 mg, 94%).

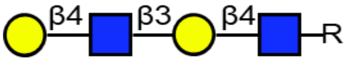

$^{1}H$ (600 MHz, $D_2O$): δ (ppm)

| | H-1 | H-2 | H-3 | H-4 | H-5 | H-6 | NHAc |
|---|---|---|---|---|---|---|---|
| GlcNAc-1 | 4.51 | 3.72 | 3.68 | 3.69 | 3.58 | 3.99, 3.83 | 2.07 – 1.99 (m, 6H) |
| Galactose-1 | 4.46 | 3.60 | 3.73 | 4.16 | 3.72 | 3.76 (4H) | - |
| GlcNAc-2 | 4.71 | 3.82 | 3.73 | 3.74 | 3.59 | 3.97, 3.85 | 2.07 – 1.99 (m, 6H) |
| Galactose-2 | 4.48 | 3.55 | 3.68 | 3.93 | 3.72 | 3.76 (4H) | - |

$^{13}C$ (150 MHz, $D_2O$): δ (ppm)

| | C-1 | C-2 | C-3 | C-4 | C-5 | C-6 | NHAc |
|---|---|---|---|---|---|---|---|
| GlcNAc-1 | 101.20 | 55.12 | 72.47 | 78.52 | 74.79 | 59.96 | 22.18 |
| Galactose-1 | 103.02 | 69.90 | 82.13 | 68.38 | 75.20 | 61.10 | - |

| | | | | | | | |
|---|---|---|---|---|---|---|---|
| GlcNAc-2 | 102.55 | 55.16 | 72.22 | 78.15 | 74.79 | 59.96 | 22.18 |
| Galactose-2 | 102.84 | 71.16 | 72.49 | 68.60 | 75.20 | 61.10 | - |

| Linker | 1 | 2 | 3 | 4 | 5 | 6 |
|---|---|---|---|---|---|---|
| H | 3.87, 3.55 | 1.58 – 1.52 (m, 2H) | 1.35 – 1.25 (m, 2H) | 1.49 (p, J = 7.3 Hz, 2H) | 3.12 (t, J = 6.8 Hz, 2H) | 5.11 (s, 2H) |
| C | 70.47 | 28.25 | 22.31 | 28.41 | 40.48 | 66.75 |

HRMS (ESI-MS): m/z calculated for $C_{41}H_{65}N_3O_{23}$ $[M+Na]^+$: 990.3902; found: 990.4101.

#### Compound 4

**4** was prepared from **3** (30.0 mg, 31.0 μmol) using the general procedure for the installation of β1,3-GlcNAc with B3GnT2. After P2 purification, **4** was obtained as a white solid (33.4 mg, 92%).

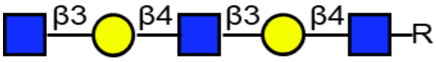

$^1$H (600 MHz, $D_2O$): δ (ppm)

| | H-1 | H-2 | H-3 | H-4 | H-5 | H-6 | NHAc |
|---|---|---|---|---|---|---|---|
| GlcNAc-1 | 4.51 (d, J = 7.7 Hz, 1H) | 3.72 | 3.69 | 3.69 | 3.58 | 3.98, 3.83 | 2.06 – 1.98 (m, 9H) |
| Galactose-1 | 4.46 (d, J = 7.7 Hz, 1H) | 3.59 | 3.73 | 4.16 (d, J = 3.3 Hz, 2H) | 3.72 | 3.76 (4H) | - |
| GlcNAc-2 | 4.70 (d, J = 8.9 Hz, 1H) | 3.81 | 3.73 | 3.73 | 3.59 | 3.97, 3.84 | 2.06 – 1.98 (m, 9H) |
| Galactose-2 | 4.47 (d, J = 7.8 Hz, 1H) | 3.59 | 3.70 | 4.16 (d, J = 3.3 Hz, 2H) | 3.73 | 3.76 (4H) | - |
| GlcNAc-3 | 4.69 (d, J = 8.7 Hz, 1H) | 3.77 | n/a | n/a | 3.46 | 3.90，3.77 | 2.06 – 1.98 (m, 9H) |

$^{13}$C (150 MHz, $D_2O$): δ (ppm)

| | C-1 | C-2 | C-3 | C-4 | C-5 | C-6 | NHAc |
|---|---|---|---|---|---|---|---|
| GlcNAc-1 | 101.07 | 55.33 | 72.51 | 78.48 | 74.70 | 60.01 | 22.20 |
| Galactose-1 | 102.90 | 69.92 | 82.12 | 68.31 | 74.98 | 61.07 | - |
| GlcNAc-2 | 102.83 | 55.16 | 72.26 | 78.35 | 74.67 | 60.03 | 22.20 |
| Galactose-2 | 102.90 | 70.06 | 82.11 | 68.31 | 74.98 | 61.07 | - |
| GlcNAc-3 | 102.83 | 55.74 | n/a | n/a | n/a | 60.43 | 22.20 |

| Linker | 1 | 2 | 3 | 4 | 5 | 6 |
|---|---|---|---|---|---|---|
| H | 3.87, 3.55 | 1.59 – 1.52 (m, 2H) | 1.35 – 1.26 (m, 2H) | 1.49 (p, J = 7.3 Hz, 2H) | 3.12 (t, J = 6.9 Hz, 2H) | 5.11 (s, 2H) |
| C | 70.44 | 28.21 | 22.33 | 28.37 | 40.30 | 66.76 |

HRMS (ESI-MS): m/z calculated for $C_{49}H_{79}N_4O_{28}$ $[M+Na]^+$: 1193.4695; found: 1193.5277.

**Compound 5**

**5** was prepared from **4** (33.4 mg, 28.5 µmol) using the general procedure for the installation of β1,4-Gal with B4GalT1 to full conversion. After P2 purification, **5** was obtained as a white solid (36.0 mg, 95%).

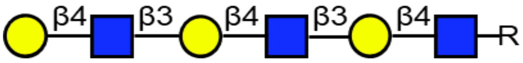

$^{1}H$ (600 MHz, $D_2O$): δ (ppm)

| | H-1 | H-2 | H-3 | H-4 | H-5 | H-6 | NHAc |
|---|---|---|---|---|---|---|---|
| GlcNAc-1 | 4.51 (d, J = 7.7 Hz, 1H) | 3.72 | 3.68 | 3.69 | 3.58 | 3.98, 3.83 | 2.06 – 1.96 (m, 9H) |
| Galactose-1 | 4.46 | 3.59 | 3.72 | 4.16 | 3.72 | 3.76 (6H) | - |
| GlcNAc-2 | 4.71 | 3.81 | 3.73 | 3.74 | 3.59 | 3.96, 3.85 | 2.06 – 1.96 (m, 9H) |
| Galactose-2 | 4.47 | 3.58 | 3.72 | 4.16 | 3.72 | 3.76 (6H) | - |
| GlcNAc-3 | 4.71 | 3.81 | 3.73 | 3.74 | 3.59 | 3.96, 3.85 | 2.06 – 1.96 (m, 9H) |
| Galactose-3 | 4.48 | 3.54 | 3.67 | 3.93 (d, J = 3.4 Hz, 1H) | 3.72 | 3.76 (6H) | - |

$^{13}C$ (150 MHz, $D_2O$): δ (ppm)

| | C-1 | C-2 | C-3 | C-4 | C-5 | C-6 | NHAc |
|---|---|---|---|---|---|---|---|
| GlcNAc-1 | 101.10 | 55.04 | 72.63 | 78.66 | 74.87 | 59.91 | 22.13 |
| Galactose-1 | 102.94 | 70.05 | 82.04 | 68.33 | 75.17 | 61.04 | - |
| GlcNAc-2 | 102.78 | 55.18 | 72.30 | 78.46 | 74.87 | 59.91 | 22.13 |
| Galactose-2 | 102.94 | 70.05 | 82.04 | 68.33 | 75.17 | 61.04 | - |
| GlcNAc-3 | 102.78 | 55.18 | 72.30 | 78.46 | 74.87 | 59.91 | 22.13 |
| Galactose-3 | 102.94 | 71.19 | 72.53 | 68.56 | 75.17 | 61.04 | - |

| Linker | 1 | 2 | 3 | 4 | 5 | 6 |
|---|---|---|---|---|---|---|
| H | 3.87, 3.55 | 1.59 – 1.52 (m, 2H) | 1.34 – 1.28 (m, 2H) | 1.49 (p, J = 7.3 Hz, 2H) | 3.15 – 3.06 (m, 2H) | 5.11 (s, 2H) |
| C | 70.40 | 28.29 | 22.38 | 28.50 | 40.47 | 66.74 |

HRMS (ESI-MS): m/z calculated for $C_{55}H_{89}N_4O_{33}$ $[M+Na]^+$: 1355.5224; found: 1355.5838.

#### Compound 6

**6** was prepared from **5** (13.0 mg, 9.7 μmol) using the general procedure for the installation of β1,3-GlcNAc with B3GnT2. After P6 and HILIC HPLC purification, **6** was obtained as a white solid (9.0 mg, 61%). Unreacted **5** was recovered.

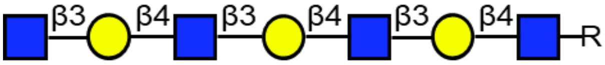

$^{1}H$ (600 MHz, $D_2O$): δ (ppm)

| | H-1 | H-2 | H-3 | H-4 | H-5 | H-6 | NHAc |
|---|---|---|---|---|---|---|---|
| GlcNAc-1 | 4.51 (d, J = 7.6 Hz, 1H) | 3.73 | 3.69 | 3.70 | 3.58 | 3.98, 3.83 | 2.06 – 1.99 (m, 12H) |
| Galactose-1 | 4.46 | 3.59 | 3.73 | 4.16 (d, J = 3.2 Hz, 3H) | 3.72 | 3.76 (6H) | - |
| GlcNAc-2 | 4.71 | 3.81 | 3.73 | 3.74 | 3.59 | 3.96, 3.85 | 2.06 – 1.99 (m, 12H) |
| Galactose-2 | 4.47 | 3.59 | 3.73 | 4.16 (d, J = 3.2 Hz, 3H) | 3.72 | 3.76 (6H) | - |
| GlcNAc-3 | 4.70 | 3.81 | 3.73 | 3.74 | 3.59 | 3.96, 3.85 | 2.06 – 1.99 (m, 12H) |
| Galactose-3 | 4.48 | 3.59 | 3.73 | 4.16 (d, J = 3.2 Hz, 3H) | 3.72 | 3.76 (6H) | - |
| GlcNAc-4 | 4.69 | 3.77 | n/a | n/a | 3.46 | 3.90, 3.78 | 2.06 – 1.99 (m, 12H) |

$^{13}C$ (150 MHz, $D_2O$): δ (ppm)

| | C-1 | C-2 | C-3 | C-4 | C-5 | C-6 | NHAc |
|---|---|---|---|---|---|---|---|
| GlcNAc-1 | 101.18 | 55.22 | 72.69 | 78.65 | 74.68 | 60.09 | 22.31 |
| Galactose-1 | 103.26 | 70.15 | 82.20 | 68.41 | 75.03 | 61.18 | - |
| GlcNAc-2 | 102.67 | 55.32 | 72.47 | 78.44 | 74.68 | 60.09 | 22.31 |
| Galactose-2 | 103.01 | 70.15 | 82.20 | 68.41 | 75.03 | 61.18 | - |
| GlcNAc-3 | 102.98 | 55.32 | 72.47 | 78.44 | 74.68 | 60.09 | 22.31 |
| Galactose-3 | 102.94 | 70.15 | 82.20 | 68.41 | 75.03 | 61.18 | - |
| GlcNAc-4 | 103.13 | 55.87 | n/a | n/a | n/a | 60.64 | 22.31 |

| Linker | 1 | 2 | 3 | 4 | 5 | 6 |
|---|---|---|---|---|---|---|
| H | 3.87, 3.55 | 1.59 – 1.52 (m, 2H) | 1.34 – 1.28 (m, 2H) | 1.49 (p, J = 7.3 Hz, 2H) | 3.12 (t, J = 6.8 Hz, 1H) | 5.12 (s, 2H) |
| C | 70.40 | 28.29 | 22.38 | 28.50 | 40.47 | 66.74 |

HRMS (ESI-MS): m/z calculated for $C_{63}H_{102}N_5O_{38}$ $[M+Na]^+$: 1558.6017; found: 1558.6687.

**Compound 7**

**7** was prepared from **2** (8.0 mg, 10.0 μmol) using the general procedure for the installation of 6-O-sulfate installation of terminal GlcNAc with CHST2. A trace amount (<5%) of di-6-O-sulfate glycan side product **7’** was also found on LC-MS. After P6 and DEAE purification, **7** was obtained as a white solid (7.1 mg, 79%) and **7’** was obtained as a white solid.

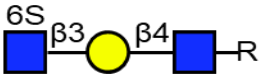

$^{1}H$ (600 MHz, $D_2O$): δ (ppm)

| | H-1 | H-2 | H-3 | H-4 | H-5 | H-6 | NHAc |
|---|---|---|---|---|---|---|---|
| GlcNAc-1 | 4.51 (d, J = 7.4 Hz, 1H) | 3.72 | 3.70 | 3.69 | 3.58 | 3.99 (d, J = 12.2 Hz, 1H)，3.83 | 2.06 – 2.00 (m, 6H) |
| Galactose | 4.47 (d, J = 7.9 Hz, 1H) | 3.60 | 3.73 | 4.20 (d, J = 3.1 Hz, 1H) | 3.72 | 3.77 – 3.73 (m, 2H) | - |
| GlcNAc-2 | 4.70 (d, J = 8.6 Hz, 1H) | 3.80 | 3.59 | 3.54 | 3.67 | 4.34 (dd, J = 11.3 Hz, 1H), 4.23 (dd, J = 11.3, 5.7 Hz, 1H) | 2.06 – 2.00 (m, 6H) |

$^{13}C$ (150 MHz, $D_2O$): δ (ppm)

| | C-1 | C-2 | C-3 | C-4 | C-5 | C-6 | NHAc |
|---|---|---|---|---|---|---|---|
| GlcNAc-1 | 101.08 | 55.01 | 72.49 | 78.70 | 74.75 | 60.15 | 22.15 |
| Galactose | 102.93 | 69.78 | 82.22 | 68.39 | 75.19 | 61.31 | - |
| GlcNAc-2 | 102.94 | 55.68 | 73.66 | 69.69 | 73.66 | 67.19 | 22.15 |

| Linker | 1 | 2 | 3 | 4 | 5 | 6 |
|---|---|---|---|---|---|---|
| H | 3.87,3.57 | 1.61 – 1.52 (m, 2H) | 1.37 – 1.24 (m, 2H) | 1.49 (p, J = 7.3 Hz, 2H) | 3.16 – 3.10 (m, 2H) | 5.12 (s, 2H) |
| C | 70.35 | 28.22 | 22.45 | 28.46 | 40.50 | 66.86 |

HRMS (ESI-MS): m/z calculated for $C_{35}H_{54}N_3O_{21}S$ $[M-H]^-$: 884.2976; found: 884.2737.

**Compound 7’**

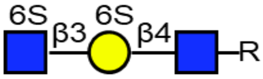

**7’**

| | H-1 | H-2 | H-3 | H-4 | H-5 | H-6 | NHAc |
|---|---|---|---|---|---|---|---|
| GlcNAc | 4.52 | 3.72 | 3.73 | 3.68 | 3.61 | 3.99, 3.82 | 2.07 – 1.98 (m, 6H) |
| Galactose-6S | 4.51 | 3.61 | 3.74 | 4.24 | 3.98 | 4.20 (2H) | - |

| | | | | | | | |
|---|---|---|---|---|---|---|---|
| GlcNAc-6S | 4.72 (d, J = 8.3 Hz, 1H) | 3.79 | 3.57 | 3.55 | 3.67 | 4.34 (d, J = 11.4 Hz, 1H), 4.26 | 2.07 – 1.98 (m, 6H) |

| | C-1 | C-2 | C-3 | C-4 | C-5 | C-6 | NHAc |
|---|---|---|---|---|---|---|---|
| GlcNAc | 100.57 | 55.19 | 72.03 | 79.46 | 74.68 | 60.65 | 22.26 |
| Galactose-6S | 103.11 | 69.59 | 82.27 | 68.81 | 72.95 | 67.92 | - |
| GlcNAc-6S | 102.74 | 55.60 | 73.60 | 69.19 | 73.70 | 67.20 | 22.26 |

| Linker | 1 | 2 | 3 | 4 | 5 | 6 |
|---|---|---|---|---|---|---|
| H | 3.87,3.57 | 1.61 – 1.52 (m, 2H) | 1.35 – 1.28 (m, 2H) | 1.52 – 1.45 (m, 2H) | 3.16 – 3.08 (m, 2H) | 5.12 (s, 2H) |
| C | 70.50 | 28.29 | 22.42 | 28.52 | 40.45 | 66.81 |

HRMS (ESI-MS): m/z calculated for $C_{35}H_{53}N_3O_{24}S_2$ $[M-2H]^{2-}$: 481.6235; found: 481.6257.

#### Compound 8

**8** was prepared from **7** (5.0 mg, 5.6 µmol) using the general procedure for the installation of β1,4-Gal with B4GalT4 to full conversion. After P6 purification, **8** was obtained as a white solid (5.9 mg, quant.).

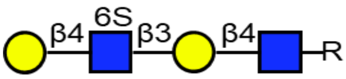

$^1H$ (600 MHz, $D_2O$): δ (ppm)

| | H-1 | H-2 | H-3 | H-4 | H-5 | H-6 | NHAc |
|---|---|---|---|---|---|---|---|
| GlcNAc-1 | 4.51 (d, J = 7.6 Hz, 1H) | 3.72 | 3.70 | 3.70 | 3.58 | 3.99 (dd, J = 12.4, 2.2 Hz, 1H), 3.83 | 2.07 – 2.00 (m, 6H) |
| Galactose-1 | 4.47 (d, J = 7.9 Hz, 1H) | 3.60 | 3.72 | 4.20 (d, J = 3.3 Hz, 1H) | 3.72 | 3.76 (4H) | - |
| GlcNAc-6S | 4.73 (d, J = 8.4 Hz, 1H) | 3.84 | 3.74 | 3.81 | 3.81 | 4.41 (d, J = 10.2 Hz, 1H), 4.32 (dd, J = 10.8, 3.9 Hz, 1H) | 2.07 – 2.00 (m, 6H) |
| Galactose-2 | 4.53 (d, J = 7.9 Hz, 1H) | 3.55 | 3.69 | 3.94 (d, J = 3.4 Hz, 1H) | 3.74 | 3.76 (4H) | - |

$^{13}C$ (150 MHz, $D_2O$): δ (ppm)

| | C-1 | C-2 | C-3 | C-4 | C-5 | C-6 | NHAc |
|---|---|---|---|---|---|---|---|
| GlcNAc-1 | 101.06 | 55.06 | 72.68 | 78.82 | 75.00 | 60.28 | 22.20 |
| Galactose-1 | 102.98 | 70.21 | 82.66 | 68.38 | 75.27 | 61.13 | - |
| GlcNAc-6S | 102.91 | 55.35 | 72.24 | 77.75 | 72.50 | 66.59 | 22.20 |
| Galactose-2 | 102.73 | 71.23 | 72.68 | 68.72 | 75.36 | 61.13 | - |

| Linker | 1 | 2 | 3 | 4 | 5 | 6 |
|---|---|---|---|---|---|---|
| H | 3.88, 3.57 | 1.58 – 1.52 (m, 2H) | 1.35 – 1.25 (m, 2H) | 1.49 (p, J = 7.2 Hz, 2H) | 3.13 (t, J = 6.8 Hz, 1H) | 5.12 (s, 2H) |
| C | 70.37 | 28.25 | 22.44 | 28.47 | 40.46 | 66.78 |

HRMS (ESI-MS): m/z calculated for $C_{41}H_{64}N_3O_{26}S$ $[M-H]^-$: 1046.3504; found: 1046.3083.

#### Compound 9

**9** was prepared from **4** (3.0 mg, 2.5 µmol) using the general procedure for the installation of 6-O-sulfate on terminal GlcNAc with CHST2. After P6 and HILIC HPLC purification, **9** was obtained as a white solid (2.4 mg, 74%).

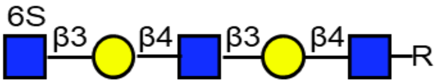

$^1$H (600 MHz, $D_2O$): δ (ppm)

| | H-1 | H-2 | H-3 | H-4 | H-5 | H-6 | NHAc |
|---|---|---|---|---|---|---|---|
| GlcNAc-1 | 4.51 (d, J = 7.5 Hz, 1H) | 3.72 | 3.70 | 3.70 | 3.59 | 3.98, 3.84 | 2.06 – 1.99 (m, 9H) |
| Galactose-1 | 4.46 | 3.59 | 3.74 | 4.16 (d, J = 3.3 Hz, 1H) | 3.72 | 3.76 (4H) | - |
| GlcNAc-2 | 4.71 | 3.80 | 3.74 | 3.74 | 3.59 | 3.98, 3.84 | 2.06 – 1.99 (m, 9H) |
| Galactose-2 | 4.48 | 3.60 | 3.74 | 4.20 (d, J = 3.2 Hz, 1H) | 3.74 | 3.76 (4H) | - |
| GlcNAc-6S | 4.70 | 3.80 | 3.59 | 3.53 | 3.67 | 4.37 – 4.31 (m, 1H), 4.23 (dd, J = 11.2, 5.8 Hz, 1H) | 2.06 – 1.99 (m, 9H) |

$^{13}$C (150 MHz, $D_2O$): δ (ppm)

| | C-1 | C-2 | C-3 | C-4 | C-5 | C-6 | NHAc |
|---|---|---|---|---|---|---|---|
| GlcNAc-1 | 101.16 | 55.27 | 72.73 | 78.75 | 74.69 | 60.05 | 22.21 |
| Galactose-1 | 102.80 | 69.95 | 82.33 | 68.35 | 75.07 | 61.27 | - |
| GlcNAc-2 | 102.88 | 55.40 | 72.33 | 78.55 | 74.87 | 60.05 | 22.21 |
| Galactose-2 | 103.20 | 69.95 | 82.33 | 68.31 | 75.27 | 61.27 | - |
| GlcNAc-6S | 103.36 | 55.40 | 73.64 | 69.75 | 73.60 | 67.26 | 22.21 |

| Linker | 1 | 2 | 3 | 4 | 5 | 6 |
|---|---|---|---|---|---|---|
| H | 3.88, 3.55 | 1.59 – 1.52 (m, 2H) | 1.35 – 1.26 (m, 2H) | 1.49 (p, J = 7.3 Hz, 2H) | 3.17 – 3.10 (m, 2H) | 5.12 (s, 2H) |

| C | 70.44 | 28.21 | 22.33 | 28.37 | 40.30 | 66.76 |
|---|---|---|---|---|---|---|

HRMS (ESI-MS): m/z calculated for $C_{49}H_{77}N_4O_{31}S$ [M-H]$^-$: 1249.4297; found: 1249.3692.

#### Compound 10

**10** was prepared from **9** (2.4 mg, 1.9 μmol) using the general procedure for the installation of β1,4-Gal with B4GalT4 to full conversion. After P6 purification, **10** was obtained as a white solid (2.7 mg, 100%).

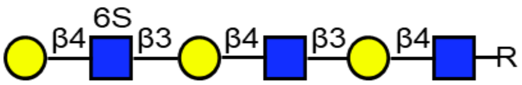

$^1$H (600 MHz, $D_2O$): δ (ppm)

| | H-1 | H-2 | H-3 | H-4 | H-5 | H-6 | NHAc |
|---|---|---|---|---|---|---|---|
| GlcNAc-1 | 4.51 | 3.73 | 3.68 | 3.69 | 3.58 | 3.97, 3.84 | 2.10 – 1.98 (m, 9H) |
| Galactose-1 | 4.46 | 3.60 | 3.72 | 4.16 | 3.72 | 3.76 (6H) | - |
| GlcNAc-2 | 4.70 | 3.80 | 3.73 | 3.73 | 3.58 | 3.97, 3.84 | 2.10 – 1.98 (m, 9H) |
| Galactose-2 | 4.48 | 3.60 | 3.72 | 4.20 | 3.72 | 3.76 (6H) | - |
| GlcNAc-6S | 4.71 | 3.83 | 3.75 | 3.80 | 3.81 | 4.40 (m, 1H), 4.32 (m, 1H) | 2.10 – 1.98 (m, 9H) |
| Galactose-3 | 4.53 | 3.53 | 3.69 | 3.94 | 3.72 | 3.76 (6H) | - |

$^{13}$C (150 MHz, $D_2O$): δ (ppm)

| | C-1 | C-2 | C-3 | C-4 | C-5 | C-6 | NHAc |
|---|---|---|---|---|---|---|---|
| GlcNAc-1 | 101.12 | 55.50 | 72.69 | 78.67 | 74.87 | 60.09 | 22.20 |
| Galactose-1 | 103.16 | 70.02 | 82.39 | 68.49 | 75.18 | 61.33 | - |
| GlcNAc-2 | 102.88 | 55.18 | 72.23 | 78.22 | 74.87 | 60.09 | 22.20 |
| Galactose-2 | 103.03 | 70.02 | 82.39 | 68.42 | 75.18 | 61.33 | - |
| GlcNAc-6S | 103.36 | 55.43 | 72.23 | 77.60 | 72.52 | 66.70 | 22.20 |
| Galactose-3 | 102.66 | 71.18 | 72.60 | 68.86 | 75.18 | 61.33 | - |

| Linker | 1 | 2 | 3 | 4 | 5 | 6 |
|---|---|---|---|---|---|---|
| H | 3.88, 3.55 | 1.59 – 1.52 (m, 2H) | 1.35 – 1.26 (m, 2H) | 1.49 (p, J = 7.3 Hz, 2H) | 3.17 – 3.10 (m, 2H) | 5.12 (s, 2H) |
| C | 70.66 | 28.24 | 22.56 | 28.53 | 40.82 | 67.15 |

HRMS (ESI-MS): m/z calculated for $C_{55}H_{87}N_4O_{36}S$ [M-H]$^-$: 1411.4826; found: 1411.4729.

**Compound 12**

**12** was prepared from **11** (1.0 mg, 1.5 μmol) using the general procedure for the selective installation of terminal α2,6-Neu5Ac using ST6Gal1. After P2 purification, **12** was obtained as a white solid (1.3 mg, 89%).

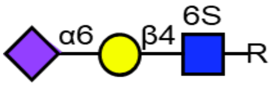

$^{1}H$ (600 MHz, $D_2O$): δ (ppm)

| | H-1 | H-2 | H-3 | H-4 | H-5 | H-6 | H-7 | H-8 | H-9 | NHAc |
|---|---|---|---|---|---|---|---|---|---|---|
| GlcNAc-6S | 4.57 (d, J = 7.8 Hz, 1H) | 3.74 | n/a | 3.66 | 3.84 | 4.45 (dd, J = 11.2, 2.0 Hz, 1H), 4.27 (dd, J = 11.1, 5.7 Hz, 1H) | - | - | - | 2.05 – 2.03 (m, 6H) |
| Galactose | 4.48 (d, J = 7.9 Hz, 1H) | 3.55 | 3.70 | 3.94 (d, J = 3.5 Hz, 1H) | n/a | 4.01, 3.55 | - | - | - | - |
| Sialic acid | - | - | 2.68 (dd, J = 12.5, 4.7 Hz, 1H), 1.73 (t, J = 12.2 Hz, 1H) | 3.68 | 3.81 | n/a | 3.57 | 3.90 | 3.89, 3.66 | 2.05 – 2.03 (m, 6H) |

$^{13}C$ (150 MHz, $D_2O$): δ (ppm)

| | C-1 | C-2 | C-3 | C-4 | C-5 | C-6 | C-7 | C-8 | C-9 | NHAc |
|---|---|---|---|---|---|---|---|---|---|---|
| GlcNAc-6S | 100.99 | 55.05 | n/a | 80.69 | 73.84 | 66.88 | - | - | - | 22.15 |
| Galactose | 103.46 | 71.14 | 72.78 | 68.53 | n/a | 63.42 | - | - | - | - |
| Sialic acid | n/a | n/a | 40.22 | n/a | 51.82 | n/a | 68.65 | 71.86 | 62.93 | 22.15 |

| Linker | 1 | 2 | 3 | 4 | 5 | 6 |
|---|---|---|---|---|---|---|
| H | 3.88, 3.60 | 1.56 (p, J = 6.8 Hz, 2H) | 1.38 – 1.27 (m, 2H) | 1.50 (p, J = 7.2 Hz, 2H) | 3.13 (t, J = 6.7 Hz, 2H) | 5.12 (s, 2H) |
| C | 70.54 | 28.29 | 22.43 | 28.52 | 40.48 | 66.72 |

HRMS (ESI-MS): m/z calculated for $C_{38}H_{58}N_3O_{24}S$ $[M-H]^-$: 972.3136; found: 972.3063.

**Compound 13**

**13** was prepared from **8** (1.0 mg, 1.0 µmol) using the general procedure for the selective installation of terminal α2,6-Neu5Ac using ST6Gal1. After P2 purification, **13** was obtained as a white solid (1.3 mg, quant.).

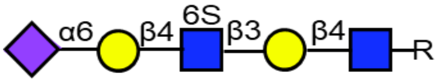

$^{1}H$ (600 MHz, $D_2O$): δ (ppm)

| | H-1 | H-2 | H-3 | H-4 | H-5 | H-6 | H-7 | H-8 | H-9 | NHAc |
|---|---|---|---|---|---|---|---|---|---|---|
| GlcNAc | 4.51 (d, J = 7.8 Hz, 1H) | 3.72 | 3.70 | 3.69 | 3.58 | 3.99, 3.83 | - | - | - | 2.08 – 1.97 (m, 9H) |
| Galactose-1 | 4.47 | 3.60 | 3.74 | 4.21 (d, J = 3.3 Hz, 1H) | n/a | 3.77 (2H) | - | - | - | - |
| GlcNAc-6S | 4.75 | 3.84 | n/a | 3.69 | 3.85 | 4.43 (d, J = 10.7 Hz, 1H), 4.29 (dd, J = 11.1, 5.8 Hz, 1H) | - | - | - | 2.08 – 1.97 (m, 9H) |
| Galactose-2 | 4.48 | 3.54 | 3.69 | 3.94 (d, J = 3.5 Hz, 1H) | n/a | 4.01, 3.55 | - | - | - | - |
| Sialic acid | - | - | 2.68 (dd, J = 12.4, 4.6 Hz, 1H), 1.73 (t, J = 12.2 Hz, 1H) | 3.69 | 3.82 | n/a | 3.57 | 3.90 | 3.89, 3.66 | 2.08 – 1.97 (m, 9H) |

$^{13}C$ (150 MHz, $D_2O$): δ (ppm)

| | C-1 | C-2 | C-3 | C-4 | C-5 | C-6 | C-7 | C-8 | C-9 | NHAc |
|---|---|---|---|---|---|---|---|---|---|---|
| GlcNAc | 101.17 | 54.99 | 72.57 | 79.47 | 74.79 | 60.15 | - | - | - | 22.15 |
| Galactose-1 | 102.55 | 69.85 | 82.53 | 68.30 | n/a | 61.34 | - | - | - | - |
| GlcNAc-6S | 102.72 | 54.87 | n/a | 79.62 | 72.98 | 66.84 | - | - | - | 22.15 |
| Galactose-2 | 103.88 | 71.03 | 72.88 | 68.44 | n/a | 63.42 | - | - | - | - |
| Sialic acid | n/a | n/a | 40.11 | n/a | 51.81 | n/a | 68.65 | 71.86 | 62.93 | 22.15 |

| Linker | 1 | 2 | 3 | 4 | 5 | 6 |
|---|---|---|---|---|---|---|
| H | 3.88, 3.57 | 1.59 – 1.52 (m, 2H) | 1.36 – 1.28 (m, 2H) | 1.49 (p, J = 7.2 Hz, 2H) | 3.13 (t, J = 6.9 Hz, 2H) | 5.12 (s, 2H) |

| | | | | | | |
|---|---|---|---|---|---|---|
| C | 69.95 | 28.34 | 22.44 | 28.52 | 40.53 | 66.77 |

HRMS (ESI-MS): m/z calculated for $C_{52}H_{80}N_4O_{34}S$ $[M-2H]^{2-}$: 668.2193; found: 668.2202.

#### Compound 14

**14** was prepared from **10** (0.9 mg, 0.6 μmol) using the general procedure for the selective installation of terminal α2,6-Neu5Ac using ST6GalL1. After P2 purification, **14** was obtained as a white solid (1.1 mg, quant.).

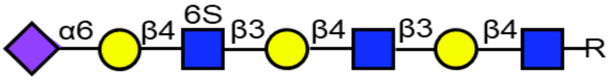

$^1$H (600 MHz, $D_2O$): δ (ppm)

| | H-1 | H-2 | H-3 | H-4 | H-5 | H-6 | H-7 | H-8 | H-9 | NHAc |
|---|---|---|---|---|---|---|---|---|---|---|
| GlcNAc-1 | 4.52 (d, J = 8.1 Hz, 1H) | 3.72 | 3.69 | 3.69 | 3.58 | 3.98, 3.84 | - | - | - | 2.08 – 2.01 (m, 12H) |
| Galactose-1 | 4.46 | 3.59 | 3.73 | 4.17 (d, J = 3.3 Hz, 1H) | n/a | 3.77 (4H) | - | - | - | - |
| GlcNAc-2 | 4.71 (d, J = 8.3 Hz, 1H) | 3.81 | 3.73 | 3.73 | 3.59 | 3.98, 3.84 | - | - | - | 2.08 – 2.01 (m, 12H) |
| Galactose-2 | 4.49 | 3.61 | 3.73 | 4.21 (d, J = 3.3 Hz, 1H) | n/a | 3.77 (4H) | - | - | - | - |
| GlcNAc-6S | 4.75 (d, J = 8.1 Hz, 1H) | 3.84 | n/a | 3.68 | 3.82 | 4.44 (d, J = 10.4 Hz, 1H), 4.29 (dd, J = 11.1, 5.7 Hz, 1H) | - | - | - | 2.08 – 2.01 (m, 12H) |
| Galactose-3 | 4.49 | 3.54 | 3.70 | 3.94 (d, J = 3.5 Hz, 1H) | n/a | 4.01, 3.55 | - | - | - | - |
| Sialic acid | - | - | 2.68 (dd, J = 12.4, 4.6 Hz, 1H), 1.74 (t, J = | 3.66 | 3.83 | n/a | 3.57 | 3.90 | 3.89, 3.66 | 2.08 – 2.01 (m, 12H) |

| | | | |
|---|---|---|---|
| | | | 12.2 Hz, 1H) |

$^{13}C$ (150 MHz, $D_2O$): δ (ppm)

| | C-1 | C-2 | C-3 | C-4 | C-5 | C-6 | C-7 | C-8 | C-9 | NHAc |
|---|---|---|---|---|---|---|---|---|---|---|
| GlcNAc-1 | 101.26 | 55.26 | 72.78 | 78.80 | 75.03 | 60.23 | - | - | - | 22.28 |
| Galactose-1 | 103.00 | 70.14 | 82.61 | 68.50 | n/a | 61.25 | - | - | - | - |
| GlcNAc-2 | 102.88 | 55.12 | 72.54 | 78.65 | 75.05 | 60.23 | - | - | - | 22.28 |
| Galactose-2 | 103.28 | 70.26 | 82.61 | 68.50 | n/a | 61.25 | - | - | - | - |
| GlcNAc-6S | 102.74 | 55.20 | n/a | 80.81 | 72.65 | 67.27 | - | - | - | 22.28 |
| Galactose-3 | 103.40 | 70.99 | 72.70 | 68.65 | n/a | 63.42 | - | - | - | - |
| Sialic acid | n/a | n/a | 40.47 | n/a | 52.22 | n/a | 68.65 | 71.86 | 62.93 | 22.28 |

| Linker | 1 | 2 | 3 | 4 | 5 | 6 |
|---|---|---|---|---|---|---|
| H | 3.87,3.57 | 1.59 – 1.52 (m, 2H) | 1.35 – 1.28 (m, 2H) | 1.50 (p, J = 7.2 Hz, 2H) | 3.13 (t, J = 6.8 Hz, 2H) | 5.12 (s, 2H) |
| C | 70.64 | 28.48 | 22.48 | 28.48 | 40.54 | 66.91 |

HRMS (ESI-MS): m/z calculated for $C_{66}H_{103}N_5O_{44}S$ $[M-2H]^{2-}$: 850.7854; found: 850.7550.

#### Compound 15

**15** was prepared from **10** (2.4 mg, 1.7 µmol) using the general procedure for the installation of α2,3-Neu5Ac with ST3Gal4. After P6 purification, **15** was obtained as a white solid (2.8 mg, 97%).

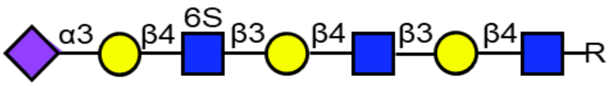

$^{1}H$ (600 MHz, $D_2O$): δ (ppm)

| | H-1 | H-2 | H-3 | H-4 | H-5 | H-6 | H-7 | H-8 | H-9 | NHAc |
|---|---|---|---|---|---|---|---|---|---|---|
| GlcNAc-1 | 4.51 (d, J = 7.6 Hz, 1H) | 3.72 | 3.69 | 3.70 | 3.59 | 3.98, 3.83 | - | - | - | 2.05 – 1.98 (m, 12H) |
| Galactose-1 | 4.46 | 3.60 | 3.73 | 4.16 (d, J = 3.3 Hz, 1H) | 3.72 | 3.76 (4H) | - | - | - | - |
| GlcNAc-2 | 4.71 | 3.81 | 3.73 | 3.74 | 3.59 | 3.98, 3.83 | - | - | - | 2.05 – 1.98 (m, 12H) |
| Galactose-2 | 4.48 | 3.60 | 3.73 | 4.20 (d, J = 3.4 | 3.72 | 3.76 (4H) | - | - | - | - |

| | | | | | | | | | | |
|---|---|---|---|---|---|---|---|---|---|---|
| | | | | Hz, 1H) | | | | | | |
| GlcNAc-6S | 4.72 | 3.84 | 3.74 | 3.81 | 3.81 | 4.41 (d, J = 11.1 Hz, 1H), 4.32 (d, J = 10.3 Hz, 1H) | - | - | - | 2.05 – 1.98 (m, 12H) |
| Galactose-3 | 4.61 (d, J = 7.8 Hz, 1H) | 3.58 | 4.13 (dd, J = 10.0, 3.0 Hz, 1H) | 3.98 | n/a | n/a | - | - | - | - |
| Sialic acid | - | - | 2.76 (dd, J = 12.4, 4.6 Hz, 1H), 1.82 (t, J = 12.1 Hz, 1H) | 3.68 | 3.86 | n/a | 3.57 | 3.90 | 3.89, 3.66 | 2.05 – 1.98 (m, 12H) |

$^{13}C$ (150 MHz, $D_2O$): δ (ppm)

| | C-1 | C-2 | C-3 | C-4 | C-5 | C-6 | C-7 | C-8 | C-9 | NHAc |
|---|---|---|---|---|---|---|---|---|---|---|
| GlcNAc-1 | 101.06 | 55.17 | 72.46 | 78.61 | 74.79 | 60.11 | - | - | - | 22.24 |
| Galactose-1 | 103.54 | 70.09 | 82.23 | 68.38 | 75.09 | 61.16 | - | - | - | - |
| GlcNAc-2 | 102.51 | 55.12 | 72.39 | 78.43 | 74.79 | 60.11 | - | - | - | 22.24 |
| Galactose-2 | 102.66 | 70.09 | 82.23 | 68.33 | 75.09 | 61.16 | - | - | - | - |
| GlcNAc-6S | 103.09 | 55.25 | 72.28 | 77.57 | 72.49 | 66.67 | - | - | - | 22.24 |
| Galactose-3 | 102.19 | 69.66 | 75.51 | 67.69 | n/a | n/a | - | - | - | - |
| Sialic acid | n/a | n/a | 39.82 | n/a | 51.87 | n/a | 68.85 | 71.73 | 62.79 | 22.24 |

| Linker | 1 | 2 | 3 | 4 | 5 | 6 |
|---|---|---|---|---|---|---|
| H | 3.88,3.57 | 1.59 – 1.52 (m, 2H) | 1.35 – 1.28 (m, 2H) | 1.49 (p, J = 7.3 Hz, 2H) | 3.12 (t, J = 6.8 Hz, 2H) | 5.12 (s, 2H) |
| C | 70.40 | 28.31 | 22.48 | 28.56 | 40.39 | 66.86 |

HRMS (ESI-MS): m/z calculated for $C_{66}H_{104}N_5O_{44}S$ $[M-H]^-$: 1702.5780; found: 1702.6213.

**Compound 16**

**16** was prepared from **4** (0.5 mg, 0.40 µmol) using the general procedure for 6-O-sulfate installation of internal galactose with CHST1. It gave a mixture of unreacted **4**, mono-6-O-sulfated glycans (2 isomers with +1 sulfate molecule weight) and di-6-O-sulfated glycan on LC-MS with HILIC column. Additional CHST1 and PAPS were added to drive the reaction further. After P6 and HILIC HPLC purification, **16** was obtained as a white solid (340 µg, 64%). Unreacted **4** and mono-6-O-sulfated glycans were recovered which could be further sulfated by CHST1 and PAPS to get **16**.

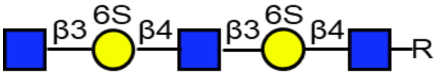

$^1$H (600 MHz, $D_2O$): δ (ppm)

| | H-1 | H-2 | H-3 | H-4 | H-5 | H-6 | NHAc |
|---|---|---|---|---|---|---|---|
| GlcNAc-1 | 4.51 | 3.72 | 3.71 | 3.67 | 3.60 | 3.97, 3.83 | 2.07 – 1.99 (m, 9H) |
| Galactose-1 | 4.51 | 3.59 | 3.75 | 4.23 (d, J = 3.2 Hz, 2H) | 3.99 | 4.22 – 4.18 (m, 4H) | - |
| GlcNAc-2 | 4.71 | 3.80 | 3.74 | 3.73 | 3.60 | 3.97, 3.83 | 2.07 – 1.99 (m, 9H) |
| Galactose-2 | 4.51 | 3.59 | 3.75 | 4.23 (d, J = 3.2 Hz, 2H) | 3.99 | 4.22 – 4.18 (m, 4H) | - |
| GlcNAc-3 | 4.70 | 3.77 | n/a | n/a | 3.46 | 3.90, 3.77 | 2.07 – 1.99 (m, 9H) |

$^{13}$C (150 MHz, $D_2O$): δ (ppm)

| | C-1 | C-2 | C-3 | C-4 | C-5 | C-6 | NHAc |
|---|---|---|---|---|---|---|---|
| GlcNAc-1 | 101.02 | 55.20 | 72.46 | 79.49 | 74.72 | 60.37 | 22.32 |
| Galactose-1 | 103.05 | 69.94 | 82.27 | 68.09 | 72.67 | 67.22 | - |
| GlcNAc-2 | 102.93 | 55.43 | 72.46 | 79.36 | 74.72 | 60.37 | 22.32 |
| Galactose-2 | 103.05 | 69.94 | 82.27 | 68.09 | 72.67 | 67.22 | - |
| GlcNAc-3 | 102.93 | 55.76 | n/a | n/a | n/a | 60.75 | 22.32 |

| Linker | 1 | 2 | 3 | 4 | 5 | 6 |
|---|---|---|---|---|---|---|
| H | 3.87, 3.55 | 1.59 – 1.52 (m, 2H) | 1.35 – 1.26 (m, 2H) | 1.49 (p, J = 7.2 Hz, 2H) | 3.13 (t, J = 7.0 Hz, 2H) | 5.12 (s, 2H) |
| C | 70.44 | 28.21 | 22.33 | 28.37 | 40.30 | 66.76 |

HRMS (ESI-MS): m/z calculated for $C_{49}H_{76}N_4O_{34}S_2$ $[M-2H]^{2-}$: 664.1896; found: 664.1631.

**Compound 17**

**17** was prepared from **5** (1.0 mg, 0.75 μmol) using the general procedure for the 6-O-sulfate installation of internal galactose with CHST1. It gave a mixture of unreacted **5**, mono-6-O-sulfated glycans (2 isomers with +1 sulfate molecule weight) and di-6-O-sulfated glycan on LC-MS with HILIC column. Additional CHST1 and PAPS were added to drive the reaction further. When the reaction was stopped, it was a mixture of a very small amount of mono-6-O-sulfated glycans (2 isomers with +1 sulfate molecule weight) and di-6-O-sulfated glycan. After P6 and HILIC HPLC purification, **17** was obtained as a white solid (1.0 mg, 89%). The mono-6-O-sulfated glycans were recovered which could be further sulfated by CHST1 and PAPS to obtain additional **17**.

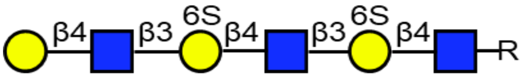

$^{1}H$ (600 MHz, $D_2O$): δ (ppm)

| | H-1 | H-2 | H-3 | H-4 | H-5 | H-6 | NHAc |
|---|---|---|---|---|---|---|---|
| GlcNAc-1 | 4.51 | 3.72 | 3.71 | 3.68 | 3.60 | 3.97, 3.84 | 2.05 – 1.99 (m, 9H) |
| Galactose-1 | 4.51 | 3.59 | 3.75 | 4.23 (d, J = 3.8 Hz, 2H) | 3.98 | 4.22 – 4.18 (m, 4H) | - |
| GlcNAc-2 | 4.71 | 3.81 | 3.73 | 3.70 | 3.60 | 3.97, 3.84 | 2.05 – 1.99 (m, 9H) |
| Galactose-2 | 4.51 | 3.59 | 3.75 | 4.23 (d, J = 3.8 Hz, 2H) | 3.98 | 4.22 – 4.18 (m, 4H) | - |
| GlcNAc-3 | 4.71 | 3.81 | n/a | 3.75 | 3.60 | 3.97, 3.84 | 2.05 – 1.99 (m, 9H) |
| Galactose-3 | 4.49 | 3.55 | 3.68 | 3.93 (d, J = 3.4 Hz, 1H) | 3.75 | 3.76 (2H) | - |

$^{13}C$ (150 MHz, $D_2O$): δ (ppm)

| | C-1 | C-2 | C-3 | C-4 | C-5 | C-6 | NHAc |
|---|---|---|---|---|---|---|---|
| GlcNAc-1 | 101.03 | 55.22 | 72.53 | 79.53 | 74.85 | 60.20 | 22.25 |
| Galactose-1 | 103.19 | 70.02 | 82.48 | 68.19 | 72.59 | 67.27 | - |
| GlcNAc-2 | 103.04 | 55.44 | 72.33 | 79.22 | 74.85 | 60.20 | 22.25 |
| Galactose-2 | 103.19 | 70.02 | 82.48 | 68.19 | 72.59 | 67.27 | - |
| GlcNAc-3 | 103.04 | 55.44 | n/a | 78.56 | 74.85 | 60.20 | 22.25 |
| Galactose-3 | 103.19 | 71.68 | 72.64 | 68.73 | 75.61 | 61.02 | - |

| Linker | 1 | 2 | 3 | 4 | 5 | 6 |
|---|---|---|---|---|---|---|
| H | 3.87, 3.55 | 1.59 – 1.52 (m, 2H) | 1.34 – 1.28 (m, 2H) | 1.49 (p, J = 7.2 Hz, 2H) | 3.12 (t, J = 6.7 Hz, 2H) | 5.12 (s, 2H) |
| C | 70.40 | 28.29 | 22.38 | 28.50 | 40.47 | 66.74 |

HRMS (ESI-MS): m/z calculated for $C_{55}H_{86}N_4O_{39}S_2$ $[M-2H]^{2-}$: 745.2161; found: 745.2121.

#### Compound 18

**18** was prepared from **6** (0.7 mg, 0.46 μmol) using the general procedure for the 6-O-sulfate installation of internal galactose with CHST1. It gave a mixture of unreacted **6**, mono-6-O-sulfated, di-6-O-sulfated and tri-6-O-sulfate glycan on LC-MS using a HILIC column. addtional CHST1 and PAPS were added to drive the reaction further until no further product was formed. When the reaction was stopped, it was a mixture of mono-6-O-sulfate glycans (isomers with +1 sulfate molecule weight) and di 6-O-sulfated glycan (isomers with +2 sulfate molecule weight) and tri-6-O-sulfate glycan. After P6 and HILIC HPLC purification, **18** was obtained as a white solid (0.4 mg, 49%). Mono-6-O-sulfate glycans and di-6-O-sulfate glycans were recovered which could be further sulfated by CHST1 and PAPS to additional **18**.

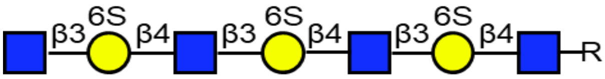

$^1H$ (600 MHz, $D_2O$): δ (ppm)

| | H-1 | H-2 | H-3 | H-4 | H-5 | H-6 | NHAc |
|---|---|---|---|---|---|---|---|
| GlcNAc-1 | 4.51 | 3.72 | 3.71 | 3.67 | 3.61 | 3.97, 3.84 | 2.07 – 2.00 (m, 12H) |
| Galactose-6S-1 | 4.51 | 3.60 | 3.75 | 4.23 (d, J = 3.0 Hz, 3H) | 3.98 | 4.20 (6H) | - |
| GlcNAc-2 | 4.71 | 3.81 | 3.74 | 3.72 | 3.61 | 3.97, 3.84 | 2.07 – 2.00 (m, 12H) |
| Galactose-6S-2 | 4.51 | 3.60 | 3.75 | 4.23 (d, J = 3.0 Hz, 3H) | 3.98 | 4.20 (6H) | - |
| GlcNAc-3 | 4.71 | 3.81 | 3.74 | 3.72 | 3.61 | 3.97, 3.84 | 2.07 – 2.00 (m, 12H) |
| Galactose-6S-3 | 4.51 | 3.60 | 3.75 | 4.23 (d, J = 3.0 Hz, 3H) | 3.98 | 4.20 (6H) | - |
| GlcNAc-4 | 4.71 | 3.77 | n/a | n/a | 3.47 | 3.90, 3.77 | 2.07 – 2.00 (m, 12H) |

$^{13}C$ (150 MHz, $D_2O$): δ (ppm)

| | C-1 | C-2 | C-3 | C-4 | C-5 | C-6 | NHAc |
|---|---|---|---|---|---|---|---|
| GlcNAc-1 | 101.11 | 55.13 | 72.17 | 79.36 | 74.71 | 60.35 | 22.27 |
| Galactose-6S-1 | 103.05 | 69.94 | 82.19 | 68.09 | 72.42 | 67.16 | - |
| GlcNAc-2 | 102.93 | 55.26 | 72.17 | 79.11 | 74.71 | 60.35 | 22.27 |
| Galactose-6S-2 | 103.05 | 69.94 | 82.19 | 68.09 | 72.42 | 67.16 | - |
| GlcNAc-3 | 102.93 | 55.26 | 72.17 | 79.11 | 74.71 | 60.35 | 22.27 |

| | | | | | | | |
|---|---|---|---|---|---|---|---|
| Galactose-6S-3 | 103.05 | 69.94 | 82.19 | 68.09 | 72.42 | 67.16 | - |
| GlcNAc-4 | 102.93 | 55.64 | n/a | n/a | n/a | 60.70 | 22.27 |

| Linker | 1 | 2 | 3 | 4 | 5 | 6 |
|---|---|---|---|---|---|---|
| H | 3.88, 3.56 | 1.60 – 1.52 (m, 2H) | 1.34 – 1.28 (m, 2H) | 1.49 (p, J = 7.3 Hz, 2H) | 3.12 (t, J = 6.9 Hz, 1H) | 5.12 (s, 2H) |
| C | 70.40 | 28.33 | 22.38 | 28.50 | 40.47 | 66.85 |

HRMS (ESI-MS): m/z calculated for $C_{63}H_{98}N_5O_{47}S_3$ $[M-3H]^{3-}$: 590.8203; found: 590.8307.

#### Compound 19

**19** was prepared from **10** (0.5 mg, 0.35 µmol) using the general procedure for the 6-O-sulfate installation of internal galactose with CHST1. When no further product was formed, the reaction was a mixture of unreacted **10** and di-6-O-sulfate glycan. After P6 and HILIC HPLC purification, **19** was obtained as a white solid (0.4 mg, 77%). Unreacted **10** was recovered.

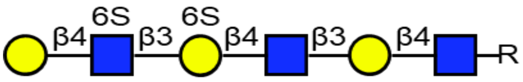

$^1$H (600 MHz, $D_2O$): δ (ppm)

| | H-1 | H-2 | H-3 | H-4 | H-5 | H-6 | NHAc |
|---|---|---|---|---|---|---|---|
| GlcNAc-1 | 4.51 | 3.72 | 3.70 | 3.70 | 3.58 | 3.99, 3.83 | 2.06 – 2.00 (m, 9H) |
| Galactose-1 | 4.46 (d, J = 7.9 Hz, 1H) | 3.60 | 3.73 | 4.16 (d, J = 3.3 Hz, 1H) | 3.73 | 3.76 (4H) | - |
| GlcNAc-2 | 4.71 (d, J = 7.2 Hz, 1H) | 3.81 | 3.73 | 3.71 | 3.63 | 3.97, 3.85 | 2.06 – 2.00 (m, 9H) |
| Galactose-6S | 4.52 | 3.61 | 3.75 | 4.24 | 3.99 | 4.23 – 4.18 (m, 2H) | - |
| GlcNAc-6S | 4.74 | 3.84 | 3.75 | 3.81 | 3.81 | 4.41 – 4.32 (m, 2H) | 2.06 – 2.00 (m, 9H) |
| Galactose-2 | 4.55 (d, J = 7.8 Hz, 1H) | 3.54 | 3.69 | 3.94 (d, J = 3.4 Hz, 1H) | 3.73 | 3.76 (4H) | - |

$^{13}$C (150 MHz, $D_2O$): δ (ppm)

| | C-1 | C-2 | C-3 | C-4 | C-5 | C-6 | NHAc |
|---|---|---|---|---|---|---|---|
| GlcNAc-1 | 101.08 | 55.16 | 72.68 | 78.86 | 75.03 | 60.23 | 22.21 |
| Galactose-1 | 102.84 | 69.97 | 82.32 | 68.51 | 75.27 | 61.01 | - |
| GlcNAc-2 | 102.79 | 55.25 | 72.58 | 79.06 | 74.58 | 60.23 | 22.21 |
| Galactose-6S | 102.39 | 69.97 | 82.64 | 68.36 | 72.89 | 67.98 | - |

| | | | | | | | |
|---|---|---|---|---|---|---|---|
| GlcNAc-6S | 102.89 | 55.25 | 72.36 | 77.46 | 72.48 | 66.42 | 22.21 |
| Galactose-2 | 102.49 | 71.37 | 72.58 | 68.76 | 75.27 | 61.01 | - |

| Linker | 1 | 2 | 3 | 4 | 5 | 6 |
|---|---|---|---|---|---|---|
| H | 3.88, 3.55 | 1.59 – 1.52 (m, 2H) | 1.35 – 1.26 (m, 2H) | 1.49 (p, J = 7.3 Hz, 2H) | 3.12 (t, J = 6.8 Hz, 1H) | 5.12 (s, 2H) |
| C | 70.46 | 28.45 | 22.46 | 28.63 | 40.58 | 66.79 |

HRMS (ESI-MS): m/z calculated for $C_{55}H_{86}N_4O_{39}S_2$ $[M-2H]^{2-}$: 745.2161; found: 745.2117.

**Compound 20**

**20** was prepared from **5** (4.0 mg, 3.0 µmol) using the general procedure for the installation of α2,3-Neu5Ac with ST3Gal4. After P6 purification, **20** was obtained as a white solid (4.6 mg, 95%).

**20** (0.3 mg, 0.18 µmol) was subjected to the general procedure for 6-O-sulfate installation of internal galactose with CHST1. Only a very small amount of a mixture of mono-6-O-sulfated glycans were detected by LC-MS and therefore no purification was attempted.

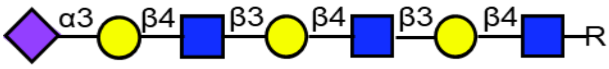

$^1$H (600 MHz, $D_2O$): δ (ppm)

| | H-1 | H-2 | H-3 | H-4 | H-5 | H-6 | H-7 | H-8 | H-9 | NHAc |
|---|---|---|---|---|---|---|---|---|---|---|
| GlcNAc-1 | 4.51 (d, J = 7.6 Hz, 1H) | 3.72 | 3.69 | 3.69 | 3.58 | 3.97, 3.84 | - | - | - | 2.05 – 1.99 (m, 12H) |
| Galactose-1 | 4.46 | 3.59 | 3.73 | 4.16 | 3.72 | 3.76 (4H) | - | - | - | - |
| GlcNAc-2 | 4.70 | 3.80 | 3.73 | 3.74 | 3.58 | 3.97, 3.84 | - | - | - | 2.05 – 1.99 (m, 12H) |
| Galactose-2 | 4.47 | 3.59 | 3.73 | 4.16 | 3.72 | 3.76 (4H) | - | - | - | - |
| GlcNAc-3 | 4.70 | 3.80 | 3.73 | 3.74 | 3.58 | 3.97, 3.84 | - | - | - | 2.05 – 1.99 (m, 12H) |
| Galactose-3 | 4.56 (d, J = 7.9 Hz, 1H) | 3.59 | 4.12 (dd, J = 9.9, 3.1 Hz, 1H) | 3.96 | n/a | n/a | - | - | - | - |
| Sialic acid | - | - | 2.76 (dd, J = 12.5, | 3.70 | 3.85 | n/a | n/a | 3.90 | 3.88, 3.65 | 2.05 – 1.99 |

| | | | | | | | | | | |
|---|---|---|---|---|---|---|---|---|---|---|
| | | | 4.6 Hz, 1H), 1.81 (t, J = 12.2 Hz, 1H) | | | | | | | (m, 12H) |

$^{13}C$ (150 MHz, $D_2O$): δ (ppm)

| | C-1 | C-2 | C-3 | C-4 | C-5 | C-6 | H-7 | H-8 | H-9 | NHAc |
|---|---|---|---|---|---|---|---|---|---|---|
| GlcNAc-1 | 101.06 | 55.18 | 72.42 | 78.62 | 74.80 | 60.13 | - | - | - | 22.20 |
| Galactose-1 | 102.97 | 69.89 | 82.24 | 68.33 | 75.40 | 61.33 | - | - | - | - |
| GlcNAc-2 | 102.87 | 55.31 | 72.32 | 78.33 | 74.80 | 60.13 | - | - | - | 22.20 |
| Galactose-2 | 102.97 | 69.89 | 82.24 | 68.33 | 75.40 | 61.33 | - | - | - | - |
| GlcNAc-3 | 102.87 | 55.31 | 72.32 | 78.33 | 74.80 | 60.13 | - | - | - | 22.20 |
| Galactose-3 | 102.63 | 69.89 | 75.83 | 67.66 | n/a | n/a | - | - | - | - |
| Sialic acid | n/a | n/a | 39.67 | n/a | 51.83 | n/a | n/a | 71.94 | 62.80 | 22.20 |

| Linker | 1 | 2 | 3 | 4 | 5 | 6 |
|---|---|---|---|---|---|---|
| H | 3.87, 3.55 | 1.58 – 1.52 (m, 2H) | 1.35 – 1.25 (m, 2H) | 1.49 (p, J = 7.3 Hz, 2H) | 3.12 (t, J = 6.7 Hz, 2H) | 5.11 (s, 2H) |
| C | 70.40 | 28.36 | 22.67 | 28.53 | 40.44 | 66.73 |

HRMS (ESI-MS): m/z calculated for $C_{66}H_{104}N_5O_{41}$ $[M-H]^-$: 1622.6212; found: 1622.5418.

#### Compound 21

**21** and **22** were prepared from **15** (0.6 mg, 0.35 μmol) using the general procedure for the 6-O-sulfate installation of internal galactose with CHST1. When no further product was observed, the reaction was a mixture of a very small amount of mono-6-*O*-sulfated glycan **15**, di-6-O-sulfate glycan **21** and tri-6-*O*-sulfate glycan **22**. After P6 and HILIC HPLC purification, **21** was obtained as a white solid (0.4 mg, 64%) as well as **22** as a white solid (0.2 mg, 31%). **21** could be further sulfated by CHST1 and PAPS treatment to give additional **22**.

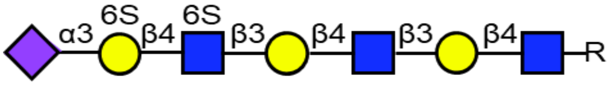

$^1H$ (600 MHz, $D_2O$): δ (ppm)

| | H-1 | H-2 | H-3 | H-4 | H-5 | H-6 | H-7 | H-8 | H-9 | NHAc |
|---|---|---|---|---|---|---|---|---|---|---|
| GlcNAc-1 | 4.51 (d, J = 7.6 Hz, 1H) | 3.72 | 3.70 | 3.69 | 3.58 | 3.97, 3.83 | - | - | - | 2.06 – 1.99 (m, 12H) |
| Galactose-1 | 4.46 | 3.59 | 3.73 | 4.16 | 3.72 | 3.76 (4H) | - | - | - | - |
| GlcNAc-2 | 4.70 | 3.82 | 3.74 | 3.74 | 3.58 | 3.97, 3.83 | - | - | - | 2.06 – 1.99 |

| | | | | | | | | | | |
|---|---|---|---|---|---|---|---|---|---|---|
| | | | | | | | | | | (m, 12H) |
| Galactose-2 | 4.48 | 3.59 | 3.73 | 4.21 (d, J = 3.4 Hz, 1H) | 3.72 | 3.76 (4H) | - | - | - | - |
| GlcNAc-6S | 4.71 | 3.82 | n/a | n/a | 3.86 | 4.45, 4.29 (dd, J = 11.1, 5.8 Hz, 1H) | - | - | - | 2.06 – 1.99 (m, 12H) |
| Galactose-6S | 4.63 (d, J = 7.9 Hz, 1H) | 3.59 | 4.15 | 4.03 (d, J = 3.1 Hz, 1H) | 3.99 | 4.19 (2H) | - | - | - | - |
| Sialic acid | - | - | 2.75 (dd, J = 12.4, 4.7 Hz, 1H), 1.82 (t, J = 12.1 Hz, 1H) | 3.68 | 3.86 | n/a | n/a | 3.91 | 3.89, 3.65 | 2.06 – 1.99 (m, 12H) |

$^{13}$C (150 MHz, $D_2O$): δ (ppm)

| | C-1 | C-2 | C-3 | C-4 | C-5 | C-6 | C-7 | C-8 | C-9 | NHAc |
|---|---|---|---|---|---|---|---|---|---|---|
| GlcNAc-1 | 101.21 | 55.16 | 72.64 | 78.56 | 74.94 | 60.11 | - | - | - | 22.20 |
| Galactose-1 | 103.07 | 69.55 | 82.49 | 68.75 | 75.04 | 61.26 | - | - | - | - |
| GlcNAc-2 | 102.68 | 55.25 | 72.34 | 78.66 | 74.94 | 60.11 | - | - | - | 22.20 |
| Galactose-2 | 102.72 | 69.55 | 82.49 | 68.70 | 75.04 | 61.26 | - | - | - | - |
| GlcNAc-6S | 102.87 | 55.25 | n/a | n/a | 72.74 | 66.85 | - | - | - | 22.20 |
| Galactose-6S | 102.52 | 69.55 | 75.31 | 67.48 | 72.97 | 67.27 | - | - | - | - |
| Sialic acid | n/a | n/a | 39.45 | n/a | 51.81 | n/a | n/a | 71.81 | 62.73 | 22.20 |

| Linker | 1 | 2 | 3 | 4 | 5 | 6 |
|---|---|---|---|---|---|---|
| H | 3.88,3.57 | 1.59 – 1.52 (m, 2H) | 1.35 – 1.27 (m, 2H) | 1.49 (p, J = 7.3 Hz, 2H) | 3.12 (t, J = 6.7 Hz, 2H) | 5.12 (s, 2H) |
| C | 70.25 | 28.34 | 22.56 | 28.63 | 40.47 | 66.89 |

HRMS (ESI-MS): m/z calculated for $C_{66}H_{103}N_5O_{47}S_2$ $[M-2H]^{2-}$: 890.7638; found: 890.7804.

#### Compound 22

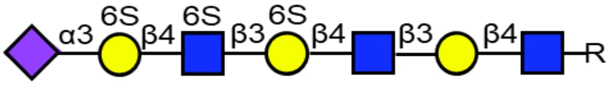

[1]H (600 MHz, $D_2O$): δ (ppm)

| | H-1 | H-2 | H-3 | H-4 | H-5 | H-6 | H-7 | H-8 | H-9 | NHAc |
|---|---|---|---|---|---|---|---|---|---|---|
| GlcNAc-1 | 4.51 | 3.72 | 3.70 | 3.69 | 3.59 | 3.97, 3.83 | - | - | - | 2.07 – 2.00 (m, 12H) |
| Galactose-1 | 4.47 (d, J = 7.8 Hz, 1H) | 3.59 | 3.74 | 4.17 (d, J = 3.4 Hz, 1H) | 3.72 | 3.76 (2H) | - | - | - | - |
| GlcNAc-2 | 4.70 (d, J = 7.7 Hz, 1H) | 3.81 | 3.73 | 3.71 | 3.63 | 3.97, 3.83 | - | - | - | 2.07 – 2.00 (m, 12H) |
| Galactose-6S-1 | 4.52 | 3.59 | 3.74 | 4.25 | 4.00 | 4.20 (4H) | - | - | - | - |
| GlcNAc-6S | 4.73 (d, J = 8.2 Hz, 1H) | 3.83 | n/a | n/a | 3.85 | 4.43 (d, J = 10.5 Hz, 1H), 4.33 (dd, J = 11.4, 5.0 Hz, 1H) | - | - | - | 2.07 – 2.00 (m, 12H) |
| Galactose-6S-2 | 4.64 (d, J = 8.0 Hz, 1H) | 3.59 | 4.16 | 4.04 | 4.00 | 4.20 (4H) | - | - | - | - |
| Sialic acid | - | - | 2.81 – 2.68 (m, 1H), 1.83 (t, J = 12.1 Hz, 1H) | 3.69 | 3.87 | n/a | n/a | 3.92 | 3.90, 3.66 | 2.07 – 2.00 (m, 12H) |

[13]C (150 MHz, $D_2O$): δ (ppm)

| | C-1 | C-2 | C-3 | C-4 | C-5 | C-6 | C-7 | C-8 | C-9 | NHAc |
|---|---|---|---|---|---|---|---|---|---|---|
| GlcNAc-1 | 101.23 | 55.28 | 72.76 | 78.83 | 74.94 | 60.29 | - | - | - | 22.27 |
| Galactose-1 | 103.07 | 69.97 | 82.39 | 68.62 | 75.10 | 61.03 | - | - | - | - |
| GlcNAc-2 | 102.98 | 55.33 | 72.39 | 79.21 | 74.80 | 60.29 | - | - | - | 22.27 |
| Galactose-6S-1 | 102.72 | 69.97 | 82.39 | 68.31 | 72.97 | 67.57 | - | - | - | - |
| GlcNAc-6S | 103.24 | 55.33 | n/a | n/a | 72.72 | 66.81 | - | - | - | 22.27 |

| | | | | | | | | | | |
|---|---|---|---|---|---|---|---|---|---|---|
| Galactose-6S-2 | 102.77 | 69.97 | 75.23 | 67.54 | 72.97 | 67.57 | - | - | - | - |
| Sialic acid | - | - | 39.79 | n/a | 51.87 | n/a | n/a | 71.74 | 62.66 | 22.27 |

| Linker | 1 | 2 | 3 | 4 | 5 | 6 |
|---|---|---|---|---|---|---|
| H | 3.88,3.57 | 1.60 – 1.53 (m, 2H) | 1.36 – 1.31 (m, 2H) | 1.49 (p, J = 7.2 Hz, 2H) | 3.12 (t, J = 6.8 Hz, 2H) | 5.12 (s, 2H) |
| C | 70.40 | 28.31 | 22.53 | 28.60 | 40.54 | 66.89 |

HRMS (ESI-MS): m/z calculated for $C_{66}H_{102}N_5O_{50}S_3$ $[M-3H]^{3-}$: 620.1590; found: 620.1808.

**Compound 23**

Extracted sialylglycopeptide was dissolved in a Tris buffer (100 mM, pH 7.5) containing $CaCl_2$ (5 mM) and $NaN_3$ (1.5 mM) to obtain a final concentration of 20 mg $mL^{-1}$. Pronase E (5-10 wt% in relation to SGP in reaction mixture) was added and the mixture was incubated at 37 °C. Every 2 d additional Pronase E (5-10 wt% in relation to SGP) was added to the reaction mixture. The reaction mixture was lyophilized after LC-MS analysis indicated nearly complete removal of the peptide, usually after 7 to 9 days. The lyophilisates were subjected to size exclusion chromatography, and the carbohydrate containing fractions were combined and further purified by high performance liquid chromatography (HPLC). Then the crude product was dissolved in an aqueous solution of acetic acid (2 M) and kept at 65 °C for 24 h. The solvent was removed in a flow of $N_2$ and the reaction was applied to P2 size exclusion chromatography. Carbohydrate-containing fractions were lyophilized and used without further purification.

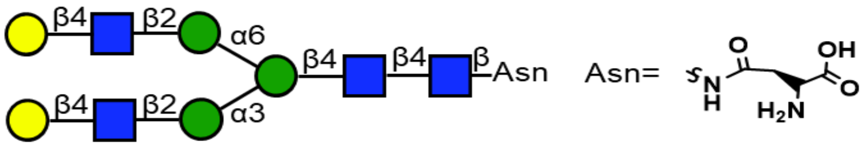

**Compound 24**

**24** was prepared from **23** (52.8 mg, 30.1 µmol) using the general procedure for the selective installation of terminal α2,6-Neu5Ac using ST6Gal1. After P2 purification, **24** was obtained as a white solid (36.8 mg, 60%).

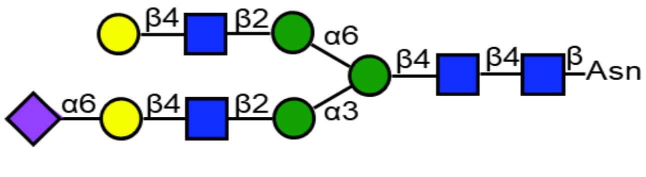

$^1H$ (600 MHz, $D_2O$): δ (ppm)

| | H-1 | H-2 | H-3 | H-4 | H-5 | H-6 | H-7 | H-8 | H-9 | NHAc |
|---|---|---|---|---|---|---|---|---|---|---|
| GlcNAc-1 | 5.08 (d, J = | 3.87 | n/a | 3.67 | 3.60 | 3.77, 3.66 | - | - | - | 2.11 – 2.01 |

| | | | | | | | | | | |
|---|---|---|---|---|---|---|---|---|---|---|
| | 9.8 Hz, 1H) | | | | | | | | | (m, 15H) |
| GlcNAc-2 | 4.63 | 3.80 | n/a | n/a | n/a | n/a | - | - | - | 2.11 – 2.01 (m, 15H) |
| Man-1 | 4.78 | 4.27 (s, 1H) | 3.80 | 3.80 | n/a | 3.97, 3.81 | - | - | - | - |
| Man-2 | 5.15 (s, 1H) | 4.21 (dd, J = 3.4, 1.6 Hz, 1H) | 3.93 | 3.54 | n/a | 3.92, 3.63 | - | - | - | - |
| Man-3 | 4.94 (s, 1H) | 4.12 (dd, J = 3.4, 1.7 Hz, 1H) | 3.91 | 3.51 | 3.63 | 3.93, 3.63 | - | - | - | - |
| GlcNAc-3 | 4.63 | 3.77 | n/a | n/a | 3.58 | 3.99, 3.85 | - | - | - | 2.11 – 2.01 (m, 15H) |
| Galactose-1 | 4.46 (d, J = 7.8 Hz, 1H) | 3.55 | 3.69 | 3.94 | n/a | 4.00, 3.55 | - | - | - | - |
| Sialic acid | - | - | 2.68 (dd, J = 12.4, 4.7 Hz, 1H), 1.73 (t, J = 12.2 Hz, 1H) | 3.68 | 3.82 | n/a | 3.57 | 3.90 | 3.89, 3.66 | 2.11 – 2.01 (m, 15H) |
| GlcNAc-3’ | 4.59 (d, J = 8.2 Hz, 1H) | 3.77 | n/a | n/a | 3.58 | 3.99, 3.85 | - | - | - | 2.11 – 2.01 (m, 15H) |
| Galactose-1’ | 4.49 (d, J = 7.9 Hz, 1H) | 3.56 | 3.69 | 3.94 | n/a | 3.77 (2H) | - | - | - | - |

$^{13}$C (150 MHz, $D_2O$): δ (ppm)

| | C-1 | C-2 | C-3 | C-4 | C-5 | C-6 | C-7 | C-8 | C-9 | NHAc |
|---|---|---|---|---|---|---|---|---|---|---|
| GlcNAc-1 | 78.15 | 53.53 | n/a | 78.79 | 76.30 | 59.92 | - | - | - | 22.27 |
| GlcNAc-2 | 101.30 | 55.03 | n/a | n/a | n/a | n/a | - | - | - | 22.27 |
| Man-1 | 100.47 | 70.22 | 80.65 | 65.58 | n/a | 65.81 | - | - | - | - |
| Man-2 | 99.66 | 76.48 | 69.33 | 67.45 | n/a | 61.72 | - | - | - | - |
| Man-3 | 97.04 | 76.35 | 69.56 | 67.45 | 74.57 | 61.60 | - | - | - | - |
| GlcNAc-3 | 99.48 | 54.84 | n/a | n/a | 74.69 | 60.19 | - | - | - | 22.27 |

| | | | | | | | | | | |
|---|---|---|---|---|---|---|---|---|---|---|
| Galactose-1 | 103.58 | 70.94 | 72.61 | 68.45 | n/a | 63.47 | - | - | - | - |
| Sialic acid | n/a | n/a | 40.10 | 68.33 | 51.88 | n/a | 68.56 | 71.74 | 62.56 | 22.27 |
| GlcNAc-3’ | 99.45 | 54.84 | n/a | n/a | 74.69 | 60.19 | - | - | - | 22.27 |
| Galactose-1’ | 103.08 | 70.94 | 72.61 | 68.45 | n/a | 60.78 | - | - | - | - |

| ASN | βCH2-Asn | αCH-Asn |
|---|---|---|
| H | 2.95 (dd, J = 17.2, 4.3 Hz, 1H), 2.87 (dd, J = 17.2, 7.0 Hz, 1H) | 4.01 |
| C | 35.00 | n/a |

HRMS (ESI-MS): m/z calculated for $C_{77}H_{125}N_7O_{56}$ $[M-2H]^{2-}$: 1021.8580; found: 1021.8320.

#### Compound 25

**25** was prepared from **24** (5.0 mg, 2.4 μmol) using the general procedure for the installation of β1,3-GlcNAc with B3GnT2. After P2 purification, **25** was obtained as a white solid (4.4 mg, 82%).

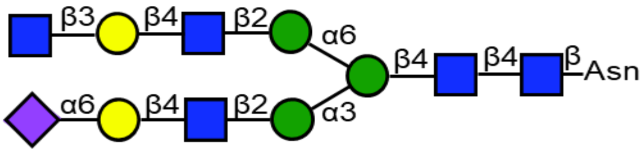

$^1$H (600 MHz, $D_2O$): δ (ppm)

| | H-1 | H-2 | H-3 | H-4 | H-5 | H-6 | H-7 | H-8 | H-9 | NHAc |
|---|---|---|---|---|---|---|---|---|---|---|
| GlcNAc-1 | 5.08 | 3.87 | n/a | 3.67 | 3.60 | 3.77, 3.65 | - | - | - | 2.12 – 1.99 (m, 18H) |
| GlcNAc-2 | 4.63 | 3.80 | n/a | n/a | n/a | n/a | - | - | - | 2.12 – 1.99 (m, 18H) |
| Man-1 | 4.78 | 4.26 (s, 1H) | 3.80 | 3.80 | n/a | 3.97, 3.81 | - | - | - | - |
| Man-2 | 5.15 (s, 1H) | 4.22 – 4.20 (m, 1H) | 3.93 | 3.54 | n/a | 3.92, 3.63 | - | - | - | - |
| Man-3 | 4.94 (s, 1H) | 4.13 – 4.09 (m, 1H) | 3.91 | 3.51 | 3.63 | 3.93, 3.63 | - | - | - | - |
| GlcNAc-3 | 4.63 | 3.77 | n/a | n/a | 3.58 | 3.99, 3.85 | - | - | - | 2.12 – 1.99 |

| | | | | | | | | | | |
|---|---|---|---|---|---|---|---|---|---|---|
| | | | | | | | | | | (m, 18H) |
| Galactose-1 | 4.46 | 3.55 | 3.69 | 3.94 | n/a | 4.01, 3.56 | - | - | - | - |
| Sialic acid | - | - | 2.68 (dd, J = 12.5, 4.6 Hz, 1H), 1.73 (t, J = 12.1 Hz, 1H) | 3.70 | 3.82 | n/a | 3.57 | 3.90 | 3.89, 3.66 | 2.12 – 1.99 (m, 18H) |
| GlcNAc-3’ | 4.59 (d, J = 8.2 Hz, 1H) | 3.77 | n/a | n/a | 3.58 | 3.99, 3.85 | - | - | - | 2.12 – 1.99 (m, 18H) |
| Galactose-1’ | 4.47 | 3.60 | 3.74 | 4.16 (d, J = 3.3 Hz, 1H) | n/a | 3.77 (2H) | - | - | - | - |
| GlcNAc-4’ | 4.70 (d, J = 8.5 Hz, 1H) | 3.78 | n/a | n/a | 3.46 | 3.90, 3.77 | - | - | - | 2.12 – 1.99 (m, 18H) |

$^{13}$C (150 MHz, $D_2O$): δ (ppm)

| | C-1 | C-2 | C-3 | C-4 | C-5 | C-6 | C-7 | C-8 | C-9 | NHAc |
|---|---|---|---|---|---|---|---|---|---|---|
| GlcNAc-1 | 78.09 | 53.53 | n/a | 78.73 | 76.25 | 59.92 | - | - | - | 22.14 |
| GlcNAc-2 | 101.27 | 54.94 | n/a | n/a | n/a | n/a | - | - | - | 22.14 |
| Man-1 | 100.41 | 70.13 | 80.55 | 65.57 | n/a | 65.80 | - | - | - | - |
| Man-2 | 99.49 | 76.40 | 69.43 | 67.39 | n/a | 61.60 | - | - | - | - |
| Man-3 | 96.92 | 76.30 | 69.43 | 67.28 | 74.32 | 61.49 | - | - | - | - |
| GlcNAc-3 | 99.56 | 54.84 | n/a | n/a | 74.39 | 60.11 | - | - | - | 22.14 |
| Galactose-1 | 103.47 | 70.91 | 72.61 | 68.45 | n/a | 63.41 | - | - | - | - |
| Sialic acid | n/a | n/a | 39.95 | n/a | 51.88 | n/a | 68.44 | 71.76 | 62.49 | 22.14 |
| GlcNAc-3’ | 99.30 | 54.84 | n/a | n/a | 74.39 | 60.11 | - | - | - | 22.14 |
| Galactose-1’ | 103.08 | 70.13 | 82.03 | 68.29 | n/a | 60.73 | - | - | - | - |
| GlcNAc-4’ | 102.81 | 54.89 | n/a | n/a | n/a | 60.15 | - | - | - | 22.14 |

HRMS (ESI-MS): m/z calculated for $C_{85}H_{138}N_8O_{61}$ $[M-2H]^{2-}$: 1123.3976; found: 1123.3771.

**Compound 26**

**26** was prepared from **25** (4.4 mg, 2.0 μmol) using the general procedure for the installation of β1,4-Gal with B4GalT1 to full conversion. After P2 purification, **26** was obtained as a white solid (4.7 mg, quant.).

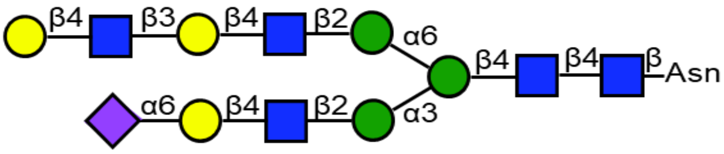

[1]H (600 MHz, $D_2O$): δ (ppm)

| | H-1 | H-2 | H-3 | H-4 | H-5 | H-6 | H-7 | H-8 | H-9 | NHAc |
|---|---|---|---|---|---|---|---|---|---|---|
| GlcNAc-1 | 5.08 (d, J = 9.7 Hz, 1H) | 3.88 | n/a | 3.67 | 3.60 | 3.77, 3.66 | - | - | - | 2.11 – 2.01 (m, 18H) |
| GlcNAc-2 | 4.63 | 3.80 | n/a | n/a | n/a | n/a | - | - | - | 2.11 – 2.01 (m, 18H) |
| Man-1 | 4.78 | 4.27 (s, 1H) | 3.80 | 3.80 | n/a | 3.97, 3.81 | - | - | - | - |
| Man-2 | 5.15 (s, 1H) | 4.21 (dd, J = 3.4, 1.6 Hz, 1H) | 3.93 | 3.54 | n/a | 3.92, 3.63 | - | - | - | - |
| Man-3 | 4.94 (s, 1H) | 4.12 (dd, J = 3.4, 1.7 Hz, 1H) | 3.91 | 3.51 | 3.63 | 3.93, 3.63 | - | - | - | - |
| GlcNAc-3 | 4.63 | 3.77 | n/a | n/a | 3.58 | 3.99, 3.85 | - | - | - | 2.11 – 2.01 (m, 18H) |
| Galactose-1 | 4.46 | 3.55 | 3.69 | 3.94 | n/a | 4.00, 3.55 | - | - | - | - |
| Sialic acid | - | - | 2.68 (dd, J = 12.4, 4.7 Hz, 1H), 1.73 (t, J = 12.1 Hz, 1H) | 3.70 | 3.83 | n/a | 3.57 | 3.90 | 3.89, 3.66 | 2.11 – 2.01 (m, 18H) |
| GlcNAc-3' | 4.59 (d, J = 8.1 Hz, 1H) | 3.77 | n/a | n/a | 3.58 | 3.99, 3.85 | - | - | - | 2.11 – 2.01 (m, 18H) |
| Galactose-1' | 4.47 | 3.61 | 3.74 | 4.17 (d, J = | n/a | n/a | - | - | - | - |

| | | | | | | | | | | |
|---|---|---|---|---|---|---|---|---|---|---|
| | | | | 2.8 Hz, 1H) | | | | | | |
| GlcNAc-4’ | 4.72 (d, J = 8.4 Hz, 1H) | 3.83 | 3.73 | 3.74 | 3.59 | 3.98, 3.84 | - | - | - | 2.11 – 2.01 (m, 18H) |
| Galactose-2’ | 4.49 (d, J = 8.0 Hz, 1H) | 3.56 | 3.69 | 3.94 | n/a | n/a | - | - | - | - |

$^{13}$C (150 MHz, $D_2O$): δ (ppm)

| | C-1 | C-2 | C-3 | C-4 | C-5 | C-6 | C-7 | C-8 | C-9 | NHAc |
|---|---|---|---|---|---|---|---|---|---|---|
| GlcNAc-1 | 78.15 | 53.70 | n/a | 78.68 | 76.42 | 59.93 | - | - | - | 22.26 |
| GlcNAc-2 | 101.39 | 55.03 | n/a | n/a | n/a | n/a | - | - | - | 22.26 |
| Man-1 | 100.47 | 70.22 | 80.65 | 65.58 | n/a | 65.81 | - | - | - | - |
| Man-2 | 99.66 | 76.48 | 69.33 | 67.45 | n/a | 61.72 | - | - | - | - |
| Man-3 | 97.04 | 76.35 | 69.56 | 67.45 | 74.57 | 61.60 | - | - | - | - |
| GlcNAc-3 | 99.56 | 54.84 | n/a | n/a | 74.69 | 60.19 | - | - | - | 22.26 |
| Galactose-1 | 103.58 | 70.98 | 72.54 | 68.57 | n/a | 63.47 | - | - | - | - |
| Sialic acid | n/a | n/a | 40.10 | n/a | 51.88 | n/a | 68.42 | 71.74 | 62.56 | 22.26 |
| GlcNAc-3’ | 99.41 | 54.84 | n/a | n/a | 74.69 | 60.19 | - | - | - | 22.26 |
| Galactose-1’ | 102.98 | 70.27 | 82.26 | 68.26 | n/a | n/a | - | - | - | - |
| GlcNAc-4’ | 102.70 | 55.09 | 72.54 | 78.54 | 74.82 | 60.07 | - | - | - | 22.26 |
| Galactose-2’ | 103.06 | 71.24 | 72.54 | 68.57 | n/a | n/a | - | - | - | - |

HRMS (ESI-MS): m/z calculated for $C_{91}H_{148}N_8O_{66}$ $[M-2H]^{2-}$: 1204.4241; found: 1204.4294.

#### Compound 27

**27** was prepared from **26** (4.7 mg, 2.0 µmol) using the general procedure for the installation of β1,3-GlcNAc with B3GnT2. After P2 purification, **27** was obtained as a white solid (4.6 mg, 88%).

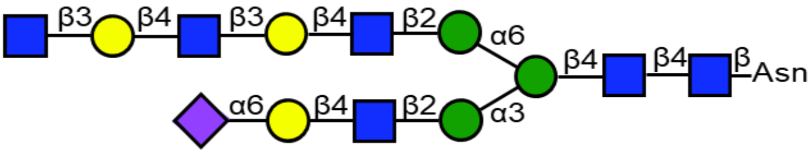

$^{1}$H (600 MHz, $D_2O$): δ (ppm)

| | H-1 | H-2 | H-3 | H-4 | H-5 | H-6 | H-7 | H-8 | H-9 | NHAc |
|---|---|---|---|---|---|---|---|---|---|---|
| GlcNAc-1 | 5.08 (d, J = 9.6 Hz, 1H) | 3.88 | n/a | 3.67 | 3.60 | 3.77, 3.66 | - | - | - | 2.10 – 2.01 (m, 21H) |
| GlcNAc-2 | 4.62 | 3.80 | n/a | n/a | n/a | n/a | - | - | - | 2.10 – 2.01 |

| | | | | | | | | | | |
|---|---|---|---|---|---|---|---|---|---|---|
| | | | | | | | | | | (m, 21H) |
| Man-1 | 4.78 | 4.26 (s, 1H) | 3.80 | 3.80 | n/a | 3.97, 3.81 | - | - | - | - |
| Man-2 | 5.15 (s, 1H) | 4.21 (d, J = 2.9 Hz, 1H) | 3.93 | 3.54 | n/a | 3.92, 3.63 | - | - | - | - |
| Man-3 | 4.94 (s, 1H) | 4.12 (d, J = 3.6 Hz, 1H) | 3.91 | 3.51 | 3.63 | 3.93, 3.63 | - | - | - | - |
| GlcNAc-3 | 4.63 | 3.77 | n/a | n/a | 3.58 | 3.99, 3.85 | - | - | - | 2.10 – 2.01 (m, 21H) |
| Galactose-1 | 4.46 | 3.55 | 3.69 | 3.94 | n/a | 4.00, 3.55 | - | - | - | - |
| Sialic acid | - | - | 2.68 (dd, J = 12.5, 4.6 Hz, 1H), 1.73 (t, J = 12.2 Hz, 1H) | 3.70 | 3.83 | n/a | 3.57 | 3.90 | 3.89, 3.66 | 2.10 – 2.01 (m, 21H) |
| GlcNAc-3’ | 4.59 (d, J = 8.2 Hz, 1H) | 3.77 | n/a | n/a | 3.58 | 3.99, 3.85 | - | - | - | 2.10 – 2.01 (m, 21H) |
| Galactose-1’ | 4.47 | 3.60 | 3.73 | 4.17 | n/a | 3.77 (4H) | - | - | - | - |
| GlcNAc-4’ | 4.71 (d, J = 8.5 Hz, 1H) | 3.82 | 3.73 | 3.73 | 3.59 | 3.98, 3.85 | - | - | - | 2.10 – 2.01 (m, 21H) |
| Galactose-2’ | 4.48 | 3.60 | 3.73 | 4.17 | n/a | 3.77 (4H) | - | - | - | - |
| GlcNAc-5’ | 4.69 (d, J = 8.5 Hz, 1H) | 3.77 | n/a | n/a | 3.46 | 3.90, 3.77 | - | - | - | 2.10 – 2.01 (m, 21H) |

$^{13}C$ (150 MHz, $D_2O$): δ (ppm)

| | C-1 | C-2 | C-3 | C-4 | C-5 | C-6 | C-7 | C-8 | C-9 | NHAc |
|---|---|---|---|---|---|---|---|---|---|---|
| GlcNAc-1 | 78.15 | 53.70 | n/a | 78.79 | 76.30 | 59.92 | - | - | - | 22.29 |
| GlcNAc-2 | 101.39 | 55.00 | n/a | n/a | n/a | n/a | - | - | - | 22.29 |
| Man-1 | 100.47 | 70.22 | 80.65 | 65.58 | n/a | 65.81 | - | - | - | - |
| Man-2 | 99.66 | 76.48 | 69.33 | 67.45 | n/a | 61.72 | - | - | - | - |
| Man-3 | 97.04 | 76.35 | 69.56 | 67.45 | 74.57 | 61.60 | - | - | - | - |
| GlcNAc-3 | 99.56 | 55.10 | n/a | n/a | 74.69 | 60.19 | - | - | - | 22.29 |
| Galactose-1 | 103.58 | 70.94 | 72.58 | 68.43 | n/a | 63.47 | - | - | - | - |
| Sialic acid | n/a | n/a | 40.10 | n/a | 51.88 | n/a | 68.56 | 71.74 | 62.56 | 22.29 |
| GlcNAc-3’ | 99.41 | 55.10 | n/a | n/a | 74.69 | 60.19 | - | - | - | 22.29 |

| | | | | | | | | | | |
|---|---|---|---|---|---|---|---|---|---|---|
| Galactose-1' | 103.11 | 70.07 | 82.18 | 68.34 | n/a | 60.81 | - | - | - | - |
| GlcNAc-4' | 102.87 | 55.09 | 72.50 | 78.47 | 74.65 | 60.11 | - | - | - | 22.29 |
| Galactose-2' | 103.24 | 70.07 | 82.09 | 68.34 | n/a | 60.81 | - | - | - | - |
| GlcNAc-5' | 102.87 | 55.00 | n/a | n/a | n/a | 60.47 | - | - | - | 22.29 |

HRMS (ESI-MS): m/z calculated for $C_{99}H_{161}N_9O_{71}$ $[M-2H]^{2-}$: 1306.4654; found: 1306.4971.

#### Compound 28

**28** was prepared from **27** (2.2 mg, 0.8 µmol) using the general procedure for the installation of 6-O-sulfate installation of terminal GlcNAc with CHST2. After P6 and HILIC HPLC purification, **28** was obtained as a white solid (1.9 mg, 84%).

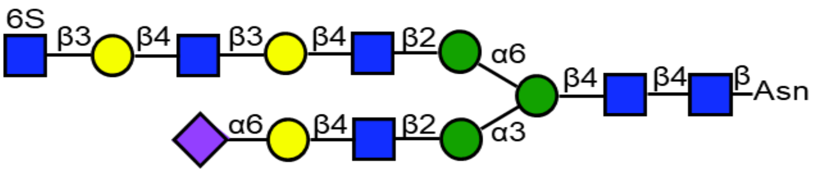

$^1H$ (600 MHz, $D_2O$): δ (ppm)

| | H-1 | H-2 | H-3 | H-4 | H-5 | H-6 | H-7 | H-8 | H-9 | NHAc |
|---|---|---|---|---|---|---|---|---|---|---|
| GlcNAc-1 | 5.08 (d, J = 9.8 Hz, 1H) | 3.87 | n/a | 3.67 | 3.60 | 3.77, 3.66 | - | - | - | 2.11 – 2.00 (m, 21H) |
| GlcNAc-2 | 4.62 | 3.80 | n/a | n/a | n/a | n/a | - | - | - | 2.11 – 2.00 (m, 21H) |
| Man-1 | 4.78 | 4.26 | 3.80 | 3.80 | n/a | 3.97, 3.81 | - | - | - | - |
| Man-2 | 5.15 (s, 1H) | 4.20 | 3.93 | 3.54 | n/a | 3.92, 3.63 | - | - | - | - |
| Man-3 | 4.94 (s, 1H) | 4.11 | 3.91 | 3.51 | 3.63 | 3.93, 3.63 | - | - | - | - |
| GlcNAc-3 | 4.61 | 3.77 | n/a | n/a | 3.58 | 3.99, 3.85 | - | - | - | 2.11 – 2.00 (m, 21H) |
| Galactose-1 | 4.46 (d, J = 7.9 Hz, 1H) | 3.55 | 3.69 | 3.94 | n/a | 4.00, 3.55 | - | - | - | - |
| Sialic acid | - | - | 2.68 (dd, J = 12.4, 4.7 Hz, 1H), 1.73 (t, J = 12.2 | 3.68 | 3.82 | n/a | 3.57 | 3.90 | 3.89, 3.66 | 2.11 – 2.00 (m, 21H) |

| | | | | | | | | | | |
|---|---|---|---|---|---|---|---|---|---|---|
| | | | Hz, 1H) | | | | | | | |
| GlcNAc-3’ | 4.58 | 3.77 | n/a | n/a | 3.58 | 3.99, 3.85 | - | - | - | 2.11 – 2.00 (m, 21H) |
| Galactose-1’ | 4.46 | 3.61 | 3.72 | 4.17 | n/a | 3.76 (4H) | - | - | - | - |
| GlcNAc-4’ | 4.70 | 3.82 | 3.73 | 3.72 | 3.59 | 3.97, 3.84 | - | - | - | 2.11 – 2.00 (m, 21H) |
| Galactose-2’ | 4.48 | 3.61 | 3.72 | 4.19 | n/a | 3.76 (4H) | - | - | - | - |
| GlcNAc-5’ | 4.69 | 3.81 | 3.58 | 3.52 | 3.67 | 4.34, 4.23 | - | - | - | 2.11 – 2.00 (m, 21H) |

$^{13}$C (150 MHz, $D_2O$): δ (ppm)

| | C-1 | C-2 | C-3 | C-4 | C-5 | C-6 | C-7 | C-8 | C-9 | NHAc |
|---|---|---|---|---|---|---|---|---|---|---|
| GlcNAc-1 | n/a | 53.53 | n/a | n/a | n/a | 59.92 | - | - | - | 22.28 |
| GlcNAc-2 | 101.30 | 55.03 | n/a | n/a | n/a | n/a | - | - | - | 22.28 |
| Man-1 | 100.47 | 70.22 | 80.65 | 65.58 | n/a | 65.81 | - | - | - | - |
| Man-2 | 99.66 | 76.48 | 69.33 | 67.45 | n/a | 61.72 | - | - | - | - |
| Man-3 | 97.04 | 76.35 | 69.56 | 67.45 | 74.57 | 61.60 | - | - | - | - |
| GlcNAc-3 | 99.48 | 54.84 | n/a | n/a | 74.69 | 60.19 | - | - | - | 22.28 |
| Galactose-1 | 103.58 | 70.94 | 72.61 | 68.45 | n/a | 63.47 | - | - | - | - |
| Sialic acid | n/a | n/a | 40.10 | 68.33 | 51.88 | n/a | 68.56 | 71.74 | 62.56 | 22.28 |
| GlcNAc-3’ | 99.41 | 55.10 | n/a | n/a | 74.69 | 60.19 | - | - | - | 22.28 |
| Galactose-1’ | 103.11 | 70.07 | 82.40 | 68.34 | n/a | 60.99 | - | - | - | - |
| GlcNAc-4’ | 102.87 | 55.09 | 72.24 | 78.45 | 74.51 | 60.09 | - | - | - | 22.28 |
| Galactose-2’ | 103.10 | 70.07 | 82.40 | 68.39 | n/a | 60.99 | - | - | - | - |
| GlcNAc-5’ | 102.87 | 55.00 | 73.58 | 69.82 | 73.50 | 67.24 | - | - | - | 22.28 |

HRMS (ESI-MS): m/z calculated for $C_{99}H_{161}N_9O_{74}S$ $[M-2H]^{2-}$: 1346.4438; found: 1346.4790.

#### Compound 29

**29** was prepared from **28** (1.9 mg, 0.7 µmol) using the general procedure for the installation of β1,4-Gal with B4GalT4 to full conversion. After P6 purification, **29** was obtained as a white solid (1.8 mg, 89%).

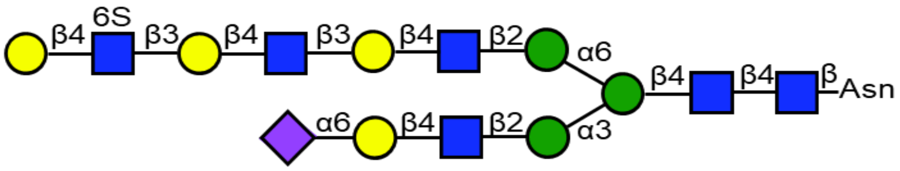

$^{1}H$ (600 MHz, $D_2O$): δ (ppm)

| | H-1 | H-2 | H-3 | H-4 | H-5 | H-6 | H-7 | H-8 | H-9 | NHAc |
|---|---|---|---|---|---|---|---|---|---|---|
| GlcNAc-1 | 5.08 (d, J = 9.7 Hz, 1H) | 3.87 | n/a | 3.67 | 3.60 | 3.77, 3.66 | - | - | - | 2.10 – 2.01 (m, 21H) |
| GlcNAc-2 | 4.62 | 3.80 | n/a | n/a | n/a | n/a | - | - | - | 2.10 – 2.01 (m, 21H) |
| Man-1 | 4.78 | 4.26 (s, 1H) | 3.80 | 3.80 | n/a | 3.97, 3.81 | - | - | - | - |
| Man-2 | 5.15 (s, 1H) | 4.21 | 3.93 | 3.54 | n/a | 3.92, 3.63 | - | - | - | - |
| Man-3 | 4.94 (s, 1H) | 4.12 (d, J = 3.6 Hz, 1H) | 3.91 | 3.51 | 3.63 | 3.93, 3.63 | - | - | - | - |
| GlcNAc-3 | 4.62 | 3.77 | n/a | n/a | 3.58 | 3.99, 3.85 | - | - | - | 2.10 – 2.01 (m, 21H) |
| Galactose-1 | 4.45 | 3.55 | 3.70 | 3.93 | n/a | 4.00, 3.55 | - | - | - | - |
| Sialic acid | - | - | 2.68 (dd, J = 12.4, 4.6 Hz, 1H), 1.73 (t, J = 12.2 Hz, 1H) | 3.68 | 3.82 | n/a | 3.57 | 3.90 | 3.89, 3.66 | 2.10 – 2.01 (m, 21H) |
| GlcNAc-3’ | 4.59 | 3.77 | n/a | n/a | 3.58 | 3.99, 3.85 | - | - | - | 2.10 – 2.01 (m, 21H) |
| Galactose-1’ | 4.47 | 3.60 | 3.73 | 4.17 (d, J = 3.2 Hz, 1H) | n/a | n/a | - | - | - | - |
| GlcNAc-4’ | 4.71 | 3.82 | 3.72 | 3.73 | 3.59 | 3.98, 3.83 | - | - | - | 2.10 – 2.01 (m, 21H) |
| Galactose-2’ | 4.48 | 3.60 | 3.73 | 4.19 | n/a | n/a | - | - | - | - |
| GlcNAc-5’ | 4.73 | 3.85 | n/a | 3.80 | 3.81 | 4.41 (d, J = 10.7 Hz, 1H), 4.32 (dd, J = 10.7, 3.4 Hz, 1H) | - | - | - | 2.10 – 2.01 (m, 21H) |

| Galactose-3' | 4.53 (d, J = 7.8 Hz, 1H) | 3.54 | 3.69 | 3.94 | n/a | n/a | - | - | - | - |
|---|---|---|---|---|---|---|---|---|---|---|

$^{13}$C (150 MHz, $D_2O$): δ (ppm)

| | C-1 | C-2 | C-3 | C-4 | C-5 | C-6 | C-7 | C-8 | C-9 | NHAc |
|---|---|---|---|---|---|---|---|---|---|---|
| GlcNAc-1 | 78.15 | 53.61 | n/a | 78.79 | 76.30 | 59.92 | - | - | - | 22.26 |
| GlcNAc-2 | 99.56 | 54.91 | n/a | n/a | n/a | n/a | - | - | - | 22.26 |
| Man-1 | 100.47 | 70.26 | 80.65 | 65.58 | n/a | 65.81 | - | - | - | - |
| Man-2 | 99.58 | 76.42 | 69.33 | 67.45 | n/a | 61.72 | - | - | - | - |
| Man-3 | 97.04 | 76.35 | 69.56 | 67.45 | 74.57 | 61.60 | - | - | - | - |
| GlcNAc-3 | 101.39 | 54.98 | n/a | n/a | 74.69 | 60.19 | - | - | - | 22.26 |
| Galactose-1 | 103.46 | 71.00 | 72.42 | 68.56 | n/a | 63.47 | - | - | - | - |
| Sialic acid | n/a | n/a | 40.10 | n/a | 51.88 | n/a | 68.56 | 71.74 | 62.56 | 22.26 |
| GlcNAc-3' | 99.47 | 55.06 | n/a | n/a | 74.82 | 60.09 | - | - | - | 22.26 |
| Galactose-1' | 103.11 | 71.18 | 82.34 | 68.33 | n/a | n/a | - | - | - | - |
| GlcNAc-4' | 102.87 | 55.21 | 72.51 | 78.52 | 74.82 | 60.09 | - | - | - | 22.26 |
| Galactose-2' | 103.10 | 71.18 | 82.34 | 68.33 | n/a | n/a | - | - | - | - |
| GlcNAc-5' | 102.87 | 55.21 | n/a | 77.65 | 72.42 | 66.56 | - | - | - | 22.26 |
| Galactose-3' | 102.63 | 71.00 | 72.51 | 68.56 | n/a | n/a | - | - | - | - |

HRMS (ESI-MS): m/z calculated for $C_{105}H_{171}N_9O_{79}S$ $[M-2H]^{2-}$: 1427.4702; found: 1427.5057.

#### Compound 30

**30** was prepared from **29** (1.2 mg, 0.42 µmol) using the general procedure for the selective installation of terminal α2,6-Neu5Ac using ST6Gal1. After P6 purification, **30** was obtained as a white solid (0.8 mg, 61%).

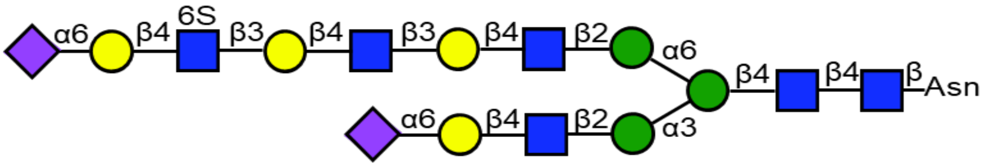

$^{1}$H (600 MHz, $D_2O$): δ (ppm)

| | H-1 | H-2 | H-3 | H-4 | H-5 | H-6 | H-7 | H-8 | H-9 | NHAc |
|---|---|---|---|---|---|---|---|---|---|---|
| GlcNAc-1 | 5.08 (d, J = 9.7 Hz, 1H) | 3.88 | n/a | 3.67 | 3.60 | 3.77, 3.66 | - | - | - | 2.11 – 2.00 (m, 24H) |
| GlcNAc-2 | 4.63 | 3.80 | n/a | n/a | n/a | n/a | - | - | - | 2.11 – 2.00 (m, 24H) |
| Man-1 | 4.78 | 4.26 (s, 1H) | 3.80 | 3.80 | n/a | n/a | - | - | - | - |

| | | | | | | | | | | |
|---|---|---|---|---|---|---|---|---|---|---|
| Man-2 | 5.15 (s, 1H) | 4.21 | 3.93 | 3.54 | n/a | 3.92, 3.63 | - | - | - | - |
| Man-3 | 4.94 (s, 1H) | 4.12 | 3.91 | 3.51 | 3.63 | 3.93, 3.63 | - | - | - | - |
| GlcNAc-3 | 4.62 | 3.77 | n/a | n/a | 3.58 | 3.99, 3.85 | - | - | - | 2.11 – 2.00 (m, 24H) |
| Galactose-1 | 4.45 | 3.54 | 3.70 | 3.94 | n/a | 4.00, 3.55 | - | - | - | - |
| Sialic acid-1 | - | - | 2.68 (dd, J = 12.5, 4.6 Hz, 1H), 1.73 (t, J = 12.2 Hz, 1H) | 3.68 | 3.82 | n/a | 3.57 | 3.90 | 3.89, 3.66 | 2.11 – 2.00 (m, 24H) |
| GlcNAc-3’ | 4.59 (d, J = 8.2 Hz, 1H) | 3.77 | n/a | n/a | 3.58 | 3.99, 3.85 | - | - | - | 2.11 – 2.00 (m, 24H) |
| Galactose-1’ | 4.47 | 3.60 | 3.73 | 4.17 (d, J = 3.1 Hz, 1H) | 3.72 | 3.76 (4H) | - | - | - | - |
| GlcNAc-4’ | 4.71 | 3.82 | 3.73 | 3.74 | 3.60 | 3.98, 3.85 | - | - | - | 2.11 – 2.00 (m, 24H) |
| Galactose-2’ | 4.48 | 3.60 | 3.73 | 4.20 | 3.72 | 3.76 (4H) | - | - | - | - |
| GlcNAc-5’ | 4.75 | 3.85 | n/a | n/a | 3.82 | 4.43, 4.29 | - | - | - | 2.11 – 2.00 (m, 24H) |
| Galactose-3’ | 4.48 | 3.54 | 3.70 | 3.94 | n/a | 4.00, 3.55 | - | - | - | - |
| Sialic acid-2 | - | - | 2.68 (dd, J = 12.5, 4.6 Hz, 1H), 1.73 (t, J = 12.2 Hz, 1H) | 3.68 | 3.82 | n/a | 3.57 | 3.90 | 3.89, 3.66 | 2.11 – 2.00 (m, 24H) |

$^{13}C$ (150 MHz, $D_2O$): δ (ppm)

| | C-1 | C-2 | C-3 | C-4 | C-5 | C-6 | C-7 | C-8 | C-9 | NHAc |
|---|---|---|---|---|---|---|---|---|---|---|
| GlcNAc-1 | 78.08 | 53.73 | n/a | 78.79 | 76.30 | 59.92 | - | - | - | 22.23 |
| GlcNAc-2 | 101.39 | 54.84 | n/a | n/a | n/a | n/a | - | - | - | 22.23 |
| Man-1 | 100.47 | 70.26 | 80.65 | 65.58 | n/a | n/a | - | - | - | - |
| Man-2 | 99.58 | 76.42 | 69.33 | 67.45 | n/a | 61.72 | - | - | - | - |
| Man-3 | 97.04 | 76.35 | 69.56 | 67.45 | 74.57 | 61.60 | - | - | - | - |

| | | | | | | | | | | |
|---|---|---|---|---|---|---|---|---|---|---|
| GlcNAc-3 | 99.52 | 54.98 | n/a | n/a | 74.69 | 60.19 | - | - | - | 22.23 |
| Galactose-1 | 103.46 | 70.95 | 72.58 | 68.53 | n/a | 63.47 | - | - | - | - |
| Sialic acid-1 | n/a | n/a | 40.02 | n/a | 52.01 | n/a | 68.56 | 71.74 | 62.56 | 22.23 |
| GlcNAc-3’ | 99.47 | 55.06 | n/a | n/a | 74.69 | 60.19 | - | - | - | 22.23 |
| Galactose-1’ | 103.11 | 70.07 | 82.27 | 68.33 | 75.04 | 61.03 | - | - | - | - |
| GlcNAc-4’ | 102.87 | 55.21 | 72.58 | 78.81 | 74.62 | 60.08 | - | - | - | 22.23 |
| Galactose-2’ | 103.10 | 70.07 | 82.53 | 68.33 | 75.04 | 61.03 | - | - | - | - |
| GlcNAc-5’ | 102.57 | 55.11 | n/a | n/a | 72.38 | 66.93 | - | - | - | 22.23 |
| Galactose-3’ | 103.20 | 70.80 | 72.48 | 68.52 | n/a | 63.47 | - | - | - | - |
| Sialic acid-2 | n/a | n/a | 40.02 | n/a | 52.01 | n/a | 68.56 | 71.74 | 62.56 | 22.23 |

HRMS (ESI-MS): m/z calculated for $C_{116}H_{187}N_{10}O_{87}S$ $[M-3H]^{3-}$: 1048.3429; found: 1048.3315.

#### Compound 31

**31** was prepared from **29** (1.8 mg, 0.6 μmol) using the general procedure for the installation of α2,3-Neu5Ac with ST3Gal4. After P6 purification, **31** was obtained as a white solid (1.8 mg, 91%).

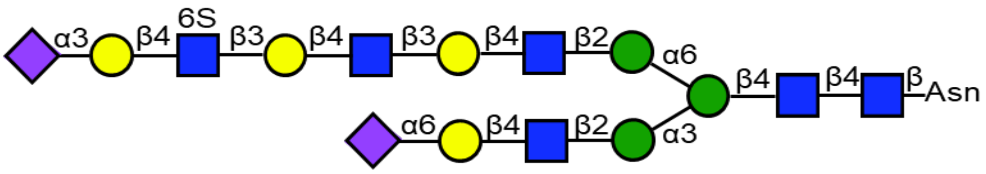

$^{1}H$ (600 MHz, $D_2O$): δ (ppm)

| | H-1 | H-2 | H-3 | H-4 | H-5 | H-6 | H-7 | H-8 | H-9 | NHAc |
|---|---|---|---|---|---|---|---|---|---|---|
| GlcNAc-1 | 5.08 (d, J = 9.8 Hz, 1H) | 3.88 | n/a | 3.67 | 3.60 | 3.77, 3.66 | - | - | - | 2.11 – 2.00 (m, 24H) |
| GlcNAc-2 | 4.63 | 3.80 | n/a | n/a | n/a | n/a | - | - | - | 2.11 – 2.00 (m, 24H) |
| Man-1 | 4.78 | 4.26 (s, 1H) | 3.80 | 3.80 | n/a | 3.97, 3.81 | - | - | - | - |
| Man-2 | 5.14 (s, 1H) | 4.21 | 3.93 | 3.54 | n/a | 3.92, 3.63 | - | - | - | - |
| Man-3 | 4.93 (s, 1H) | 4.12 | 3.91 | 3.51 | 3.63 | 3.93, 3.63 | - | - | - | - |
| GlcNAc-3 | 4.63 | 3.77 | n/a | n/a | 3.58 | 3.99, 3.85 | - | - | - | 2.11 – 2.00 (m, 24H) |
| Galactose-1 | 4.45 | 3.54 | 3.70 | 3.94 | n/a | 4.00, 3.55 | - | - | - | - |

| | | | | | | | | | | |
|---|---|---|---|---|---|---|---|---|---|---|
| Sialic acid-1 | - | - | 2.68 (dd, J = 12.3, 4.6 Hz, 1H), 1.73 (t, J = 12.2 Hz, 1H) | 3.69 | 3.82 | n/a | 3.57 | 3.90 | 3.89, 3.66 | 2.11 – 2.00 (m, 24H) |
| GlcNAc-3’ | 4.59 | 3.77 | n/a | n/a | 3.58 | 3.99, 3.85 | - | - | - | 2.11 – 2.00 (m, 24H) |
| Galactose-1’ | 4.47 | 3.60 | 3.73 | 4.17 (d, J = 3.1 Hz, 1H) | 3.72 | 3.76 (4H) | - | - | - | - |
| GlcNAc-4’ | 4.71 | 3.82 | 3.73 | 3.74 | 3.60 | 3.98, 3.85 | - | - | - | 2.11 – 2.00 (m, 24H) |
| Galactose-2’ | 4.48 | 3.60 | 3.73 | 4.20 | 3.72 | 3.76 (4H) | - | - | - | - |
| GlcNAc-5’ | 4.72 | 3.85 | n/a | 3.81 | 3.82 | 4.41 (d, J = 10.8 Hz, 1H), 4.32 (d, J = 10.6 Hz, 1H) | - | - | - | 2.11 – 2.00 (m, 24H) |
| Galactose-3’ | 4.61 | 3.57 | 4.13 | 3.98 | n/a | n/a | - | - | - | - |
| Sialic acid-2 | - | - | 2.76 (dd, J = 12.6, 4.4 Hz, 1H), 1.81 (t, J = 12.1 Hz, 1H) | 3.67 | 3.87 | n/a | n/a | n/a | n/a | 2.11 – 2.00 (m, 24H) |

$^{13}C$ (150 MHz, $D_2O$): δ (ppm)

| | C-1 | C-2 | C-3 | C-4 | C-5 | C-6 | C-7 | C-8 | C-9 | NHAc |
|---|---|---|---|---|---|---|---|---|---|---|
| GlcNAc-1 | 78.15 | 53.61 | n/a | 78.79 | 76.30 | 59.92 | - | - | - | 22.23 |
| GlcNAc-2 | 101.39 | 54.84 | n/a | n/a | n/a | n/a | - | - | - | 22.23 |
| Man-1 | 100.47 | 70.26 | 80.65 | 65.58 | n/a | 65.81 | - | - | - | |
| Man-2 | 99.58 | 76.42 | 69.33 | 67.45 | n/a | 61.72 | - | - | - | |
| Man-3 | 97.04 | 76.35 | 69.56 | 67.45 | 74.57 | 61.60 | - | - | - | |
| GlcNAc-3 | 99.56 | 54.98 | n/a | n/a | 74.69 | 60.19 | - | - | - | 22.23 |
| Galactose-1 | 103.46 | 70.78 | 72.58 | 68.53 | n/a | 63.47 | - | - | - | |

| | | | | | | | | | | |
|---|---|---|---|---|---|---|---|---|---|---|
| Sialic acid-1 | n/a | n/a | 40.10 | n/a | 51.88 | n/a | 68.56 | 71.74 | 62.56 | 22.23 |
| GlcNAc-3’ | 99.47 | 55.06 | n/a | n/a | 74.69 | 60.19 | - | - | - | 22.23 |
| Galactose-1’ | 103.11 | 70.07 | 82.53 | 68.33 | 75.12 | 61.22 | - | - | - | |
| GlcNAc-4’ | 102.87 | 55.21 | 72.58 | 78.81 | 74.62 | 60.08 | - | - | - | 22.23 |
| Galactose-2’ | 103.10 | 70.07 | 82.53 | 68.33 | 75.12 | 61.22 | - | - | - | |
| GlcNAc-5’ | 102.87 | 55.21 | n/a | 77.75 | 72.46 | 66.53 | - | - | - | 22.23 |
| Galactose-3’ | 102.18 | 69.01 | 75.41 | 67.57 | n/a | n/a | - | - | - | |
| Sialic acid-2 | n/a | n/a | 39.60 | n/a | 51.76 | n/a | n/a | n/a | n/a | 22.23 |

HRMS (ESI-MS): m/z calculated for $C_{116}H_{188}N_{10}O_{87}S$ $[M-2H]^{2-}$: 1573.0179; found: 1573.0514.

#### Compound 32

**32** and **33** were prepared from **31** (1.4 mg, 0.44 μmol) using the rate control procedure for the 6-O-sulfate installation of internal galactose with CHST1. When no further product formation was observed, LC-MS analysis indicated a mixture of very small amount of mono-6-O-sulfated glycan **31**, and substantial quantities of di-6-O-sulfated glycan **32** and tri-6-O-sulfated glycan **33**. After P6 and DEAE purification, **32** was obtained as a white solid (1.0 mg, 70%), and **33** as a white solid (0.2 mg, 14%). Unreacted **31** was recovered as a white solid (0.1 mg).

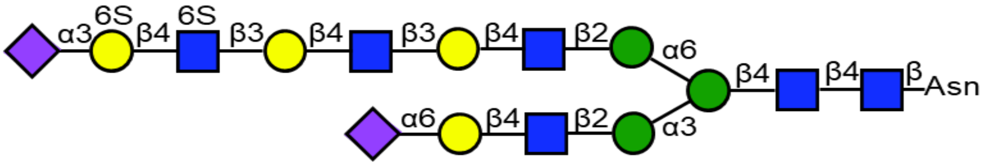

$^1$H (600 MHz, $D_2O$): δ (ppm)

| | H-1 | H-2 | H-3 | H-4 | H-5 | H-6 | H-7 | H-8 | H-9 | NHAc |
|---|---|---|---|---|---|---|---|---|---|---|
| GlcNAc-1 | 5.08 (d, J = 8.9 Hz, 1H) | 3.88 | n/a | 3.67 | 3.60 | 3.77, 3.66 | - | - | - | 2.12 – 2.00 (m, 24H) |
| GlcNAc-2 | 4.63 | 3.80 | n/a | n/a | n/a | n/a | - | - | - | 2.12 – 2.00 (m, 24H) |
| Man-1 | 4.78 | 4.26 (s, 1H) | 3.80 | 3.80 | n/a | n/a | - | - | - | - |
| Man-2 | 5.14 (s, 1H) | 4.20 | 3.93 | 3.54 | n/a | 3.92, 3.63 | - | - | - | - |
| Man-3 | 4.93 (s, 1H) | 4.12 | 3.91 | 3.51 | 3.63 | 3.93, 3.63 | - | - | - | - |
| GlcNAc-3 | 4.62 | 3.77 | n/a | n/a | 3.58 | 3.99, 3.85 | - | - | - | 2.12 – 2.00 (m, 24H) |

| | | | | | | | | | | |
|---|---|---|---|---|---|---|---|---|---|---|
| Galactose-1 | 4.45 | 3.55 | 3.70 | 3.93 | n/a | 4.00, 3.55 | - | - | - | - |
| Sialic acid-1 | - | - | 2.68, 1.73 (t, J = 11.8 Hz, 1H) | 3.69 | 3.81 | n/a | 3.57 | 3.90 | 3.89, 3.66 | 2.12 – 2.00 (m, 24H) |
| GlcNAc-3’ | 4.59 | 3.77 | n/a | n/a | 3.58 | 3.99, 3.85 | - | - | - | 2.12 – 2.00 (m, 24H) |
| Galactose-1’ | 4.47 | 3.59 | 3.73 | 4.17 | 3.73 | 3.76 (4H) | - | - | - | - |
| GlcNAc-4’ | 4.71 | 3.80 | 3.73 | 3.74 | 3.59 | 3.98, 3.85 | - | - | - | 2.12 – 2.00 (m, 24H) |
| Galactose-2’ | 4.48 | 3.59 | 3.73 | 4.21 | 3.73 | 3.76 (4H) | - | - | - | - |
| GlcNAc-5’ | 4.72 | 3.82 | n/a | n/a | 3.85 | 4.44, 4.30 | - | - | - | 2.12 – 2.00 (m, 24H) |
| Galactose-3’ | 4.63 | 3.58 | 4.15 | 4.03 | 3.99 | 4.19 (2H) | - | - | - | - |
| Sialic acid-2 | - | - | 2.75, 1.82 (t, J = 11.6 Hz, 1H) | 3.67 | 3.87 | n/a | n/a | n/a | n/a | 2.12 – 2.00 (m, 24H) |

$^{13}C$ (150 MHz, $D_2O$): δ (ppm)

| | C-1 | C-2 | C-3 | C-4 | C-5 | C-6 | C-7 | C-8 | C-9 | NHAc |
|---|---|---|---|---|---|---|---|---|---|---|
| GlcNAc-1 | 78.15 | 53.61 | n/a | 78.79 | 76.30 | 59.92 | - | - | - | 22.23 |
| GlcNAc-2 | 101.39 | 54.84 | n/a | n/a | n/a | n/a | - | - | - | 22.23 |
| Man-1 | 100.47 | 70.23 | 80.65 | 65.58 | n/a | n/a | - | - | - | - |
| Man-2 | 99.58 | 76.47 | 69.33 | 67.45 | n/a | 61.72 | - | - | - | - |
| Man-3 | 97.04 | 76.39 | 69.56 | 67.45 | 74.57 | 61.60 | - | - | - | - |
| GlcNAc-3 | 99.56 | 54.98 | n/a | n/a | 74.69 | 60.19 | - | - | - | 22.23 |
| Galactose-1 | 103.46 | 70.89 | 72.78 | 68.67 | n/a | n/a | - | - | - | - |
| Sialic acid-1 | n/a | n/a | 40.10 | n/a | 51.74 | n/a | 68.56 | 71.74 | 62.56 | 22.23 |
| GlcNAc-3’ | 99.47 | 55.06 | n/a | n/a | 74.69 | 60.19 | - | - | - | 22.23 |
| Galactose-1’ | 103.11 | 69.97 | 82.45 | 68.60 | 75.15 | 61.23 | - | - | - | - |
| GlcNAc-4’ | 102.87 | 55.21 | 72.46 | 78.86 | 74.71 | 60.09 | - | - | - | 22.23 |
| Galactose-2’ | 103.10 | 69.97 | 82.45 | 68.53 | 75.15 | 61.23 | - | - | - | - |
| GlcNAc-5’ | 102.87 | 55.21 | n/a | n/a | 72.05 | 66.87 | - | - | - | 22.23 |
| Galactose-3’ | 102.54 | 69.47 | 75.30 | 67.36 | 72.89 | 67.18 | - | - | - | - |
| Sialic acid-2 | n/a | n/a | 39.41 | n/a | 51.63 | n/a | n/a | n/a | n/a | 22.23 |

HRMS (ESI-MS): m/z calculated for $C_{116}H_{187}N_{10}O_{90}S_2$ $[M-3H]^{3-}$: 1074.9951; found: 1074.6877.

**Compound 33**

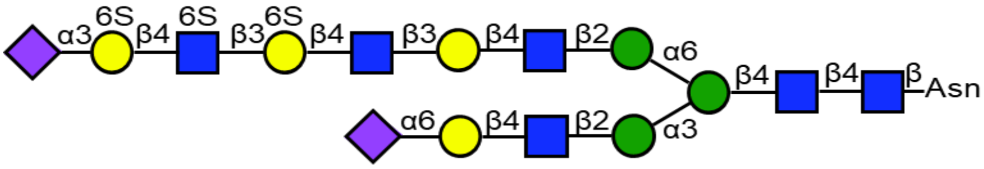

$^1$H (600 MHz, $D_2O$): δ (ppm)

| | H-1 | H-2 | H-3 | H-4 | H-5 | H-6 | H-7 | H-8 | H-9 | NHAc |
|---|---|---|---|---|---|---|---|---|---|---|
| GlcNAc-1 | 5.08 (d, J = 9.6 Hz, 1H) | 3.88 | n/a | 3.67 | 3.60 | 3.77, 3.66 | - | - | - | 2.11 – 2.00 (m, 24H) |
| GlcNAc-2 | 4.63 | 3.80 | n/a | n/a | n/a | n/a | - | - | - | 2.11 – 2.00 (m, 24H) |
| Man-1 | 4.79 | 4.26 (s, 1H) | 3.80 | 3.80 | n/a | n/a | - | - | - | - |
| Man-2 | 5.14 (s, 1H) | 4.20 | 3.93 | 3.54 | n/a | 3.92, 3.63 | - | - | - | - |
| Man-3 | 4.93 (s, 1H) | 4.12 | 3.91 | 3.51 | 3.63 | 3.93, 3.63 | - | - | - | - |
| GlcNAc-3 | 4.62 | 3.80 | n/a | n/a | 3.58 | 3.99, 3.85 | - | - | - | 2.11 – 2.00 (m, 24H) |
| Galactose-1 | 4.45 | 3.54 | 3.69 | 3.93 | n/a | 4.00, 3.55 | - | - | - | - |
| Sialic acid-1 | - | - | 2.68 (dd, J = 12.4, 4.7 Hz, 1H), 1.73 (t, J = 12.1 Hz, 1H) | 3.70 | 3.82 | n/a | 3.57 | 3.90 | 3.89, 3.66 | 2.11 – 2.00 (m, 24H) |
| GlcNAc-3’ | 4.59 (d, J = 8.2 Hz, 1H) | 3.77 | n/a | n/a | 3.58 | 3.99, 3.85 | - | - | - | 2.11 – 2.00 (m, 24H) |
| Galactose-1’ | 4.47 | 3.59 | 3.75 | 4.17 | 3.74 | 3.78 (2H) | - | - | - | - |
| GlcNAc-4’ | 4.70 | 3.81 | 3.72 | 3.71 | n/a | 3.99, 3.85 | - | - | - | 2.11 – 2.00 (m, 24H) |

| | | | | | | | | | | |
|---|---|---|---|---|---|---|---|---|---|---|
| Galactose-2’ | 4.52 (d, J = 7.9 Hz, 1H) | 3.59 | 3.75 | 4.25 | 4.00 | 4.20 (4H) | - | - | - | - |
| GlcNAc-5’ | 4.74 (d, J = 8.3 Hz, 1H) | 3.83 | n/a | n/a | 3.84 | 4.44, 4.33 (dd, J = 11.3, 5.0 Hz, 1H) | - | - | - | 2.11 – 2.00 (m, 24H) |
| Galactose-3’ | 4.64 | 3.59 | 4.16 | 4.04 | 4.00 | 4.20 (4H) | - | - | - | - |
| Sialic acid-2 | - | - | 2.75 (dd, J = 12.5, 4.6 Hz, 1H), 1.82 (t, J = 12.0 Hz, 1H) | 3.67 | 3.87 | n/a | n/a | n/a | n/a | 2.11 – 2.00 (m, 24H) |

$^{13}C$ (150 MHz, $D_2O$): δ (ppm)

| | C-1 | C-2 | C-3 | C-4 | C-5 | C-6 | C-7 | C-8 | C-9 | NHAc |
|---|---|---|---|---|---|---|---|---|---|---|
| GlcNAc-1 | 78.15 | n/a | n/a | n/a | n/a | 59.92 | - | - | - | 22.27 |
| GlcNAc-2 | 101.53 | 54.84 | n/a | n/a | n/a | n/a | - | - | - | 22.27 |
| Man-1 | 100.54 | n/a | 80.65 | n/a | n/a | n/a | - | - | - | - |
| Man-2 | 99.58 | 76.31 | 69.33 | n/a | n/a | 61.72 | - | - | - | - |
| Man-3 | 96.99 | 76.26 | 69.56 | n/a | n/a | 61.60 | - | - | - | - |
| GlcNAc-3 | 99.56 | 54.98 | n/a | n/a | 74.69 | 60.19 | - | - | - | 22.27 |
| Galactose-1 | 103.46 | 70.87 | 72.66 | 68.84 | n/a | n/a | - | - | - | - |
| Sialic acid-1 | n/a | n/a | 40.06 | n/a | 51.88 | n/a | n/a | n/a | 62.56 | 22.27 |
| GlcNAc-3’ | 99.47 | 55.06 | n/a | n/a | 74.69 | 60.19 | - | - | - | 22.27 |
| Galactose-1’ | 103.31 | 69.30 | 82.29 | 68.60 | 74.77 | 60.87 | - | - | - | - |
| GlcNAc-4’ | 102.87 | 55.21 | 72.59 | 79.07 | n/a | 60.17 | - | - | - | 22.27 |
| Galactose-2’ | 102.76 | 69.30 | 82.29 | 67.89 | 72.74 | 67.28 | - | - | - | - |
| GlcNAc-5’ | 102.87 | 55.21 | n/a | n/a | 72.37 | 66.59 | - | - | - | 22.27 |
| Galactose-3’ | 102.46 | 69.30 | 75.23 | 67.41 | 72.74 | 67.28 | - | - | - | - |
| Sialic acid-2 | n/a | n/a | 39.45 | n/a | 51.79 | n/a | n/a | n/a | n/a | 22.27 |

HRMS (ESI-MS): m/z calculated for $C_{116}H_{187}N_{10}O_{93}S_3$ $[M-3H]^{3-}$: 1101.6474; found: 1101.6758.

#### Cbz deprotection by hydrogenation over $Pd(OH)_2$ for microarray development

#### Compound 34

**34** was prepared from **12** (1.3 mg, 0.23 µmol) using the general procedure for Cbz deprotection using $Pd(OH)_2$. After purification, **34** was obtained as a white solid (0.4 mg, 21%).

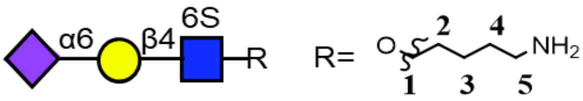

$^{1}$H (600 MHz, $D_2O$): δ (ppm)

| | H-1 | H-2 | H-3 | H-4 | H-5 | H-6 | H-7 | H-8 | H-9 | NHAc |
|---|---|---|---|---|---|---|---|---|---|---|
| GlcNAc-6S | 4.59 (d, J = 7.6 Hz, 1H) | 3.76 | n/a | 3.67 | 3.84 | 4.46-4.44 (m, 1H), 4.27 (dd, J = 11.1, 5.8 Hz, 1H) | - | - | - | 2.07 – 2.01 (m, 6H) |
| Galactose | 4.49 (d, J = 7.9 Hz, 1H) | 3.55 | 3.69 | 3.94 (d, J = 3.5 Hz, 1H) | n/a | 4.01, 3.54 | - | - | - | - |
| Sialic acid | - | - | 2.68 (dd, J = 12.4, 4.7 Hz, 1H), 1.73 | 3.69 | 3.81 | n/a | 3.57 | 3.90 | 3.89, 3.65 | 2.07 – 2.01 (m, 6H) |

$^{13}$C (150 MHz, $D_2O$): δ (ppm)

| | C-1 | C-2 | C-3 | C-4 | C-5 | C-6 | C-7 | C-8 | C-9 | NHAc |
|---|---|---|---|---|---|---|---|---|---|---|
| GlcNAc-6S | 101.09 | 54.89 | n/a | 80.71 | 73.82 | 67.09 | - | - | - | 22.35 |
| Galactose | 103.64 | 71.01 | 72.65 | 68.58 | n/a | 63.51 | - | - | - | - |
| Sialic acid | n/a | n/a | 40.27 | n/a | 51.89 | n/a | 68.56 | 71.91 | 62.82 | 22.35 |

| Linker | 1 | 2 | 3 | 4 | 5 |
|---|---|---|---|---|---|
| H | 3.90, 3.66 | 1.63 (p, J = 6.9 Hz, 2H) | 1.49 – 1.37 (m, 2H) | 1.69 (p, J = 7.8 Hz, 2H) | 3.07 – 2.91 (m, 2H) |
| C | 70.32 | 28.23 | 22.15 | 26.42 | 39.44 |

HRMS (ESI-MS): m/z calculated for $C_{30}H_{52}N_3O_{22}S$ $[M-H]^-$: 838.2768; found: 838.2776.

**Compound 35**

**35** was prepared from **13** (1.3 mg, 0.97 µmol) using the general procedure for Cbz deprotection using $Pd(OH)_2$ reduction. After purification, **35** was obtained as a white solid (0,6 mg, 51%).

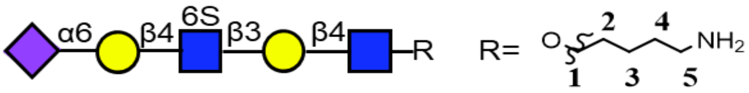

[1]H (600 MHz, $D_2O$): δ (ppm)

| | H-1 | H-2 | H-3 | H-4 | H-5 | H-6 | H-7 | H-8 | H-9 | NHAc |
|---|---|---|---|---|---|---|---|---|---|---|
| GlcNAc | 4.53 (d, J = 7.5 Hz, 1H) | 3.73 | 3.70 | 3.70 | 3.60 | 4.00, 3.84 | - | - | - | 2.07 – 2.03 (m, 9H) |
| Galactose-1 | 4.47 | 3.60 | 3.74 | 4.21 (d, J = 3.2 Hz, 1H) | n/a | 3.77 (2H) | - | - | - | - |
| GlcNAc-6S | 4.75 (d, J = 8.2 Hz, 1H) | 3.84 | n/a | 3.69 | 3.85 | 4.44 (dd, J = 11.2, 2.1 Hz, 1H), 4.28 (dd, J = 11.1, 5.9 Hz, 1H) | - | - | - | 2.07 – 2.03 (m, 9H) |
| Galactose-2 | 4.49 | 3.55 | 3.71 | 3.94 (d, J = 3.5 Hz, 1H) | n/a | 4.01, 3.55 | - | - | - | - |
| Sialic acid | - | - | 2.68 (dd, J = 12.4, 4.6 Hz, 1H), 1.74 (t, J = 12.2 Hz, 1H) | 3.67 | 3.82 | n/a | 3.57 | 3.91 | 3.89, 3.65 | 2.07 – 2.03 (m, 9H) |

[13]C (150 MHz, $D_2O$): δ (ppm)

| | C-1 | C-2 | C-3 | C-4 | C-5 | C-6 | C-7 | C-8 | C-9 | NHAc |
|---|---|---|---|---|---|---|---|---|---|---|
| GlcNAc | 101.24 | 54.99 | 72.53 | 78.47 | 74.85 | 60.16 | - | - | - | 22.16 |
| Galactose-1 | 103.49 | 69.59 | 82.25 | 68.27 | n/a | 61.23 | - | - | - | - |
| GlcNAc-6S | 102.60 | 54.67 | n/a | 80.53 | 73.99 | 66.93 | - | - | - | 22.16 |
| Galactose-2 | 102.60 | 70.89 | 72.53 | 68.21 | n/a | 63.31 | - | - | - | - |
| Sialic acid | n/a | n/a | 40.10 | n/a | 51.76 | n/a | 68.64 | 72.15 | 62.74 | 22.16 |

| Linker | 1 | 2 | 3 | 4 | 5 |
|---|---|---|---|---|---|
| H | 3.91, 3.63 | 1.61 (p, J = 6.8 Hz, 2H) | 1.46 – 1.36 (m, 2H) | 1.71 – 1.65 (m, 2H) | 3.00 (t, J = 7.6 Hz, 2H) |

| | | | | | |
|---|---|---|---|---|---|
| C | 69.66 | 28.13 | 22.18 | 26.44 | 39.26 |

HRMS (ESI-MS): m/z calculated for $C_{44}H_{74}N_4O_{32}S$ $[M-2H]^{2-}$: 601.2009; found: 601.1941.

#### Compound 36

**36** was prepared from **14** (1.1 mg, 0.64 µmol) using the general procedure for Cbz deprotection using $Pd(OH)_2$ reduction. After purification, **36** was obtained as a white solid (0.5 mg, 50%).

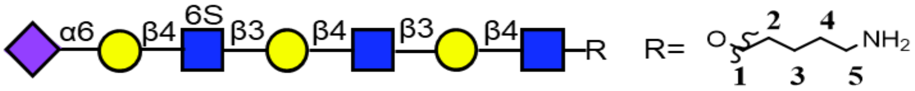

$^1$H (600 MHz, $D_2O$): δ (ppm)

| | H-1 | H-2 | H-3 | H-4 | H-5 | H-6 | H-7 | H-8 | H-9 | NHAc |
|---|---|---|---|---|---|---|---|---|---|---|
| GlcNAc-1 | 4.53 (d, J = 7.3 Hz, 1H) | 3.73 | 3.70 | 3.70 | 3.60 | 3.98, 3.84 | - | - | - | 2.07 – 2.01 (m, 12H) |
| Galactose-1 | 4.46 | 3.60 | 3.73 | 4.17 (d, J = 3.2 Hz, 1H) | n/a | 3.77 (4H) | - | - | - | - |
| GlcNAc-2 | 4.71 (d, J = 8.4 Hz, 1H) | 3.82 | 3.72 | 3.73 | 3.60 | 3.98, 3.84 | - | - | - | 2.07 – 2.01 (m, 12H) |
| Galactose-2 | 4.49 | 3.60 | 3.73 | 4.22 – 4.19 (m, 1H) | n/a | 3.77 (4H) | - | - | - | - |
| GlcNAc-6S | 4.75 | 3.85 | n/a | 3.69 | 3.85 | 4.44 (d, J = 10.7 Hz, 1H), 4.28 (dd, J = 11.1, 5.9 Hz, 1H) | - | - | - | 2.07 – 2.01 (m, 12H) |
| Galactose-3 | 4.49 | 3.54 | 3.72 | 3.94 (d, J = 3.2 Hz, 1H) | n/a | 4.01, 3.55 | - | - | - | - |
| Sialic acid | - | - | 2.68 (dd, J = 12.4, 4.6 Hz, 1H), 1.74 | 3.70 | 3.82 | n/a | 3.57 | 3.91 | 3.89, 3.65 | 2.07 – 2.01 (m, 12H) |

$^{13}$C (150 MHz, $D_2O$): δ (ppm)

| | C-1 | C-2 | C-3 | C-4 | C-5 | C-6 | C-7 | C-8 | C-9 | NHAc |
|---|---|---|---|---|---|---|---|---|---|---|
| GlcNAc-1 | 101.28 | 55.19 | 72.38 | 78.64 | 74.86 | 60.16 | - | - | - | 22.20 |
| Galactose-1 | 102.96 | 70.00 | 82.41 | 68.41 | n/a | 61.21 | - | - | - | - |
| GlcNAc-2 | 102.88 | 55.19 | 72.57 | 78.36 | 74.86 | 60.16 | - | - | - | 22.20 |
| Galactose-2 | 103.39 | 70.00 | 82.41 | 68.41 | n/a | 61.21 | - | - | - | - |
| GlcNAc-6S | 102.69 | 55.08 | n/a | 80.84 | 73.86 | 67.10 | - | - | - | 22.20 |
| Galactose-3 | 103.28 | 70.93 | 72.50 | 68.41 | n/a | 63.46 | - | - | - | - |
| Sialic acid | n/a | n/a | 40.24 | n/a | 52.02 | n/a | 68.64 | 72.15 | 62.82 | 22.20 |

| Linker | 1 | 2 | 3 | 4 | 5 |
|---|---|---|---|---|---|
| H | 3.91,3.62 | 1.61 (p, J = 6.8 Hz, 2H) | 1.46 – 1.36 (m, 2H) | 1.71 – 1.65 (m, 2H) | 3.00 (t, J = 7.7 Hz, 2H) |
| C | 70.31 | 28.30 | 22.26 | 26.50 | 39.60 |

HRMS (ESI-MS): m/z calculated for $C_{58}H_{97}N_5O_{42}S$ $[M-2H]^{2-}$: 783.7670; found: 783.7381.

#### Compound 37

**37** was prepared from **16** (0.3 mg, 0.24 µmol) using the general procedure for Cbz deprotection using $Pd(OH)_2$ reduction. After purification, **37** was obtained as a white solid (140 µg, 48%).

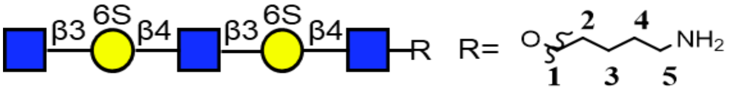

$^1H$ (600 MHz, $D_2O$): δ (ppm)

| | H-1 | H-2 | H-3 | H-4 | H-5 | H-6 | NHAc |
|---|---|---|---|---|---|---|---|
| GlcNAc-1 | 4.53 | 3.74 | 3.73 | 3.68 | 3.60 | 3.97, 3.83 | 2.07 – 1.99 (m, 9H) |
| Galactose-1 | 4.51 | 3.59 | 3.75 | 4.24 – 4.22 (m, 2H) | 3.98 | 4.22 – 4.18 (m, 4H) | - |
| GlcNAc-2 | 4.71 | 3.80 | 3.75 | 3.72 | 3.60 | 3.97, 3.83 | 2.07 – 1.99 (m, 9H) |
| Galactose-2 | 4.51 | 3.59 | 3.75 | 4.24 – 4.22 (m, 2H) | 3.98 | 4.22 – 4.18 (m, 4H) | - |
| GlcNAc-3 | 4.70 | 3.77 | n/a | n/a | 3.46 | 3.90, 3.77 | 2.07 – 1.99 (m, 9H) |

$^{13}C$ (150 MHz, $D_2O$): δ (ppm)

| | C-1 | C-2 | C-3 | C-4 | C-5 | C-6 | NHAc |
|---|---|---|---|---|---|---|---|
| GlcNAc-1 | 101.11 | 55.47 | 72.06 | 79.28 | 74.35 | 60.20 | 22.07 |
| Galactose-1 | 103.05 | 69.94 | 82.42 | 68.02 | 72.67 | 67.32 | - |
| GlcNAc-2 | 102.93 | 55.05 | 72.06 | 79.07 | 74.35 | 60.20 | 22.07 |
| Galactose-2 | 103.05 | 69.94 | 82.42 | 68.02 | 72.67 | 67.32 | - |

| GlcNAc-3 | 102.93 | 55.58 | n/a | n/a | 75.96 | 60.47 | 22.07 |
|---|---|---|---|---|---|---|---|

| Linker | 1 | 2 | 3 | 4 | 5 |
|---|---|---|---|---|---|
| H | 3.91,3.62 | 1.60 (p, J = 6.9 Hz, 2H) | 1.46 – 1.36 (m, 2H) | 1.71 – 1.65 (m, 2H) | 3.00 (t, J = 7.6 Hz, 2H) |
| C | 69.90 | 28.30 | 22.12 | 26.50 | 39.15 |

HRMS (ESI-MS): m/z calculated for $C_{41}H_{70}N_4O_{32}S_2$ $[M-2H]^{2-}$: 597.1713; found: 597.1634.

#### Compound 38

**38** was prepared from **18** (0.4 mg, 0.23 µmol) using the general procedure for Cbz deprotection using $Pd(OH)_2$ reduction. After purification, **38** was obtained as a white solid (120 µg, 32%).

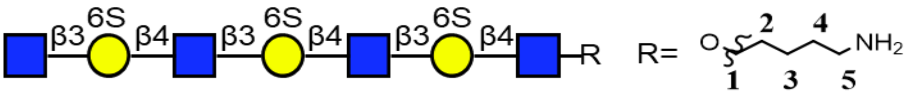

$^1$H (600 MHz, $D_2O$): δ (ppm)

| | H-1 | H-2 | H-3 | H-4 | H-5 | H-6 | NHAc |
|---|---|---|---|---|---|---|---|
| GlcNAc-1 | 4.53 | 3.74 | 3.72 | 3.69 | 3.61 | 3.97, 3.84 | 2.04 (s, 12H) |
| Galactose-6S-1 | 4.51 | 3.60 | 3.75 | 4.23 | 3.98 | 4.20 (6H) | - |
| GlcNAc-2 | 4.71 | 3.80 | 3.75 | 3.73 | 3.61 | 3.97, 3.84 | 2.04 (s, 12H) |
| Galactose-6S-2 | 4.51 | 3.60 | 3.75 | 4.23 | 3.98 | 4.20 (6H) | - |
| GlcNAc-3 | 4.71 | 3.80 | 3.75 | 3.73 | 3.61 | 3.97, 3.84 | 2.04 (s, 12H) |
| Galactose-6S-3 | 4.51 | 3.60 | 3.75 | 4.23 | 3.98 | 4.20 (6H) | - |
| GlcNAc-4 | 4.71 | 3.77 | n/a | n/a | 3.46 | 3.91, 3.76 | 2.04 (s, 12H) |

$^{13}$C (150 MHz, $D_2O$): δ (ppm)

| | C-1 | C-2 | C-3 | C-4 | C-5 | C-6 | NHAc |
|---|---|---|---|---|---|---|---|
| GlcNAc-1 | 100.83 | 55.50 | 72.43 | 79.13 | 74.79 | 60.12 | 22.22 |
| Galactose-6S-1 | 102.59 | 69.80 | 82.10 | 68.33 | 72.55 | 67.19 | - |
| GlcNAc-2 | 102.72 | 55.25 | 71.91 | 79.38 | 74.79 | 60.12 | 22.22 |
| Galactose-6S-2 | 102.59 | 69.80 | 82.10 | 68.33 | 72.55 | 67.19 | - |
| GlcNAc-3 | 102.72 | 55.25 | 71.91 | 79.38 | 74.79 | 60.12 | 22.22 |
| Galactose-6S-3 | 102.59 | 69.80 | 82.10 | 68.33 | 72.55 | 67.19 | - |
| GlcNAc-4 | 102.72 | 55.25 | n/a | n/a | 75.89 | 60.61 | 22.22 |

| Linker | 1 | 2 | 3 | 4 | 5 |
|---|---|---|---|---|---|
| H | 3.91, n/a | 1.60 (q, J = 7.0 Hz, 2H) | 1.46 – 1.37 (m, 2H) | 1.68 (p, J = 7.8 Hz, 2H) | 3.00 (t, J = 7.7 Hz, 2H) |

| | | | | | |
|---|---|---|---|---|---|
| C | 69.53 | 28.25 | 22.25 | 26.51 | 39.44 |

HRMS (ESI-MS): m/z calculated for $C_{55}H_{92}N_5O_{45}S_3$ $[M-3H]^{3-}$: 546.1414; found: 546.1389.

#### Compound 39

**39** was prepared from **21** (0.4 mg, 0.20 µmol) using the general procedure for Cbz deprotection using $Pd(OH)_2$ reduction. After purification, **39** was obtained as a white solid (150 µg, 45%).

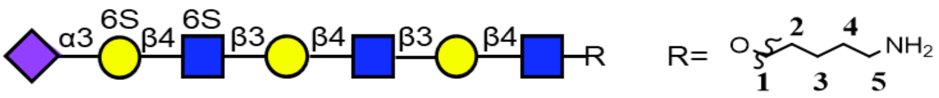

$^{1}H$ (600 MHz, $D_2O$): δ (ppm)

| | H-1 | H-2 | H-3 | H-4 | H-5 | H-6 | H-7 | H-8 | H-9 | NHAc |
|---|---|---|---|---|---|---|---|---|---|---|
| GlcNAc-1 | 4.53 | 3.73 | 3.70 | 3.70 | 3.59 | 3.97, 3.84 | - | - | - | 2.06 – 1.99 (m, 12H) |
| Galactose-1 | 4.46 | 3.59 | 3.73 | 4.16 | 3.72 | 3.76 | - | - | - | - |
| GlcNAc-2 | 4.71 | 3.81 | 3.73 | 3.72 | 3.59 | 3.97, 3.84 | - | - | - | 2.06 – 1.99 (m, 12H) |
| Galactose-2 | 4.48 | 3.59 | 3.73 | 4.21 | 3.72 | 3.76 | - | - | - | - |
| GlcNAc-6S | 4.71 | 3.81 | n/a | n/a | 3.86 | 4.45, 4.29 | - | - | - | 2.06 – 1.99 (m, 12H) |
| Galactose-6S | 4.63 (d, J = 7.6 Hz, 1H) | 3.59 | 4.15 | 4.03 | n/a | 4.19 | - | - | - | - |
| Sialic acid | - | - | 2.76 (dd, J = 12.5, 4.6 Hz, 1H), 1.82 (t, J = 12.1 Hz, 1H) | 3.68 | 3.86 | n/a | n/a | n/a | n/a | 2.06 – 1.99 (m, 12H) |

$^{13}C$ (150 MHz, $D_2O$): δ (ppm)

| | C-1 | C-2 | C-3 | C-4 | C-5 | C-6 | C-7 | C-8 | C-9 | NHAc |
|---|---|---|---|---|---|---|---|---|---|---|
| GlcNAc-1 | 101.21 | 55.00 | 72.52 | 78.46 | 74.87 | 59.95 | - | - | - | 2.06 – 1.99 |

| | | | | | | | | | | |
|---|---|---|---|---|---|---|---|---|---|---|
| | | | | | | | | | | (m, 3H) |
| Galactose-1 | 103.07 | 69.72 | 82.60 | 68.72 | 75.18 | 61.04 | - | - | - | - |
| GlcNAc-2 | 102.78 | 55.11 | 72.22 | 78.55 | 74.87 | 59.95 | - | - | - | 2.06 – 1.99 (m, 3H) |
| Galactose-2 | 103.07 | 69.72 | 82.60 | 68.57 | 75.18 | 61.04 | - | - | - | - |
| GlcNAc-6S | 102.78 | 55.11 | n/a | n/a | n/a | n/a | - | - | - | 2.06 – 1.99 (m, 3H) |
| Galactose-6S | 102.42 | 69.55 | 75.14 | 67.63 | n/a | 67.06 | - | - | - | - |
| Sialic acid | n/a | n/a | n/a | n/a | n/a | n/a | n/a | n/a | n/a | 2.06 – 1.99 (m, 3H) |

| Linker | 1 | 2 | 3 | 4 | 5 |
|---|---|---|---|---|---|
| H | 3.91,3.62 | 1.61 (p, J = 6.8 Hz, 2H) | 1.46 – 1.36 (m, 2H) | 1.71 – 1.65 (m, 2H) | 3.04 – 2.95 (m, 2H) |
| C | 69.90 | 28.08 | 22.25 | 26.41 | 39.26 |

HRMS (ESI-MS): m/z calculated for $C_{58}H_{97}N_5O_{45}S_2$ $[M-2H]^{2-}$: 823.7454; found: 823.7531.

**8) Microarray Procedure**

**Protein Design and Expression**

The pA-LS, containing domain B of protein A (pA) of Staphylococcus aureus (amino acid 212-270, UniProt accession number P38507) and 6,7-dimethyl-8-ribityllumazine synthase (LS) of Aquifex aeolicus (GenBank accession number WP_010880027.1), was constructed in a pUC57 plasmid by GenScript USA, Inc. Besides pA and LS, the pA-LS sequence contained a N-terminal Gly-Ser linker and a streptavidin tag II (WSHPQFEK).[4] The pA-LS was ligated into an expression vector, containing a CD5 signal sequence. pCDNA5-Siglec plasmids were kindly provided by Matthew Macauley, University of Alberta.[5]

Recombinant trimeric IAV hemagglutinin ectodomain proteins (HA) were cloned into the pCD5 expression vector (an example is addgene plasmid #182546)[6] in frame with a GCN4 trimerization motif (KQIEDKIEEIESKQKKIENEIARIKK), a superfolder GFP[7] and the Twin-Strep-tag (WSHPQFEKGGGSGGGSWSHPQFEK); IBA, Germany). Mutations in HAs were generated by site-directed mutagenesis.

The proteins were expressed by poly-ethylenimine I (PEI)-transfecting 40-60% confluent HEK293S GnTI(-) cells. Before addition to the cells, the DNA/PEI mix was incubated on Dulbecco's Modified Eagle Medium (DMEM) for 20 min and 1/3 of the medium was removed from the cell dishes. At 6 h post-transfection, the medium was replaced with 293 SFM II medium (Gibco) supplemented with Primatone (3.0 g/L), bicarbonate (3.6 g/L), glucose (2.0 g/L), valproic acid (0.4 g/L), glutaMAX (1%), and DMSO (1.5%). Cells were incubated for 5 days at 37 °C and 5% $CO_2$ before supernatants were collected. Proteins containing superfolder GFP were quantified by measuring fluorescence (excitation 480 nm; emission 520 nm) with the POLARstar Omega (BMG Labtech). Protein expression was checked by western blotting using a StrepMAB-Classic HRP antibody (IBA Lifesciences). All proteins were purified using Strep-Tactin Sepharose beads (IBA Lifesciences) and subsequently analyzed on SDS-PAGE gels, which were stained with Coomassie blue.

**Glycan Microarray Binding Studies**

Siglecs, and HAs, both at 50 µg/ml, were either premixed with pA-LS or pre-complexed with human anti-streptag and goat anti-human-Alexa555 (#A21433, Thermo Fisher Scientific) antibodies in a 4:2:1 molar ratio respectively in 50 µL PBS with 0.1% Tween-20. Biotinylated lectins (5 µg/mL) were pre-complexed with streptavidin-Alexa555 (#S32355, Thermo Fisher Scientific) in a 5:1 weight ratio. The following biotinylated lectins from Vector Laboratories were used: MAL-I (B-1315-2), MAL-II (B-1265-1, and SNA (B-1305-2). The mixtures were incubated on ice for 15 min and afterward incubated on the surface of the array for 90 min in a humidified chamber. The siglec-pA-LS complexes where subsequently detected with human anti-streptag (10 µg/mL) and thereafter with goat-anti-human-alexa647 (5 µg/mL) with washes in between as the final was as described next. Slides were rinsed successively with PBS-T (0.1% Tween-20), PBS, and deionized water. After washing successively with PBS-T (0.1% Tween-20), PBS, and deionized water, a mixture of 10 µg/mL goat anti-mouse IgM-HRP (#1021-05, Southern Biotech) and 5 µg/mL donkey anti-goat IgG-Alexa555 (#A21432, Thermo Fisher

Scientific) in 40 μL PBS with 0.1% Tween-20 was incubated on the slide for 90 min in a humidified chamber. Afterward, the slides were rinsed successively with PBS-T (0.1% Tween-20), PBS, and deionized water. The arrays were dried by centrifugation and immediately scanned as described previously (1). Processing of the six replicates was performed by removing the highest and lowest replicate and subsequently calculating the mean value and standard deviation over the four remaining replicates.

### 10) NMR Spectra

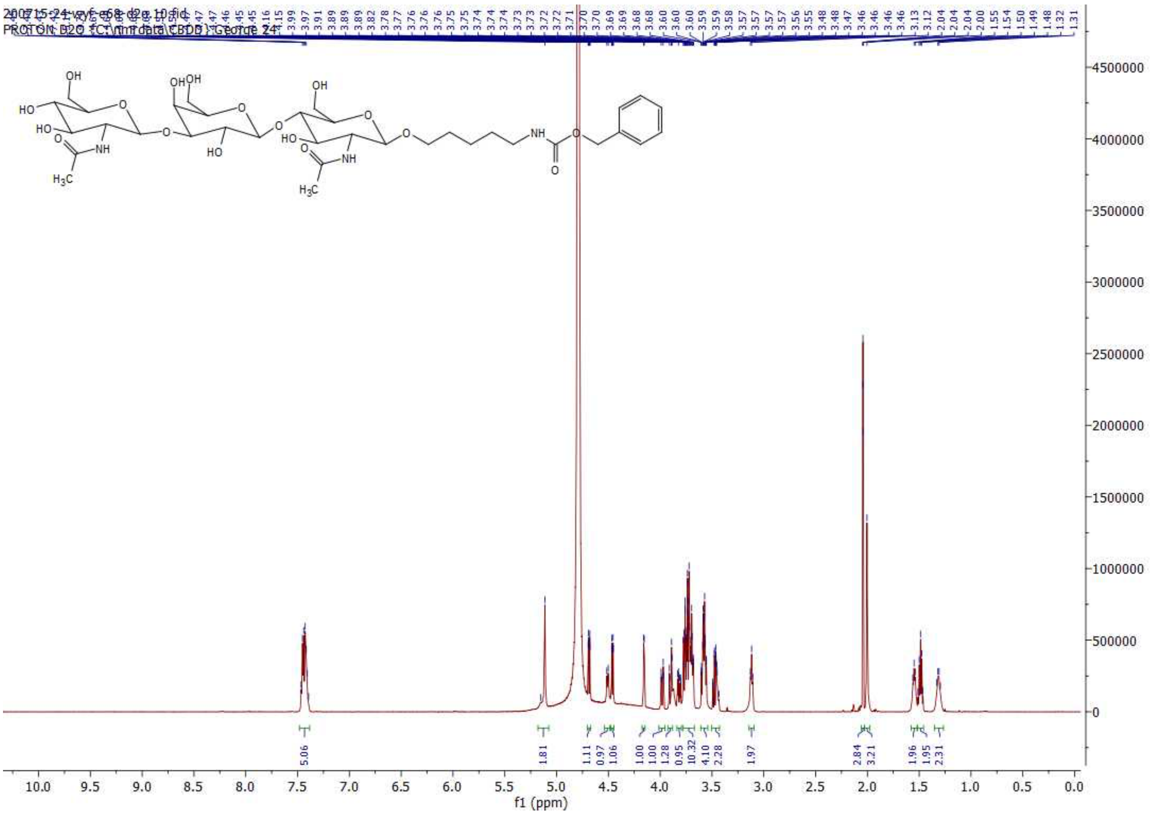

[1]H NMR of 2; 600MHz; $D_2O$

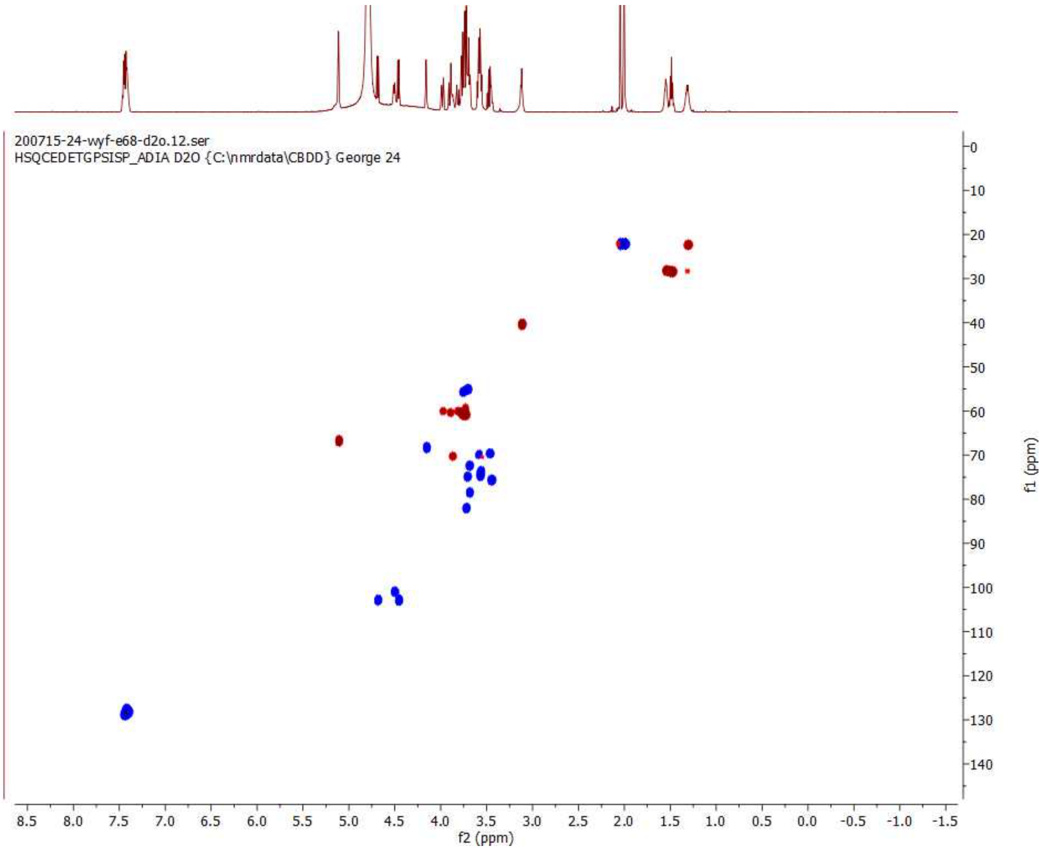

HSQC of 2; 600 MHz/150 MHz, $D_2O$

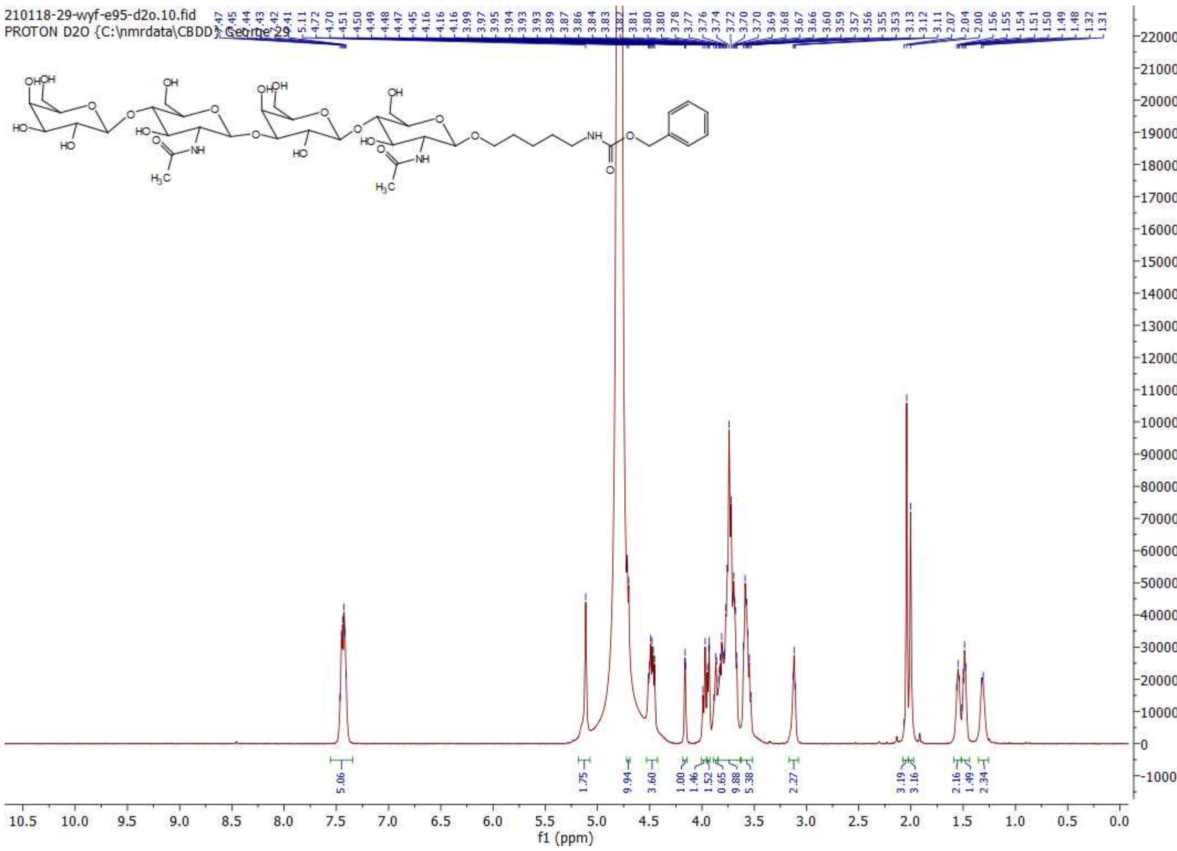

$^1$H NMR of 3; 600MHz; $D_2O$

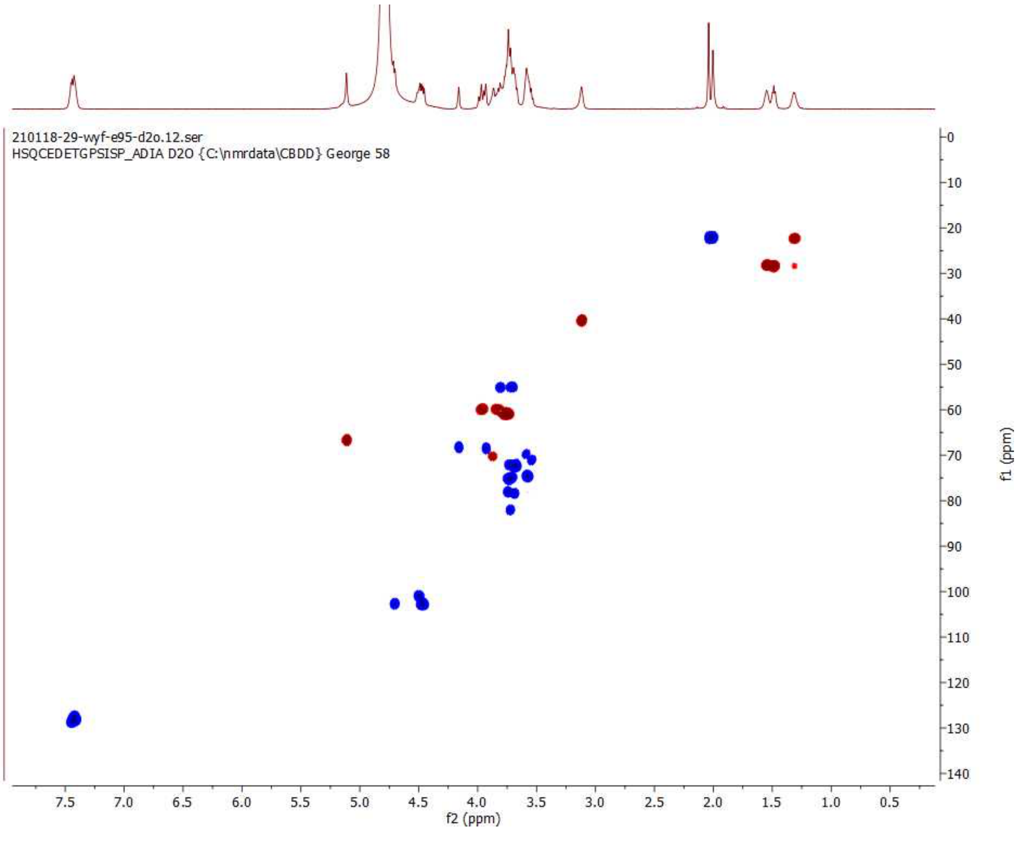

HSQC of 3; 600 MHz/150 MHz, $D_2O$

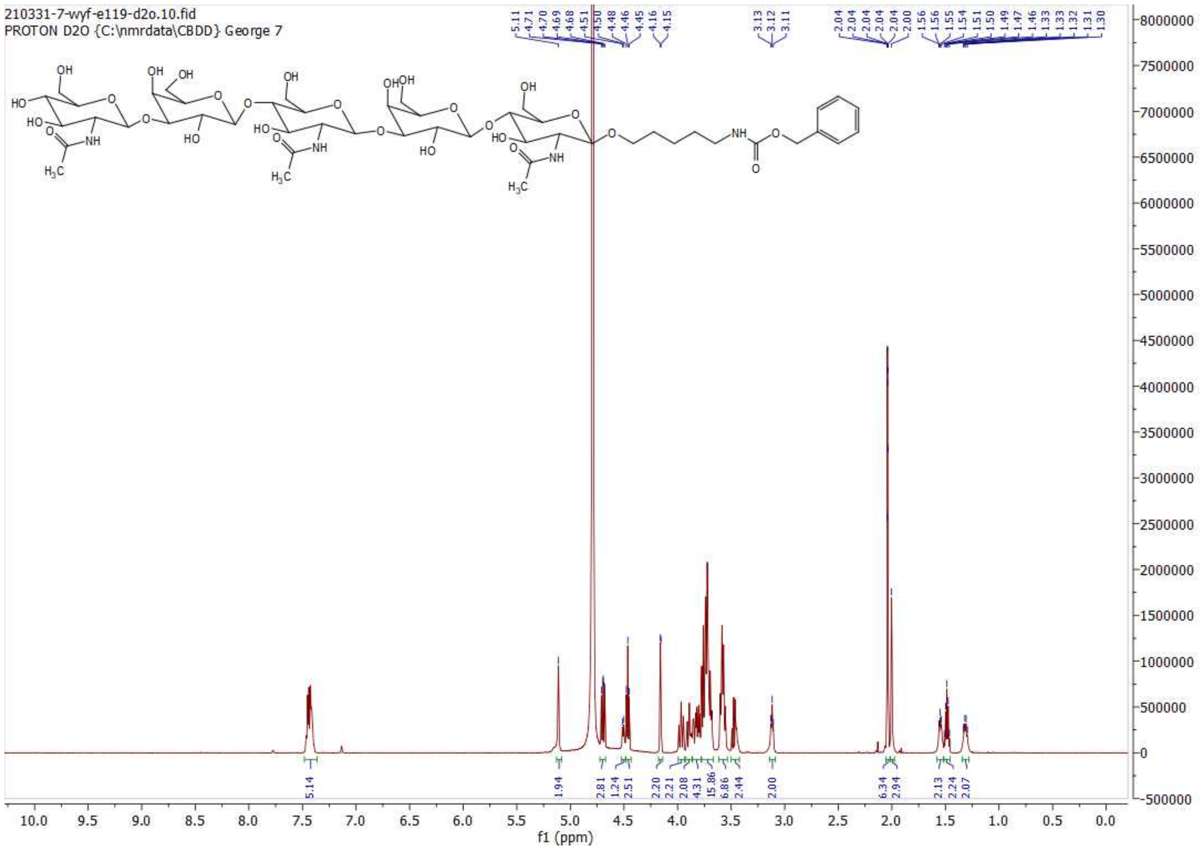

**[1]H NMR of 4; 600MHz; $D_2O$**

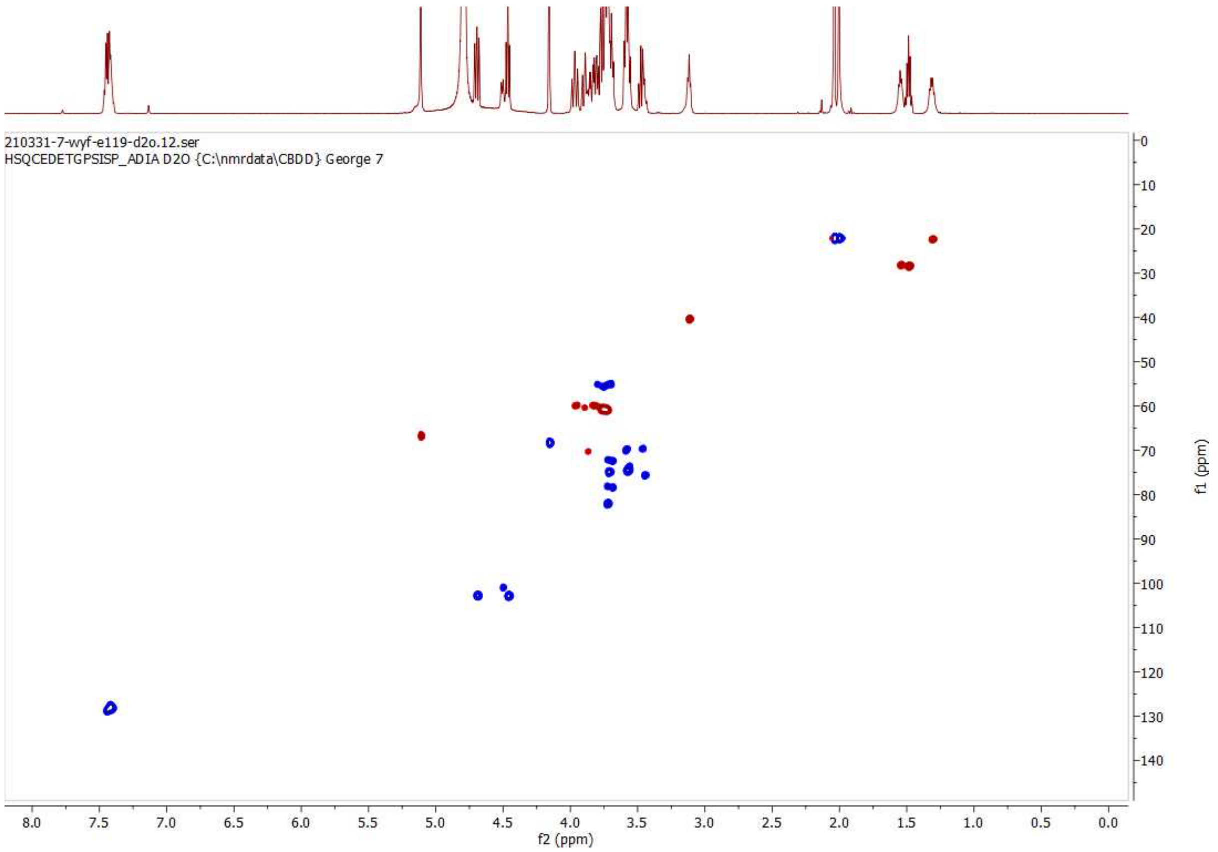

**HSQC of 4; 600 MHz/150 MHz, $D_2O$**

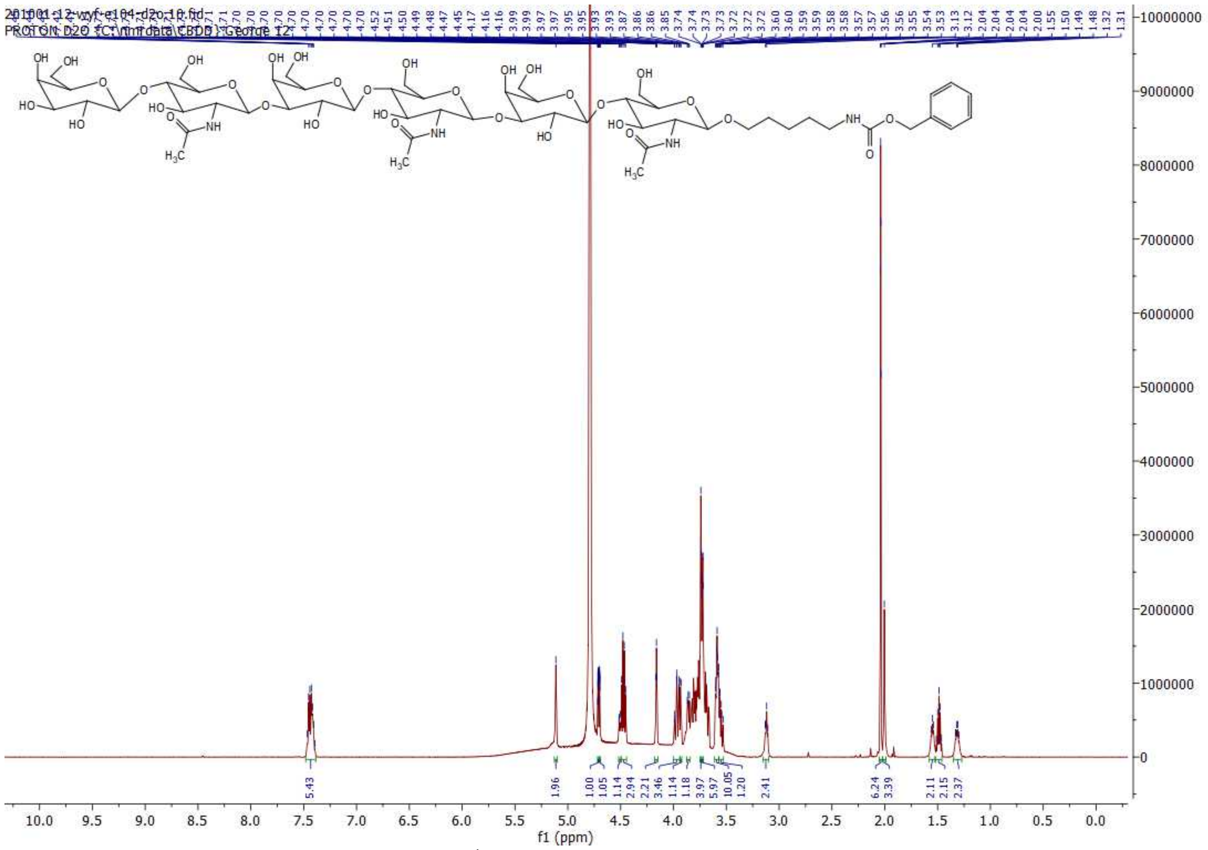

**[1]H NMR of 5; 600MHz; D₂O**

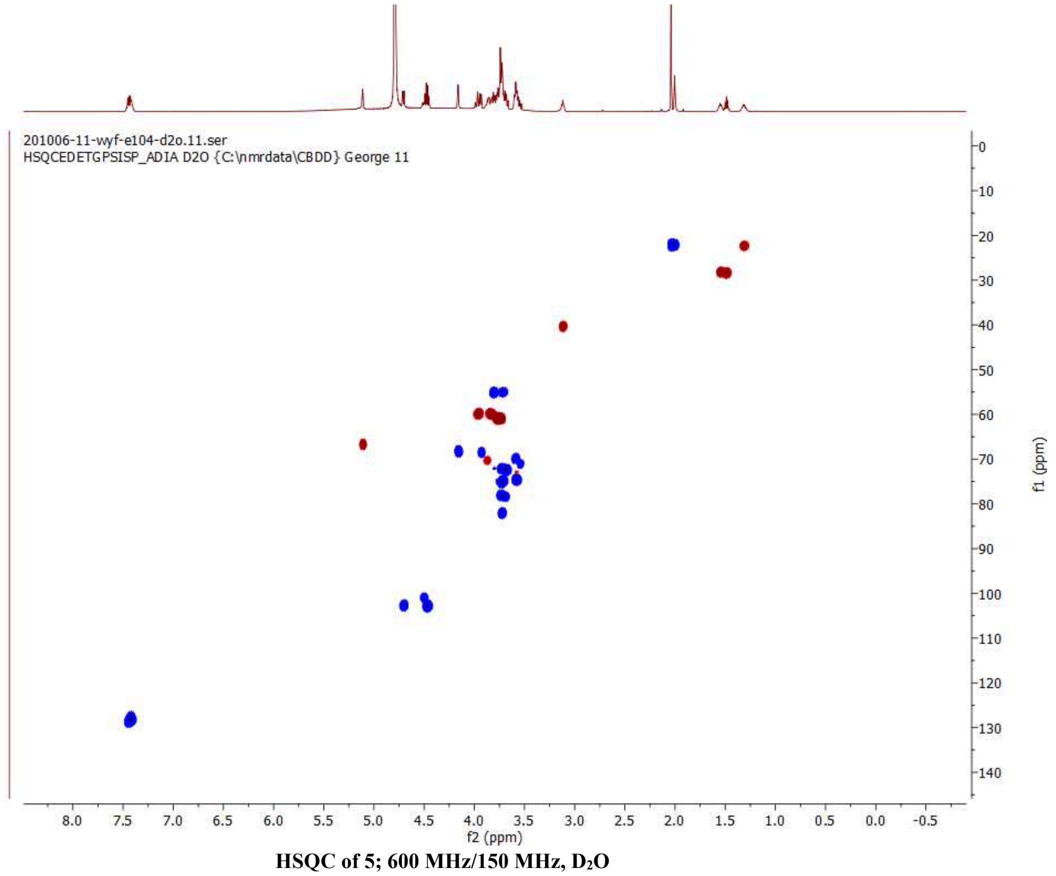

**HSQC of 5; 600 MHz/150 MHz, D₂O**

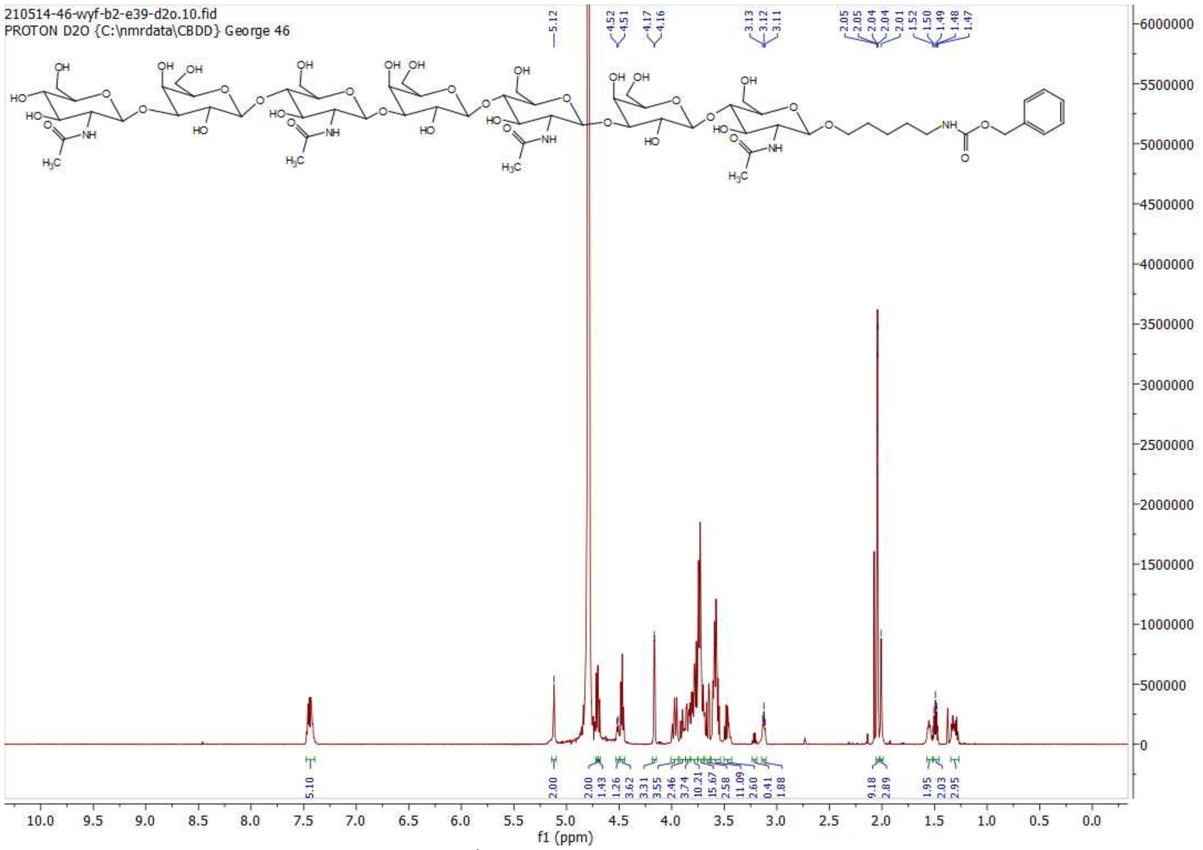

$^1H$ NMR of 6; 600MHz; $D_2O$

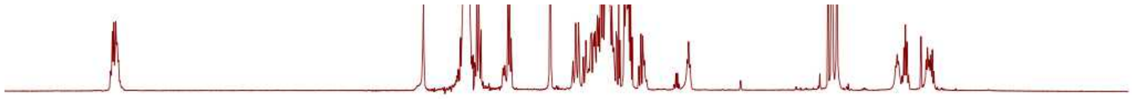

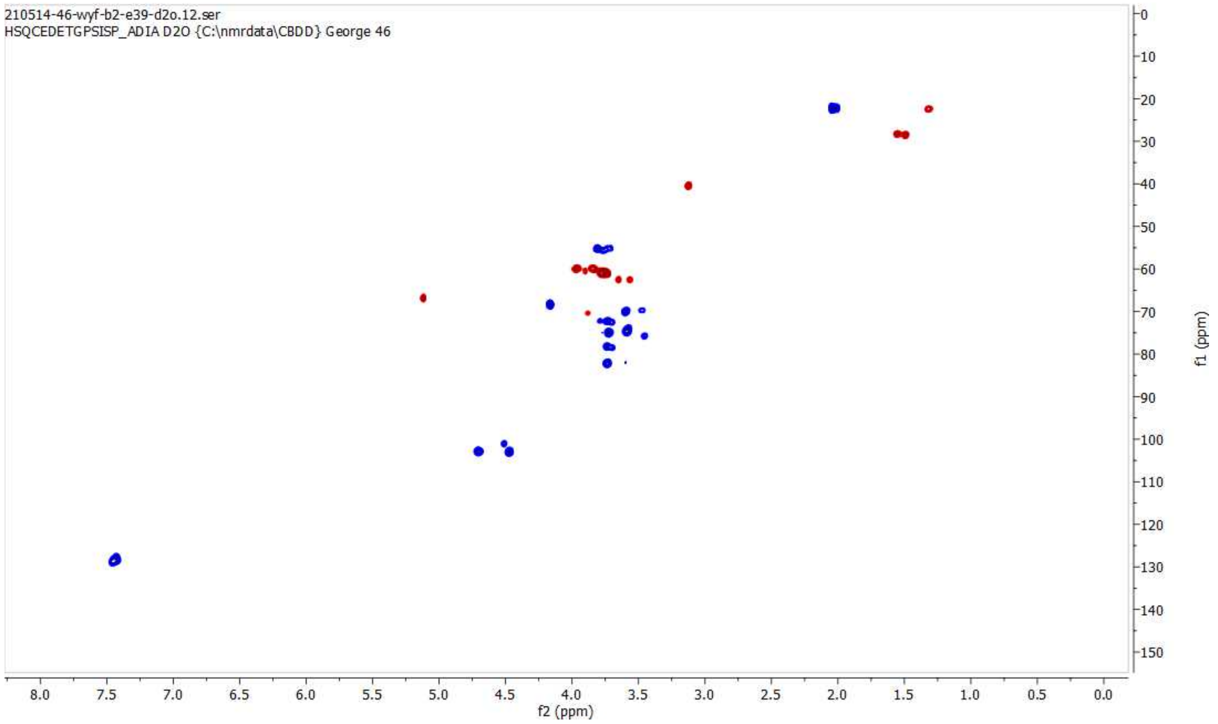

HSQC of 6; 600 MHz/150 MHz, $D_2O$

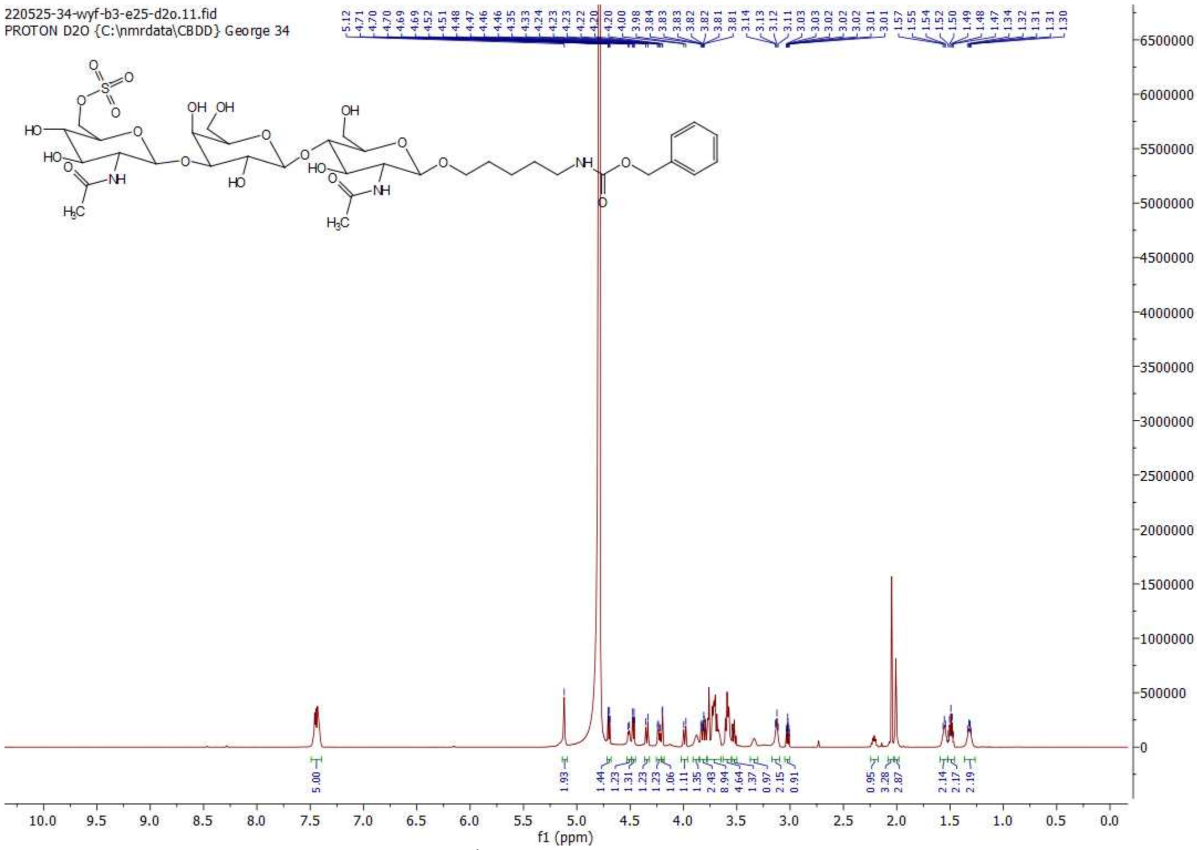

**[1]H NMR of 7; 600MHz; $D_2O$**

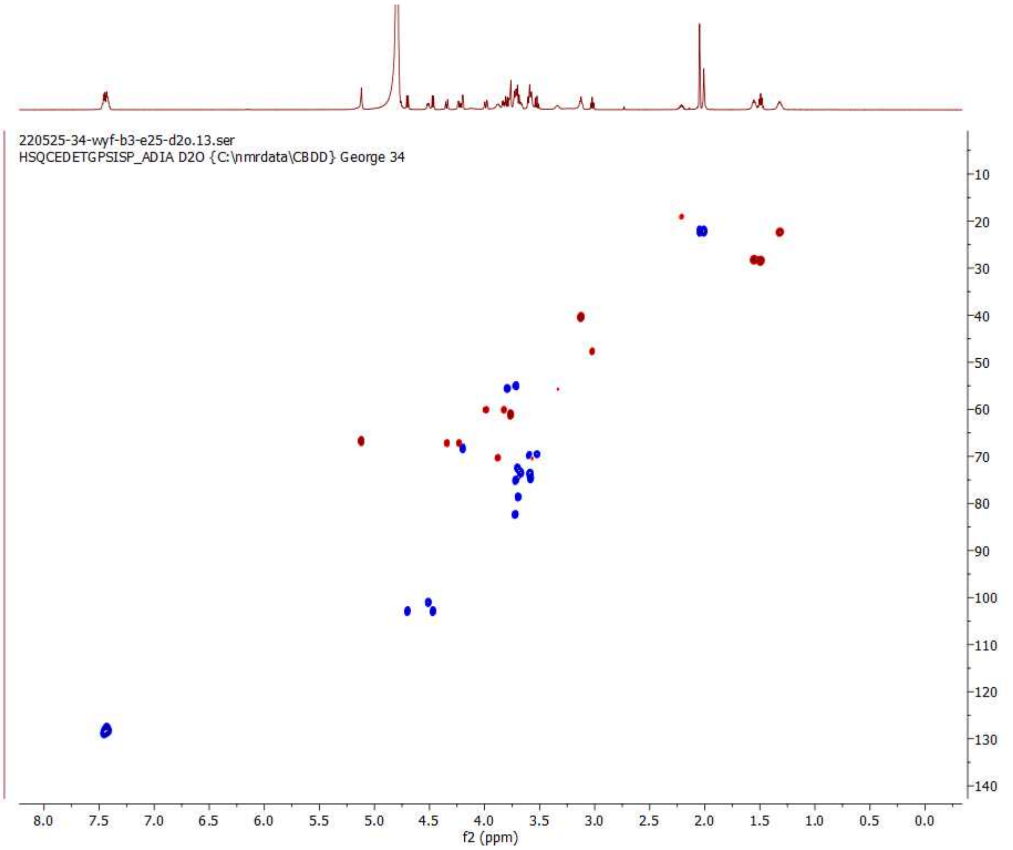

**HSQC of 7; 600 MHz/150 MHz, $D_2O$**

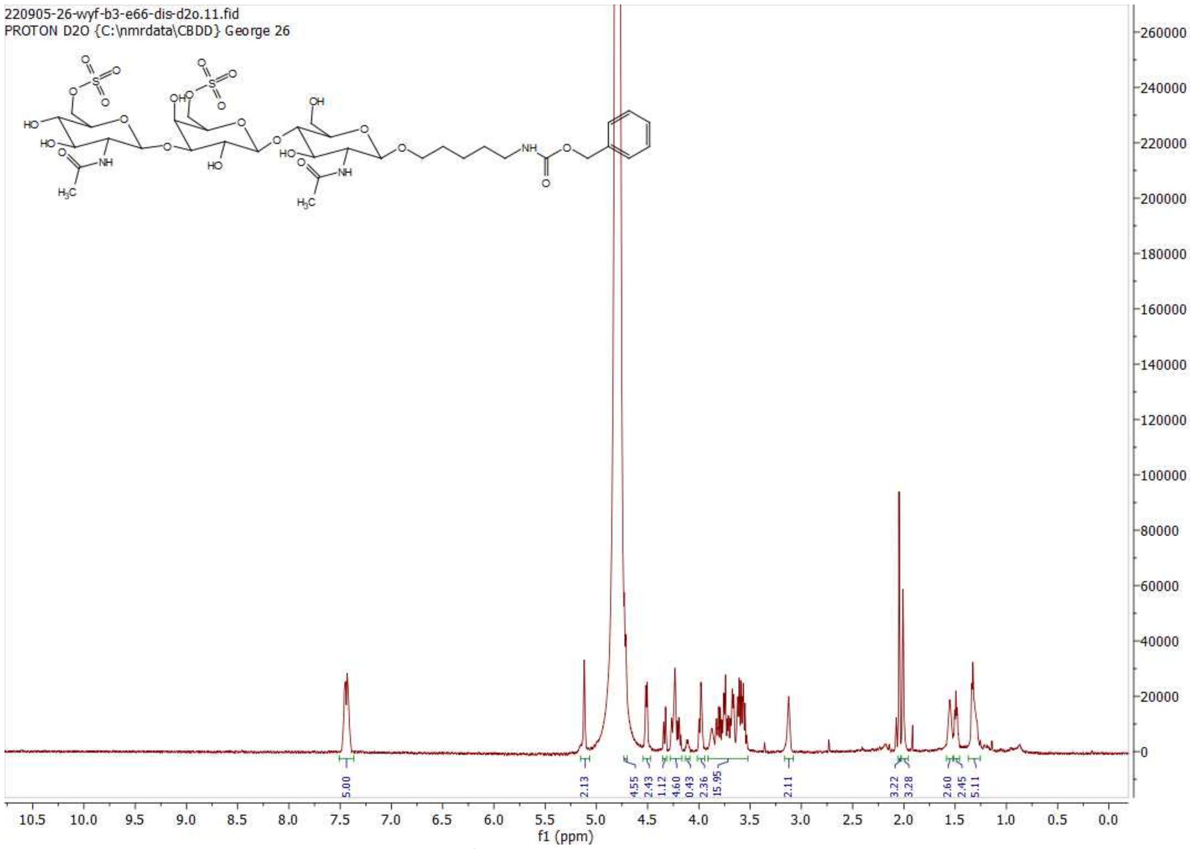

[1]H NMR of 7’; 600MHz; $D_2O$

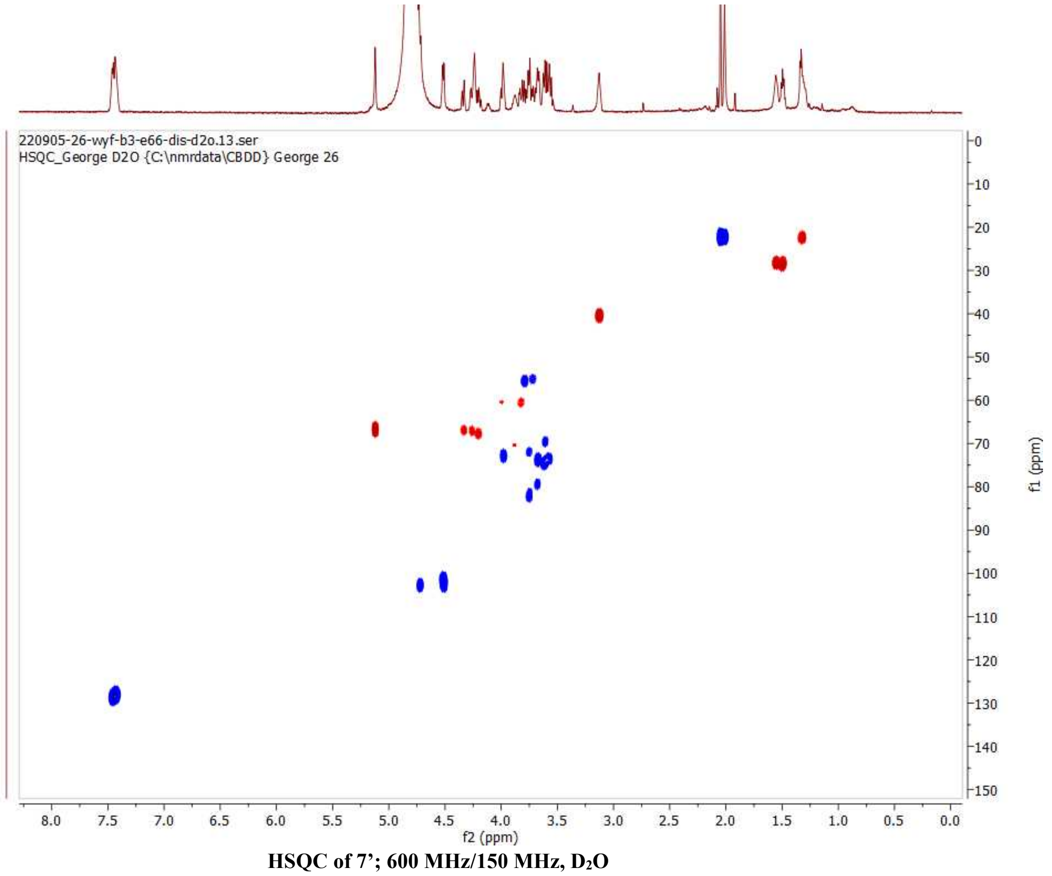

HSQC of 7’; 600 MHz/150 MHz, $D_2O$

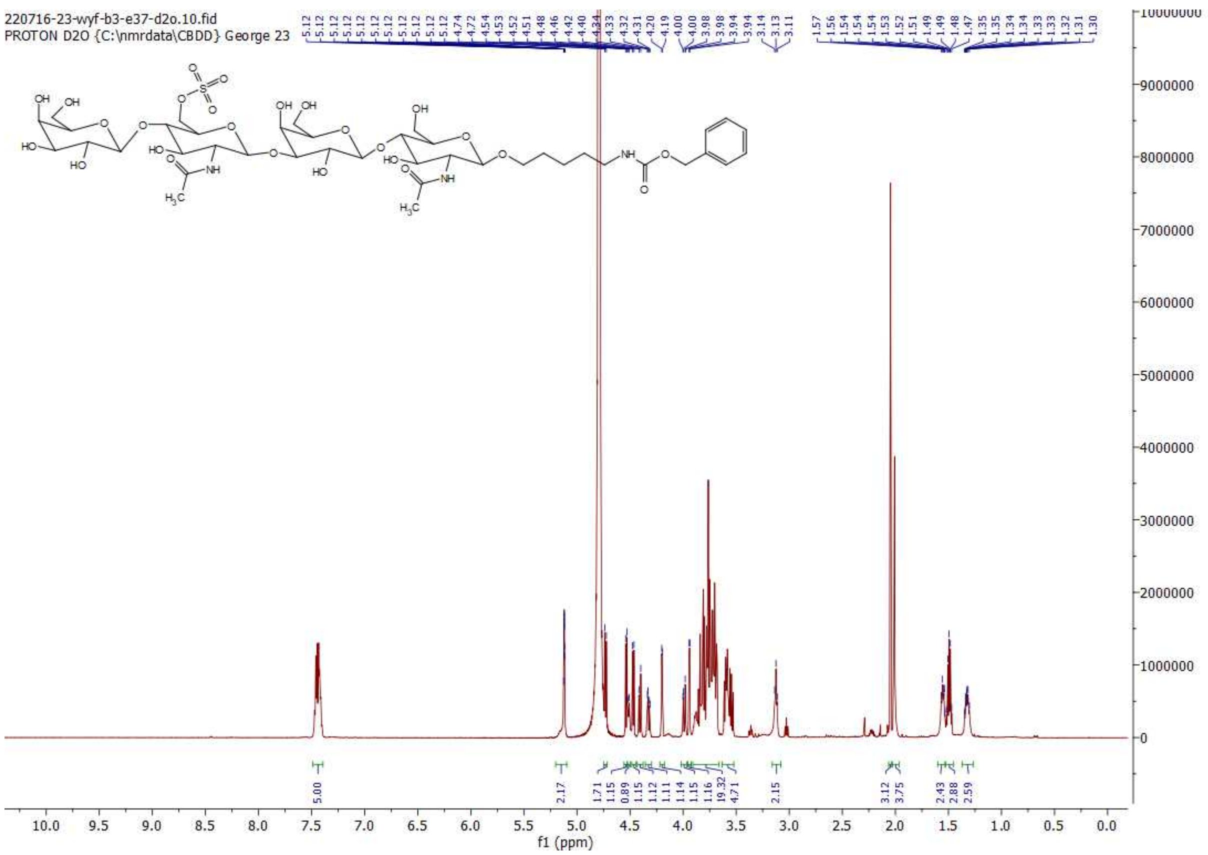

**[1]H NMR of 8; 600MHz; $D_2O$**

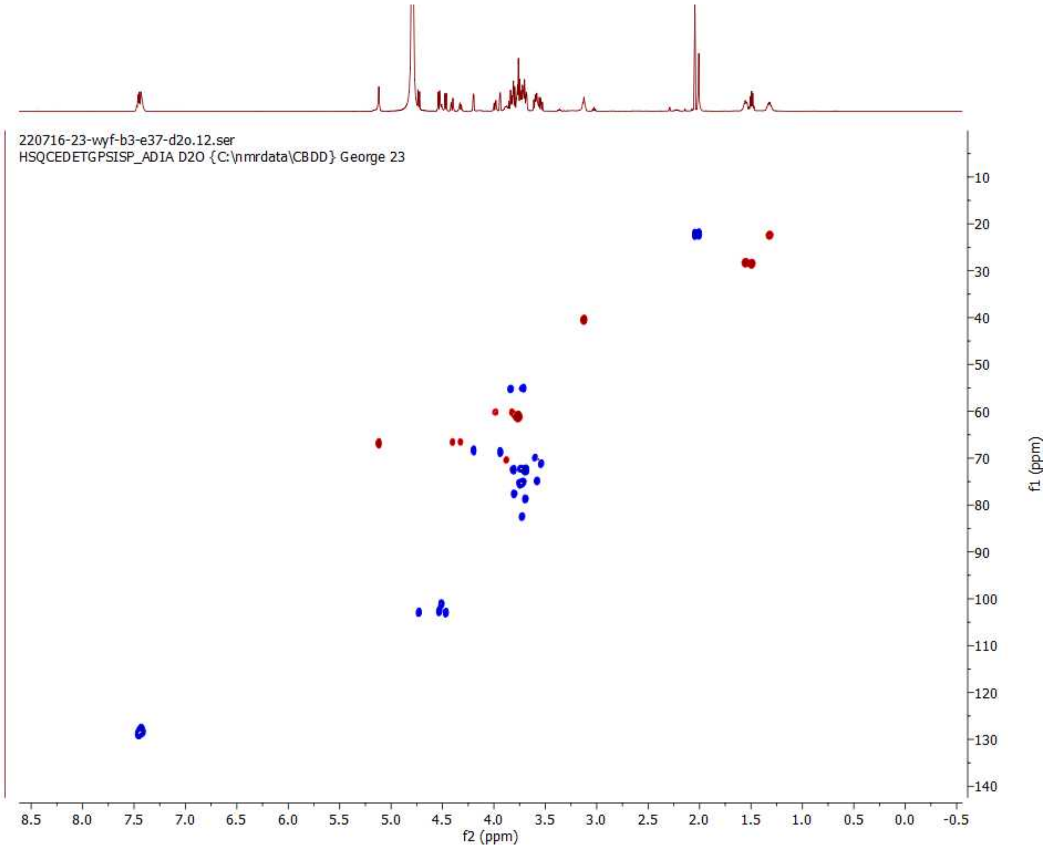

**HSQC of 8; 600 MHz/150 MHz, $D_2O$**

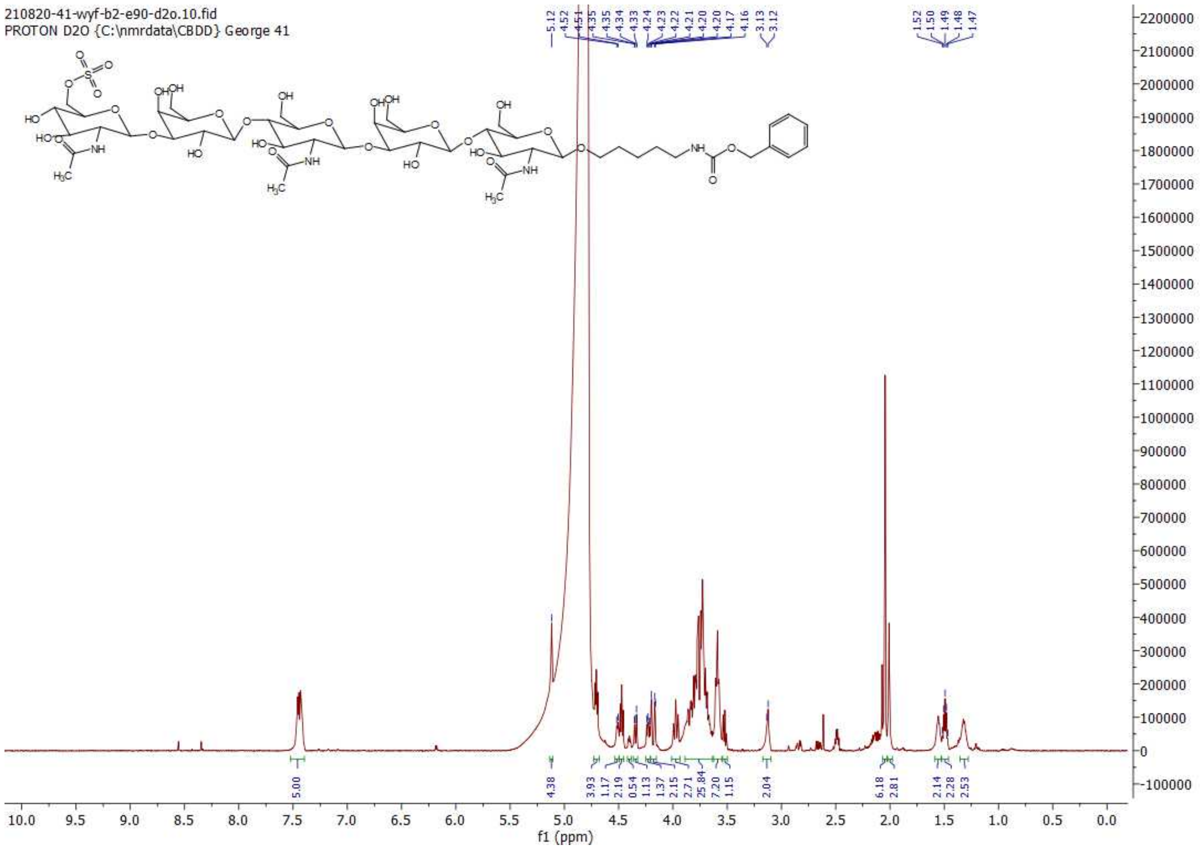

$^1$H NMR of 9; 600MHz; $D_2O$

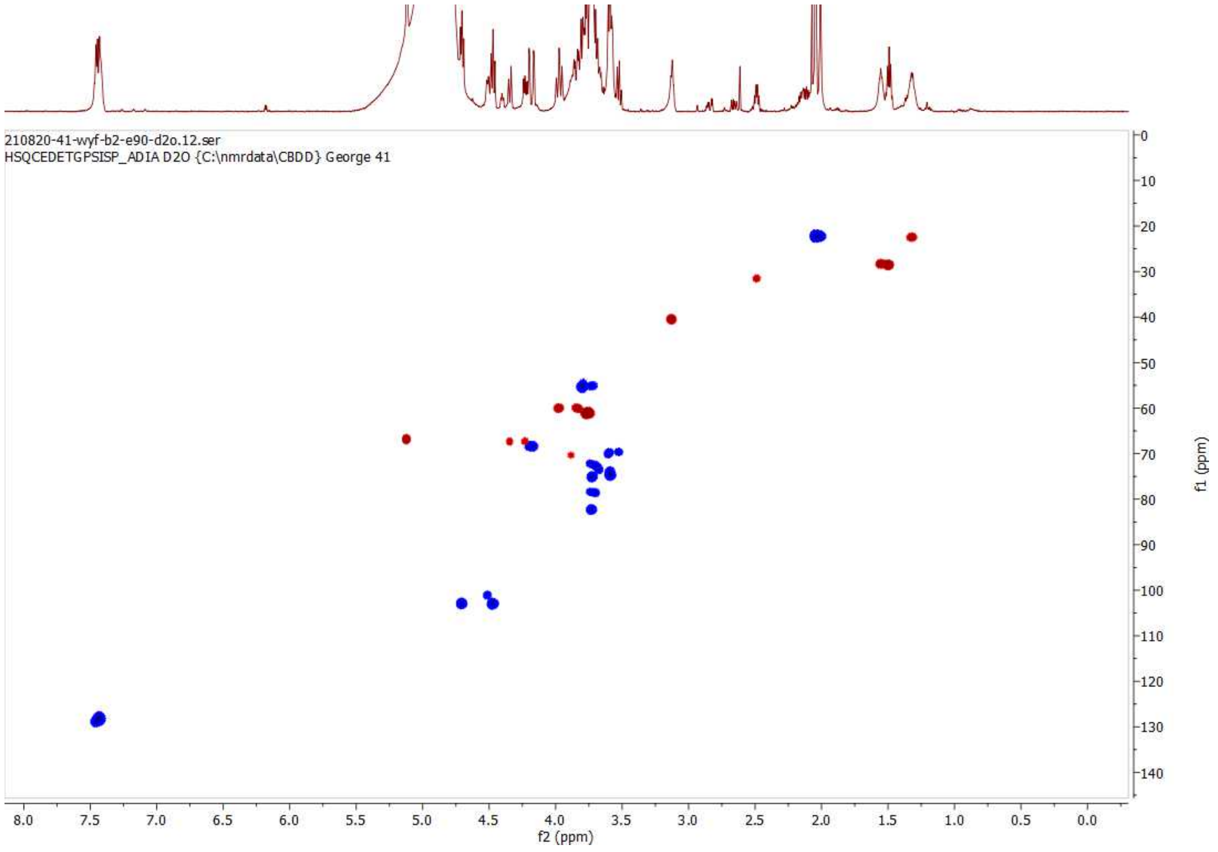

HSQC of 9; 600 MHz/150 MHz, $D_2O$

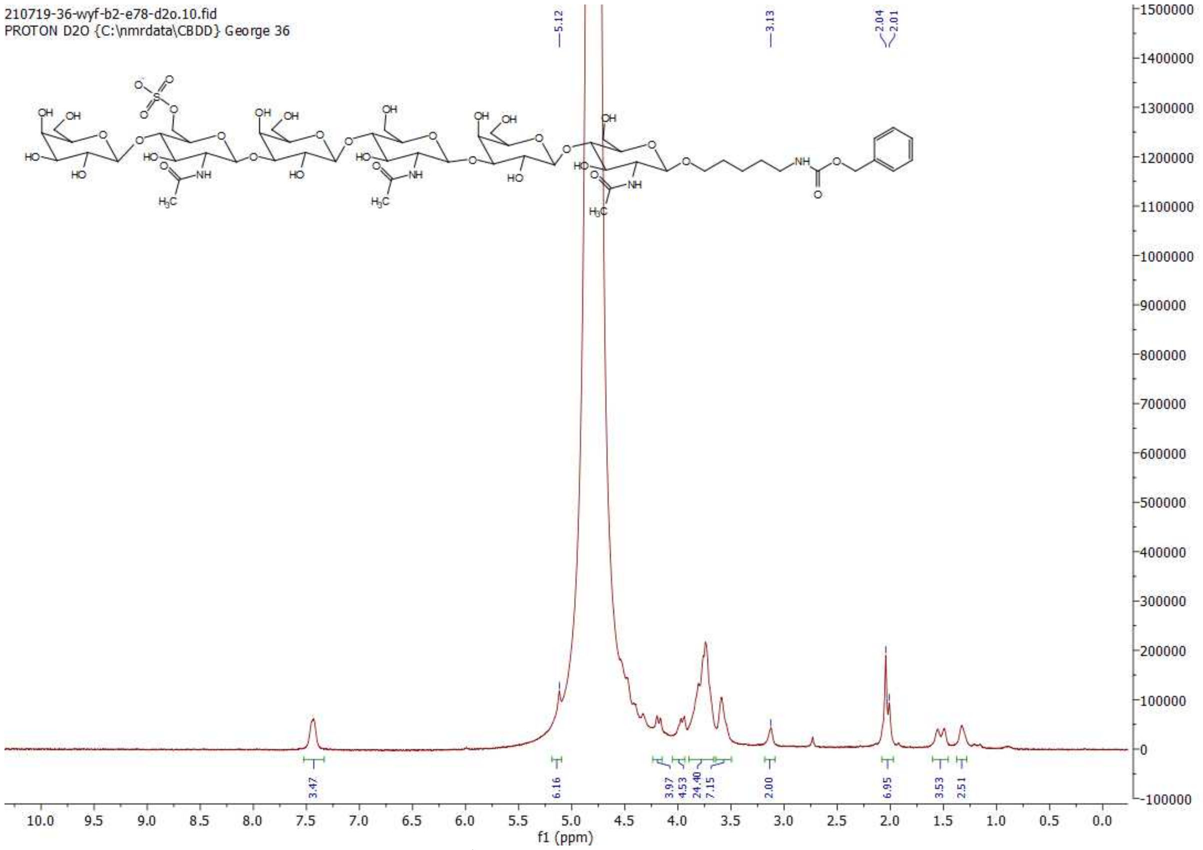

$^{1}$H NMR of 10; 600MHz; $D_2O$

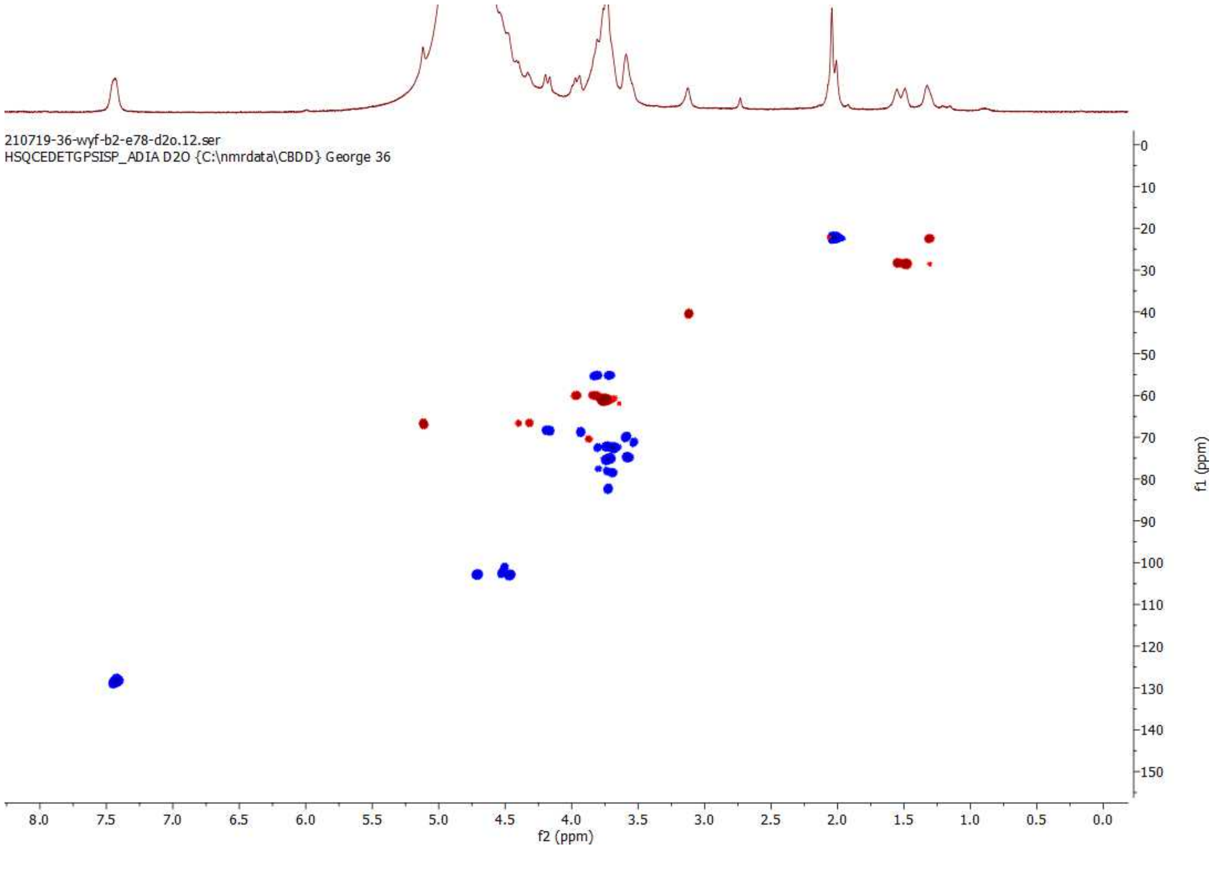

HSQC of 10; 600 MHz/150 MHz, $D_2O$

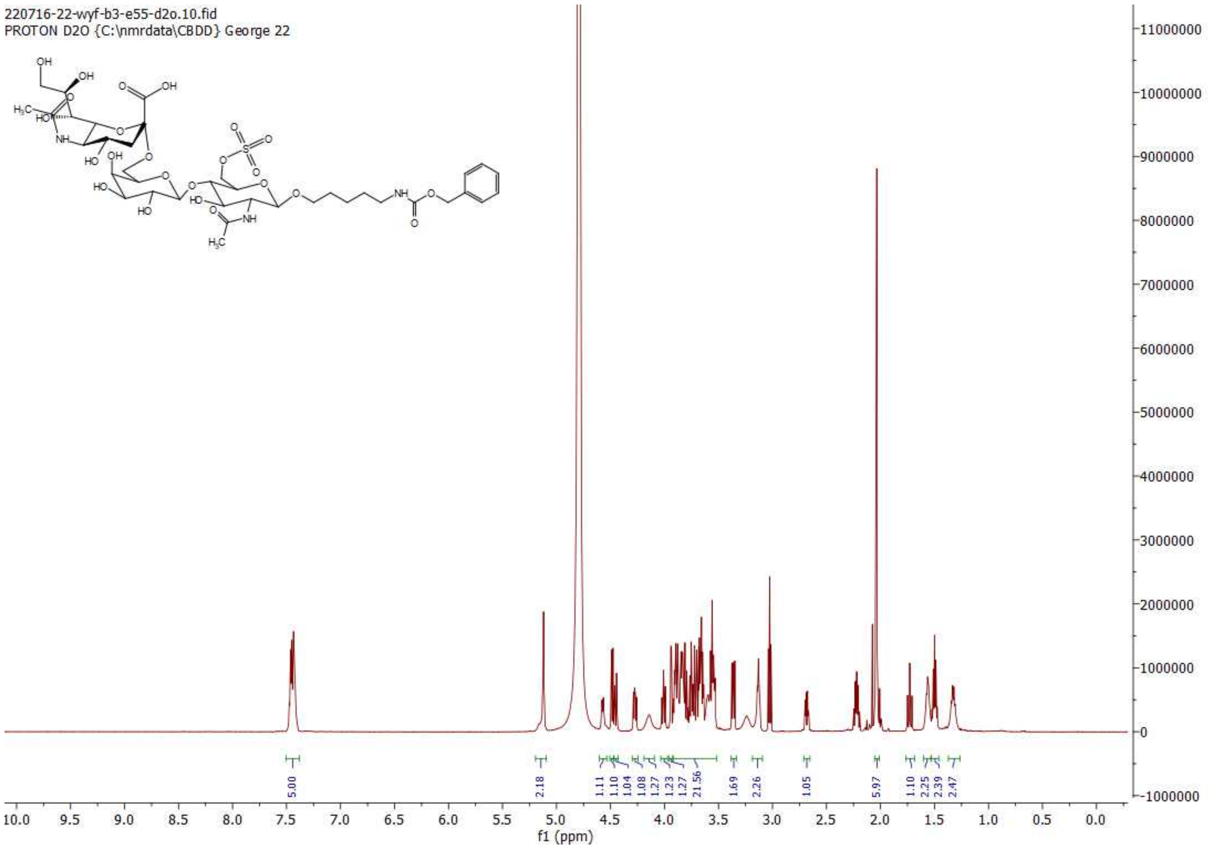

**[1]H NMR of 12; 600MHz; $D_2O$**

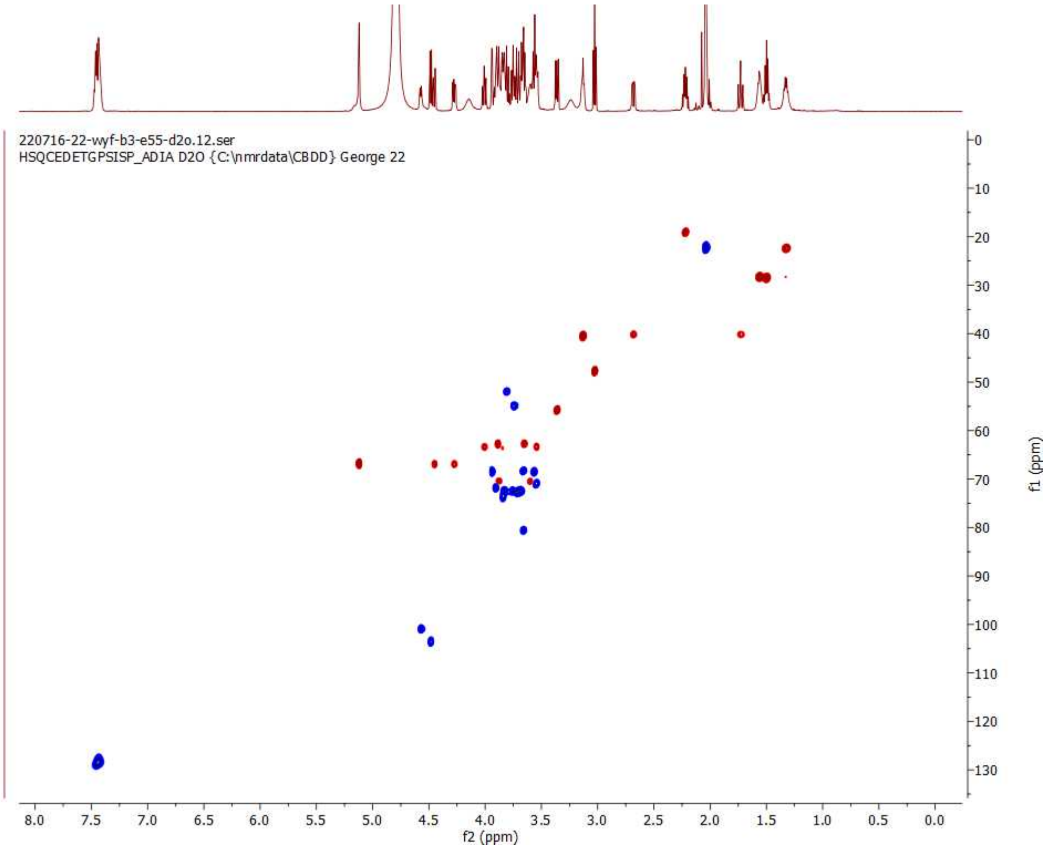

**HSQC of 12; 600 MHz/150 MHz, $D_2O$**

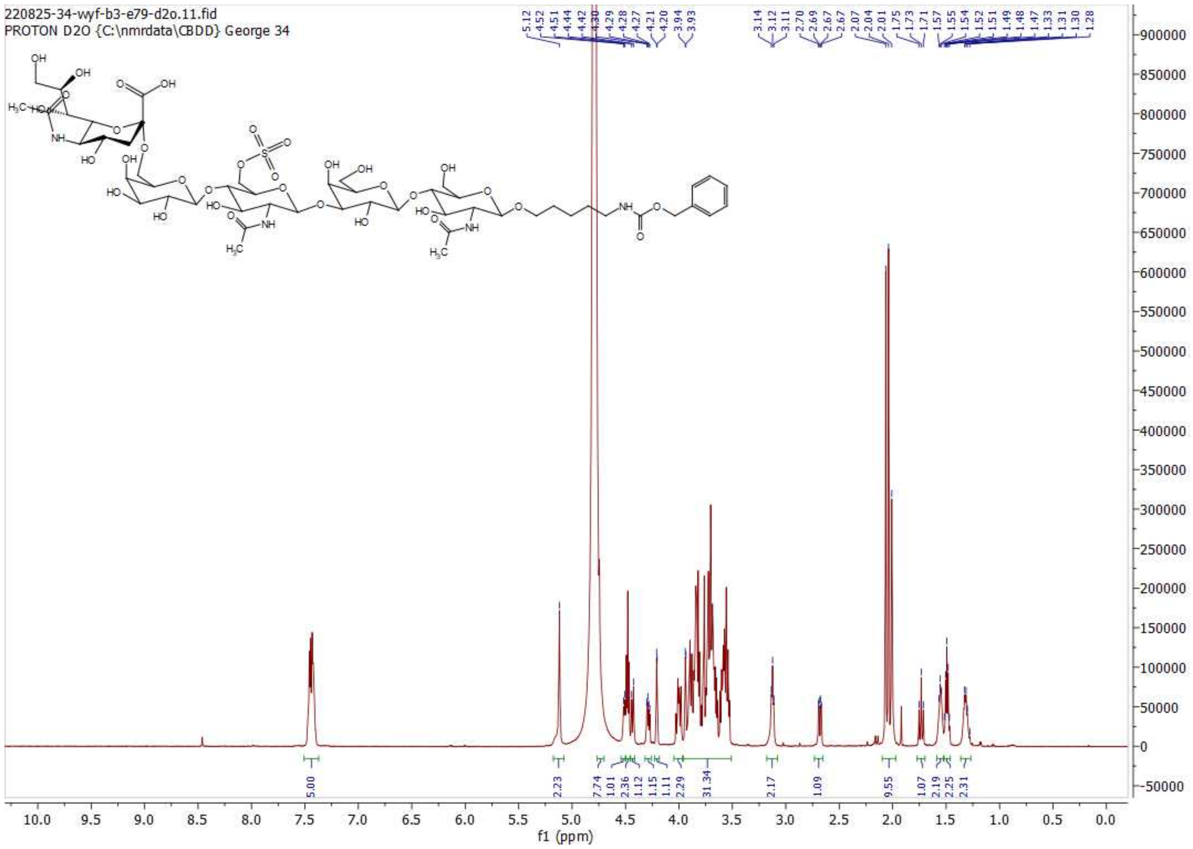

[1]H NMR of 13; 600MHz; $D_2O$

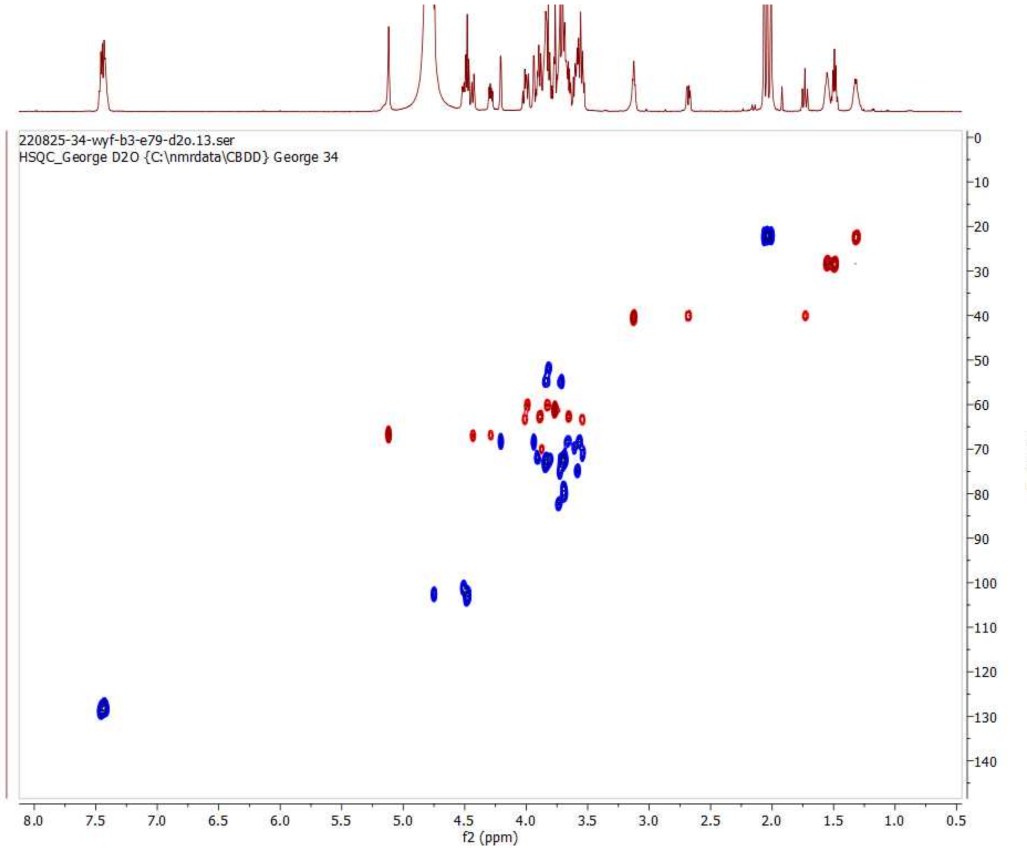

HSQC of 13; 600 MHz/150 MHz, $D_2O$

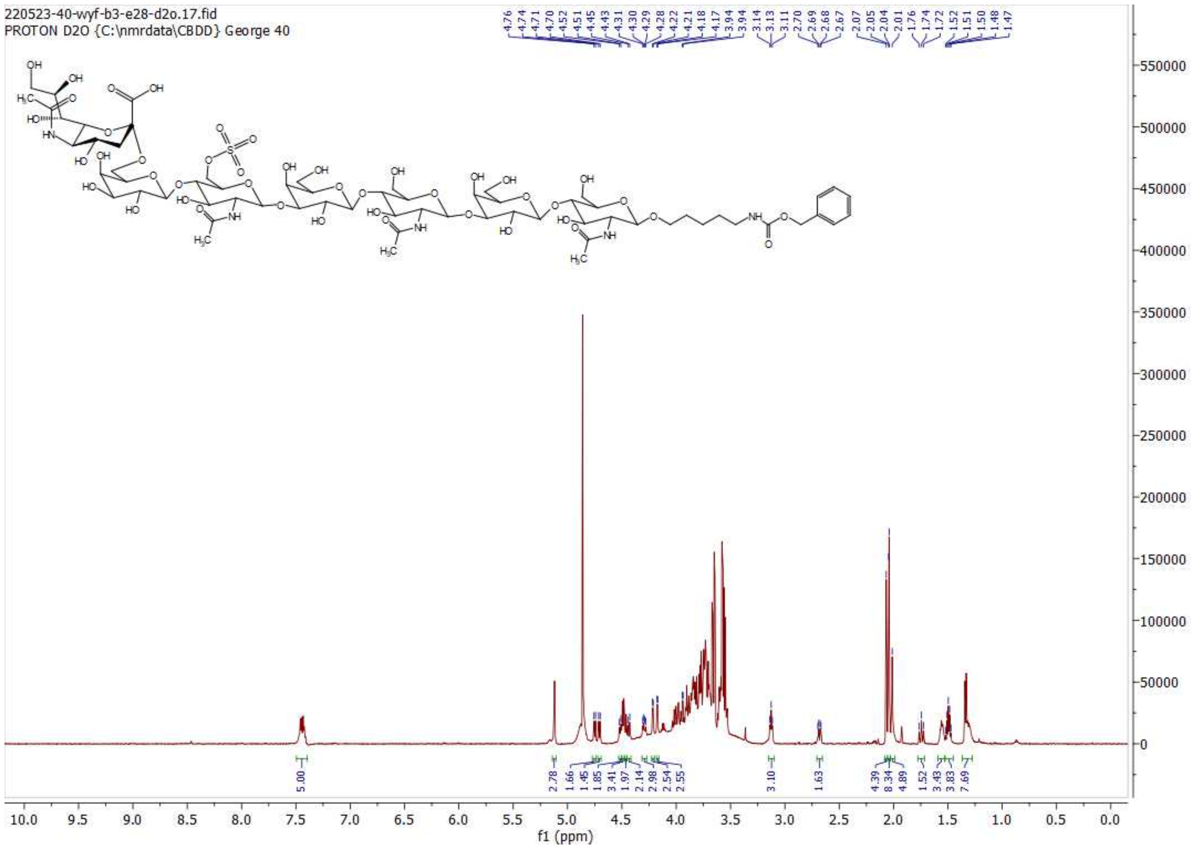

**[1]H NMR of 14; 600MHz; $D_2O$**

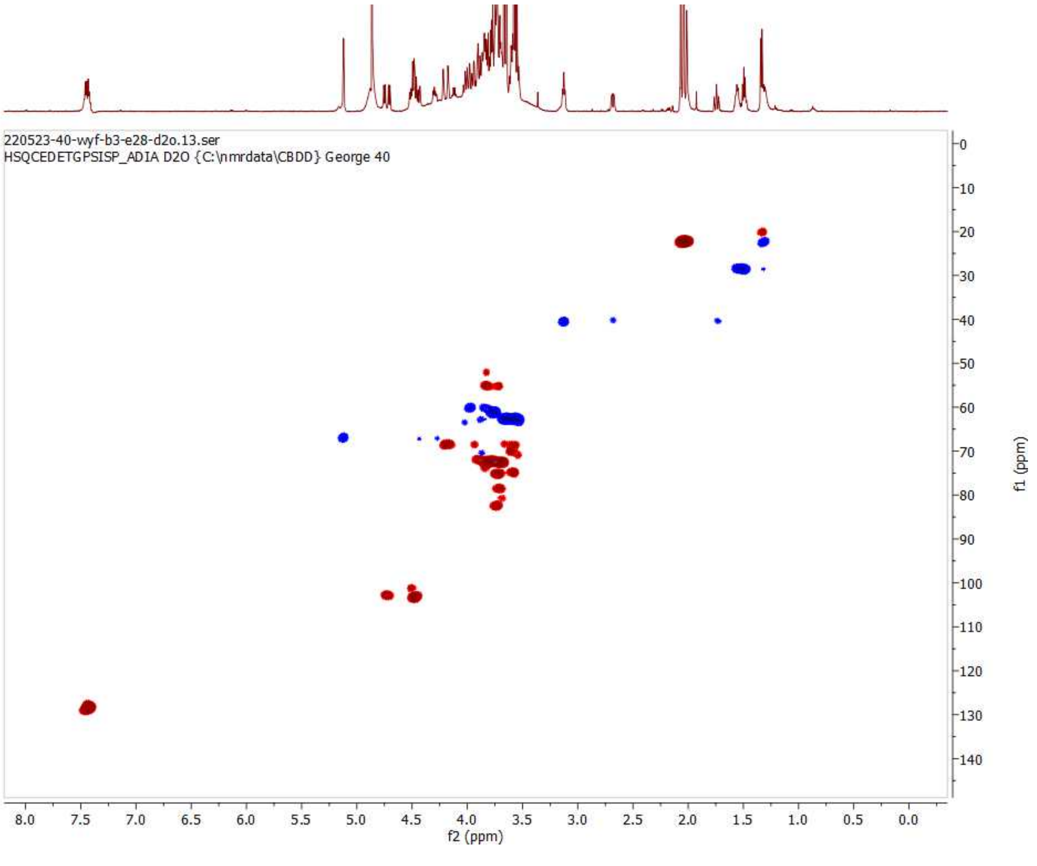

**HSQC of 14; 600 MHz/150 MHz, $D_2O$**

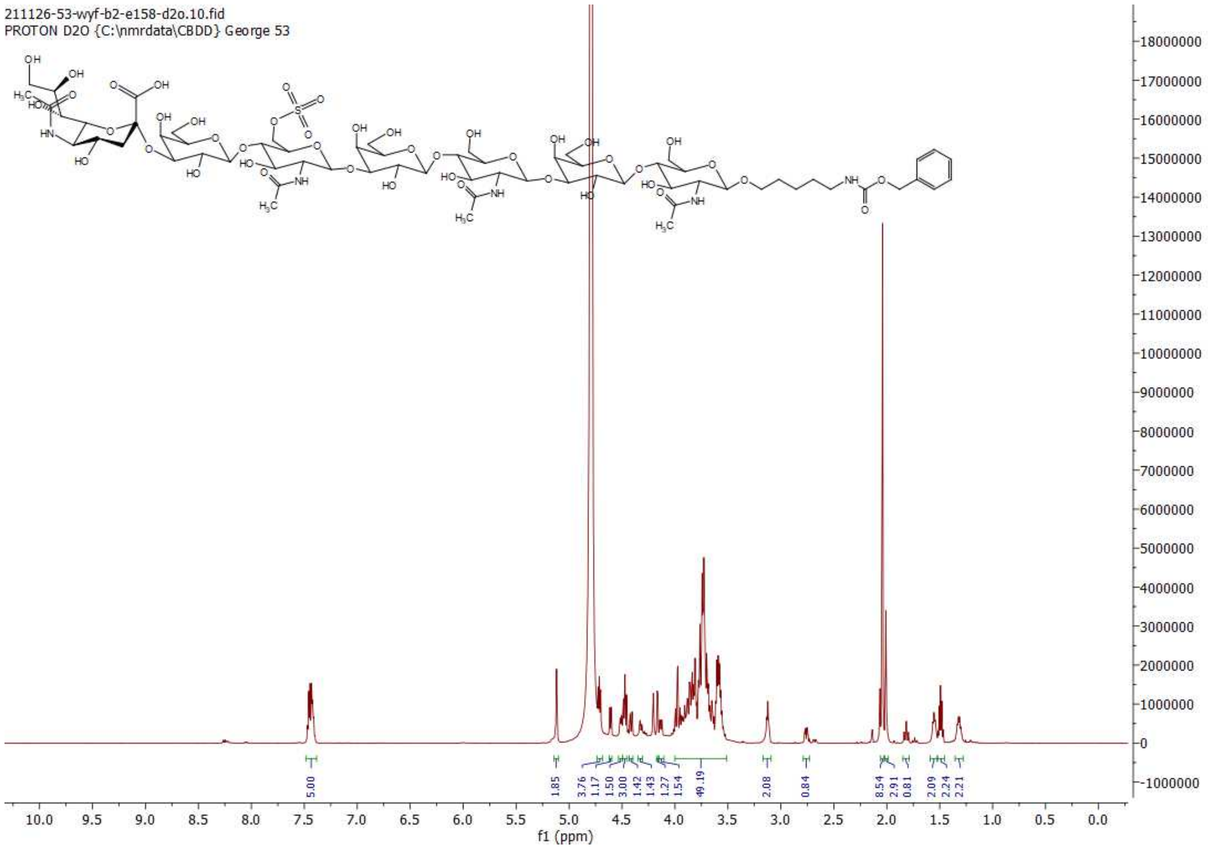

**[1]H NMR of 15; 600MHz; $D_2O$**

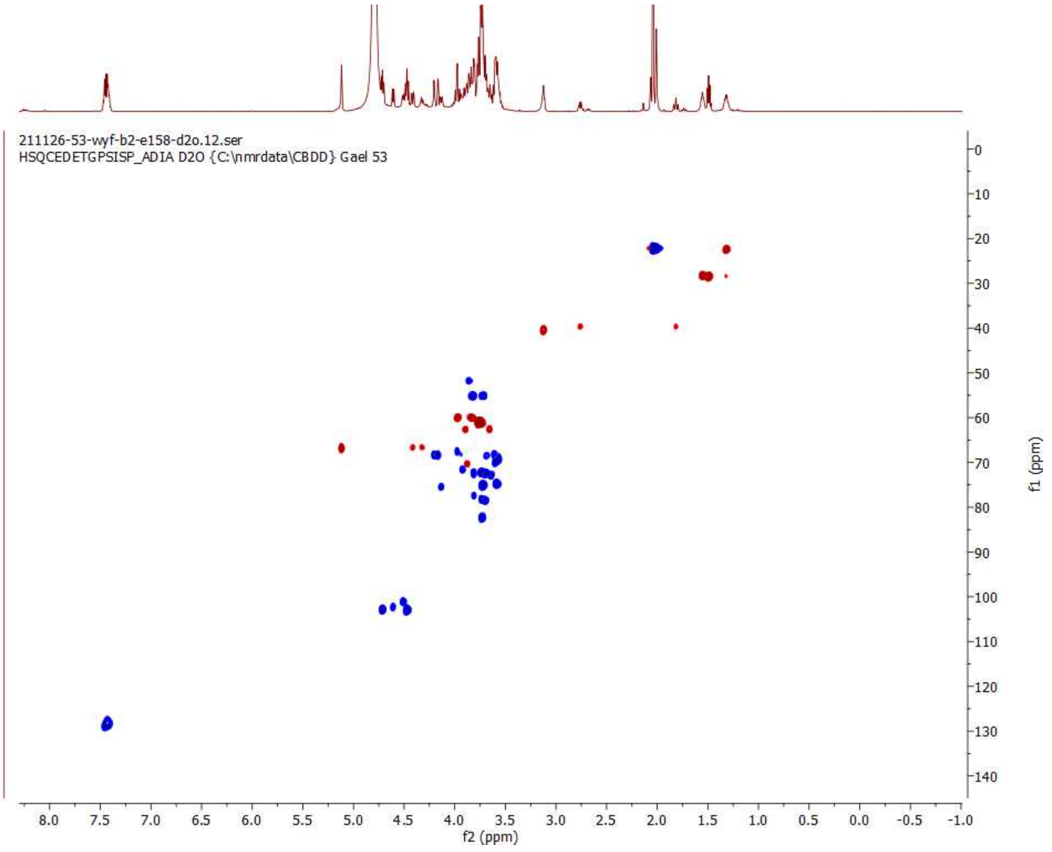

**HSQC of 15; 600 MHz/150 MHz, $D_2O$**

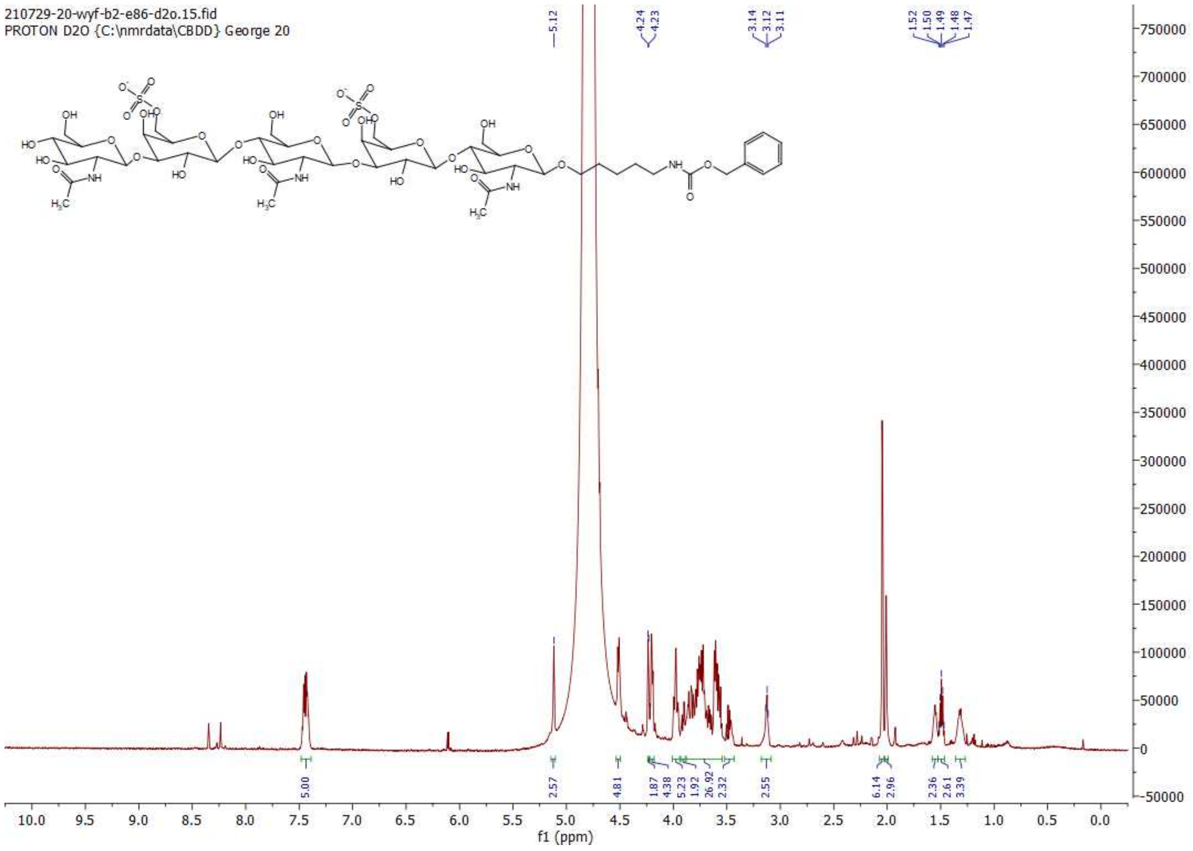

$^{1}$H NMR of 16; 600MHz; $D_2O$

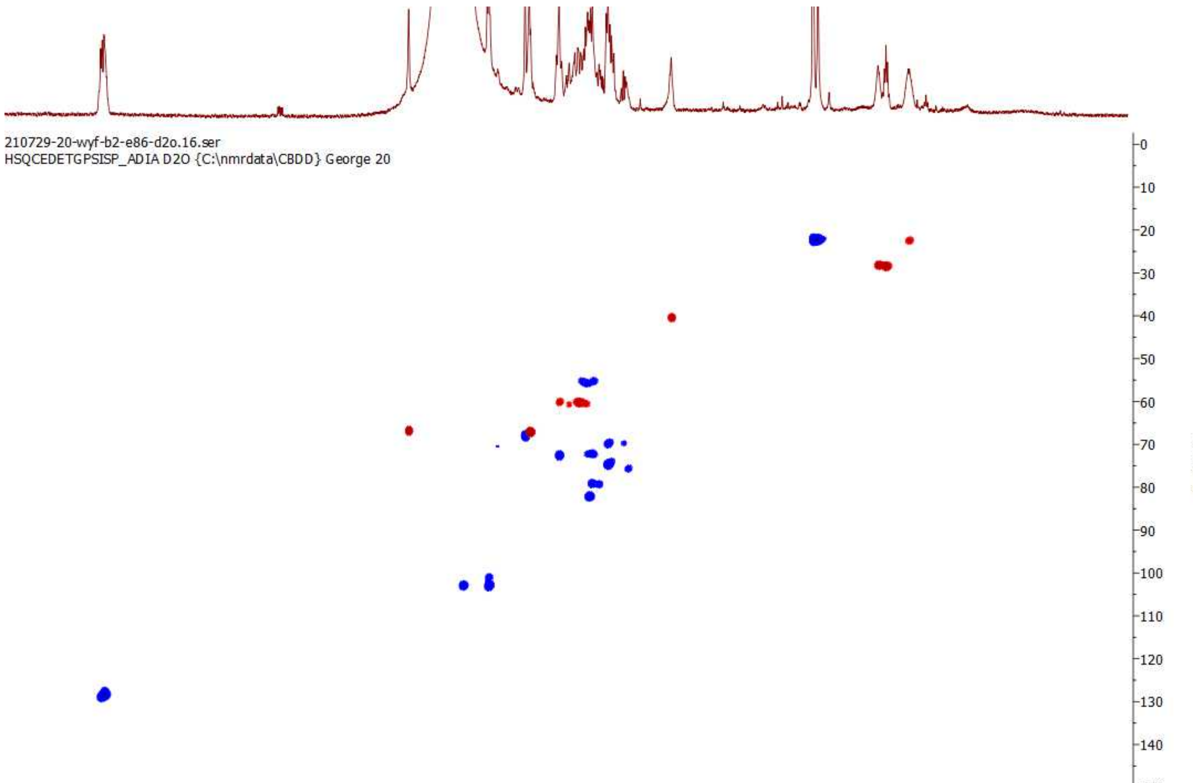

HSQC of 16; 600 MHz/150 MHz, $D_2O$

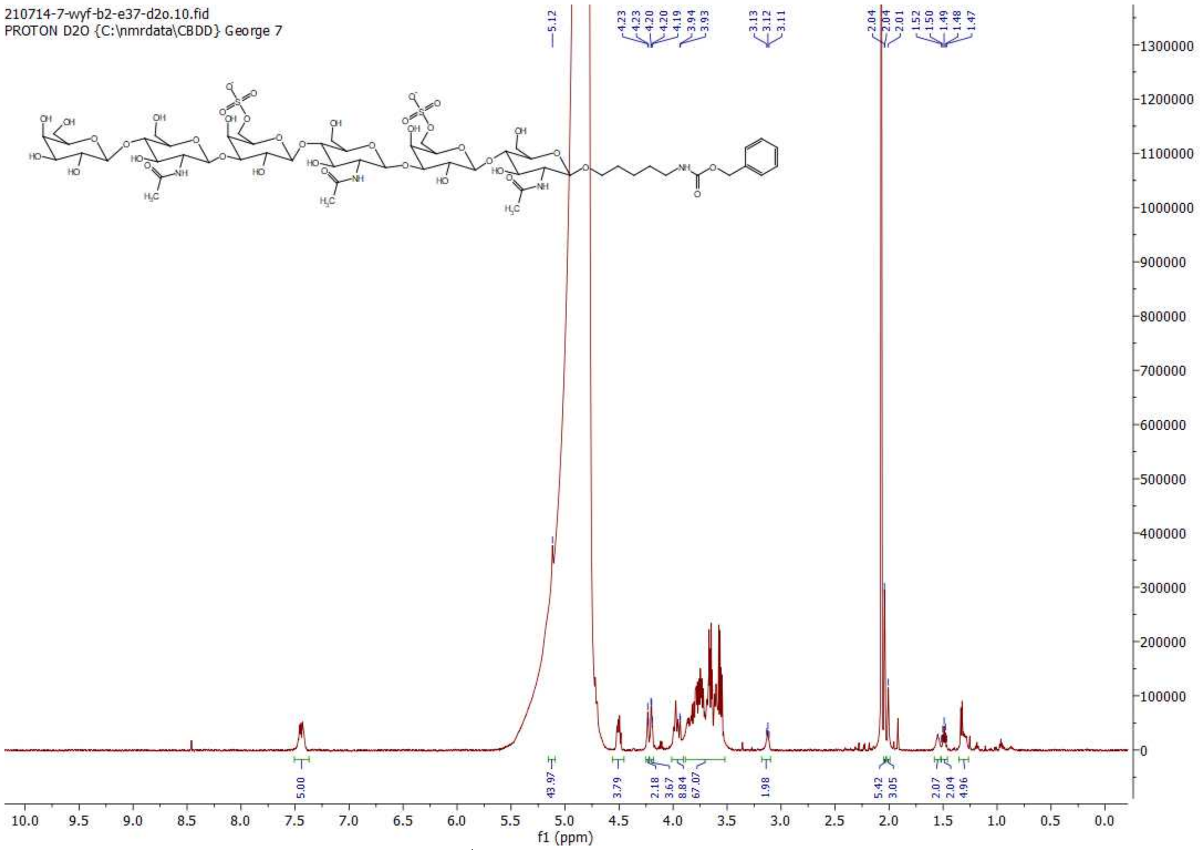

**[1]H NMR of 17; 600MHz; $D_2O$**

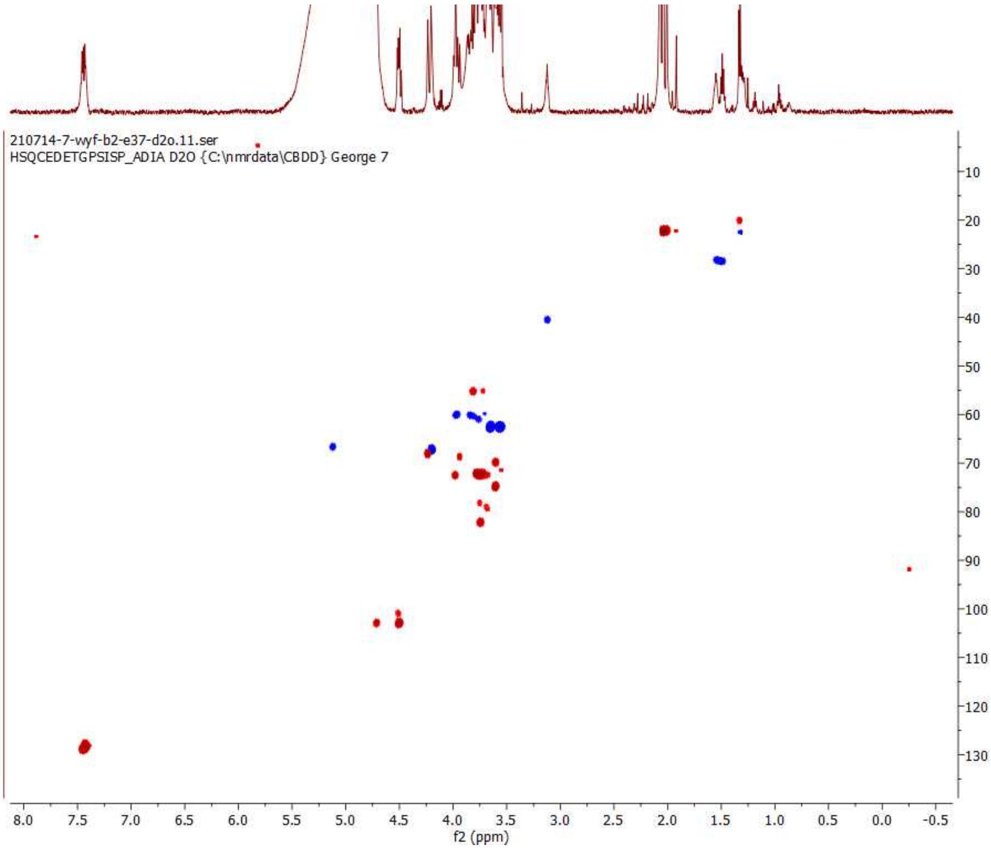

**HSQC of 17; 600 MHz/150 MHz, $D_2O$**

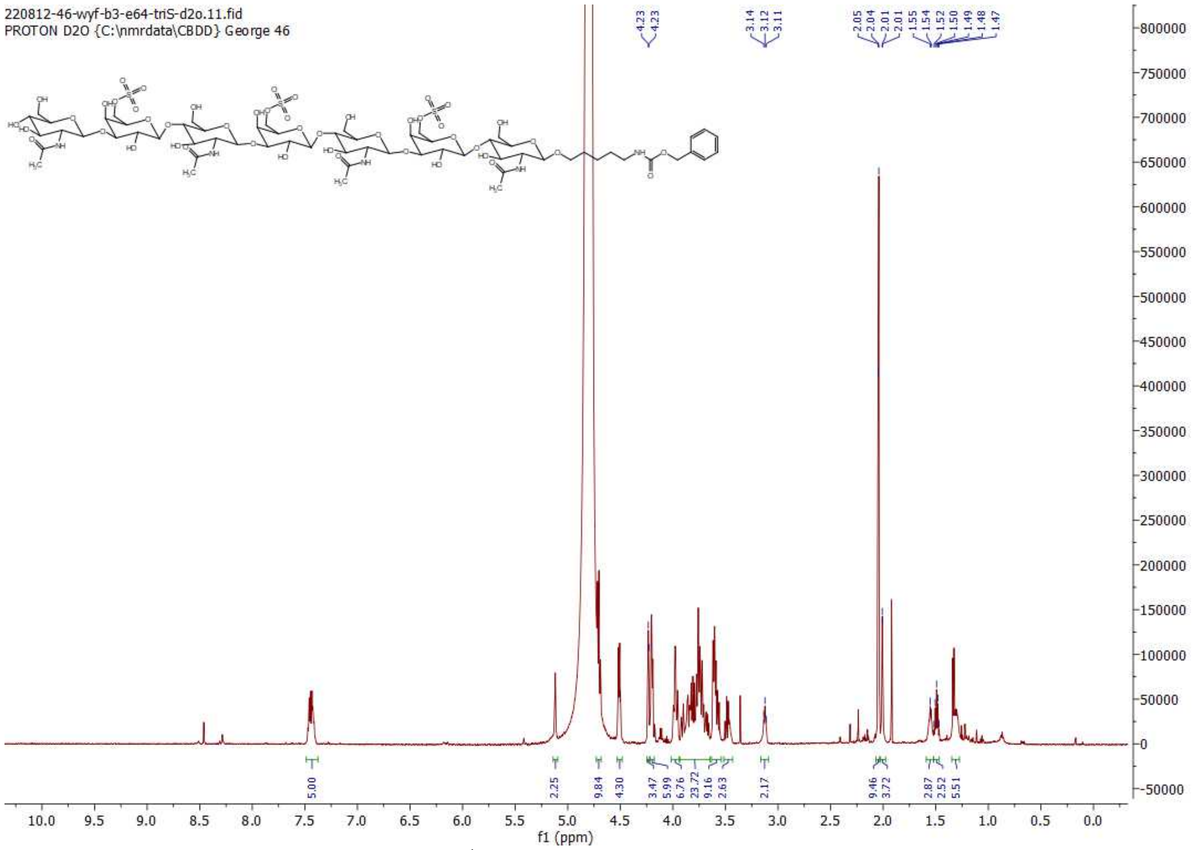

$^{1}$H NMR of 18; 600MHz; $D_2O$

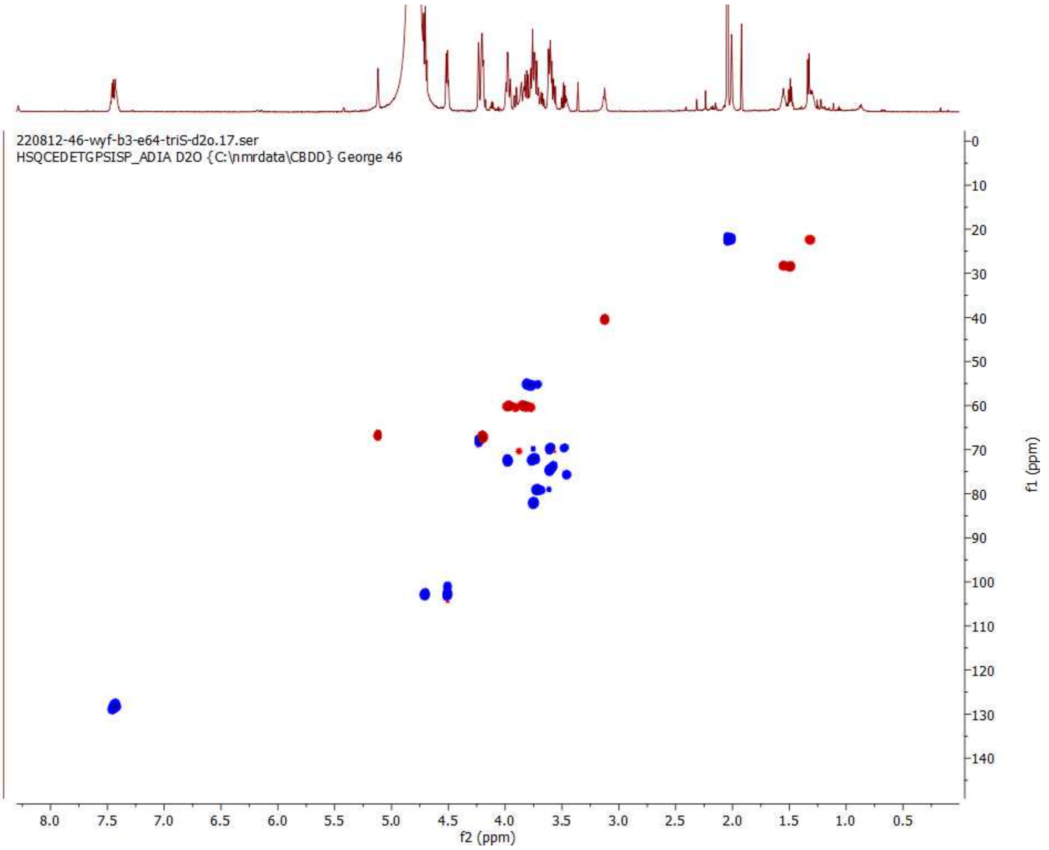

HSQC of 18; 600 MHz/150 MHz, $D_2O$

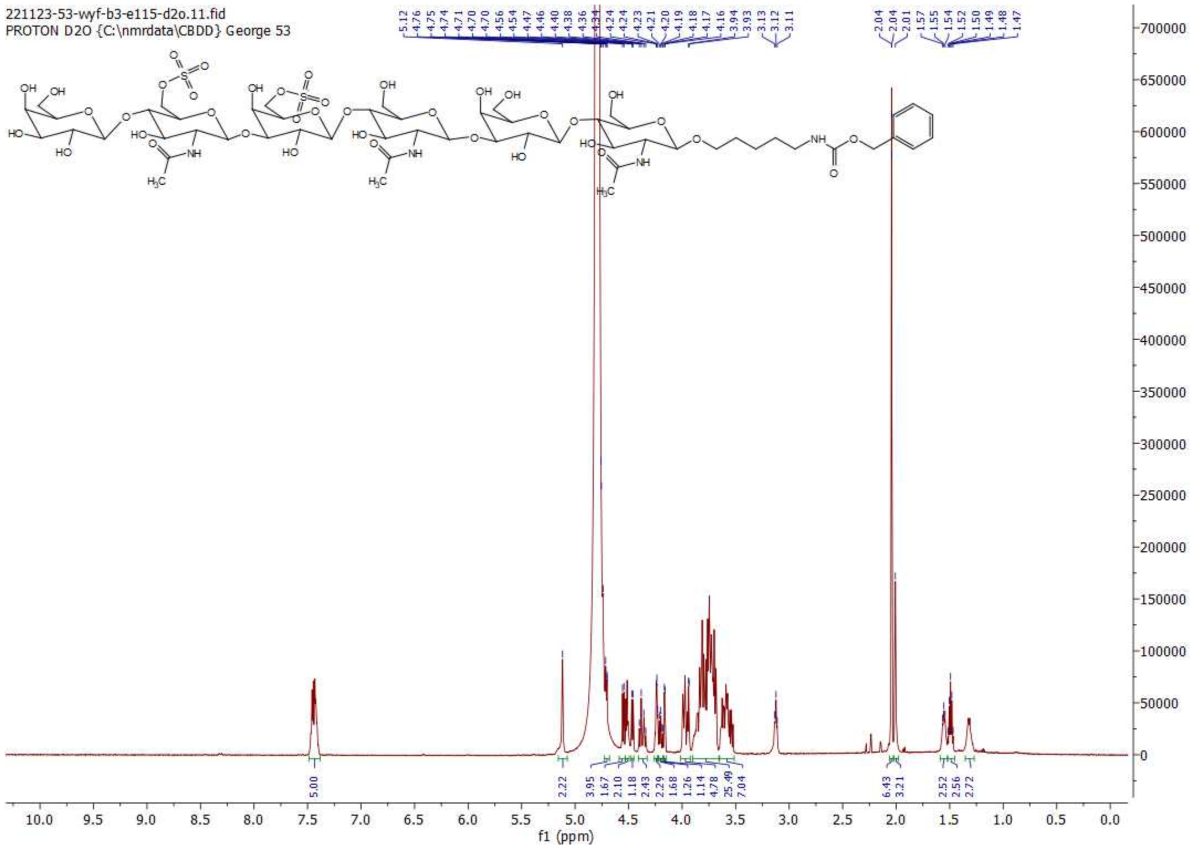

**$^{1}H$ NMR of 19; 600MHz; $D_2O$**

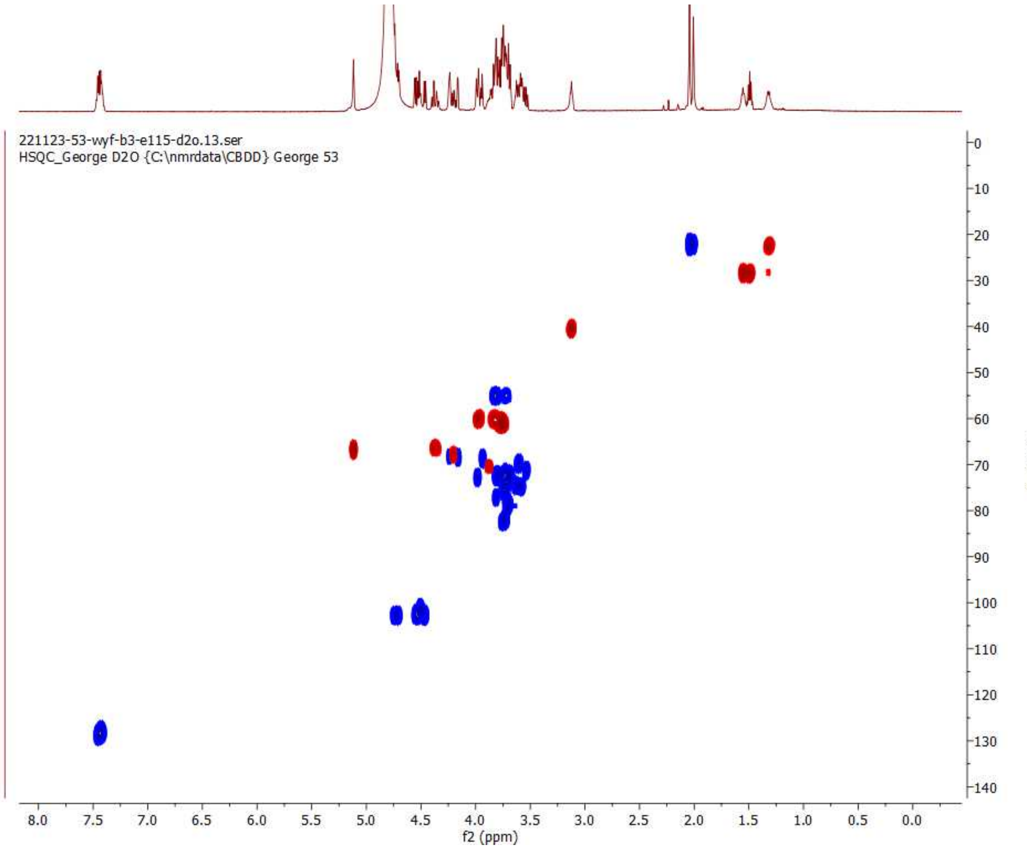

**HSQC of 19; 600 MHz/150 MHz, $D_2O$**

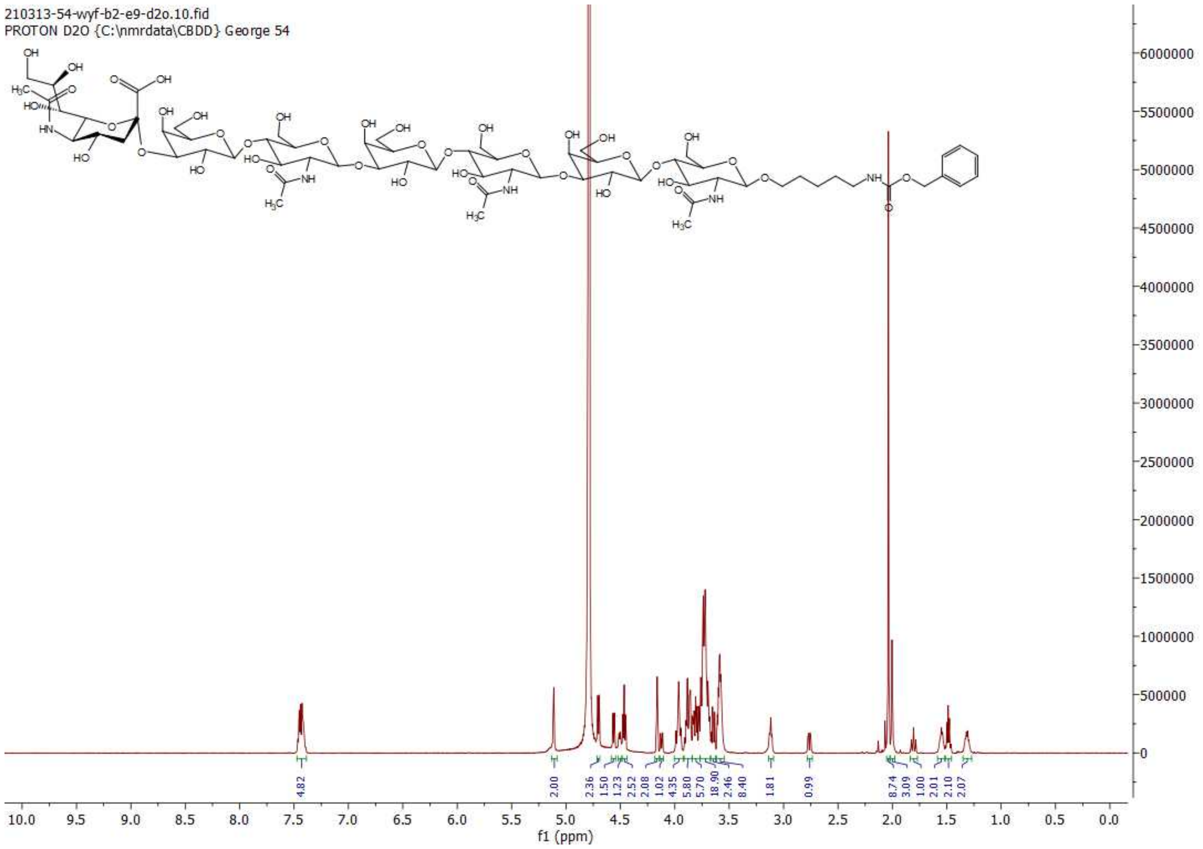

**[1]H NMR of 20; 600MHz; $D_2O$**

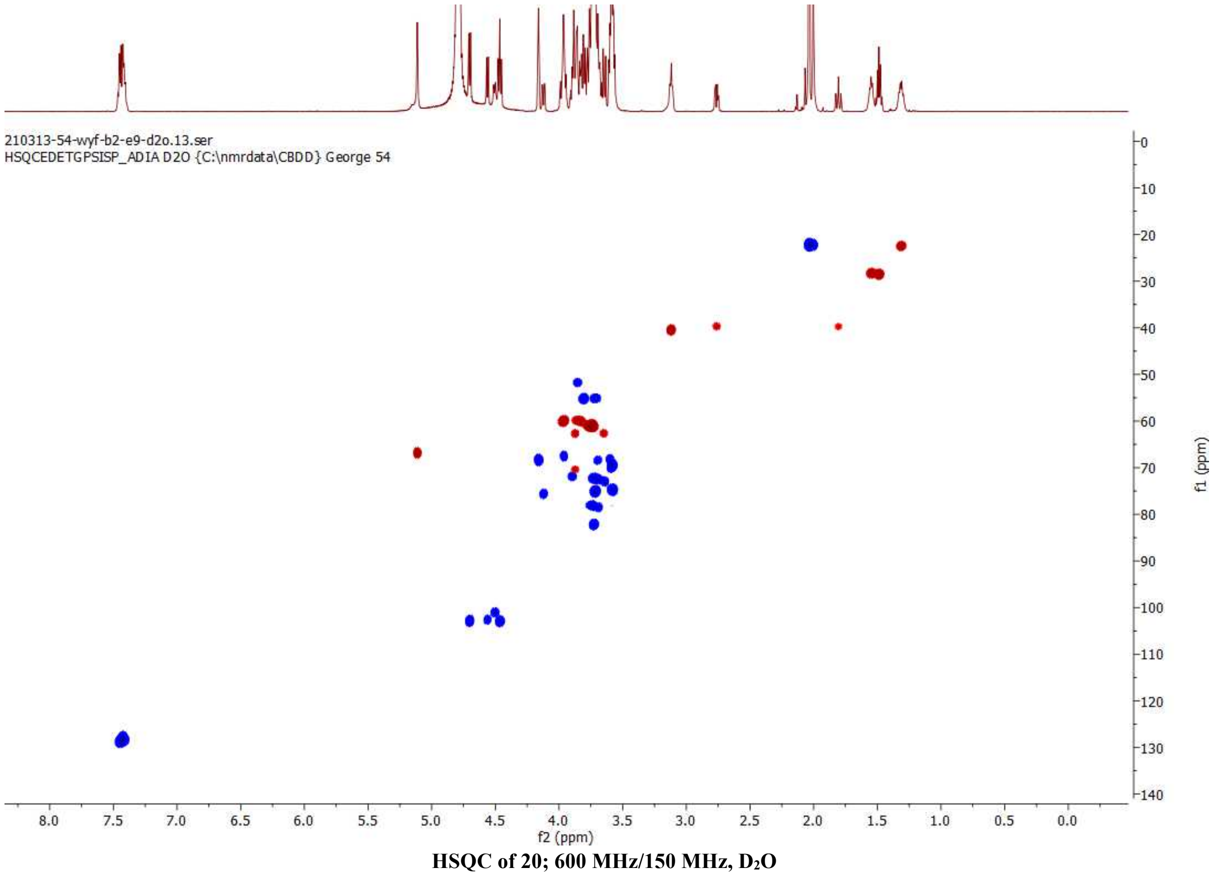

**HSQC of 20; 600 MHz/150 MHz, $D_2O$**

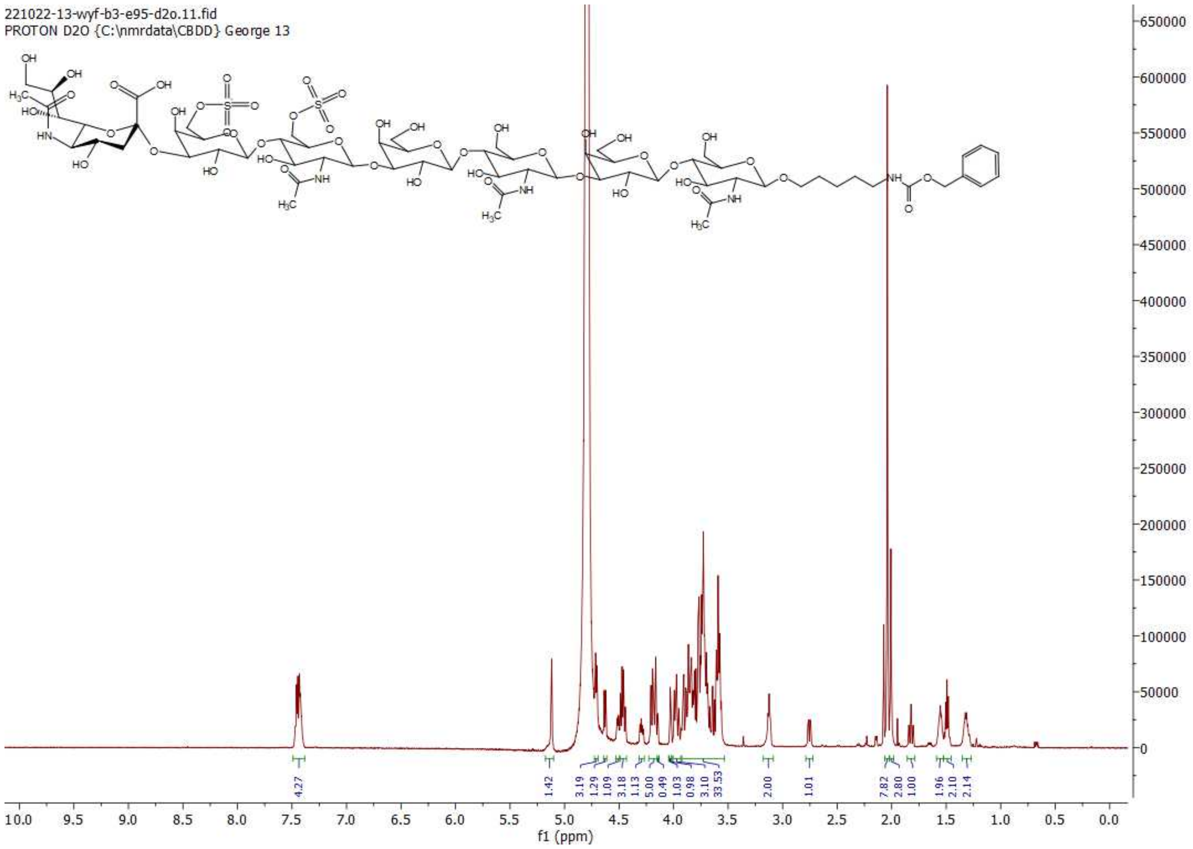

$^{1}$H NMR of 21; 600MHz; $D_2O$

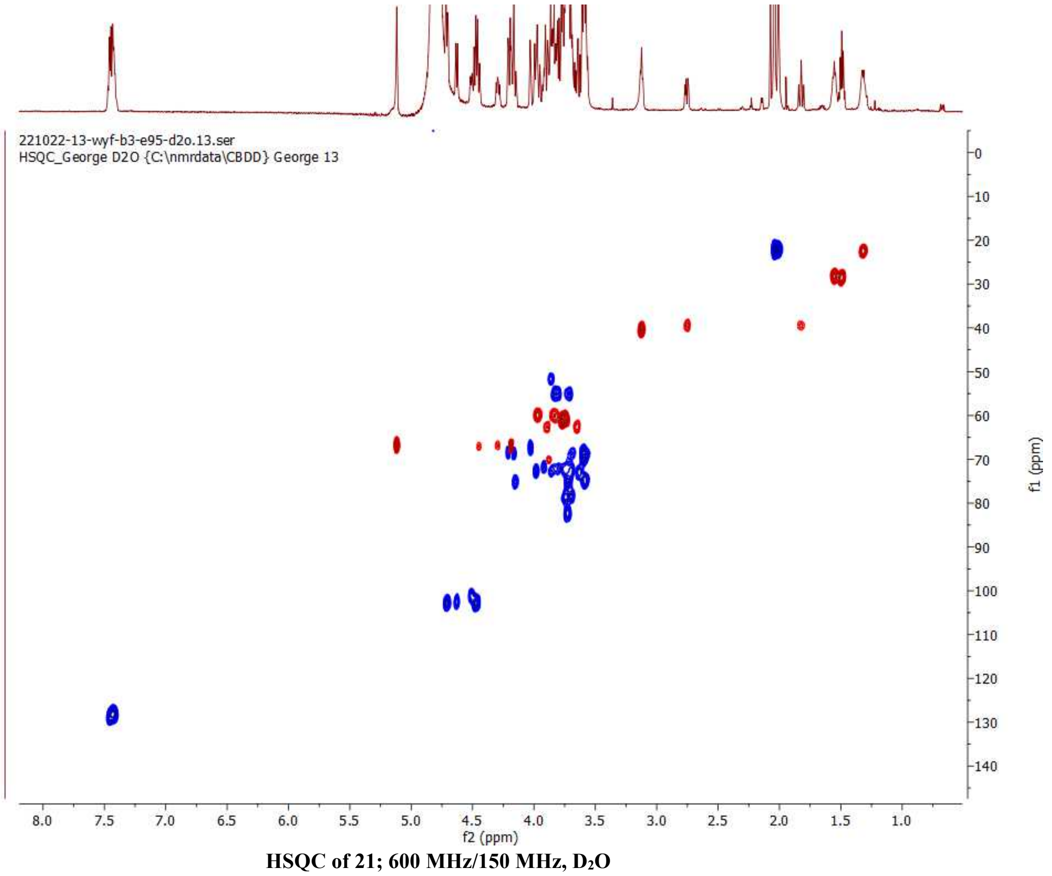

HSQC of 21; 600 MHz/150 MHz, $D_2O$

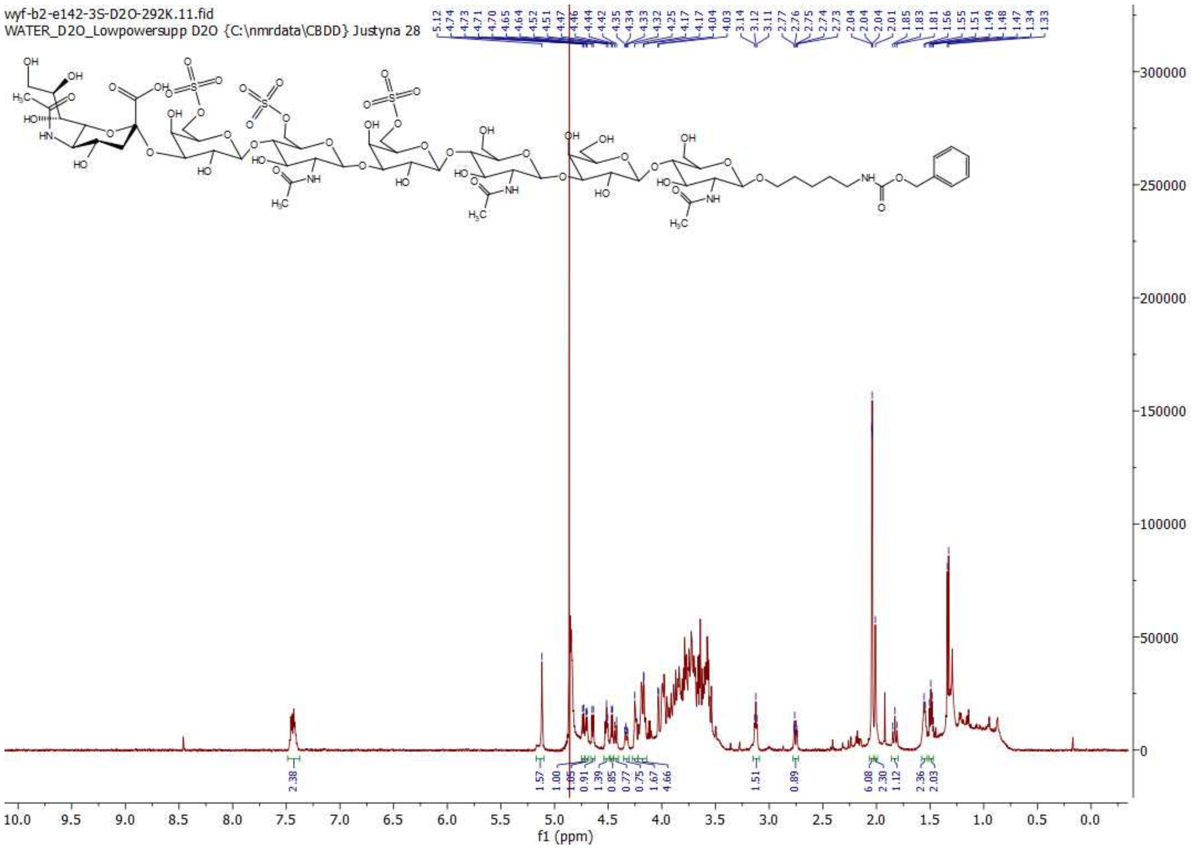

$^{1}$H NMR of 22; 600MHz; $D_2O$

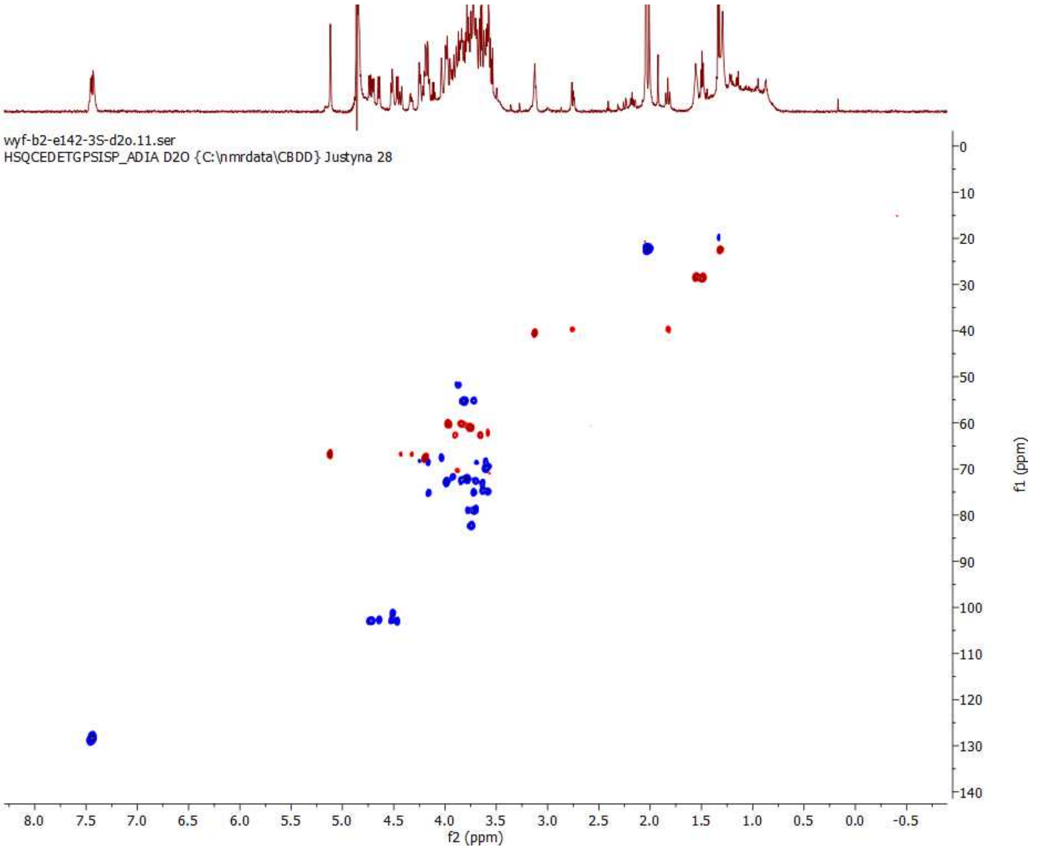

HSQC of 22; 600 MHz/150 MHz, $D_2O$

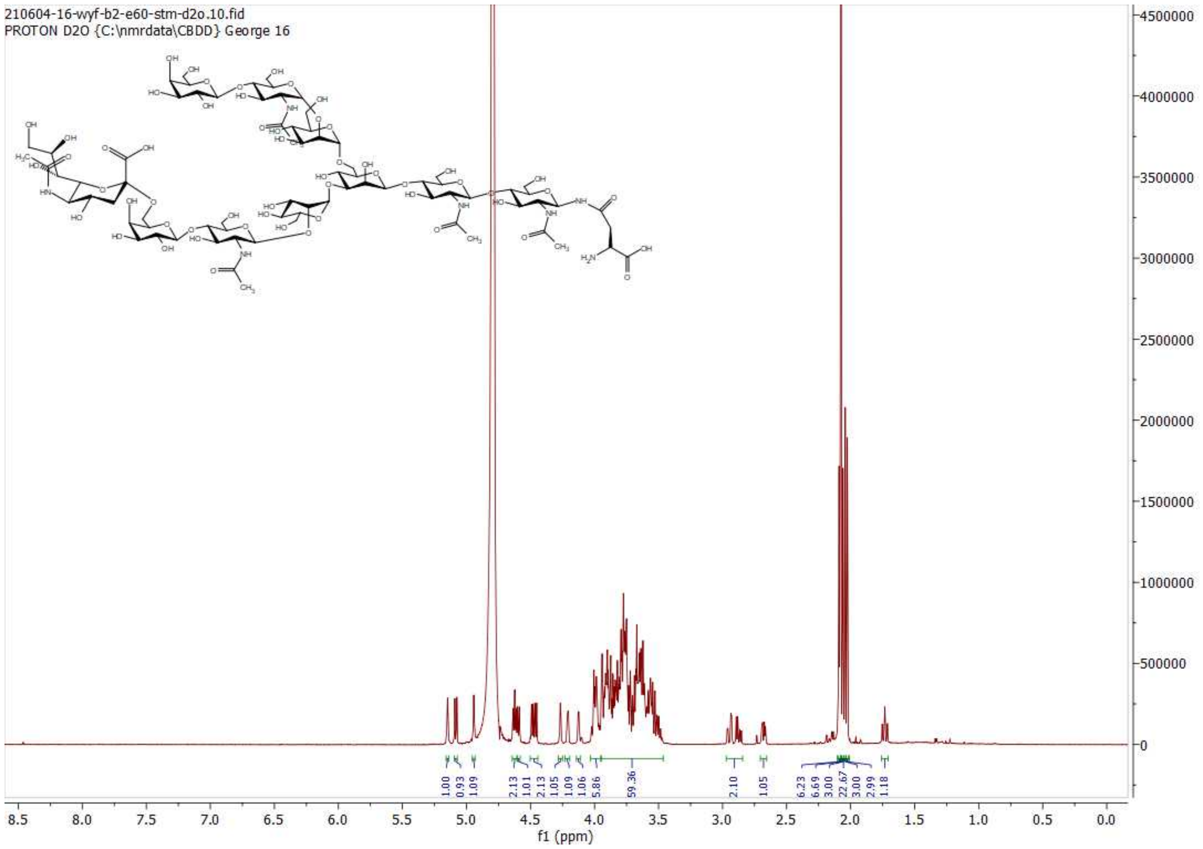

$^{1}$H NMR of 24; 600MHz; $D_2O$

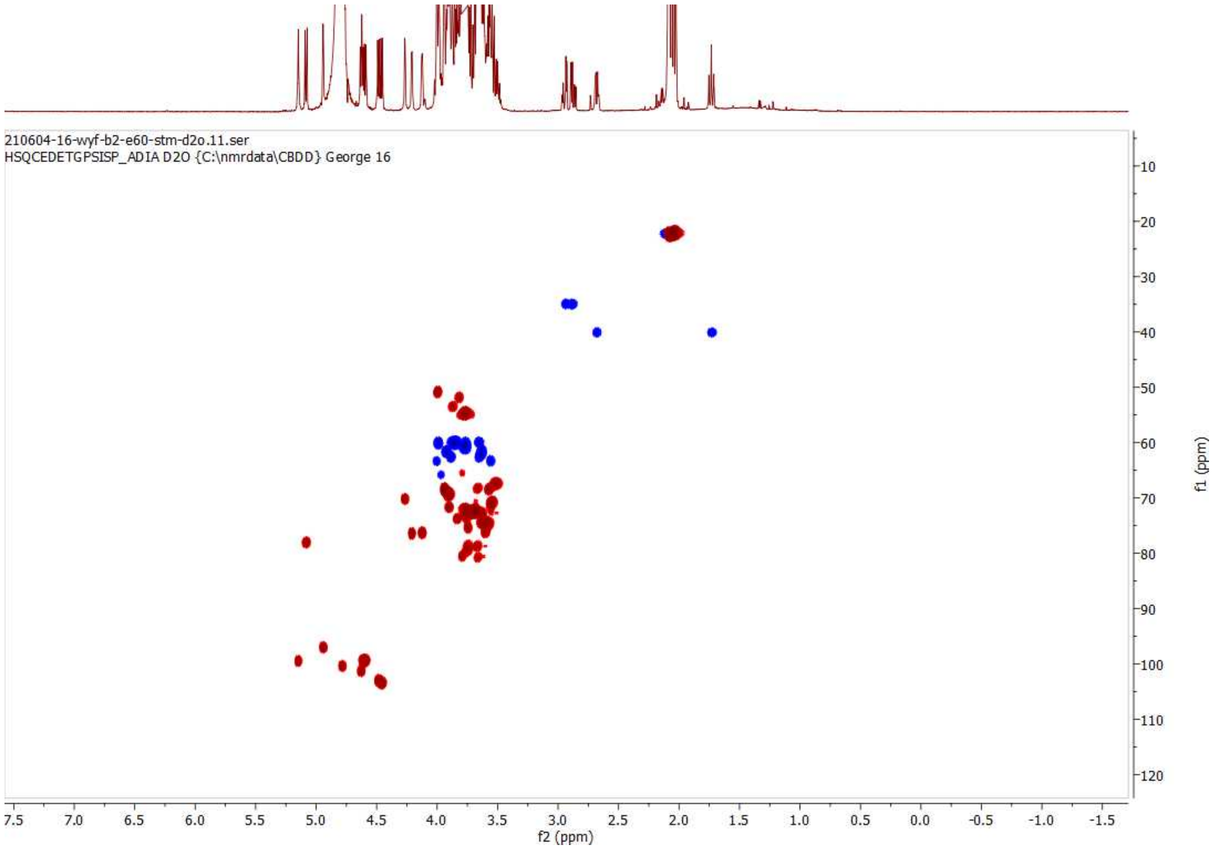

HSQC of 24; 600 MHz/150 MHz, $D_2O$

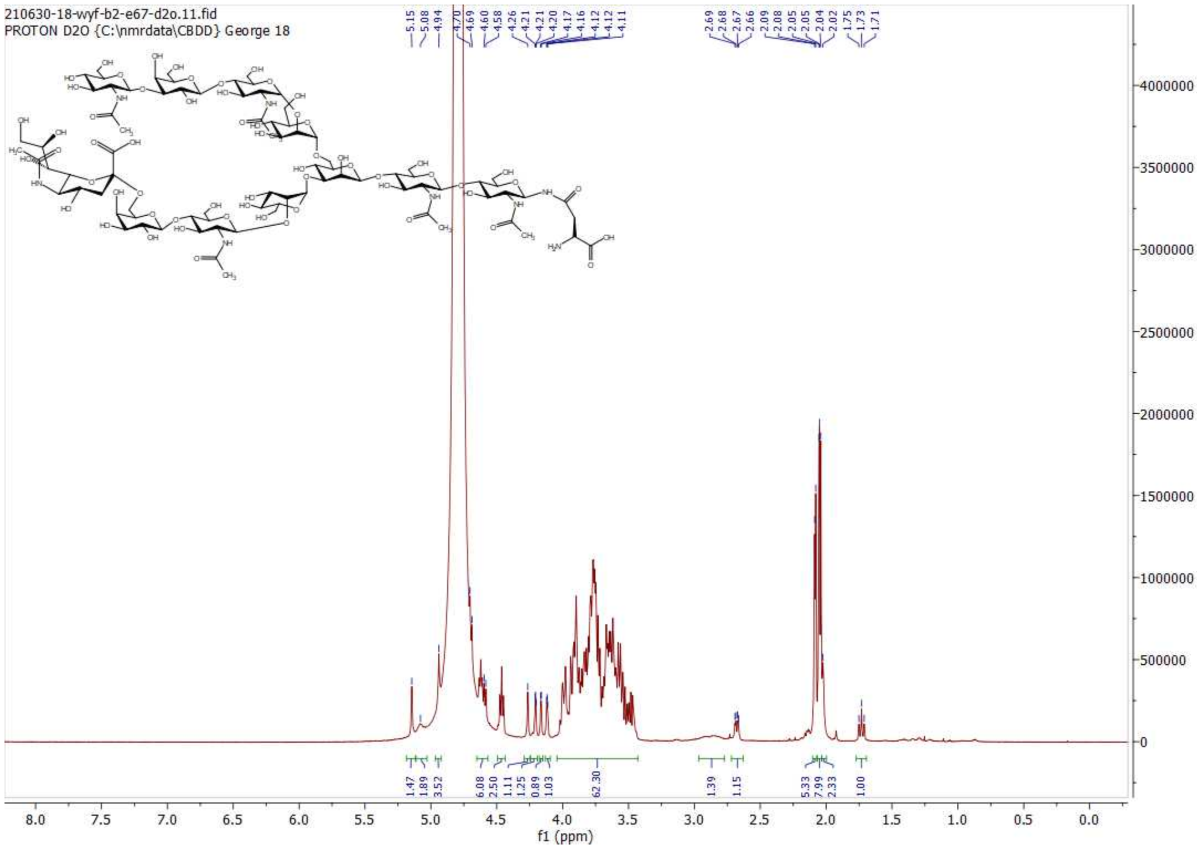

$^1$H NMR of 25; 600MHz; $D_2O$

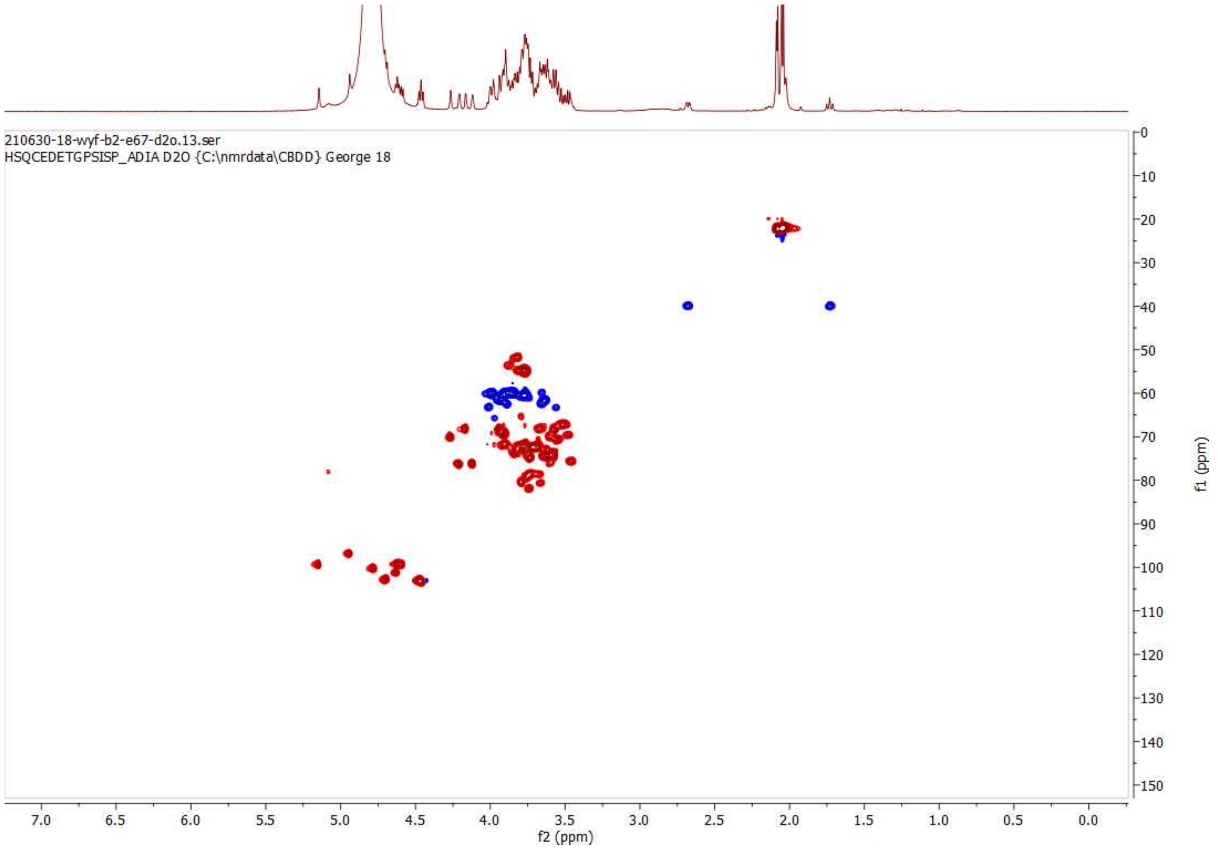

HSQC of 25; 600 MHz/150 MHz, $D_2O$

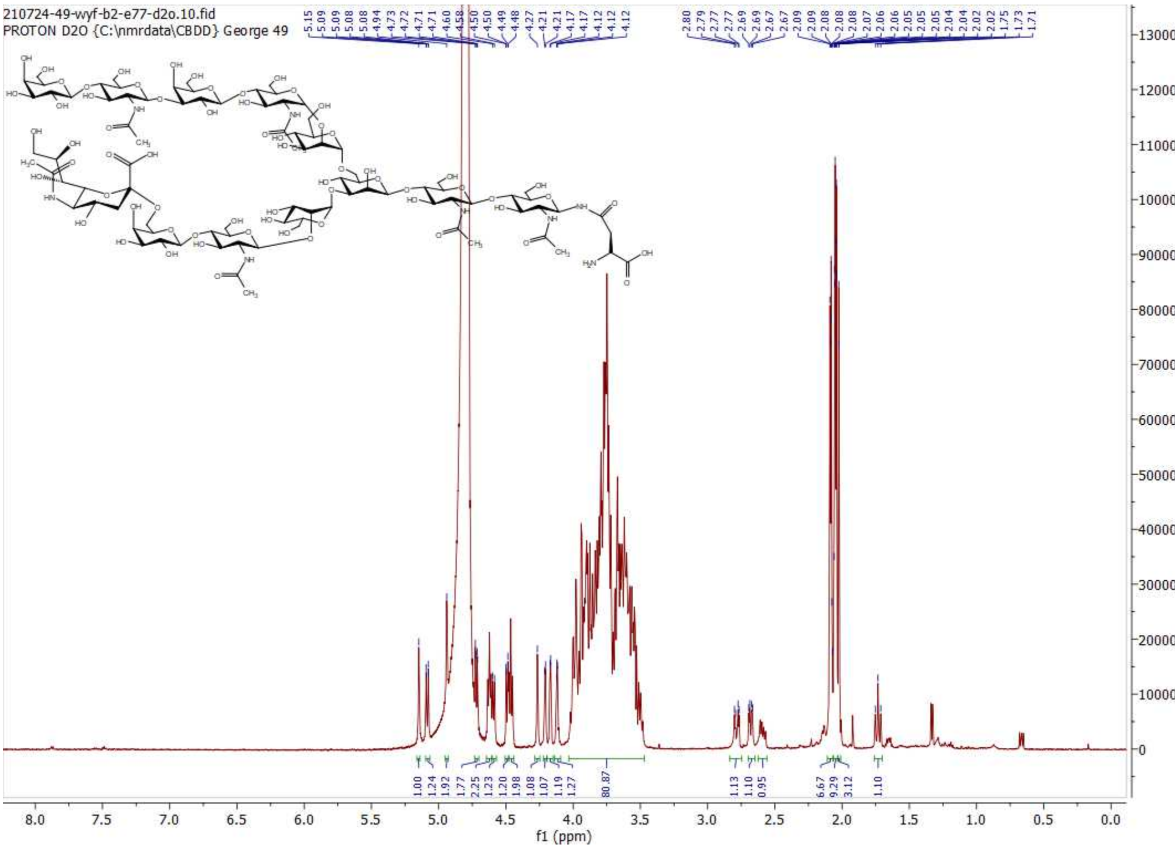

$^1$H NMR of 26; 600MHz; $D_2O$

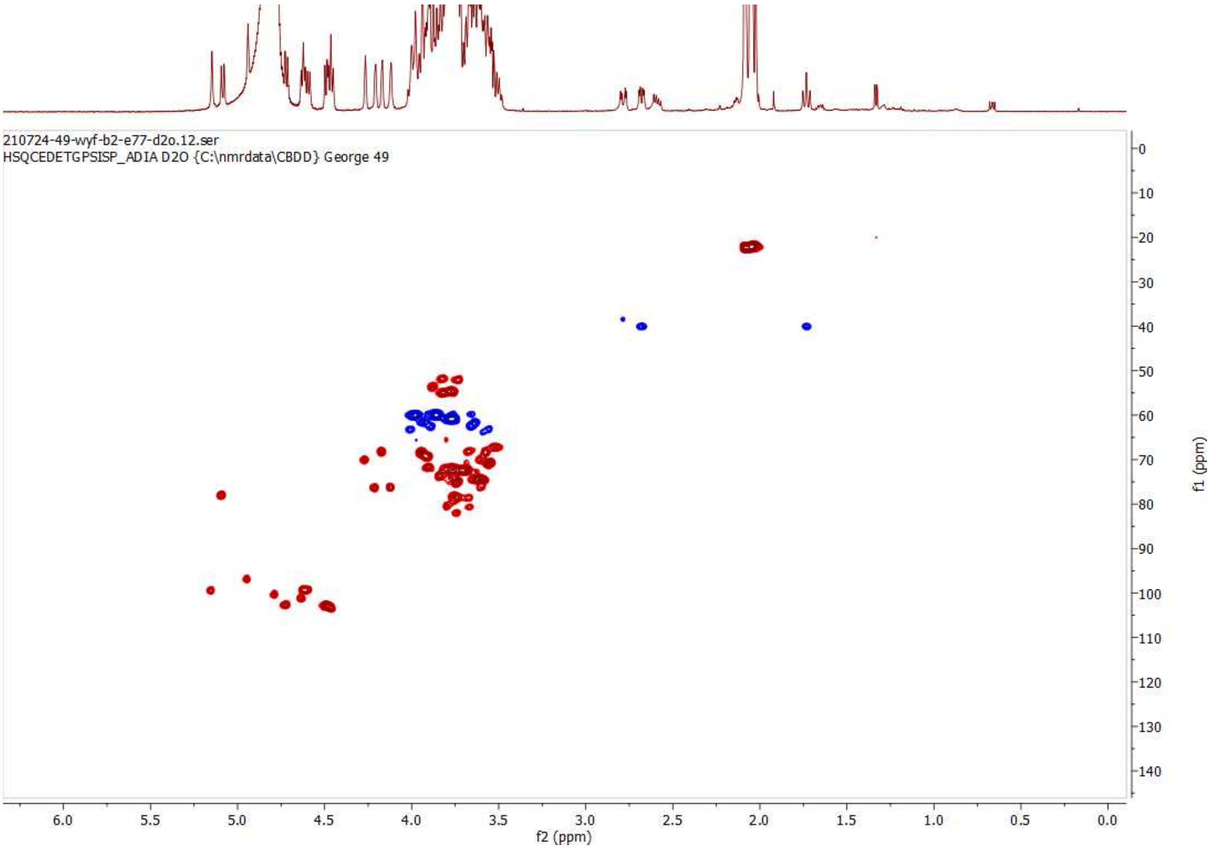

HSQC of 26; 600 MHz/150 MHz, $D_2O$

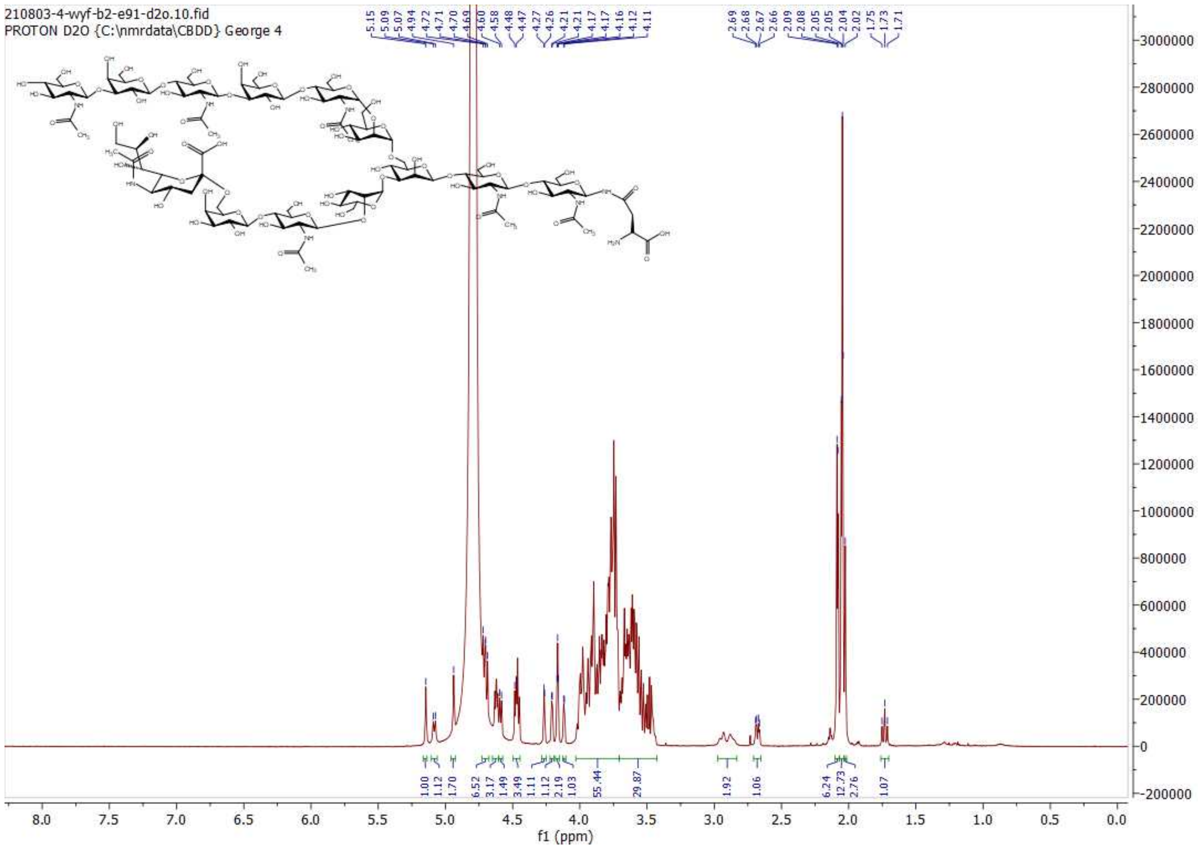

**[1]H NMR of 27; 600MHz; $D_2O$**

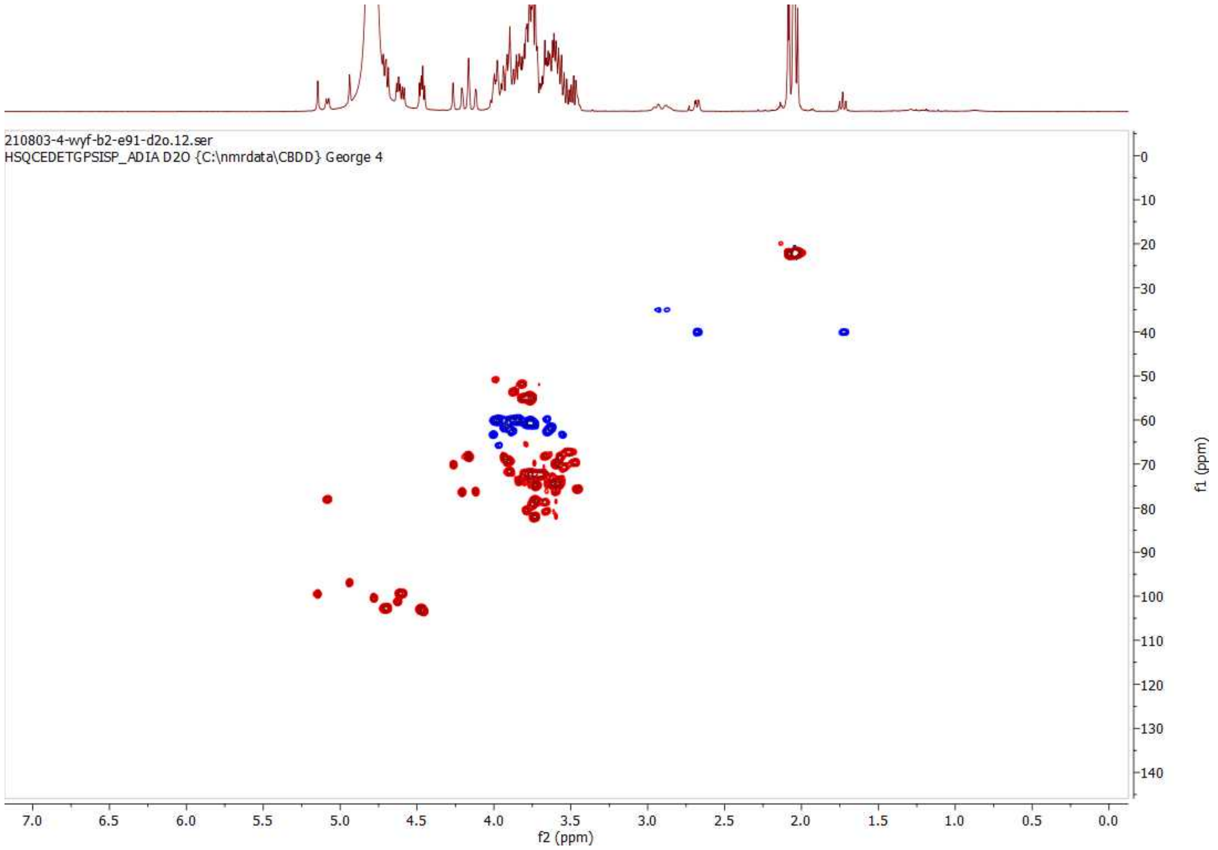

**HSQC of 27; 600 MHz/150 MHz, $D_2O$**

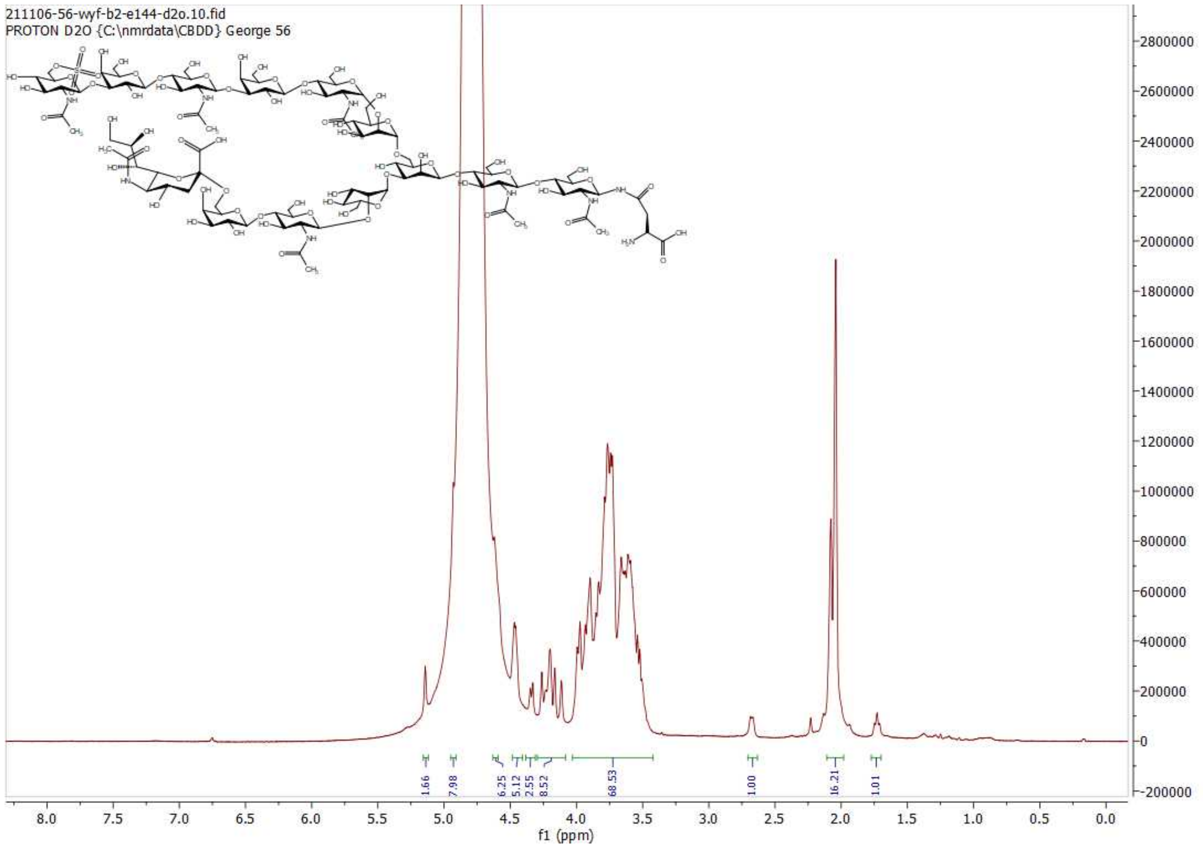

**[1]H NMR of 28; 600MHz; D$_2$O**

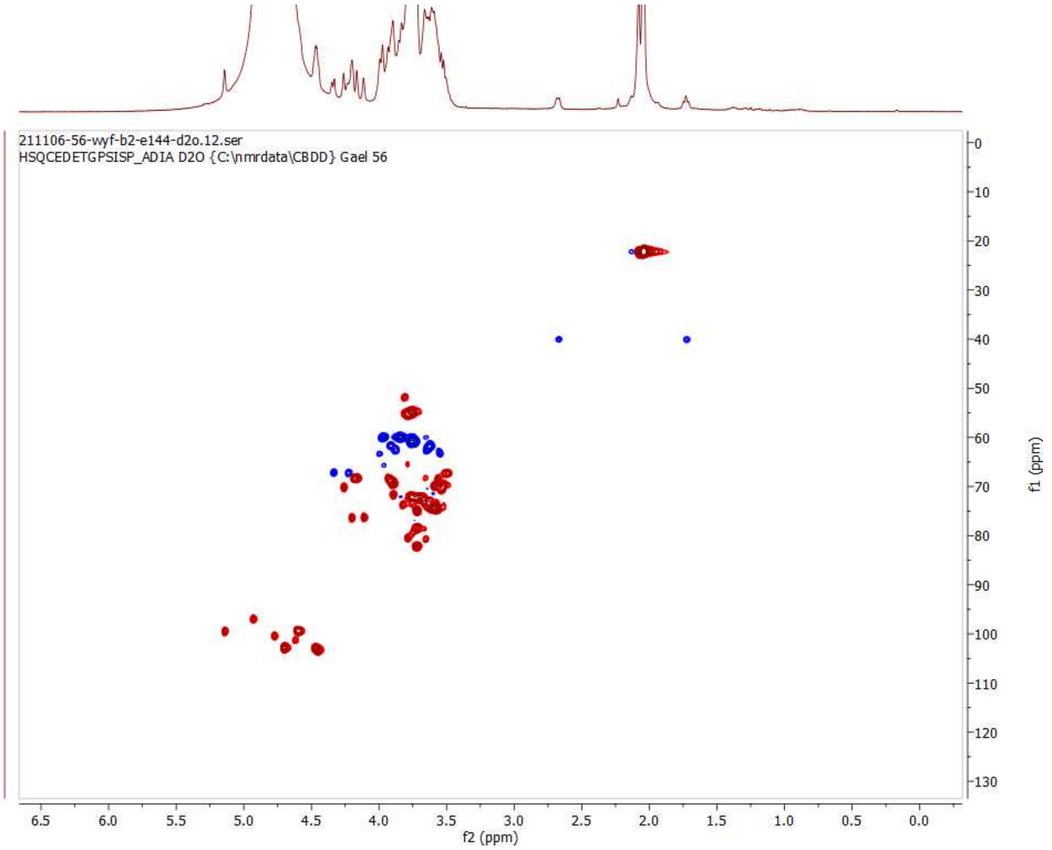

**HSQC of 28; 600 MHz/150 MHz, D$_2$O**

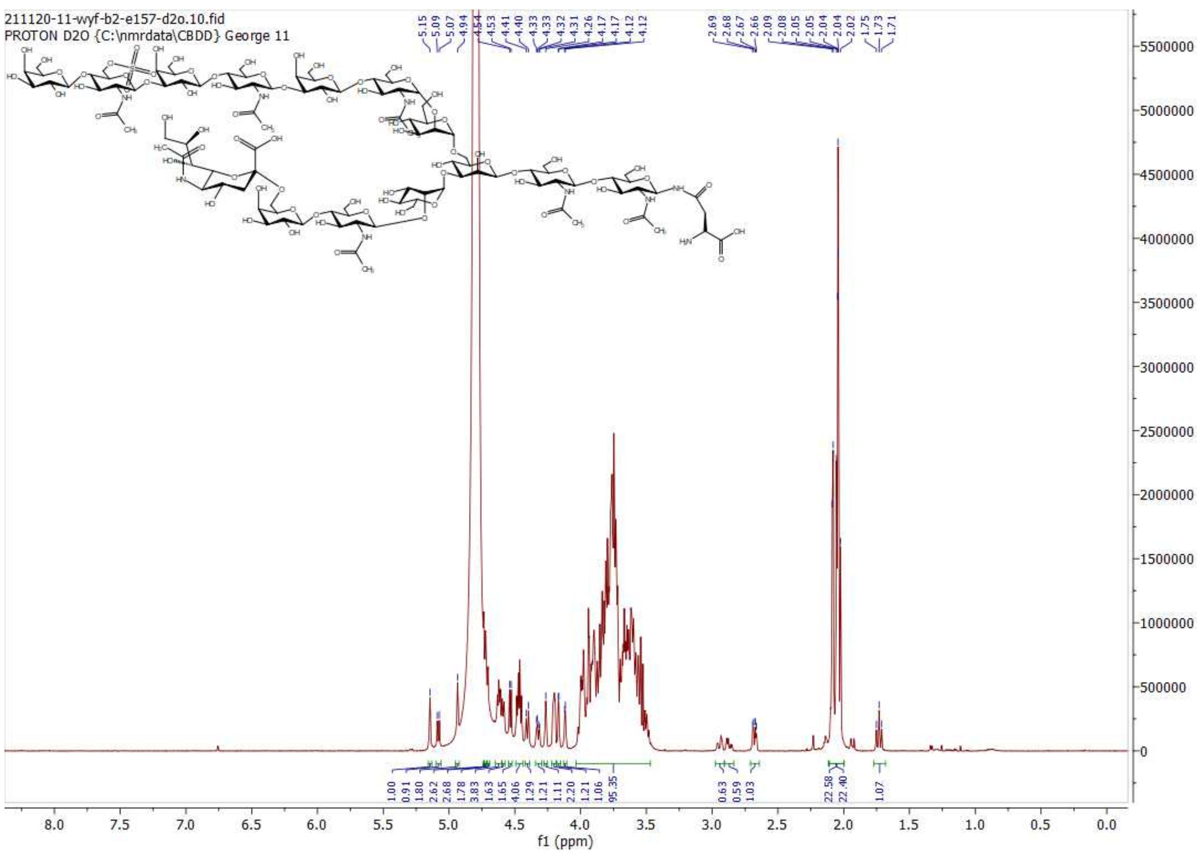

$^{1}$H NMR of 29; 600MHz; $D_2O$

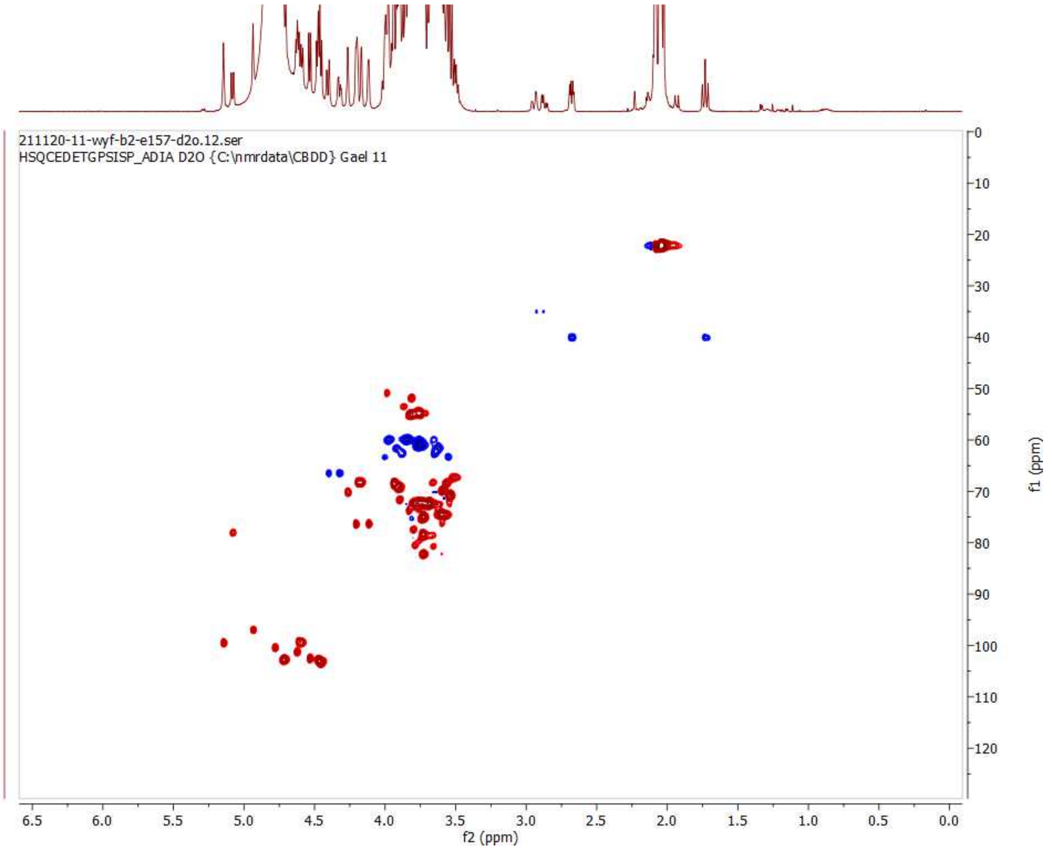

HSQC of 29; 600 MHz/150 MHz, $D_2O$

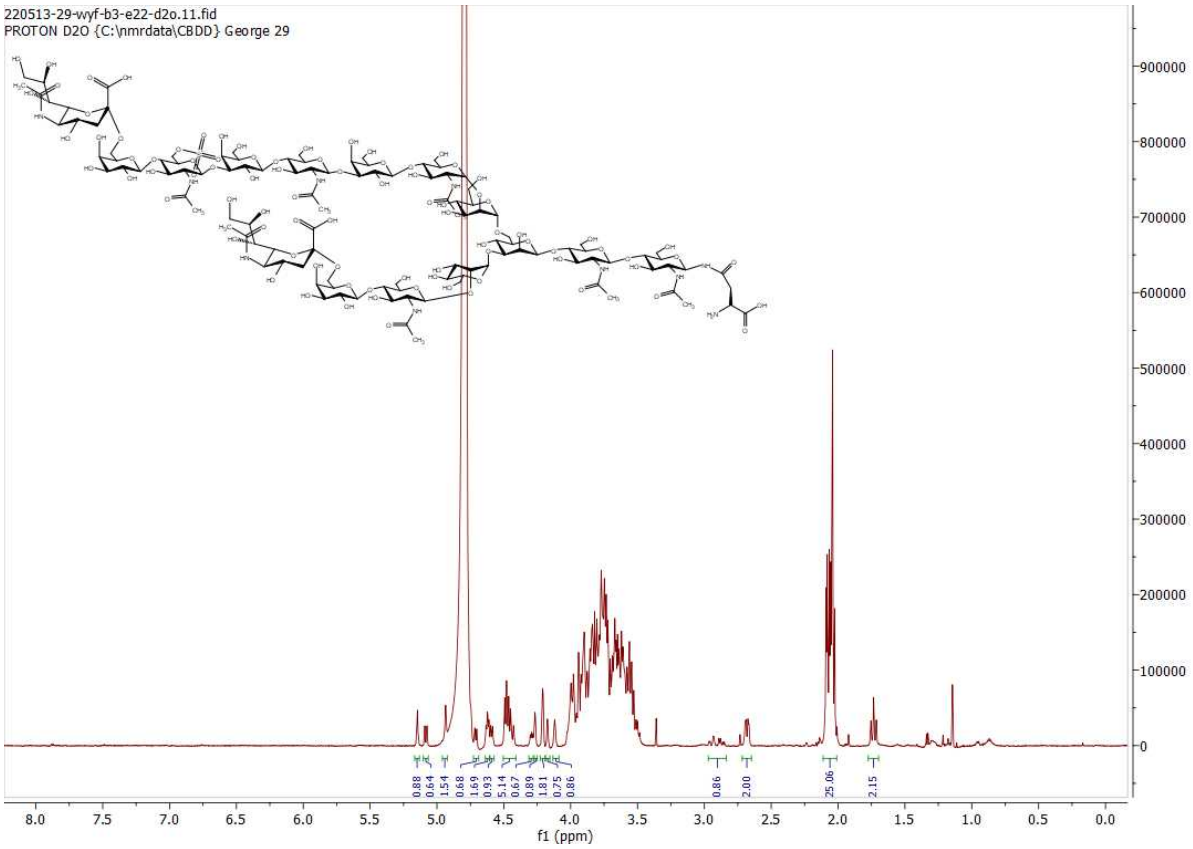

**[1]H NMR of 30; 600MHz; $D_2O$**

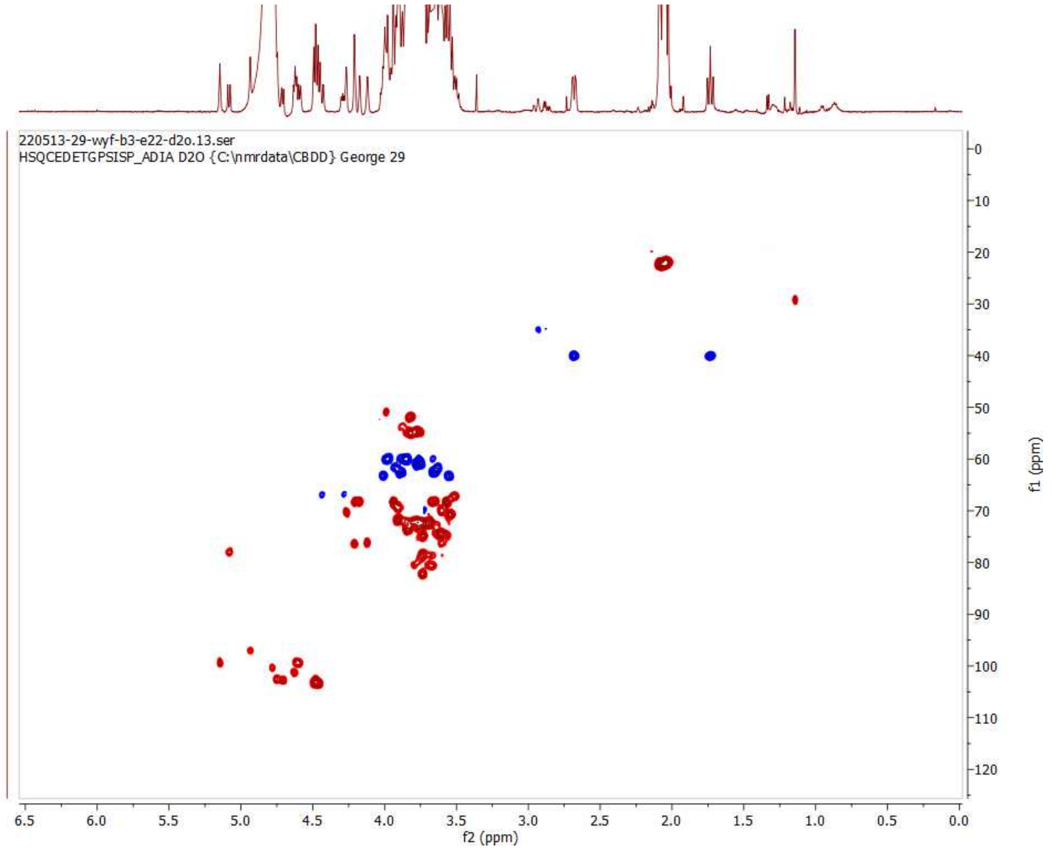

**HSQC of 30; 600 MHz/150 MHz, $D_2O$**

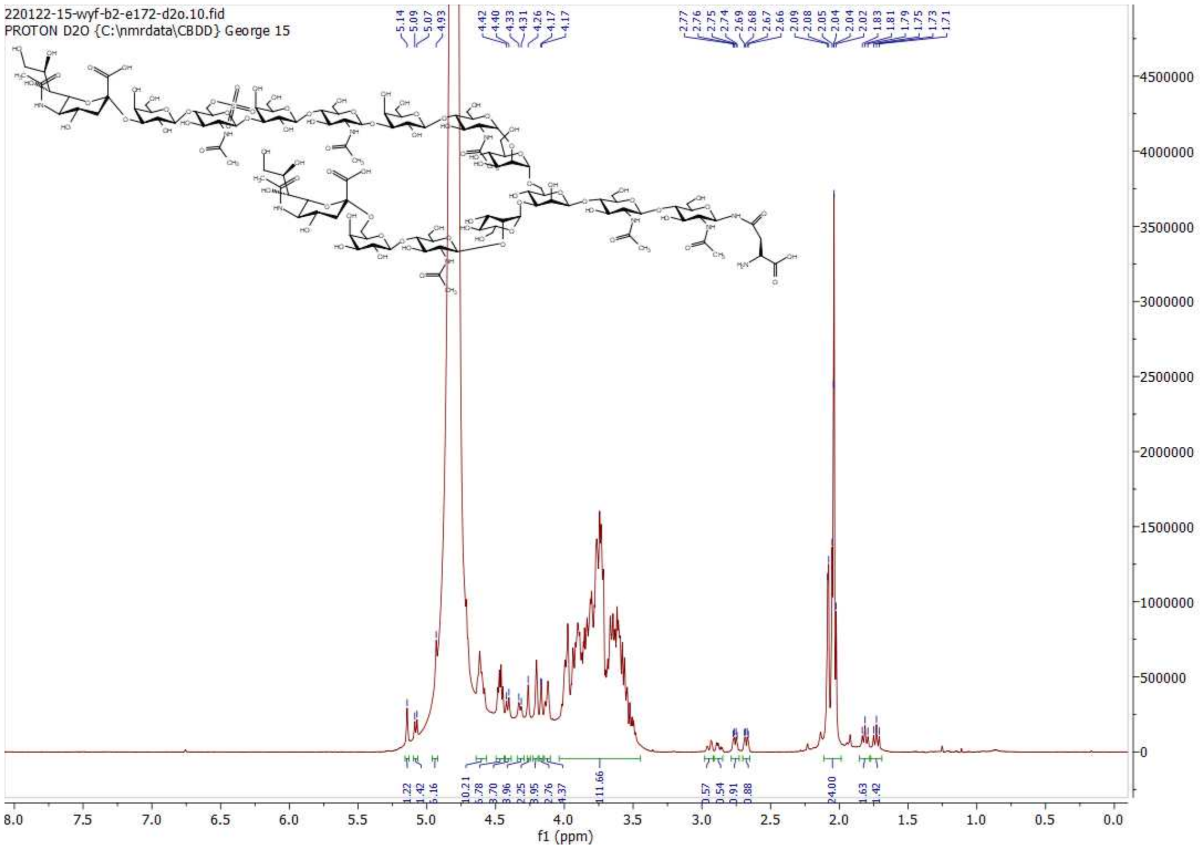

**[1]H NMR of 31; 600MHz; $D_2O$**

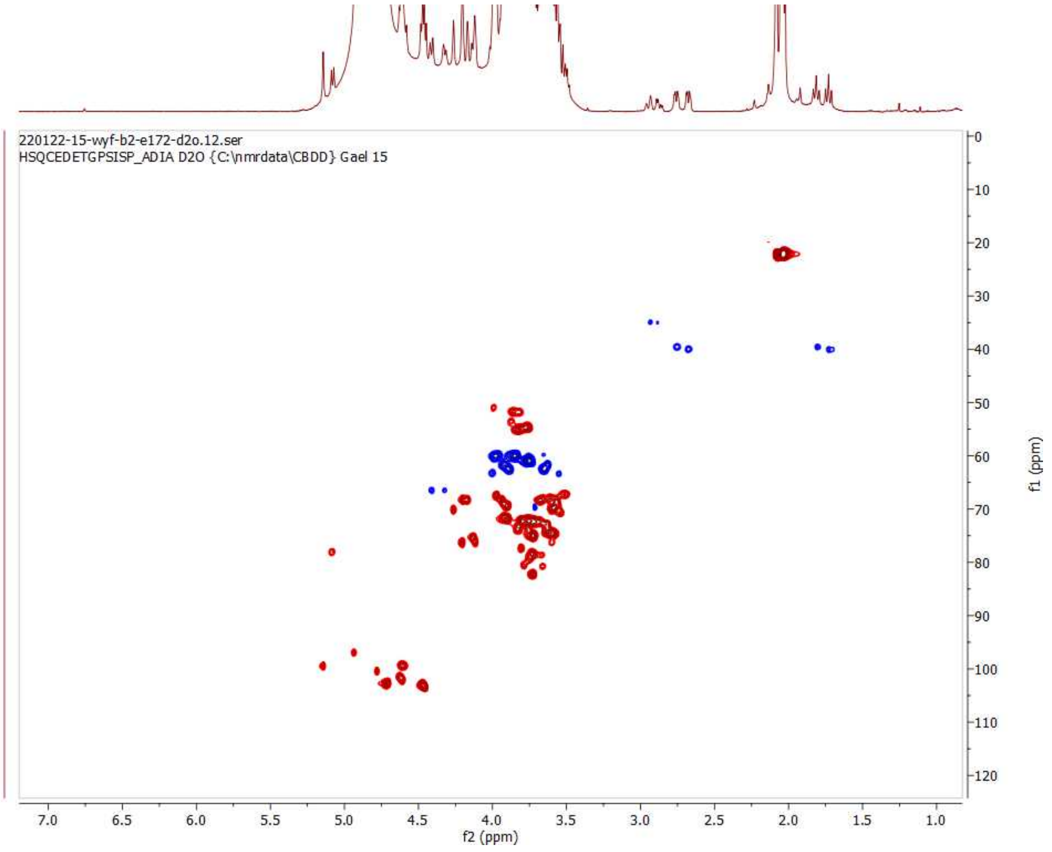

**HSQC of 31; 600 MHz/150 MHz, $D_2O$**

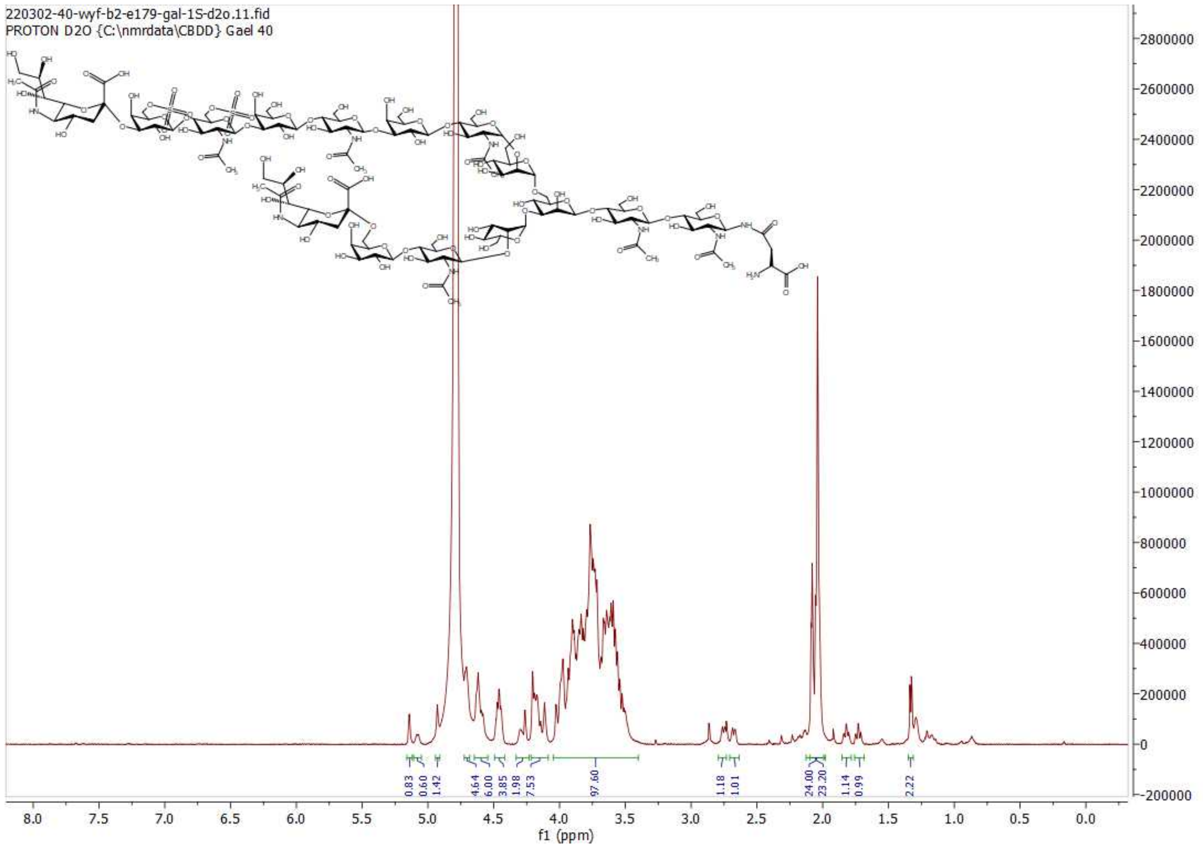

$^1$H NMR of 32; 600MHz; $D_2O$

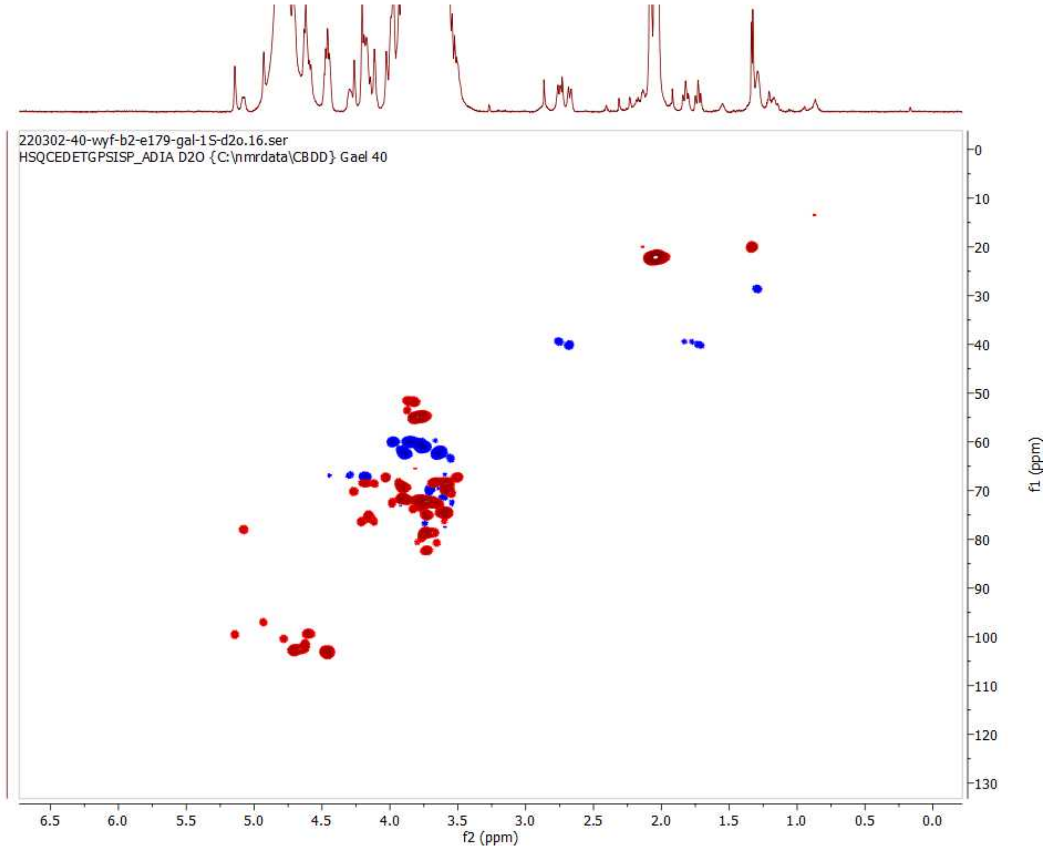

HSQC of 32; 600 MHz/150 MHz, $D_2O$

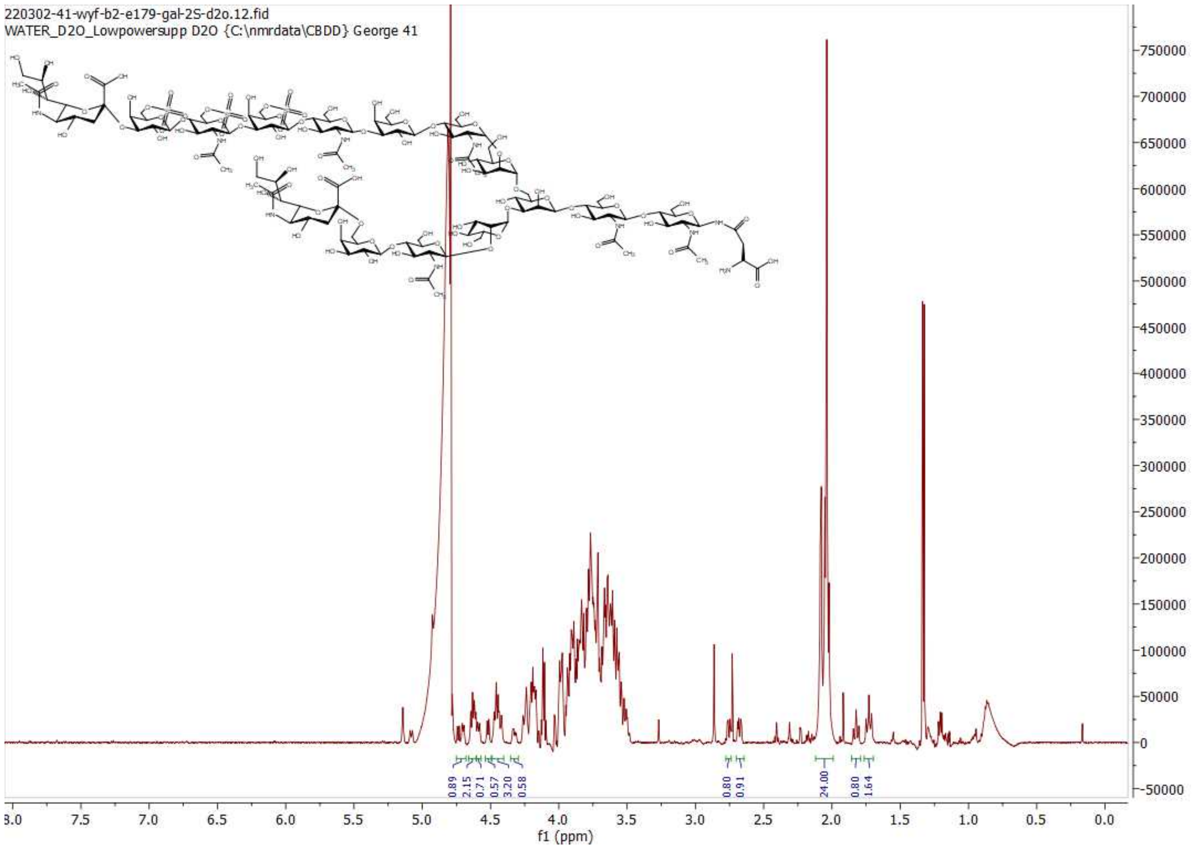

**[1]H NMR of 33; 600MHz; $D_2O$**

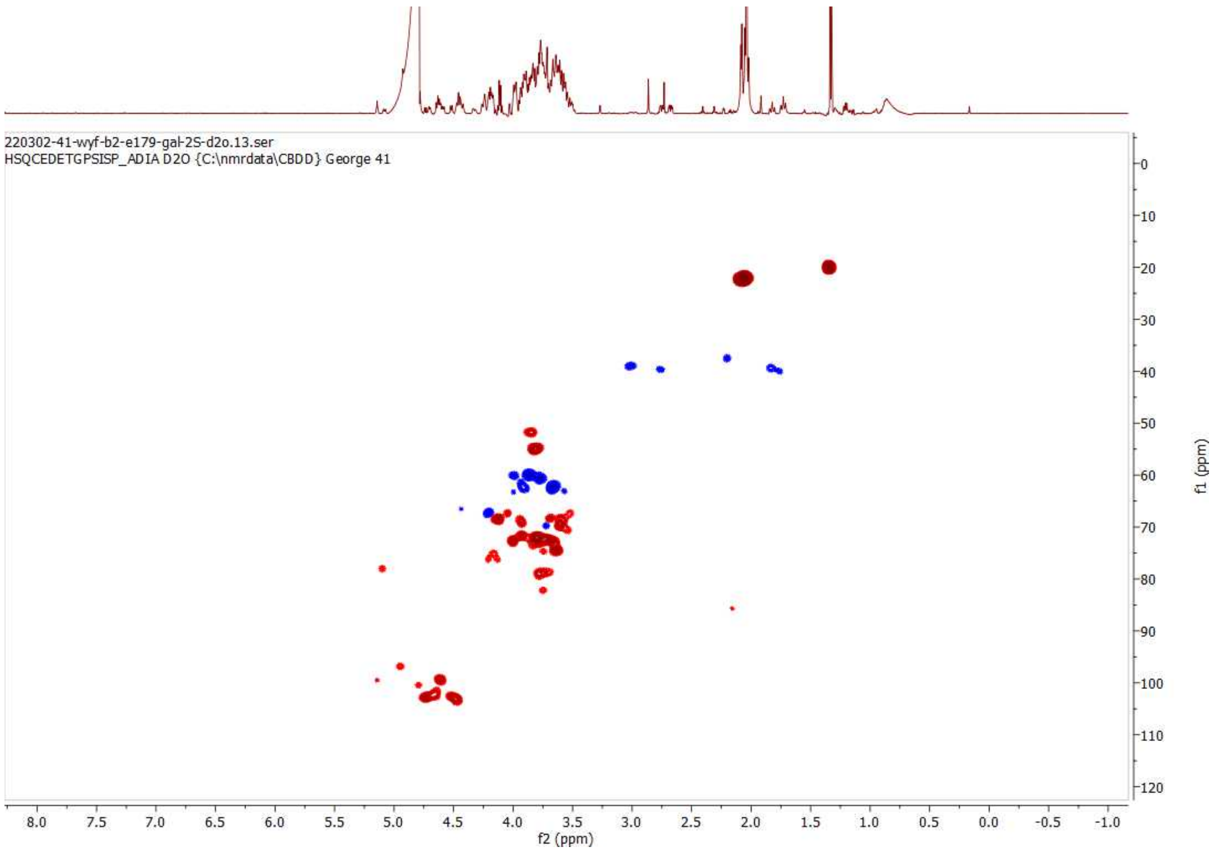

**HSQC of 33; 600 MHz/150 MHz, $D_2O$**

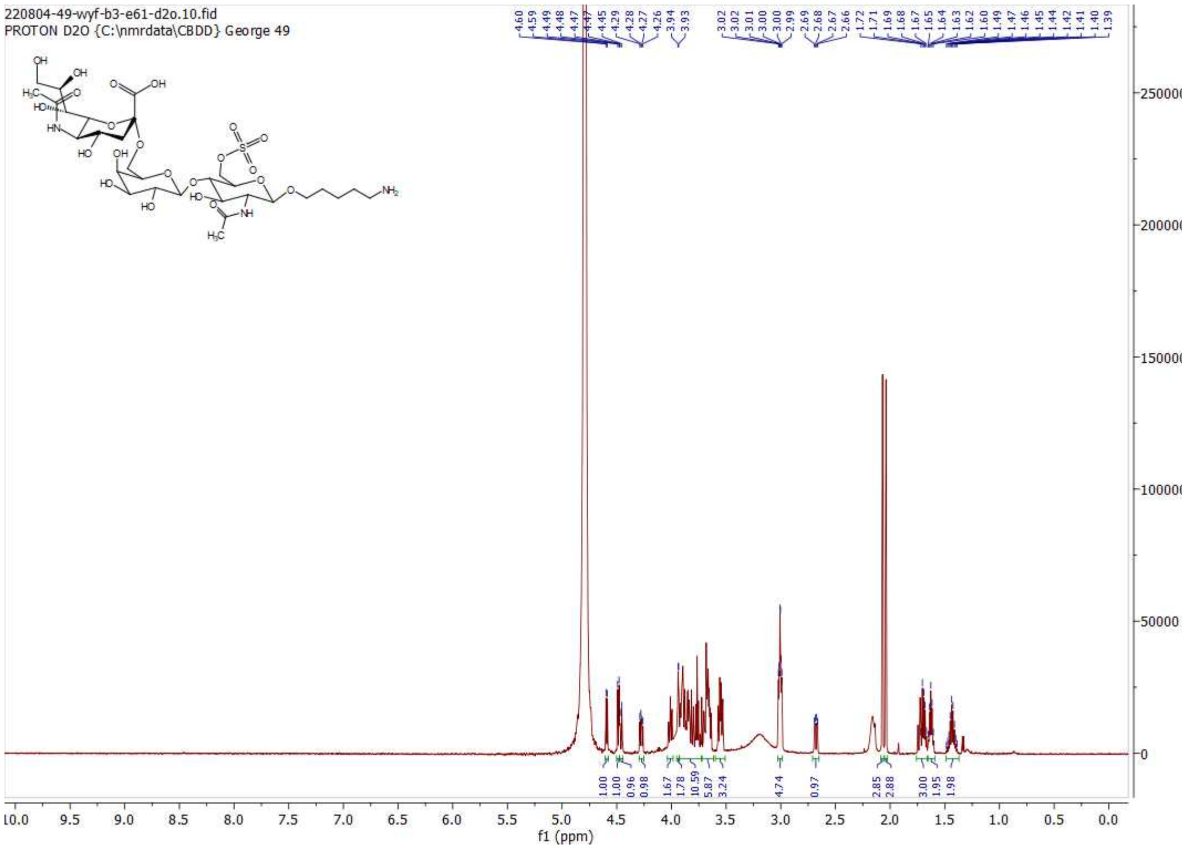

**$^{1}$H NMR of 34; 600MHz; $D_2O$**

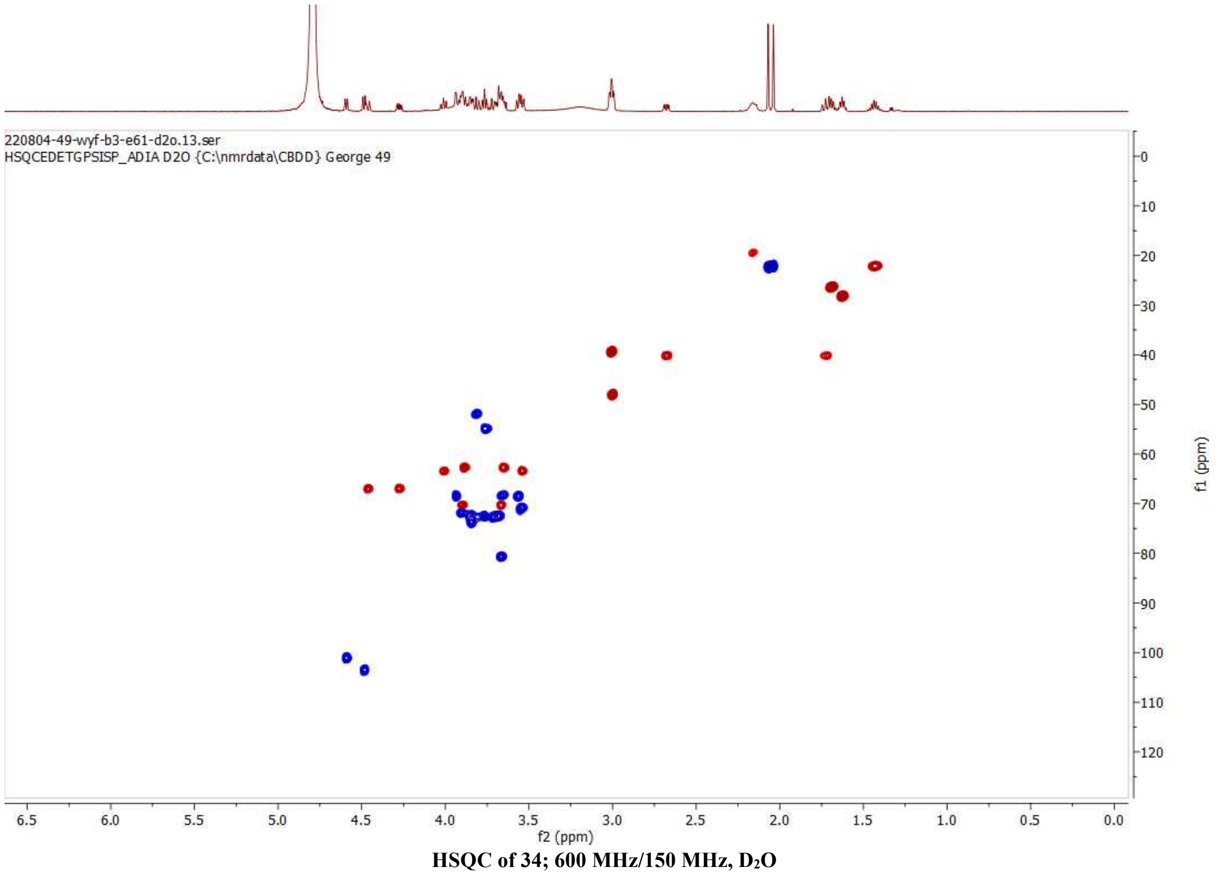

**HSQC of 34; 600 MHz/150 MHz, $D_2O$**

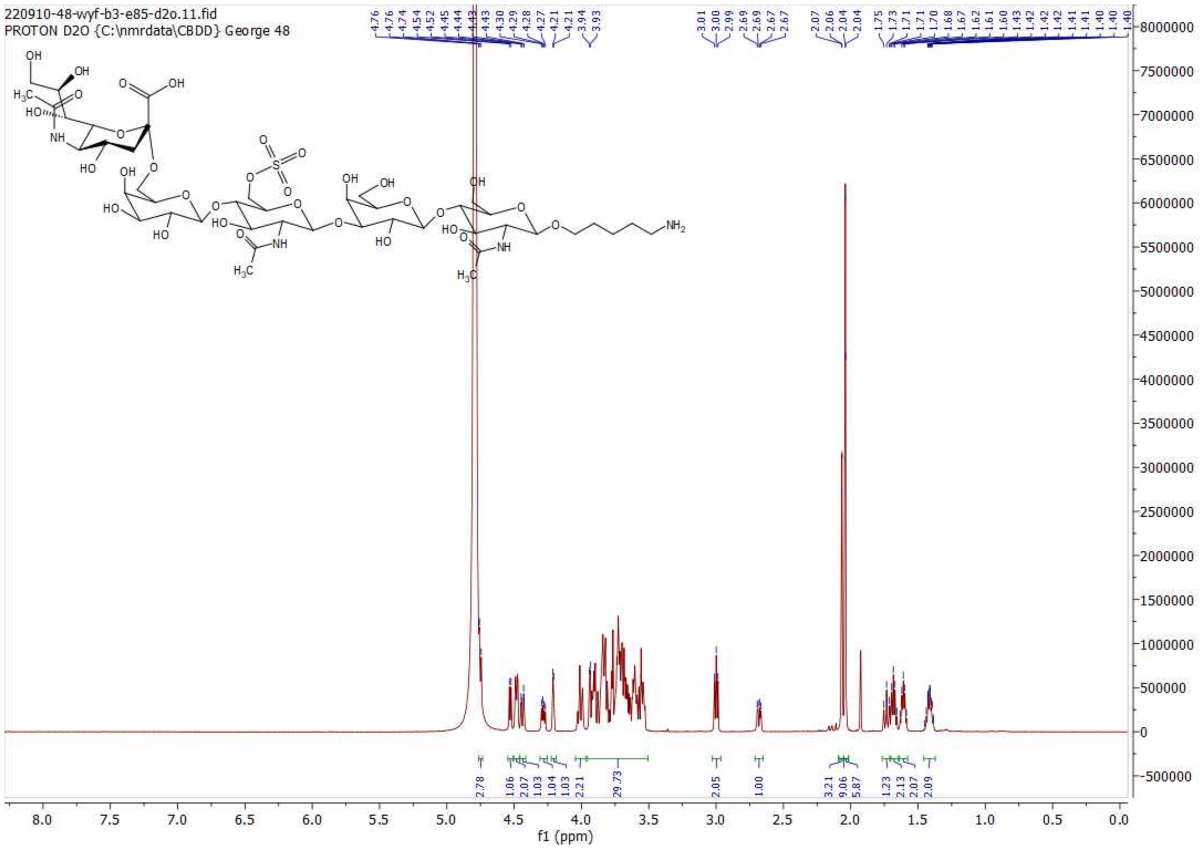

$^{1}H$ NMR of 35; 600MHz; $D_2O$

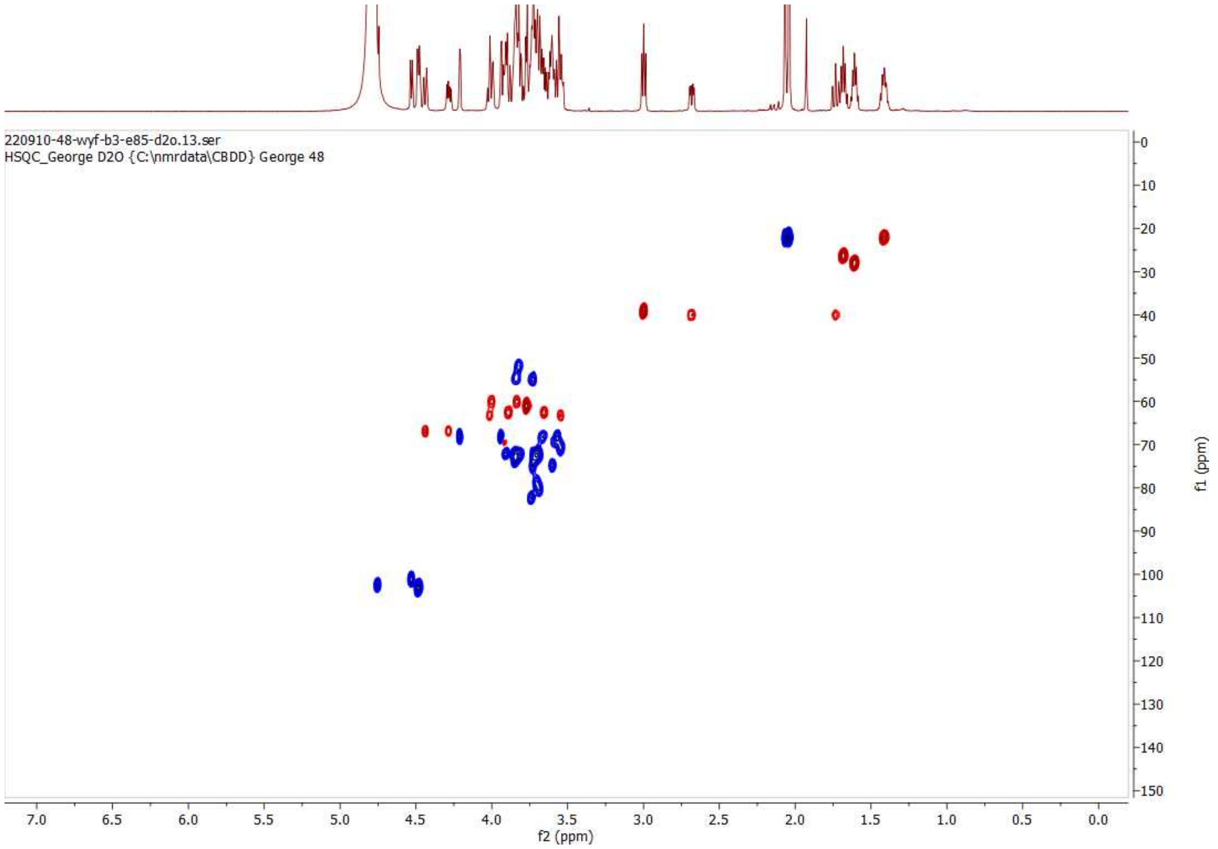

HSQC of 35; 600 MHz/150 MHz, $D_2O$

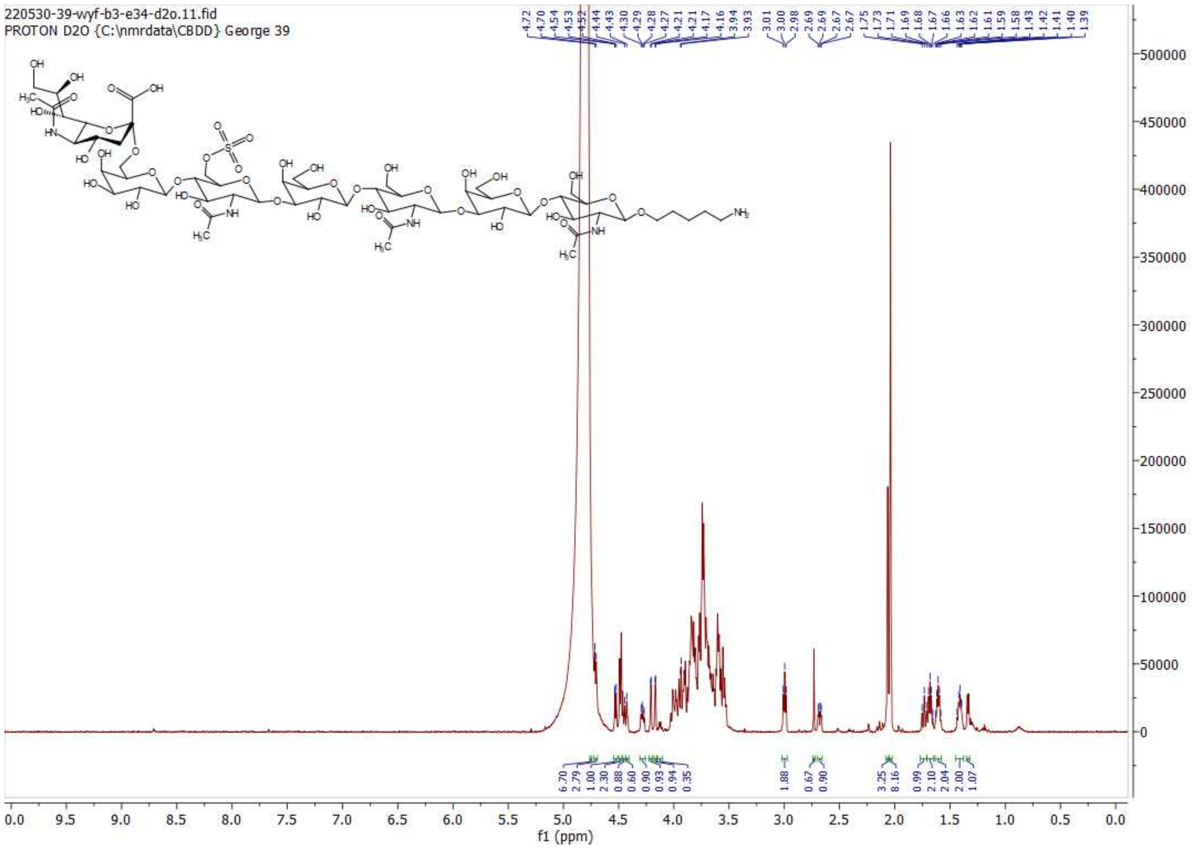

$^1$H NMR of 36; 600MHz; $D_2O$

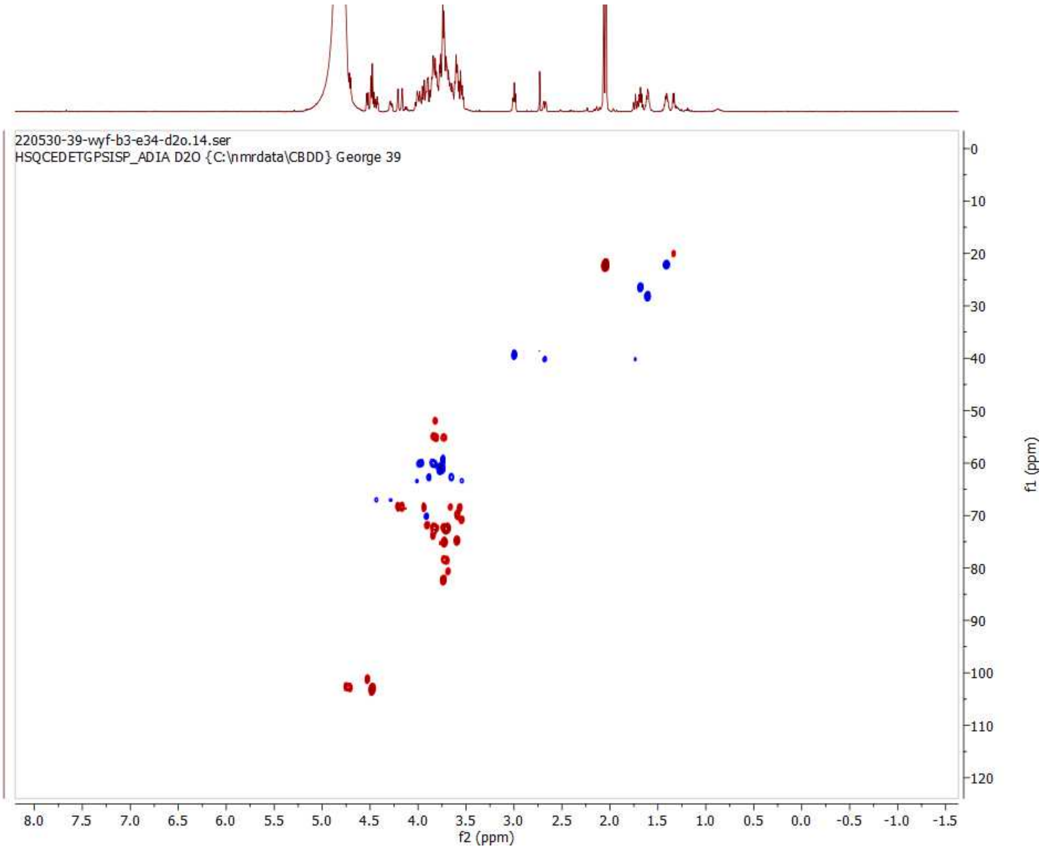

HSQC of 36; 600 MHz/150 MHz, $D_2O$

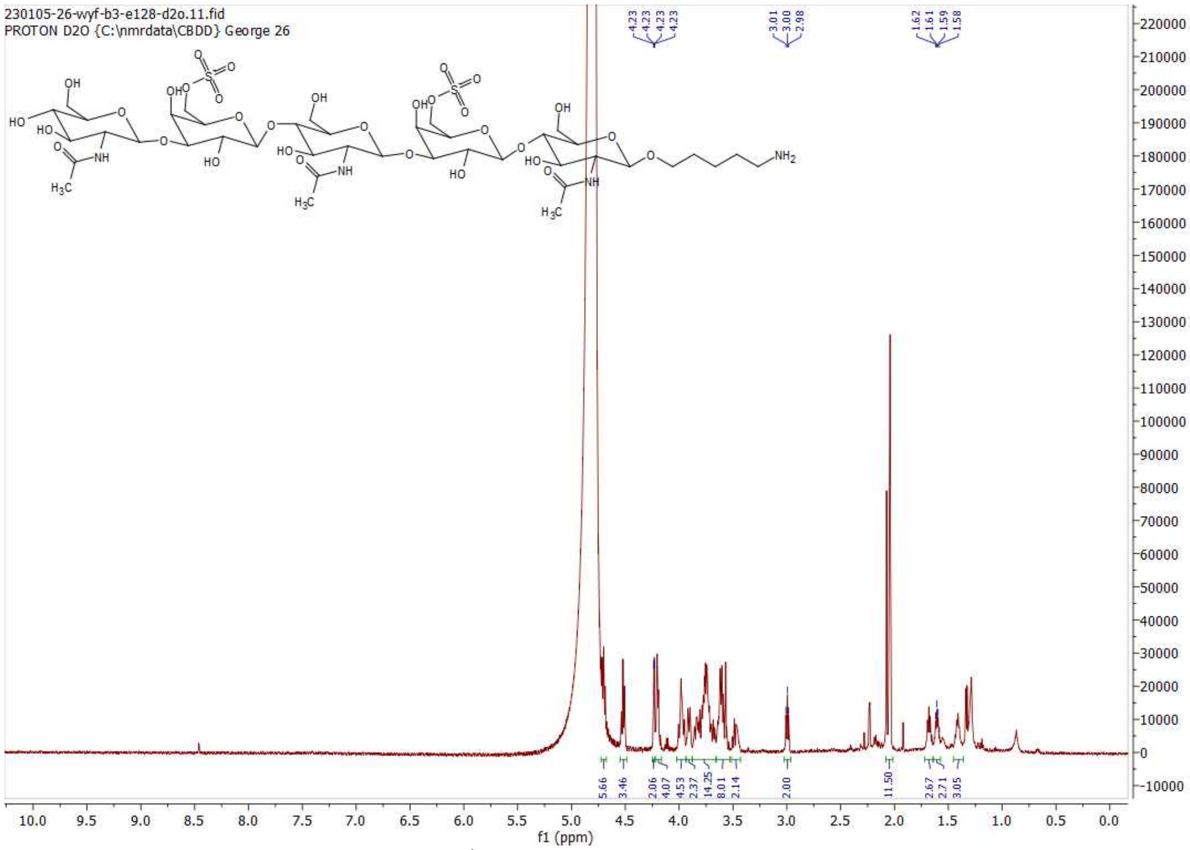

**[1]H NMR of 37; 600MHz; $D_2O$**

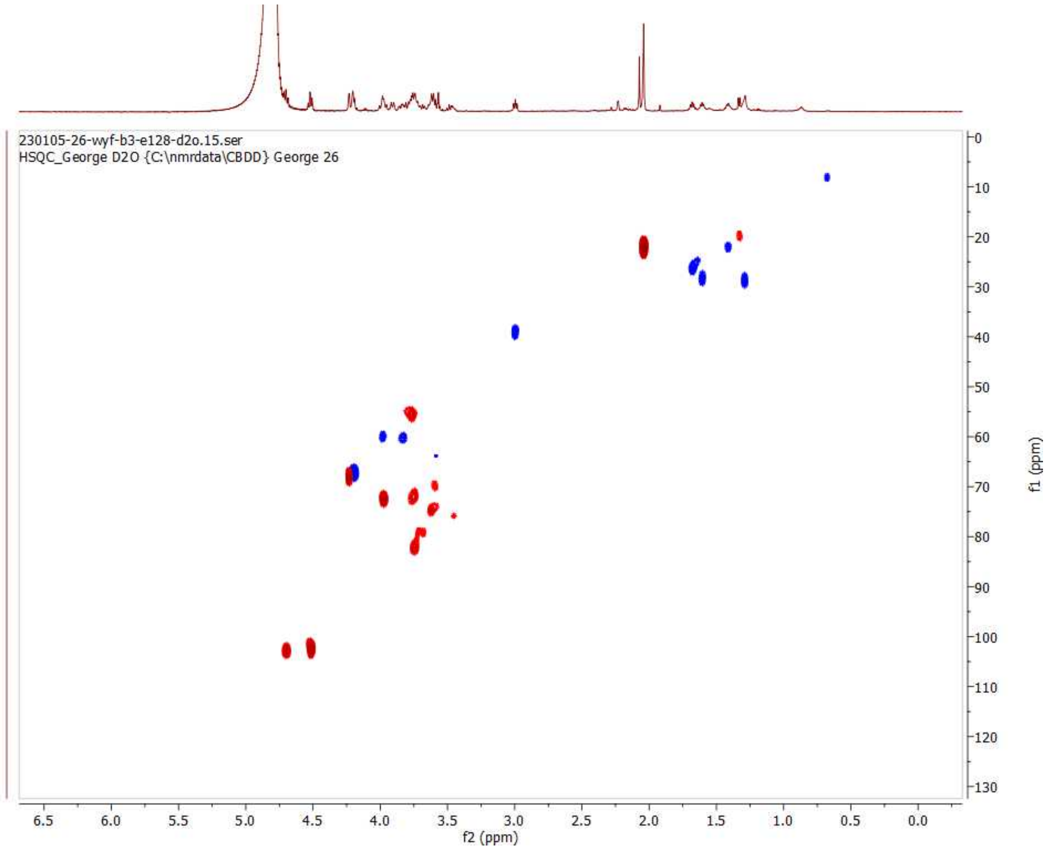

**HSQC of 37; 600 MHz/150 MHz, $D_2O$**

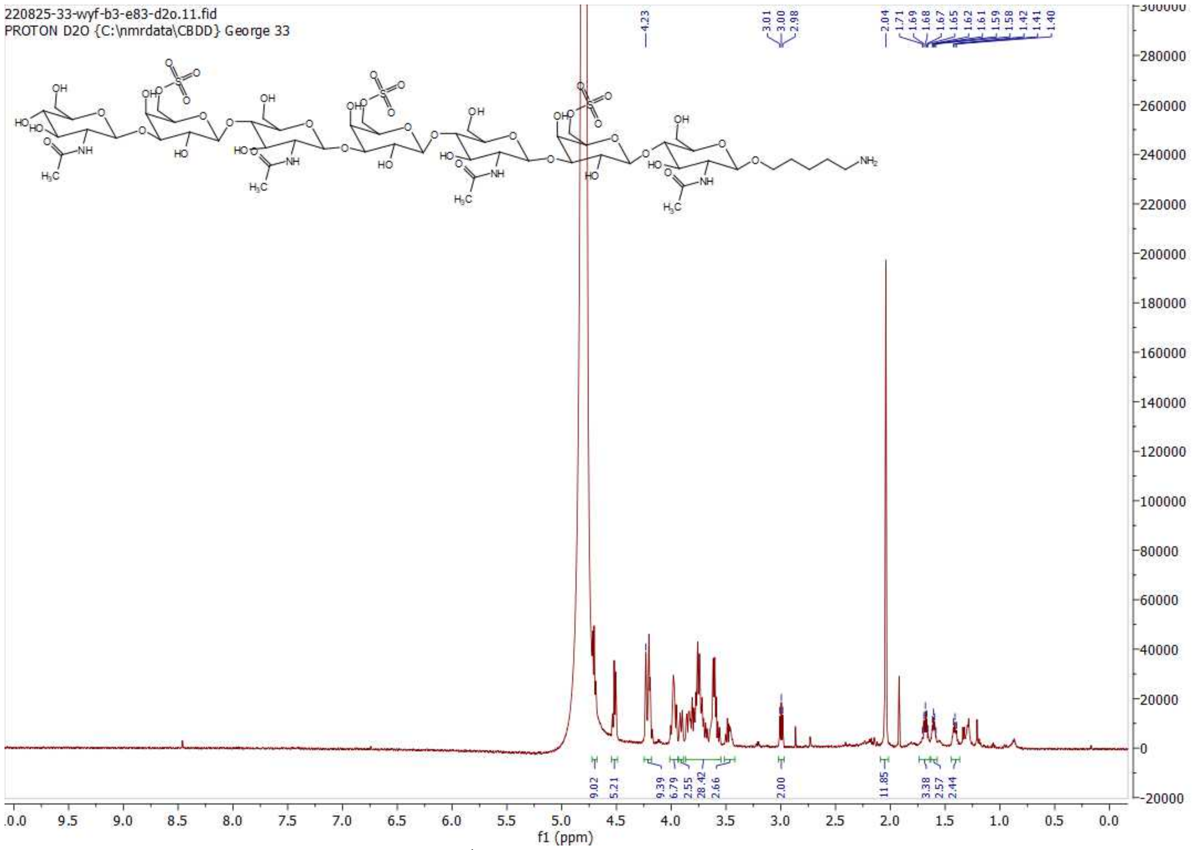

[1]H NMR of 38; 600MHz; $D_2O$

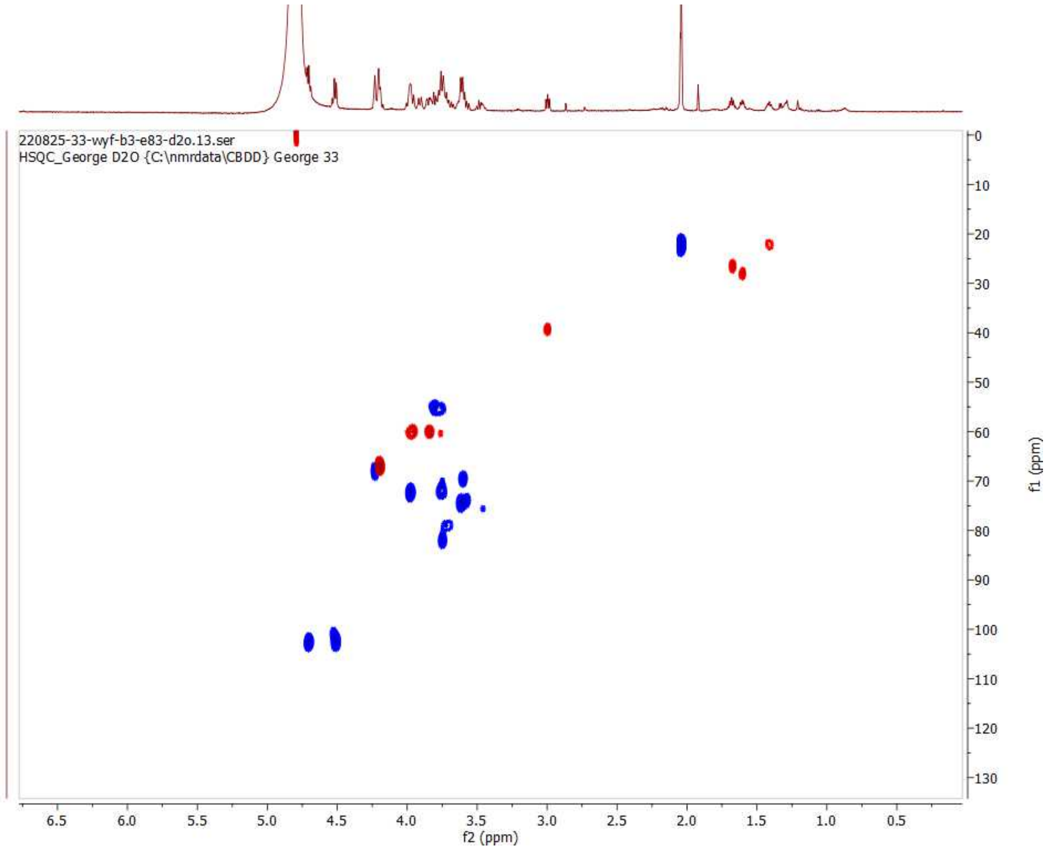

HSQC of 38; 600 MHz/150 MHz, $D_2O$

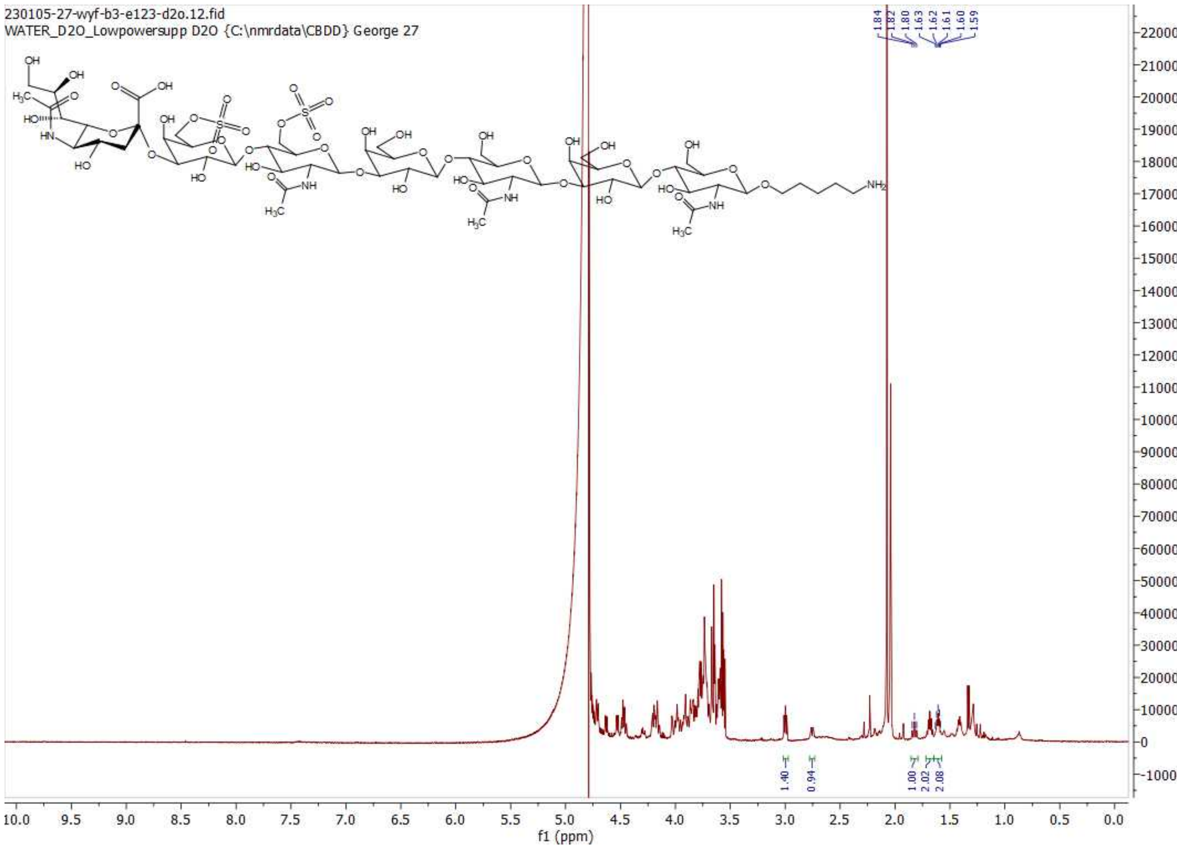

$^{1}$H NMR of 39; 600MHz; $D_2O$

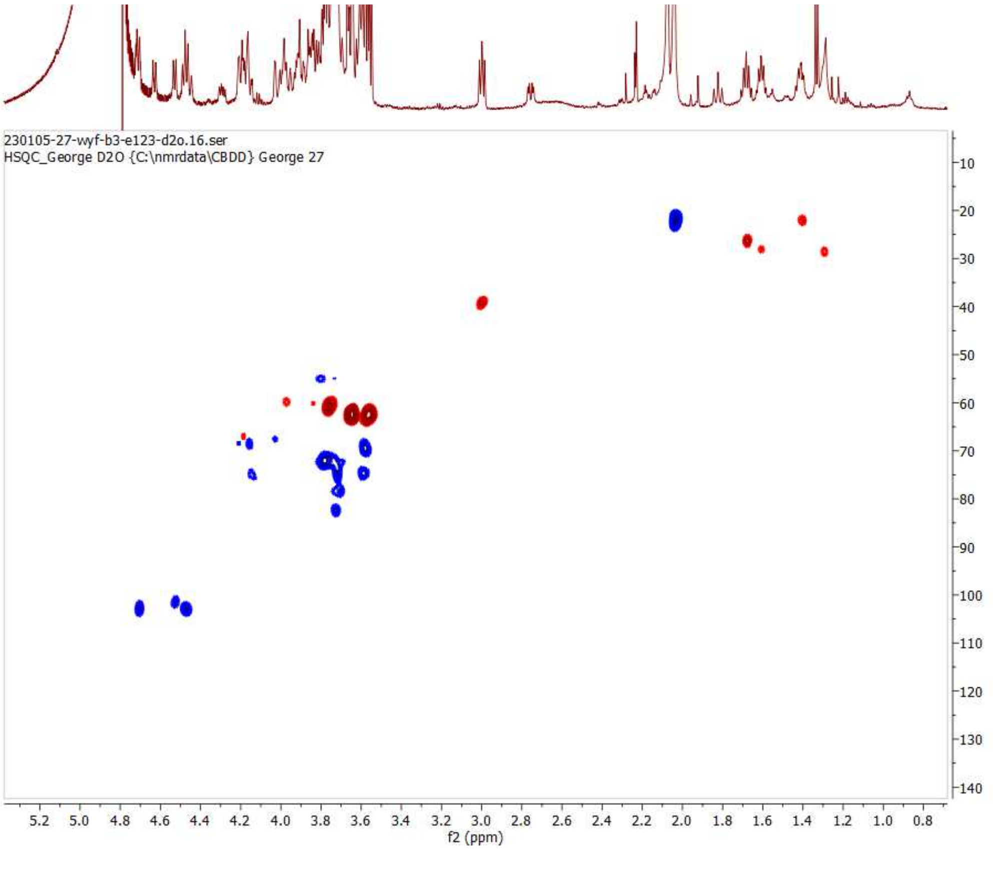

HSQC of 39; 600 MHz/150 MHz, $D_2O$
